# Core–sheath coupling controls flagellar curvature and motility in *Leptospira*

**DOI:** 10.64898/2026.02.02.703089

**Authors:** F San Martin, MR Brady, L Fule, L Nouchikian, A Rodriguez, S Mondino, N Larrieux, EA Wunder, AI Ko, M Rey, J Chamot-Rooke, R Duran, F Trajtenberg, M Picardeau, CV Sindelar, A Buschiazzo

## Abstract

Spirochete pathogens are among the most invasive bacteria known, causing syphilis, Lyme disease, and leptospirosis. Their tissue penetration depends on periplasmic flagellar filaments that, unlike other bacterial flagella, are encased in a spirochete-specific multi-protein sheath and deform the cell body into motile waves. How these filaments achieve the mechanical properties needed for invasive motility has remained unclear. Here we determine complete atomic structures of the *Leptospira* endoflagellar filament, revealing an elaborate sheath of 9 to 12 distinct asymmetrically arranged proteins. We show that the flagellin variant forming the filament core determines sheath composition, producing curvatures ranging from ∼3.5 µm^-1^ to ∼5 µm^-1^. The lower-curvature architecture, employed by pathogenic *Leptospira interrogans*, proves essential for motility in viscous environments and during infection. Thus, *Leptospira* achieves environment-specific motility through modular core–sheath coupling, linking atomic-scale structural plasticity to large-scale changes in swimming behaviour. Conservation of key sheath components suggests this mechanism may extend across spirochetes.

## Introduction

Spirochete bacteria encompass some of the most invasive bacterial pathogens known [1], including the causative agents of leptospirosis (*Leptospira* spp.), syphilis (*Treponema pallidum*), and Lyme disease (*Borrelia burgdorferi*). Their ability to rapidly disseminate through connective tissue, blood, and organs, including crossing biological barriers like the blood-brain barrier, relies on a distinctive swimming mechanism powered by periplasmic flagella (endoflagella). Unlike conventional bacterial flagella that extend freely into the aqueous medium [2], spirochete endoflagellar filaments are confined within the periplasm, where they transmit motor-generated torque along micron-scale lengths to deform the spiral-shaped cell body [3, 4]. Spirochete flagellar filaments precisely tune curvature and stiffness to enable motility across environments spanning orders of magnitude in viscosity, from fluids to host tissues [3, 5].

Spirochete pathogens cause serious zoonoses, vector-borne diseases, and sexually transmitted infections that affect millions worldwide each year. Among spirochetes, pathogenic *Leptospira* species cause >1 million cases of leptospirosis and nearly 60,000 deaths annually [6]. This globally important zoonosis is usually transmitted through contact with water contaminated with urine from animal reservoirs [7]. The disease ranges from mild self-limiting illness to severe manifestations including hemorrhage, jaundice, and renal failure (Weil’s disease). Beyond its medical relevance, *Leptospira* offers a powerful system for dissecting structure-function relationships in spirochete motility [8]: it possesses a single flagellum at each cell pole, providing structural simplicity compared to other spirochetes, while the saprophytic model *L. biflexa* enables rapid culture, high biomass yields, and tractable genetic manipulation. This combination enables detailed structural studies and direct comparison between pathogenic and non-pathogenic species.

Understanding flagellar structure is critical because spirochete motility is a key determinant of virulence. *Leptospira* mutants lacking functional flagella show severely attenuated infection in animal models [9, 10], and spirochetal motility is linked to unique adaptations in their flagellar motor, hook and filament [11, 12]. Unlike exoflagellated bacteria, whose filaments consist of flagellin alone [3], spirochetal filaments are covered by a proteinaceous sheath. Until recently, the only known sheath component was FlaA, conserved across the phylum [3]; however, the discovery of two additional, *Leptospira*-specific sheath components (FcpA and FcpB) revealed that sheath composition was more complex [10, 13]. Structural studies of *Leptospira* filaments by cryo-electron tomography identified an asymmetric, uneven distribution of sheath proteins covering the flagellin core in native supercoiled configurations [14]. Subsequently, high resolution structures of mutant *Leptospira* filaments uncovered the role that individual sheath components (FlaA2, FcpA and FcpB) play to induce filament supercoiling, demonstrating that an asymmetric protein arrangement is essential for flagellar function [15]. However, how the complete sheath is organized in wild-type filaments, and how it cooperates with the flagellin core to control filament curvature and motility, have remained unknown.

Here we present complete atomic structures of endoflagellar filaments from *L. biflexa* and from the pathogen *L. interrogans*, integrating cryo-electron microscopy (cryo-EM), quantitative and crosslinking mass spectrometry (MS), and X-ray crystallography. We show that the sheath architecture is more elaborate than previously anticipated, with 9 to 12 distinct proteins asymmetrically enveloping the flagellin core. Furthermore, comparison of the wild-type pathogen cryo-EM structure with that of a *flaB1^-^* knockout mutant (deleting the main core flagellin) reveals that swapping core isoforms (from FlaB1 to FlaB4) reorganises the sheath and substantially increases the filament curvature, abrogating virulence and tissue penetration. These findings reveal how modular core-sheath coupling links atomic-scale structural plasticity to large-scale flagellar mechanics and invasive motility.

## Results

### Leptospira biflexa filaments display a highly elaborate sheath organization

Cryo-EM structures of wild-type (*wt*) *Leptospira biflexa* were determined at 3.5 Å resolution [16]. Density modification [17] improved map details achieving a final resolution of 3.2 Å. (Fig. 1a,b). The atomic structure was solved using an integrative strategy (Extended Data Fig. 1) that combined cryo-EM single particle analysis (Supplementary Table 1; Extended Data Fig. 2), crosslinking and quantitative MS (Table 1; Fig. 1c; Extended Data Fig. 3; Supplementary Tables 2 & 3), X-ray crystallographic structures of FcpA and FcpB [14], and AlphaFold3 predictions. This process was iterated to obtain the final structure: comparative proteomics identified candidate proteins including unannotated gene products, crosslinking data prioritized these by proximity [18], and cryo-EM density fitting provided validation. Each confirmed component enabled re-interrogation of crosslinking interaction networks, progressively expanding the catalogue of filament constituents.

**Fig. 1.**
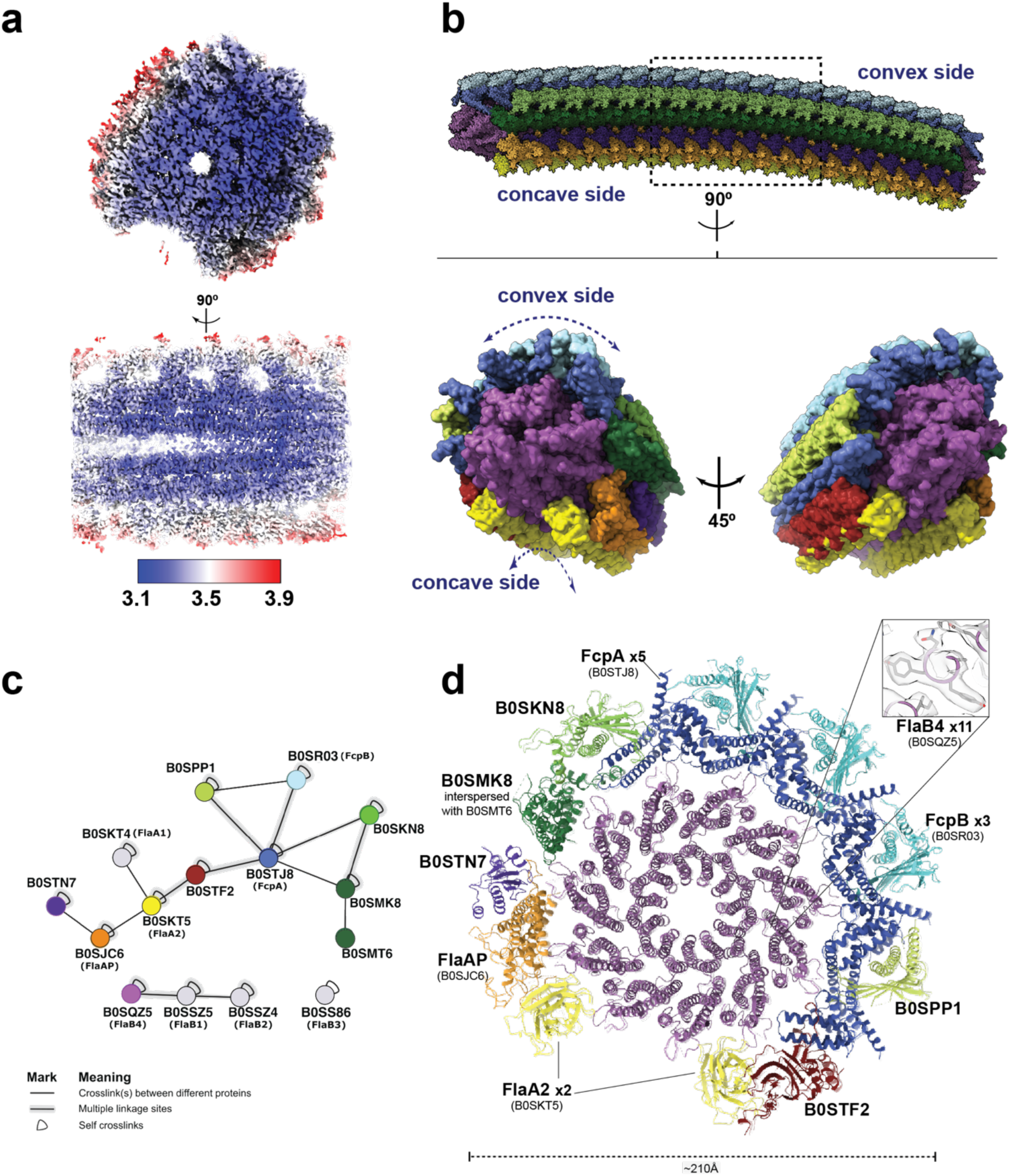
Integrative 3D structure determination of *Leptospira biflexa* flagellar filaments. **(a)** Cryo-EM map of a native *L. biflexa* filament coloured by local resolution (scale bar: 3.1-3.9 Å), viewed from one end (top) and from the side (bottom), contoured at 0.05 threshold. **(b)** Cryo-EM reconstruction shown as a molecular surface, coloured by protein identity. Top: side view with convex and concave sides indicated; dashed box marks the size of the reconstructed volume (∼7 repeat layers), extended at both ends by superposition of terminal layers to visualize overall curvature more clearly. Bottom: end-on views of the refined volume, rotated 45° relative to each other to reveal lateral sheath organisation surrounding the FlaB4 core. **(c)** Crosslinking mass spectrometry interaction network. Nodes represent identified filament proteins, coloured as in panels (b,d) and labelled with UniProt IDs (gene names in parentheses for annotated proteins). Edges indicate inter-protein crosslinks; thick edges denote multiple linkage sites; self-crosslinks shown as loops. Core flagellins (bottom) lack inter-protein crosslinks with sheath components, consistent with glycan-mediated core-sheath interactions. An interactive version of this network is available at https://www.xiview.org/network.php?upload=42923-71803-52415-45391-89719, allowing exploration of individual crosslinks and protein properties. **(d)** Refined atomic model shown as cartoon, viewed as a transverse cross-section representing one helical layer (∼52 Å). Proteins labelled with UniProt IDs and copy numbers per layer. Inset: FlaB4 core cryo-EM density (semi-transparent surface) with fitted atomic model (sticks)

**Table 1.**
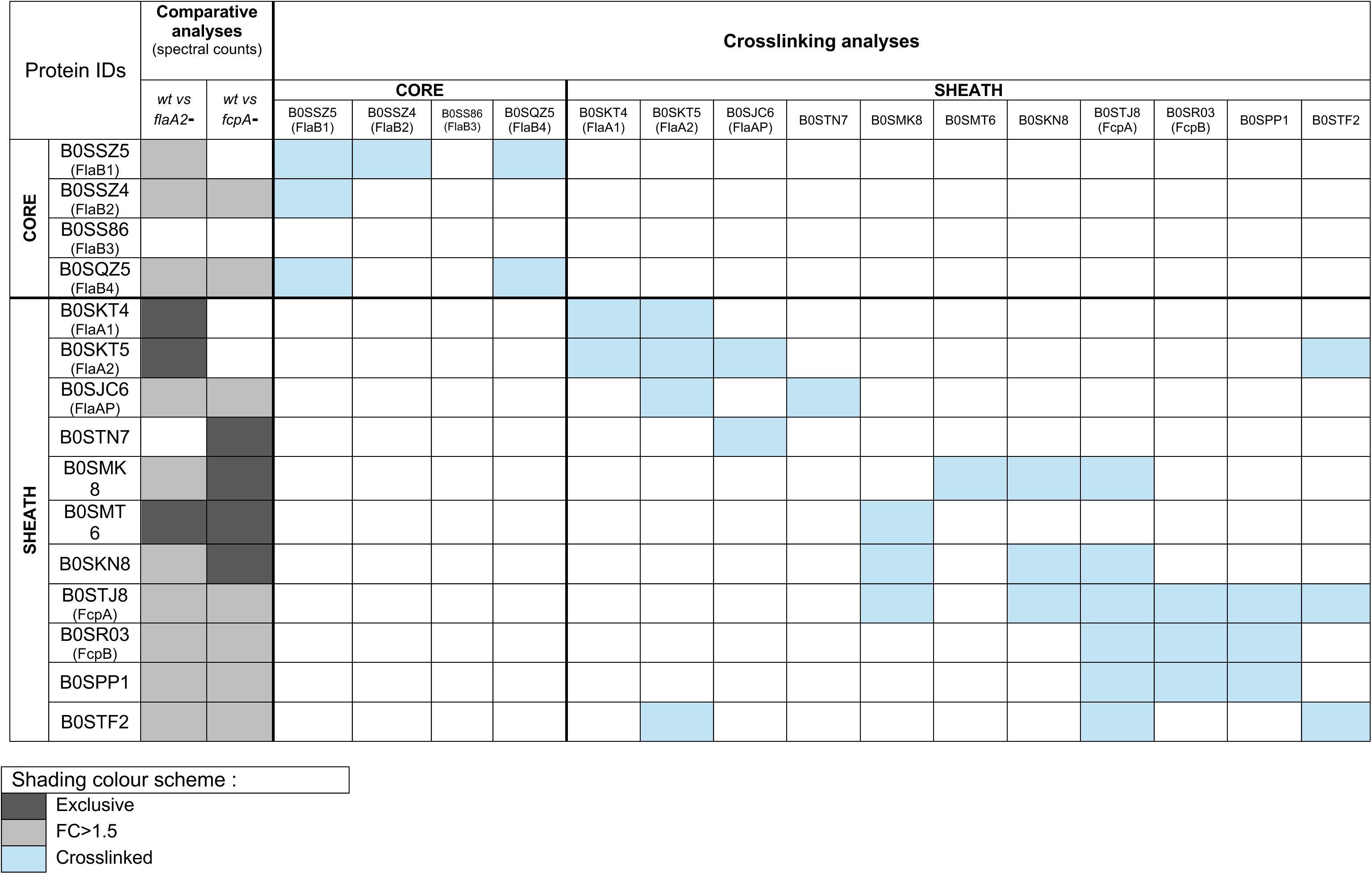
Proteins identified by comparative proteomics and crosslinking mass spectrometry analyses of Leptospira biflexa flagellar filaments. *\*when within the same protein species, crosslinks between different subunits in the filament are shown (inter-crosslinks)* See Supplementary Tables 2 & 3 for full details; and https://www.xiview.org/network.php?upload=42923-71803-52415-45391-89719 for interactive analysis

The refined *L. biflexa* filament model comprises a core of flagellin (FlaB) protomers coated with 11 distinct sheath proteins and one additional unidentified component, distributed with a radially asymmetric organization and a ∼52 Å periodicity along the filament (Fig. 1b,d). This architecture starkly contrasts with previously reported, core-only bacterial flagellar filament structures, and more than doubles the known number of sheath components in spirochetes, including *Leptospira*.

The core follows the canonical bacterial flagellar architecture, with FlaB protomers polymerized helically (twist 65.4°; rise 4.73 Å) into 11 protofilaments enclosing a ∼20 Å central channel. Inter-protomer contacts are established along three principal subsidiary helices: the 11-start (corresponding to the longitudinal protofilaments), the right-handed 6-start, and the left-handed −5-start [19]. Although *Leptospira* expresses four FlaB isoforms [20], *L. biflexa wt* filaments display FlaB4 as the major core component, with distinctive side chains and glycosylation sites visible in the cryo-EM density.

The sheath displays a heterogeneous organization, with distinct proteins defining different surfaces of the assembly (Fig. 1d; Extended Data Fig. 4; Supplementary Table 4), overall organized in two concentric tiers around the core. The inner tier establishes direct contacts with FlaB, primarily through glycans [15]: FlaA2 moieties coat the concave surface, each associated with distinct partner proteins, FlaAP on one side and B0STF2 on the other; FcpA protomers line the convex surface; and a mixed protofilament of alternating B0SMK8 and B0SMT6 (structurally similar to FlaAP and showing inter-crosslinks between them by MS, see Fig 1c) completes the core-contacting shell. An outer tier of proteins caps this first layer without direct core contact: FcpB moieties sit atop FcpA on the convex side, as do B0SPP1 and B0SKN8 (which share an FcpB-like fold); B0STN7 caps the junction between FlaAP and B0SMK8; and density for one additional outer-tier component, capping FlaA2 and FlaAP, could not be interpreted. Overall, the sheath proteins are organised with a helical pseudosymmetry matching the core’s, with complementary ridges and troughs mediating core-sheath contacts. Notably, a groove remains at the concave surface between the two FlaA2 protofilaments, as reported in earlier tomography studies [14]. Although MS data unambiguously detected FlaA1 in purified samples, including crosslinks to FlaA2 (Fig. 1c; Table 1), and the groove could accommodate two FlaA1 units per sheath layer, no cryo-EM density could be attributed to this protein, prompting further investigation of this poorly characterized sheath component.

### FlaA1 structure reveals a new Carbohydrate Binding Module family and calcium-dependent stability

While FlaA2 structure has been determined by cryo-EM [15], FlaA1 remained uncharacterized. The two paralogues share only ∼25% sequence identity, and FlaA1 is ∼60 residues larger. The puzzling loss of FlaA1 from *L. biflexa* filaments motivated its X-ray crystallographic analysis. Recombinant FlaA1 required iodoacetamide (IAA) and calcium for solubility; after extensive crystallization trials, the structure from *L. borgpetersenii* FlaA1 (UniProt Q04UP1) was solved at 1.85 Å resolution (Fig. 2a,b; Supplementary Table 5). FlaA1 exhibits a bent jelly-roll-like β-sandwich fold creating a deep concave groove opposite a two-faceted convex side (Fig. 2c). The concave face comprises a 5-stranded antiparallel β-sheet, while the convex side has two partly discontinuous antiparallel β-sheets (four and six strands). Fold and topology identify FlaAs as Carbohydrate Binding Modules (CBM) [21], though not belonging to any known CBM family [22]. Crucially, a calcium cation near the domain’s triangular vertex (Fig. 2c) is octahedrally coordinated by four protein oxygens and two water molecules. The cation stabilizes two β-strands on each convex facet explaining its role in protein stability. Re-examination of the *L. biflexa* cryo-EM map revealed density consistent with calcium at the equivalent position in FlaA2 (Fig. 2d), confirming that calcium-dependent fold stabilization is conserved across the FlaA family.

**Fig. 2.**
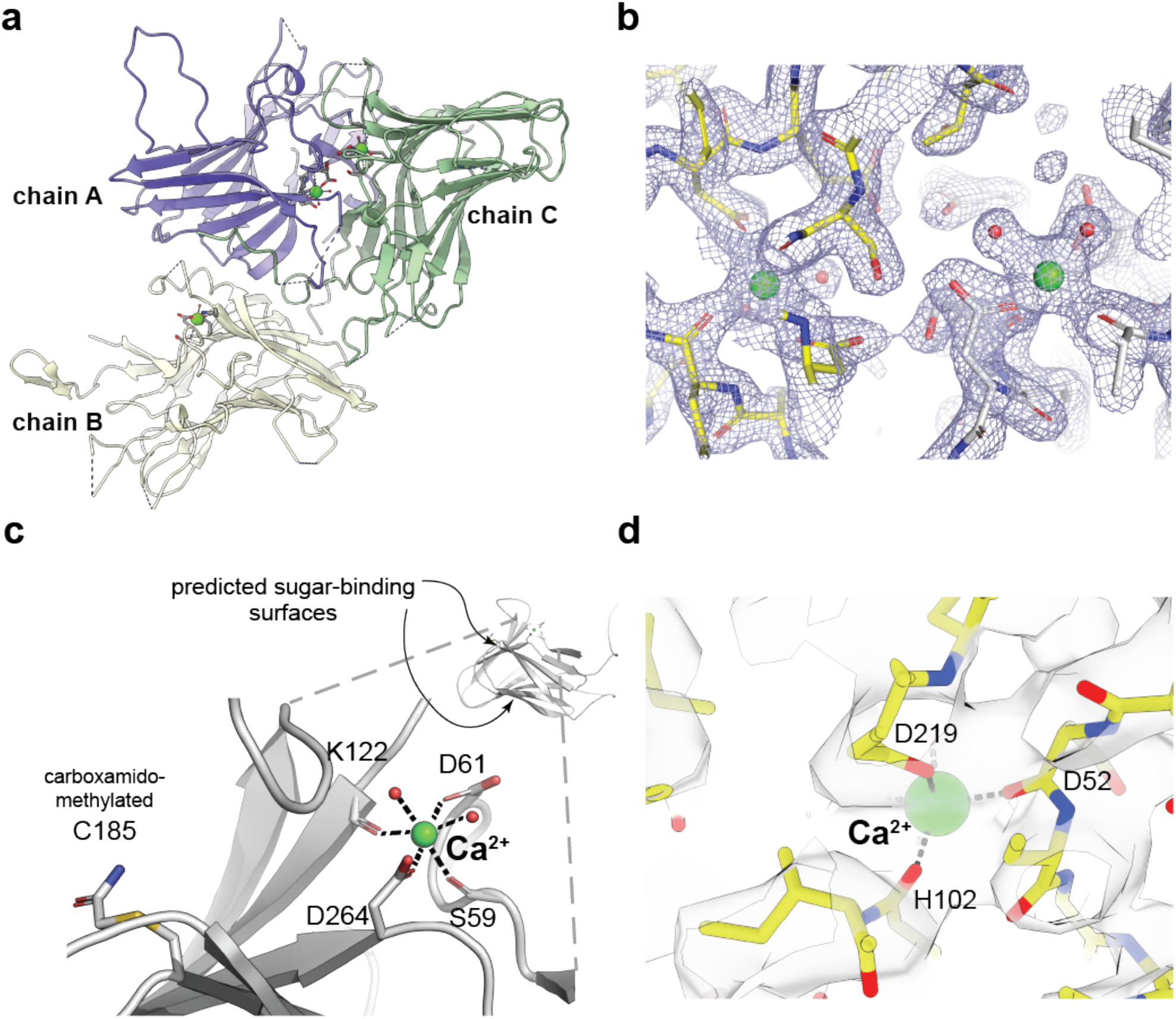
FlaA1 structure reveals a calcium-dependent carbohydrate-binding module. **(a)** Crystal structure of *L. borgpetersenii* FlaA1 at 1.85 Å resolution. The asymmetric unit contains three monomers (chains A, B, C) shown as cartoons in different colours. Ca^2+^ ions are shown as green spheres. **(b)** Representative σA-weighted 2mFo-DFc electron density map contoured at 1.2σ, showing well-resolved Ca^2+^ coordination and surrounding residues. **(c)** Close-up of the Ca^2+^-binding site showing octahedral coordination by Asp61 and Asp264 (side chain oxygens), Ser59 and Lys122 (backbone carbonyls), and two water molecules. The carboxamidomethylated Cys185 marks a predicted sugar-binding surface. Dashed lines connect to a zoom-out view showing the position of the Ca^2+^ site within the overall bent jelly-roll fold, where the locations of two putative carbohydrate-binding grooves are marked. **(d)** Retrospective analysis of *L. biflexa* FlaA2 cryo-EM density at the structurally equivalent Ca^2+^ binding site. Density consistent with bound calcium is observed, with coordinating residues Asp52, His102, and Asp219 labelled.

Superposition with characterized CBM structures suggests potential sugar-binding sites in FlaA1: one in its concave groove (Fig. 2c), another on a convex facet harbouring a groove with Cys185 on its floor. IAA reacted with this solvent-exposed Cys185 during crystallization, producing its stable carboxamidomethylated derivative. Within assembled filaments, FlaA2 concave crevices bind sugar moieties decorating FlaB core protomers [15] (confirmed here in native filaments, see Extended Data Fig. 6c,d), validating the functional relevance of CBM-type recognition in core-sheath interactions. Whether FlaA1 is incorporated within native filaments remained to be determined.

### *L. interrogans* filaments exhibit a closed sheath and reveal architecture plasticity

The apparent loss of FlaA1 during *L. biflexa* filament preparations for cryo-EM imaging, together with proteomic data indicating distinct FlaB isoform profiles across species [23], prompted us to determine the native structure of flagellar filaments from pathogenic *L. interrogans*. Cryo-EM reconstruction yielded a 4.4 Å resolution map, with further density modification improving resolution to 4.2 Å (Fig. 3a; Supplementary Table 1; Extended Data Fig. 5). The atomic model was built by fitting AlphaFold-predicted orthologues into density (Fig. 3b), with *L. interrogans* proteome data [15] and our *L. biflexa* comparative proteomics analysis guiding the assignments.

**Fig. 3.**
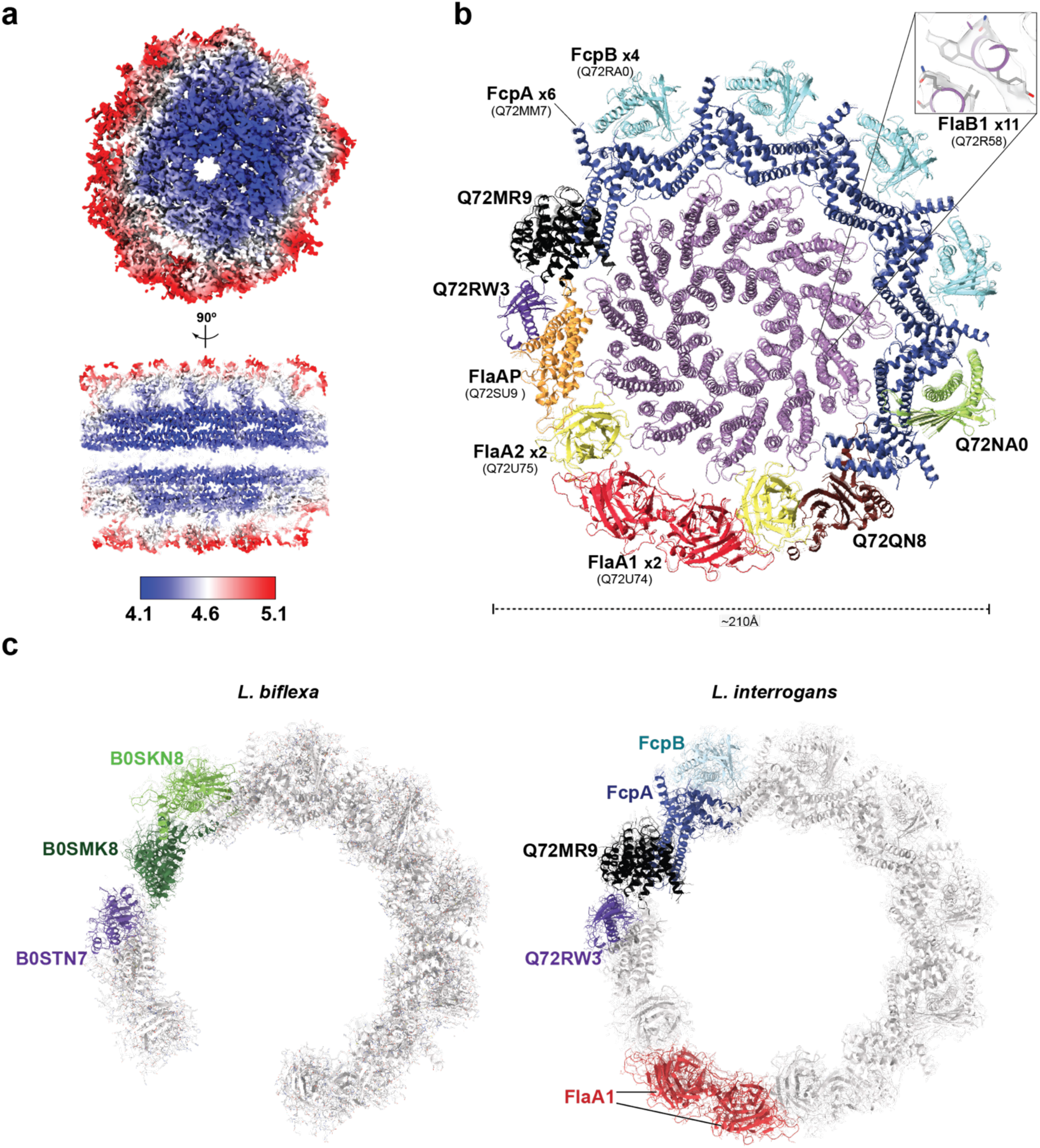
Native *Leptospira interrogans* filaments exhibit a fully closed sheath cylinder. **(a)** Cryo-EM map coloured by local resolution (scale bar: 4.1–5.1 Å), viewed from one end (top) and from the side (bottom), contoured at 0.04 threshold. **(b)** Refined atomic model shown as cartoon in transverse cross-section. Orthologous proteins are coloured as in Fig. 1. Novel *L. interrogans*-specific components are labelled distinctly. Inset: close-up of FlaB1 core density with fitted model. **(c)** Side-by-side comparison of lateral sheath regions in *L. biflexa* (left) and *L. interrogans* (right), highlighting species-specific differences. *L. biflexa* features B0SKN8, B0SMK8, and B0STN7; *L. interrogans* features an additional unit of FcpA and FcpB, the HEAT-repeat protein Q72MR9, Q72RW3, and FlaA1 occupying the lateral groove.

Based on differential cryo-EM features at large side chains and diagnostic glycosylation sites, the *L. interrogans* core is predominantly FlaB1, contrasting with FlaB4 in the saprophyte. The groove on the concave face of the sheath (open in *L. biflexa*) is occupied in the pathogen by two FlaA1 protomers per sheath layer, completing the cylinder (Fig. 3b). The crystallographic monomer fitted unambiguously into this density (Extended Data Fig. 6a), resulting in a dimeric arrangement consistent with FlaA1-FlaA1 and FlaA1-FlaA2 crosslinks (Table 1). The 2-fold axis relating both monomers is perpendicular to the filament, such that the two FlaA1 protofilaments run antiparallel, each contacting distinct surfaces on their neighbouring FlaA2 partners (Extended Data Fig. 6a). Interactions with FlaA2 protofilaments open the dimer angle relative to the crystal structure (Extended Data Fig. 6b). The larger FlaA1 sugar-binding groove interfaces with FlaA2 rather than the core, while the second putative sugar-binding crevice is exposed on the filament surface, forming a continuous outward-facing groove with the dyad-related FlaA1 neighbour (Extended Data Fig. 6c). FlaA2 concave crevices bind FlaB *O*-glycans (Extended Data Fig. 6d) as in *L. biflexa* [3]. Intriguingly, the completed cylinder forms a flattened ∼20 Å lateral channel bounded by FlaA1 (Extended Data Fig. 6e).

The *L. interrogans* sheath comprises 9 distinct proteins (Fig. 3b,c). The convex side possesses six FcpA and four FcpB units per repeat, compared to five FcpA and three FcpB in *L. biflexa*. However, the *L. biflexa*-specific protein B0SKN8 exhibits a FcpB-like fold, making the convex region compositionally similar except for the extra FcpA moiety in *L. interrogans*. The two species show the greatest architectural divergence in one sector of the sheath (Fig. 3c). In *L. interrogans*, an extra FcpA unit bridges the convex outer tier FcpB array to Q72MR9, a HEAT repeat protein [24] that directly contacts the core (Extended Data Fig. 4d). The HEAT protein then further contacts FlaAP on the concave side, capped by the outer tier Q72RW3. In *L. biflexa*, the position of the HEAT protein is instead occupied by the mixed B0SMK8/B0SMT6 protofilament (both FlaAP paralogues). Although *L. interrogans* expresses orthologues of these FlaAP-like proteins [23], neither was observed in its filament structure. This sector of the sheath thus represents a zone of compositional plasticity, where proteins are being differentially incorporated in each *Leptospira* species. The observed species-specific correlation between sheath organization and core compositions prompted us to investigate whether these two architectural features are coupled.

### Core flagellin identity is coupled to sheath architecture, dictating filament curvature

Extensive *O*-glycosylation of FlaB protomers on their outer surfaces dominantly mediates the native core:sheath interaction network [15] in both species. To pinpoint whether the distinct FlaB isoforms that dominate their respective cores exert an influence on sheath organization and stability, we generated a *flaB1* deletion mutant (*flaB1^-^*) in *L. interrogans*. This mutant grew normally *in vitro* and swam indistinguishably from *L. interrogans wt* when observed by darkfield microscopy in liquid medium (Supplementary Video 1). However, motility assays across viscosity conditions revealed striking differences (Fig. 4a). In low-viscosity medium, *L. interrogans flaB1^-^*cells swam faster than *wt* (5.5 vs 3.9 µm/s), but this relationship reversed as viscosity increased: in 1% methylcellulose, mutant velocity dropped to 2.5 µm/s while *wt* increased to 5 µm/s. Consistently, the mutant showed severely impaired dispersion in semi-solid agar, dropping to 29% of *wt*, comparable to the non-motile *L. interrogans flaA2^-^* control (Fig. 4b, Extended Data Fig. 7a). This viscosity-dependent motility defect translated to impaired translocation across epithelial monolayers (Fig. 4c) and, critically, complete attenuation of virulence in the hamster model of acute infection (Extended Data Fig. 7b). Thus, while the *flaB1^-^* mutant assembles functional flagella, their mechanical properties are not optimized for the viscous environments encountered during tissue invasion.

**Fig. 4.**
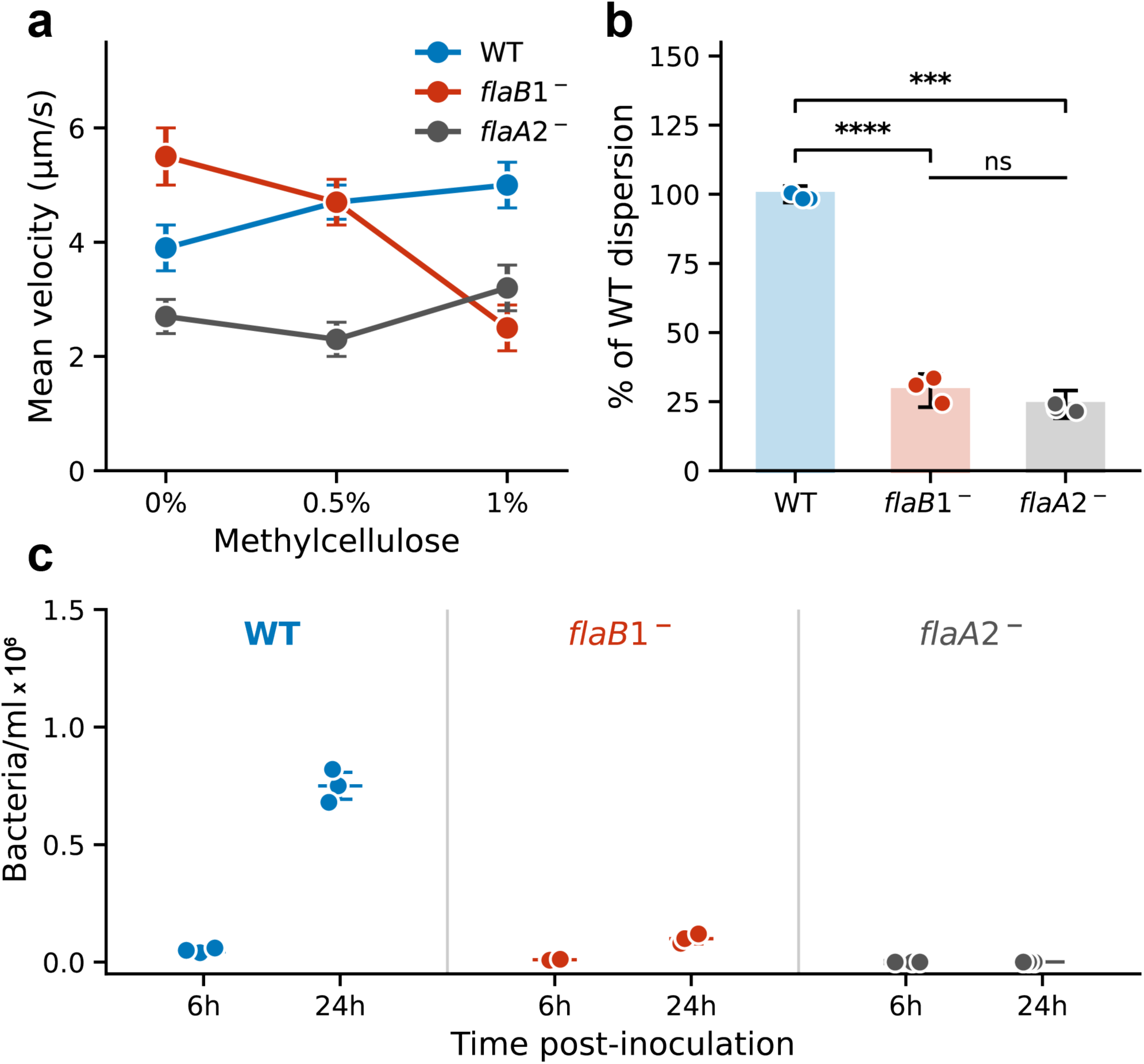
*L. interrogans flaB1^-^* mutant reveals viscosity-dependent motility defects and impaired tissue invasion. **(a)** Swimming velocity of *L. interrogans* wild-type (WT, blue), *flaB1^-^* mutant (pink), and non-motile *flaA2^-^* control (grey) measured in liquid EMJH medium supplemented with increasing concentrations of methylcellulose (MC: 0%, 0.5%, 1%). Data represent mean ± s.d. from ∼60 tracked bacteria per condition. **(b)** Dispersion in semi-solid agar (0.8%) after 10 days at 30°C, normalized to WT. Data represent mean ± s.d. from ≥3 independent experiments. **** p < 0.0001; *** p < 0.001 (one-way ANOVA with *post-hoc* test). **(c)** • Translocation across MDCK-1 epithelial monolayers in Transwell® assays. Bacteria in the lower chamber were quantified at 6 h and 24 h post-inoculation. The *flaB1^-^*mutant shows ∼8.7-fold reduction in translocation compared to WT at 24 h, with no detectable crossing at 6 h. Data represent mean ± 1 sd from 3 replicates.

The stark phenotypic differences prompted structural characterization of the *flaB1^-^* mutant flagellar filament to identify underlying architectural bases. Cryo-EM reconstruction of *flaB1^-^*filaments at 4.3 Å resolution (Supplementary Table 1; Extended Data Fig. 8) revealed that FlaB4 is indeed the dominant core component. Position 126 (serine) is glycosylated in all major FlaB isoforms. Position 137 provides a diagnostic distinction: FlaB4 carries threonine (O-glycosylated), whereas FlaB1 carries glutamine, which cannot be glycosylated (Extended Data Fig. 9a). The *flaB1^-^* map confirms FlaB4 identity through unambiguous glycan density at position 137 and differential densities at other diagnostic side chains (Extended Data Fig. 9b). The mutant thus provides an *L. interrogans* background with a FlaB4-dominated core.

Beyond its diagnostic value, position 137 has direct consequences for sheath selectivity. At the I–J protofilament interface, the Thr137 *O*-glycan on FlaB4 contacts and stabilizes B0SMK8 at the core–sheath boundary (Fig. 5a); this contact is conserved in *L. biflexa wt* and *L. interrogans flaB1^-^* (orthologue Q72Q50). In *wt L. interrogans*, Q137 cannot be glycosylated, abolishing this contact; the conserved Ser126 glycan then repositions to bridge across protofilaments I and J, eliminating the seam between them (Fig. 5a). A single glycosylation site thus controls both sheath protein selectivity and seam distribution.

**Fig. 5.**
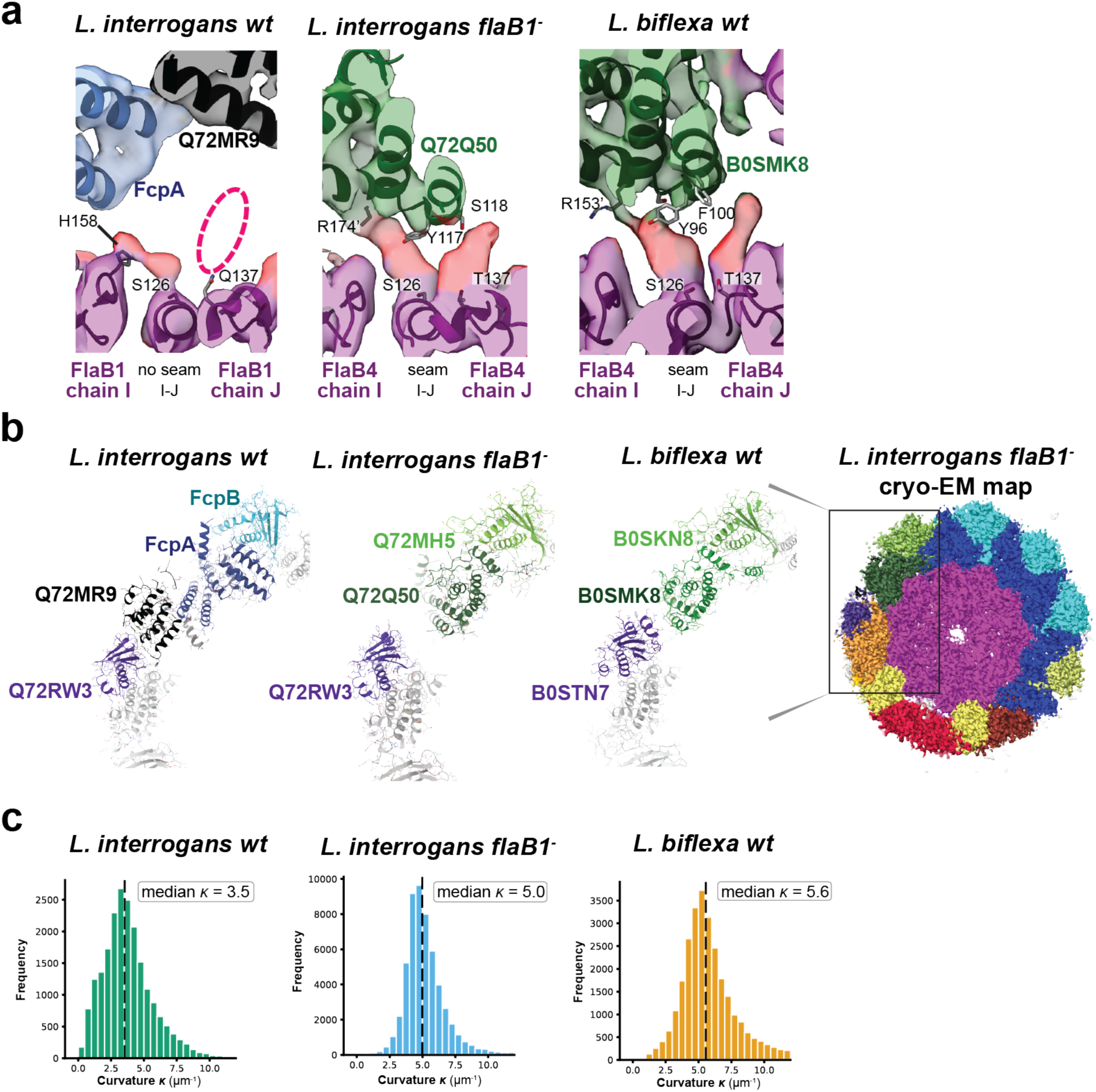
Core-driven regulation of *Leptospira* filament curvature. **(a)** Cryo-EM density of *L. interrogans wt* (left), *L. interrogans flaB1^-^* (centre), and *L. biflexa wt* (right), coloured according to fitted models. Glycan densities are coloured red. The view is equivalent across the strains, focused on the I–J core protofilament interface. Gln137 on FlaB1 (*L. interrogans wt*) cannot be glycosylated (dashed circle), creating a looser sheath interaction and no seam; Thr137 on FlaB4 (*flaB1^-^* and *L. biflexa wt*) is *O*-glycosylated, directly stabilizing B0SMK8 orthologues and introducing a seam within the core. **(b)** Cryo-EM map of *L. interrogans flaB1-* (solid surface, right), coloured according to best-fitting atomic model chains (Extended Data Fig. 8c). The boxed area highlights the variable sheath region, compared across all three strains (left). The *flaB1-*sheath architecture resembles *L. biflexa* rather than parental *L. interrogans*. **(c)** Distribution of filament curvature (κ) for the three strains. Histograms show individual curvature measurements along filament trajectories; dashed lines indicate median values. The curvature of *flaB1-* closely matches *L. biflexa wt* rather than parental *L. interrogans wt*, correlating with sheath architectures shown above.

This selectivity extends across the entire sheath: the *flaB1^-^* organization resembles *L. biflexa* rather than parental *L. interrogans* (Fig. 5b). The extra FcpA and HEAT repeat protein (Q72MR9) characteristic of *L. interrogans wt* are absent in the *flaB1^-^* density, replaced by *L. biflexa*-type architecture (Extended Data Fig. 9c). The refined *flaB1^-^* atomic model fits well into both the *flaB1^-^* and *L. biflexa wt* maps, but leaves unoccupied density in *L. interrogans wt* where species-specific proteins reside (Fig. 5b; Extended Data Fig. 9c).

These architectural similarities extend to mechanical properties. Quantitative curvature analyses [25] revealed substantial differences among the three strains (Fig. 5c). *L. interrogans wt* filaments exhibited the lowest curvature (median κ = 3.5 μm^-1^; n = 20,977), while *L. biflexa wt* displayed significantly higher values (median κ = 5.6 μm^-1^; n = 28,853; Δκ = 2.1 μm^-1^). Critically, the *flaB1^-^* mutant closely matched *L. biflexa* (median κ = 5.0 μm⁻¹; n = 55,579) rather than parental *L. interrogans*. This demonstrates that FlaB isoform identity, coupled with corresponding sheath architecture, primarily dictates filament curvature. The residual difference between FlaB4-containing assemblies (Δκ = 0.6 μm^-1^) likely reflects species-specific variations in minor core proteins and/or post-translational modifications.

What structural features underlie this curvature modulation? The strict helical symmetry of FlaB cores is subtly broken at certain protofilament interfaces creating seams, which consist of ∼2.3 Å translational shifts (Extended Data Fig. 10a). *L. biflexa* exhibits four seams, whereas *L. interrogans* displays only two (Extended Data Fig. 10b,c). Beyond the shared seams at interfaces K-A and B-C (each contacting a FlaA2 moiety on the sheath concave side), *L. biflexa* has two additional seams: at A-B, directly beneath the position where FlaA1 is absent, and at I-J (Fig. 5a; Extended Data Fig. 10c). B0SMK8 and FlaAP, adjacent paralogues with inverted quasi-symmetric core contacts [26], differ in that only B0SMK8 induces a seam. FlaB protomers display continuous conformational variation (Extended Data Fig. 10d); seams provide a structural mechanism to accommodate this flexibility while maintaining overall helical architecture.

Together, these observations establish that core flagellin identity is directly coupled to sheath organization, seam distribution, and filament curvature, revealing an integrated architectural system that tunes flagellar mechanical properties essential for *Leptospira* motility in environments of varying viscosity.

## Discussion

We report high-resolution structures of native endoflagellar filaments from saprophytic and pathogenic *Leptospira* species, revealing unprecedented architectural complexity in bacterial flagella. These structures comprise helically symmetric flagellin cores enveloped by 9–12 distinct sheath proteins organised along radially asymmetric architectures. Combined with genetic and phenotypic analyses, our results demonstrate that core flagellin isoform identity is coupled to sheath organization, together dictating filament curvature. Critically, this architectural plasticity proves essential for motility in viscous environments and for virulence, establishing a mechanistic framework linking protein asymmetry to the tissue-penetrating capabilities of these pathogens.

### Viscosity-dependent motility and virulence

The *L. interrogans flaB1^-^* mutant provides a striking demonstration that filament architecture is tuned for specific environmental conditions. In low-viscosity liquid, the mutant swims faster than *wt*, yet this advantage reverses dramatically in viscous media, culminating in loss of dispersion in semi-solid agar and complete virulence attenuation in the mutant strain. Thus, higher-curvature FlaB4-dominated filaments appear suited to low-resistance aquatic environments, while lower-curvature FlaB1-dominated filaments are optimized for the viscous conditions encountered during tissue invasion. Flagellar architecture has similarly been linked to motility in viscous environments in externally flagellated bacteria [27]. Lower-curvature filaments are expected to generate smaller-amplitude body deformations, which may improve swimming efficiency and reduce energetic costs under high-viscosity conditions. The architectural complexity we describe here provides a structural basis for such tunability. Whether *Leptospira* actively regulate FlaB isoform expression in response to environmental cues remains unexplored.

### Mechanisms of curvature modulation

Previous work established that asymmetric sheath components promote filament curvature in *L. biflexa*: progressive loss of FcpA and FcpB leads to straightening, with wild-type filaments exhibiting curvature of ∼5 µm^-1^ [14]. Our structures now reveal how this curvature is achieved at atomic resolution and demonstrate an additional layer of regulation through core flagellin identity. Classical models of flagellar supercoiling proposed discrete L-state and R-state flagellin conformations mixed in varying stoichiometries [19, 28, 29]. However, elastic deformation within each state (quasi-equivalence) also contributes substantially to filament flexibility [26], and recent cryo-EM structures revealed that flagellin adopts a continuum of conformations rather than two discrete states [30]. Our *Leptospira* structures are consistent with this view: FlaB protomers display continuous conformational variation, accommodated by seams where helical symmetry is locally broken [15]. The similar curvatures of FlaB4-containing *L. biflexa wt* and *L. interrogans flaB1^-^* mutant filaments (Δκ = 0.6 μm^-1^), contrasting with the substantially lower curvature of FlaB1-dominated *L. interrogans wt* (Δκ = 2.1 μm^-1^ from *L. biflexa*), demonstrate that core flagellin identity, coupled with corresponding sheath architecture, primarily dictates filament curvature. The radially asymmetric sheath creates differential mass distribution around the filament axis, altering area moments of inertia that will likely generate unbalanced force vectors [31]. This is analogous to eccentric loading in elastic rods, which produces characteristic bending moments [32]. A similar principle has been demonstrated in asymmetrically decorated actin filaments where protein interactions modify bending stiffness [33]. Introducing a fully symmetric sheath (as in the *L. interrogans flaA2^-^*mutant) produces straight filaments [15], confirming that sheath architecture templates curvature. The asymmetric sheath architecture thus creates the mechanical context for curvature, though the detailed relationship between sheath-core contact geometry and local symmetry breaking remains to be established.

### Core-sheath coupling

Our data reveal a hierarchical, unidirectional relationship in which core identity determines sheath architecture, which in turn dictates curvature. Perturbing the sheath does not alter core composition, as shown by *flaA2^-^* mutants in both *L. interrogans* [15] and *L. biflexa* (CVS, unpublished observations). These mutants assemble symmetric sheaths yet retain their native FlaB isoform dominance, producing straight, non-motile filaments. In contrast, perturbing the core reorganizes the sheath. The *L. interrogans flaB1^-^* mutant, in which FlaB4 replaces FlaB1, assembles a *L. biflexa*-like sheath architecture and exhibits correspondingly increased curvature, while remaining functional and tuned for different mechanical requirements. Thus, core identity acts upstream, selecting among functional sheath configurations, while loss of sheath asymmetry collapses the system outside its physiological operating range.

What molecular features enable this selectivity? Position 137 provides a direct structural answer (Fig. 5a): the Thr137 *O*-glycan on FlaB4 stabilizes B0SMK8 at the I–J interface, whereas Gln137 on FlaB1 cannot be glycosylated, abolishing this contact and triggering Ser126 glycan rearrangement that eliminates the I–J seam. Electrostatic complementarity reinforces this selectivity: the FlaB1 core exposes a more cationic surface than FlaB4, particularly at loop 142-RGSR-145, further disfavouring B0SMK8 binding while permitting FcpA substitution.

### Conservation beyond *Leptospira*

Comparative structural analysis revealed clusters of structurally similar sheath proteins within species (orthologue variants) and across Spirochete genera (paralogues), despite sequence divergence (Supplementary Table 6). Only some of the proteins from each cluster were observed in our cryo-EM structures yet clearly detected by LC-MS/MS (Supplementary Table 3). This suggests that additional/alternative components may be incorporated at different lifecycle stages or at precise filament locations.

FlaA proteins are conserved phylum-wide, often as multiple paralogues, with *Treponema* and *Brachyspira* also possessing FlaA2-associated proteins suggesting conserved roles in asymmetric sheath assembly. In contrast, FcpA and FcpB are unique to *Leptospira* [10, 12, 13], likely representing genus-specific adaptations for enhanced torque transmission through a single filament per cell pole.

Spirochetal FlaA proteins define a new carbohydrate-binding module family characterized by a bent jelly-roll topology, calcium-dependent fold stabilization, and conserved sugar-recognition grooves. FlaA2 binding to FlaB *O*-glycans demonstrates functional CBM activity [15]. Intriguingly, FlaA1 exposes a putative sugar-binding groove on the filament surface, with consecutive dimers spaced ∼50 Å apart, within the range of peptidoglycan oligosaccharide spacing [34], suggesting possible filament–cell wall interactions.

### Conclusions

This study introduces the most complex flagellar filament architecture characterized to date in bacteria, revealing how protein asymmetry creates mechanically tuned assemblies. The demonstration that core flagellin identity is coupled to sheath organization, together dictating curvature, provides a structural basis for environment-specific motility. How the asymmetric filament couples mechanically to the crosslinked spirochetal hook during torque transmission remains an open question. The complete virulence attenuation of the *flaB1^-^* mutant establishes that architectural tunability is essential for tissue invasion by pathogenic *Leptospira*, and may represent a conserved strategy enabling spirochetes to optimize motility across the diverse environments encountered during their complex lifecycles.

## Methods

### Bacterial strains and culture conditions

*Leptospira biflexa* serovar Patoc strain Patoc I (*wt*), Patoc *fcpA^-^* knockout mutant [10], and Patoc *flaA2^-^* (this study); *L. interrogans* serovar Manilae strain L495 (*wt*), Manilae *flaA2⁻* [9], and Manilae *flaB1⁻* (this study) strains; were all cultured in liquid Ellinghausen-McCullough-Johnson-Harris (EMJH) medium at 30°C [35]. Cultures were grown to mid-logarithmic phase (optical density ∼0.3 at 420 nm) before harvest for flagellar purification.

### Generation of *Leptospira* mutant strains

The *L. interrogans* serovar Manilae *flaB1^-^* mutant (disruption of gene LIMLP_09410) was obtained by transposon mutagenesis using Himar1 as previously described [36]. The transposon insertion site was confirmed by PCR with primers flaB1F (5’-TATCTAAGAAGCTGCAGAAC-3’) and flaB1R (5’-GATCATTAATCACAACTTAG-3’) flanking the transposon insertion sites. Whole genome sequencing was also performed to rule out off-target modifications. Briefly genomic DNA from *L. interrogans* serovar Manilae wild-type (WT) and *flaB1^−^* strains was extracted with the Maxwell 16 cell DNA purification kit (Promega) and sequenced by Illumina next-generation sequencing. Sequence Reads were cleaned using fqCleaner and mapped to the reference genome of *Leptospira interrogans* serovar Manilae strain UP-MMC-NIID LP (accession numbers CP011931, CP011932, CP011933) using the Burrows-Wheeler Alignment tool version 0.7.5a. The detection of single nucleotide polymorphisms (SNPs) and insertions or deletions (Indels) was performed utilizing the Genome Analysis Toolkit (GATK2), adhering to the recommended practices by the Broad Institute. For targeted mutagenesis of *flaA2* in *Leptospira biflexa*, a suicide vector carrying a kanamycin resistance cassette flanked by 1 kb of the *flaA2* flanking sequences was synthesized by GeneArt (Life Technologies, Grand Island, NY, USA), pretreated by UV, and used to transform *L. biflexa* as previously described [37]. Cells were selected on EMJH agar containing 50 μg/mL kanamycin, and double-crossover mutants were identified by PCR.

### Motility assays

#### Dispersion in semi-solid medium

Mid-logarithmic phase *Leptospira* cultures (approximately 2×10⁸ bacteria/mL) were inoculated (3 µL) into 0.8% soft-agar medium supplemented with EMJH and incubated for 10 days at 30°C. Dispersion halo diameters were measured manually and normalized to *wt* values. *L. interrogans flaA2^-^* was used as a non-motile control. Experiments were performed in at least three independent replicates per strain.

#### Velocity measurements in varying viscosity

Late-logarithmic phase cultures were diluted in fresh EMJH medium with or without methylcellulose (0%, 0.5%, or 1% from a 2% stock solution in water) to obtain approximately 20 bacteria per field of view. A 5 µL droplet of diluted culture was deposited on a glass slide and covered with a coverslip elevated by silicone spots to minimize compression against the glass surface and Brownian motion. Slides were observed using an Olympus BX53 darkfield microscope connected to a Hamamatsu Orca Flash camera. Multiple 20-second videos were recorded using CellSens Dimension software (Olympus). Approximately 60 bacteria per strain were manually tracked to determine mean swimming velocity.

### Epithelial translocation assay

The ability of *Leptospira* strains to cross epithelial barriers was assessed using a Transwell® assay as previously described [38]. Briefly, canine kidney epithelial cells (MDCK-1) were seeded onto Transwell® cell culture inserts (3.0 µm porosity) and cultured at 37°C with 5% CO₂ in Eagle’s Minimum Essential Medium (EMEM) supplemented with 10% inactivated fetal bovine serum and 2 mM L-glutamine until confluent. Cell layer integrity was verified using trypan blue exclusion. Inserts were washed with EMJH/EMEM mixture (1:2 ratio), and mid-logarithmic *Leptospira* (4×10⁷ bacteria/mL) were added to upper chambers. Bacteria that successfully crossed the cell layer were collected from lower chambers and counted manually using a Petroff-Hausser chamber at 6 h and 24 h post-inoculation. Generation time of *L. interrogans* (∼20 h) was considered for bacterial load calculations at the 24 h timepoint.

### Animal infection model

Four-week-old Golden Syrian hamsters (RjHan:AURA, Janvier Labs; n = 4 per group) were infected by intraperitoneal injection with 10⁶ *Leptospira interrogans* serovar Manilae (*wt* or *flaB1^-^*) counted with a Petroff-Hausser counting chamber. Animals were monitored daily and euthanized by CO_2_ inhalation upon reaching predefined endpoint criteria (signs of distress and morbidity). Survival was recorded for 20 days post-infection. All animal procedures were approved by the Institut Pasteur Animal Care and Use Committee (CETEA #220016) and the French Ministry of Agriculture, in accordance with European Union Directive 2010/63/EU for the protection of animals used for scientific purposes.

### Flagellar filaments purification

Flagellar filaments from *L. biflexa* and *L. interrogans* strains were purified using reported protocols [10]. Purified filaments were assessed by SDS-PAGE and negative-stain transmission electron microscopy to confirm purity and structural integrity.

### Single particle analysis cryo-electron microscopy

#### Sample preparation

Purified flagellar filaments at ∼0.5-1.0 mg/mL were applied to Quantifoil R1.2/1.3 Cu300 mesh holey carbon grids; prior to sample application all grids were plasma discharged in a Gatan Model 950 Solarus Advanced Plasma System using H_2_O_2_. Grids were blotted for 6 seconds, with the blotting force parameter set to 2, at 18 - 22 °C and 100% humidity using a Vitrobot Mark IV® (Thermo Fisher Scientific) before plunging into liquid ethane.

#### Data collection

Cryo-EM data were collected on a Titan Krios operated at 300 kV equipped with a Quantum LS Model 1967 energy filter (20 eV slit width) and K3 direct electron detector. Micrographs were recorded in super-resolution counting mode at nominal magnification of 81000× (calibrated pixel size of 1.068 Å for all samples, with a super-resolution pixel size 0.534 Å) using Serial-EM [39]. Total electron doses of ∼25 and ∼60 e⁻/Å², for *L. biflexa* and *L. interrogans* samples, respectively, were fractionated across 42 frames with a frame time of ∼0.1 s. Defocus values ranged from -1.5 to -2.6 μm. Total dataset sizes: 1985, 1081, 1094 micrographs for *L. biflexa wt*, *L. interrogans wt* and *L. interrogans flaB1^-^*, respectively.

#### Image processing

Cryo-EM refinement and 3D reconstruction was performed using cryoSPARC [16] using a specialized workflow to account for supercoil deviations from strict helical symmetry. CryoSPARC version 3.3.2 was used for all except the final stages of refinement, which used the updated version 4.6.2 where noted below. All three structure refinements followed the same general protocol (Extended Data Fig. 2, 5 & 8a). After initial micrograph processing (full-frame motion correction, patch CTF estimation), the filament tracer tool was first applied for particle picking using the template-free option, followed by one or more rounds of 2D classification and manual particle selection. The filament tracer tool was then run a second time using the final manually selected 2D classes as templates. Particles identified by this latter step were initially extracted with 192x192 box size and 2.136 Å nominal pixel size, followed by multiple rounds of 2D classification and manual selection to eliminate false positive junk particles such as high-contrast carbon edges. The resulting particles were subjected one or more times to non-symmetrized helical refinement (option *‘maximum symmetry order to apply during reconstruction’* set to 1), starting with a generic sheath-less flagellar filament reference volume filtered to 20 Å resolution, resulting in reported resolutions of ∼6 Å. The resulting refined particle alignments were then used to re-extract 384 box-size,1.068 Å nominal pixel-size image segments. Following additional rounds of helical refinement, 2D classification, local/global CTF refinement and/or anisotropic magnification correction, the heterogenous refinement job was used to generate 3-5 structure classes. Particles belonging to artefactual volumes or structures with missing sheath densities were then discarded. Classes that were kept included volumes that were rotated and translated with respect to each other by the bacterial flagellar pseudo-symmetry operator (+/- ∼26 Å z-shift and +/- ∼32.7° z-rotation). The final stage of structure refinement utilized the ‘non-uniform refinement’ job, which overcame local minima due to pseudo-symmetric false alignments but also shifted some nearby particles on top of each other. This latter effect introduced particle duplications between the two gold standard half data sets, artificially boosting FSC resolution estimates. This resolution artifact was subsequently eliminated by running a ‘remove duplicates’ job, followed by a ‘local refinement’ job where additional particle duplications were prevented by restricting particle translation and rotation changes to +/- 7.5 Å (15 Å extent) and +/- 5° (10°), respectively (recentring of rotations and shifts after each iteration was disabled). Reported final map resolutions correspond to the final, auto-tightened mask generated by cryoSPARC. Density modification was subsequently performed using RESOLVE [17] from the Phenix package [40] to improve map interpretability for structure building. This required removing low-frequency background solvent fluctuations from 3D map corners, which was accomplished by using relion_image_handler [41] to extract the central 256^3^ zone from final cryoSPARC output raw 384^3^ half-maps. An estimated protein occupancy of 0.52451 was input to RESOLVE based on identified protein components, flagellar geometry and volume size. Final map gold-standard FSC resolutions were 3.5 Å (176,558 particles) for *L. biflexa*, 4.4 Å (35,760 particles) for *L. interrogans wt*, and 4.3 Å (53,501 particles) for *L. interrogans flaB1^-^*mutant. Estimated resolution improvements reported by resolve were ∼0.15-0.2 Å. Local resolution was calculated using the Phenix package (phenix.local resolution) [40].

#### Model building

FlaB models were predicted with AlphaFold3 [42], rigid-body fitted into the cryo-EM maps with ChimeraX v1.10.1[43], and initially refined interactively using Coot v1.1.18 [44]. Sheath proteins were initially fitted within cryo-EM maps similarly, using: (i) crystal structures of FcpA and FcpB [14], including required sequence substitutions for each *Leptospira* species; (ii) AlphaFold3 predictions for remaining components; and (iii) crosslinking mass spectrometry data to decide in ambiguous cases. For the *L. interrogans flaB1^-^* mutant, initial sheath protein models were derived from the *L. biflexa* structure given the architectural similarity revealed by cross-correlation analysis (see Results), with sequence substitutions for *L. interrogans* orthologues where applicable. Models were further refined interactively with Coot using intra-chain self-restraints (6 Å sphere), and real-space chain refinement with Geman-McClure restraints (α parameter 0.01). A conservative approach was followed for all structures with backrub rotamers, deleting side chains, residues or short loops in regions where the cryo-EM density was not reliable (undetectable density at sensible contour levels). Model validation was performed continuously within Coot. Secondary structure restraints were calculated from the starting models using phenix.secondary_structure_restraints as implemented in Phenix v1.21.2-5419 [40]. These extra restraints were included in first real-space refinement cycles with phenix.real_space_refine v2.0-5936 using Ramachandran restraints [45]. Servalcat v0.4.128 [46] was then used to further refine atomic models in reciprocal space against the final unsharpened and unweighted cryo-EM half-maps. External geometric restraints based on initial structures (ProSMART v0.860 [47]) was used throughout Servalcat refinement. Final validation was done with MolProbity [48] implemented within Phenix. Checkmysequence v1.5.3 was instrumental to validate sequence assignment and correct map interpretation errors [49].

### Quantitative filament curvature analyses

Helical filament coordinates were extracted from RELION star files following helical refinement using custom bash scripts (Supplementary Data 1 & 2). Particle positions were corrected for origin shifts by subtracting *rlnOriginXAngst* and *rlnOriginYAngst* values from micrograph coordinates (in pixels) multiplied by pixel size (3.39 Å for *L. biflexa* and *L. interrogans wt*, 3.54 Å for *L. interrogans flaB1^-^*), yielding actual 2D trajectory positions in Ångstroms. Three-dimensional coordinates were reconstructed from 2D positions and tilt angles (*rlnAngleTilt*) by iteratively calculating z-coordinates as z(n) = z(n-1) + cos[(θ(n) + θ(n-1))/2] × d(n,n-1), where d represents the Euclidean distance between consecutive particles in the xy-plane.

Local curvature (κ) and torsion (τ) were computed using the method of Crenshaw [25] with a separation parameter of 10 particles. For each filament segment spanning 2×sep positions, the algorithm constructs tangent (T), normal (N), and binormal (B) vectors to determine local geometric properties. Curvature κ* at position i is calculated as κ*(i) = 2·arctan(sin φ / cos φ) / (|C| + |D|), where φ is the angle between vectors C and D spanning the segment, and the final curvature is the average of consecutive κ* values multiplied by 10^4^ for convenient scaling. Torsion τ is calculated as the angle between consecutive binormal vectors along the trajectory.

Individual curvature measurements were treated as sampling points of the intrinsic filament curvature, with random perturbations introduced during cryo-preservation averaged out across high n. *L. biflexa wt* (median κ = 5.6 μm^-1^, IQR 4.5 - 7.1; n = 28,853 from 409 filaments), *L. interrogans flaB1⁻* (median κ = 5.0 μm^-1^, IQR 4.3 - 6.0; n = 55,579 from 1,148 filaments), and *L. interrogans wt* (median κ = 3.5 μm^-1^, IQR 2.5 – 4.8; n = 20,977 from 309 filaments), were compared using non-parametric methods. Overall differences were assessed using Kruskal-Wallis test (H = 15,798, p<<0.0001); pairwise comparisons used Mann-Whitney U tests (all p<<0.0001). Given the large sample sizes, effect sizes provide more informative measures of difference magnitude than p-values: Cohen’s d was 0.83 (large) for *L. biflexa vs L. interrogans wt*, 0.60 (medium) for *flaB1^-^ vs L. interrogans wt*, and 0.25 (small) for *L. biflexa vs flaB1^-^*, indicating that the FlaB4-dominated strains (*L. biflexa wt* and *L. interrogans flaB1^-^*) cluster together while FlaB1-dominated *L. interrogans wt* is set apart. Bootstrap 95% confidence intervals for median differences confirmed non-overlapping intervals for all comparisons (Δκ = 2.02 [1.98–2.05], 1.44 [1.41–1.47], and 0.58 [0.55–0.60] μm^-1^, respectively). All statistical analyses were performed in Python 3.12 using SciPy and NumPy.

### Comprehensive mass spectrometry and crosslinking network analyses

#### Label-free quantitative comparative proteomics

Purified flagellar filaments (∼20 µg) were embedded into polyacrylamide gels by short electrophoresis in the presence of SDS, and Coomassie-stained to excise the protein band. Gels were destained, reduced with dithiothreitol, alkylated with iodoacetamide, and digested overnight with sequencing-grade trypsin. (Promega) as described previously [50]. Peptides were extracted, with 60% acetonitrile, 0.1% trifluoroacetic acid, dried, desalted with ZipTip® C18 tips (Millipore) and resuspended in 0.1 % formic acid. Samples were analysed by LC–MS/MS on an UltiMate 3000 nano-HPLC (Thermo Fisher Scientific) coupled to a Q-Exactive Plus mass spectrometer (Thermo Fisher Scientific). Peptide separation was performed on an Easy-Spray analytical column (PepMapTM RSLC, C18, 75 µm X 50 cm, 2 µm particle size), using a 90-min 1–35 % acetonitrile separation gradient at 200 nL.min^-1^ (mobile phases: 0.1% formic acid in water, 0.1% formic acid in acetonitrile). Mass analysis was carried in a data-dependent mode: acquisition of full MS scans in a range of m/z from 200 to 2000; followed by HCD fragmentation of the 12 most intense ions in each segment, using a stepped normalized collision energy of 25, 30 and 35 and dynamic exclusion list. Protein identification and label-free quantification were performed with PatternLab for Proteomics V [51]. A target-reverse *L. biflexa* serovar Patoc database (downloaded from UniProt on 12/2021) was used for protein identification. Oxidation of methionine was included as a variable, and carbamidomethylation as a fixed modification. Initial mass tolerance for the measured precursor m/z was 35 ppm. Peptide spectrum matches were filtered to achieve <10 ppm of tolerance for precursor, and less than 1% of false discovery rate (FDR) at the protein level. Differential abundance analyses separately compared *wild-type L. biflexa* with *fcpA^-^*and with *flaA2^-^* mutants using spectrum counting. PatternLab for Proteomics was used to statistically identify proteins exclusively detected in each condition (p<0.05) and to pinpoint proteins with an FC > 2 (q < 0.05).

#### Crosslinking mass spectrometry

Purified *Leptospira* filaments were treated with NNP9 following established protocols [52], including some adaptations. Briefly, NNP9 was used at 5 nmol per 10 µg protein, for 1 h at 4°C to capture native protein:protein contacts. Reactions were quenched with ammonium bicarbonate, and crosslinked complexes were purified by enhanced filter-aided sample preparation (eFASP) on photocleavable agarose beads. After on-filter tryptic digestion, peptides were released by UV irradiation (365 nm), dried, and resuspended for LC–MS/MS analysis on a Q-Exactive HF mass spectrometer (Thermo Fisher Scientific). Crosslinked peptides were separated on a home-made C18 nanocolumn (75 µm ID, solid phase: Aeris C18, 1.7 µm, Phenomenex) using a 90 min hydrophobic gradient in presence of 0.1% formic acid (from 8% to 30% B [80% acetonitrile, 1% formic acid]) in 70 min, follow by a step up to 60 % B in 20 min). Data-dependent acquisition was performed in positive polarity at 60,000 resolution (both MS and MS/MS) from 300 to 1800 m/z in MS and 200 to 2000 m/z in MS/MS, excluding singly and doubly charged species. Crosslinks were identified using Mass Spec Studio [53] with ±10 ppm mass tolerance and filtered to 1 % FDR after removal of peptides identified in less than 2 MS/MS events. Crosslink spectrum matches were then carefully checked manually. Validated crosslinked residue pairs were mapped onto structural models using PyMOL/PyXlinkViewer, and interaction networks were visualized in xiVIEW [54] (https://www.xiview.org), enabling integrative interpretation of filament architecture and inter-subunit connectivity. The crosslinking interaction network (Fig. 1c) was exported for interactive web-based exploration publicly accessible at https://www.xiview.org/network.php?upload=42923-71803-52415-45391-89719; allowing users to explore individual crosslinked residue pairs, filter by confidence scores, and examine protein-protein connectivity.

### FlaA1 X-ray crystallographic analysis

#### FlaA1 expression and purification

*L. borgpetersenii* FlaA1 (UniProt Q04UP1) was expressed in plasmid pET32a [55] from a truncated construct (lacking the first 49 and last 16 residues) with a TEV-cleavable N-terminal 6xHisTag, in *E. coli* Lemo21 cells cultured in TB medium. The protein was overexpressed for 18 h at 20°C with 1 mM IPTG in the presence of ampicillin (100 μg/L) and cloramphenicol (34 μg/L) under agitation (220 rpm). FlaA1 was purified in the presence of 10mM CaCl_2_ by standard nickel affinity chromatography (HisTrap FF, Cytiva), TEV digestion and size exclusion chromatography (Superdex S75 10/300, Cytiva). Purified protein was treated 1 h with 10 mM dithiothreitol, followed by 1 h with 55 mM iodoacetamide at RT.

#### Crystallization and structure determination

Robotic screening using commercial sparse-matrix kits led to several hits. Optimized crystals using hanging-drop vapour diffusion grew in 0.1 M sodium cacodylate pH 6.5, 1 M ammonium sulphate, with 2 μL protein (50 mg/mL) mixed with 2 μL reservoir solution. Crystals were cryoprotected in reservoir solution supplemented with 25% (v/v) glycerol before flash-freezing in liquid nitrogen. Diffraction data were collected at Diamond Light Source beamline I04 at 100 K. Data were processed using XDS [56] and scaled with Aimless [57]. The structure was solved by molecular replacement using Phaser [58] with an AlphaFold3-predicted model as search probe. The structure was interactively rebuilt using Coot 1.1.18 and refined with phenix.refine 1.21.2-5419 [59] to 1.85 Å resolution in space group P2_1_2_1_2_1_. Data processing and model refinement statistics are detailed in Supplementary Table 5. Coordinates and structure factors have been deposited in the PDB (accession code pdb_00009cyn).

### Comprehensive Structural Homology Analysis and Cross-Spirochete Comparison

Structural homology searches were carried out with FoldSeek [60] installed locally, using all known *Leptospira* filament protein structures as queries against: (i) entire proteomes from *L. biflexa* (NCBI Taxonomy Identifier 456481) and *L. interrogans* (taxid 267671) as targets to identify paralogues, fetched from UniProt and associated AlphaFold Database entries; and, (ii) *Treponema pallidum* (taxid 243276), *Borrelia burgdorferi* (taxid 224326), and *Brachyspira hyododysenteriae* (taxid 1266923) entire proteomes, to identify spirochetal orthologues. Predicted structures from entire proteomes were fetched using a script (Supplementary Data 3). FoldSeek “easy-search” routine was used with default search parameters, except for not prefiltering (exhaustive-search); using an automatically optimized number of iterations; and, including average LDDT and probability prob in the output table (to better identify domain similarities and not only entire protein fold similarities). A specific E-value threshold was not used, contrast line between hits and noise was preferred instead. Structural families resulted in ≥0.7 estimated probability for query and target to be homologous. Sequence identities within families were calculated using clustalW [61]. Protein copy numbers for *L. interrogans* were extracted from quantitative proteomic datasets [23] if available.

## Data availability

Atomic coordinates of filament models and corresponding cryo-EM maps have been deposited in the worldwide Protein Data Bank under accession codes pdb_000010LM / EMD-75270 (*L. biflexa wt*); pdb_000010LK / EMD-75268 (*L. interrogans wt*); pdb_000010LL / EMD-75269 (*L. interrogans flaB1^-^*). Atomic coordinates and X-ray diffraction structure factors have been deposited in the wwPDB under accession code pdb_00009cyn (FlaA1 crystal structure). The mass spectrometry proteomics data have been deposited to the ProteomeXchange Consortium via the PRIDE [62] partner repository with the dataset identifiers PXD072058 (for *L. biflexa wt* and *flaA2^-^* strains); PXD030741 (*L. biflexa fcpA^-^* strain) [15]; and PXD073467 (for crosslinking data). Whole genome sequencing data of generated *Leptospira* mutant strains are available at NCBI under BioProject accession # PRJNA1416034.

## Acknowledgments

This research was supported by the Institut Pasteur and Institut Pasteur de Montevideo through the Pasteur International Unit ‘Leptospirosis Pasteur NETwork’ (LEPNET); FOCEM (MERCOSUR, COF 03/11); ANII (Uruguay) grant FCE_1_2025_1_186367 (SM, AB); ANR (France) grant LEPTOMOVE 18-CE15-0027-01 (MP, AB); National Institutes of Health (USA) grants P01 AI 168148 (MP), R01 AI052473 (AIK), R21AI163663 (EAWJ), and R01AI182354 (EAWJ). This work is part of the PhD of Fabiana San Martin (ANII and CAP fellowships), Lenka Fule (ANR LEPTOMOVE grant) and Megan Brady (NIH grant T32GM8283). We gratefully acknowledge Dr L Zarantonelli and the UMPI laboratory IPMontevideo for the microbiology BSL2 facilities; and Dr. Shenping Wu, Dr. Jianfeng Lin and the Yale Cryo-EM Resource and Yale Center for Research Computing facility, for expert support and facilities’ maintenance.

## Extended Data and Supplementary Information

**Extended Data Fig. 1.**
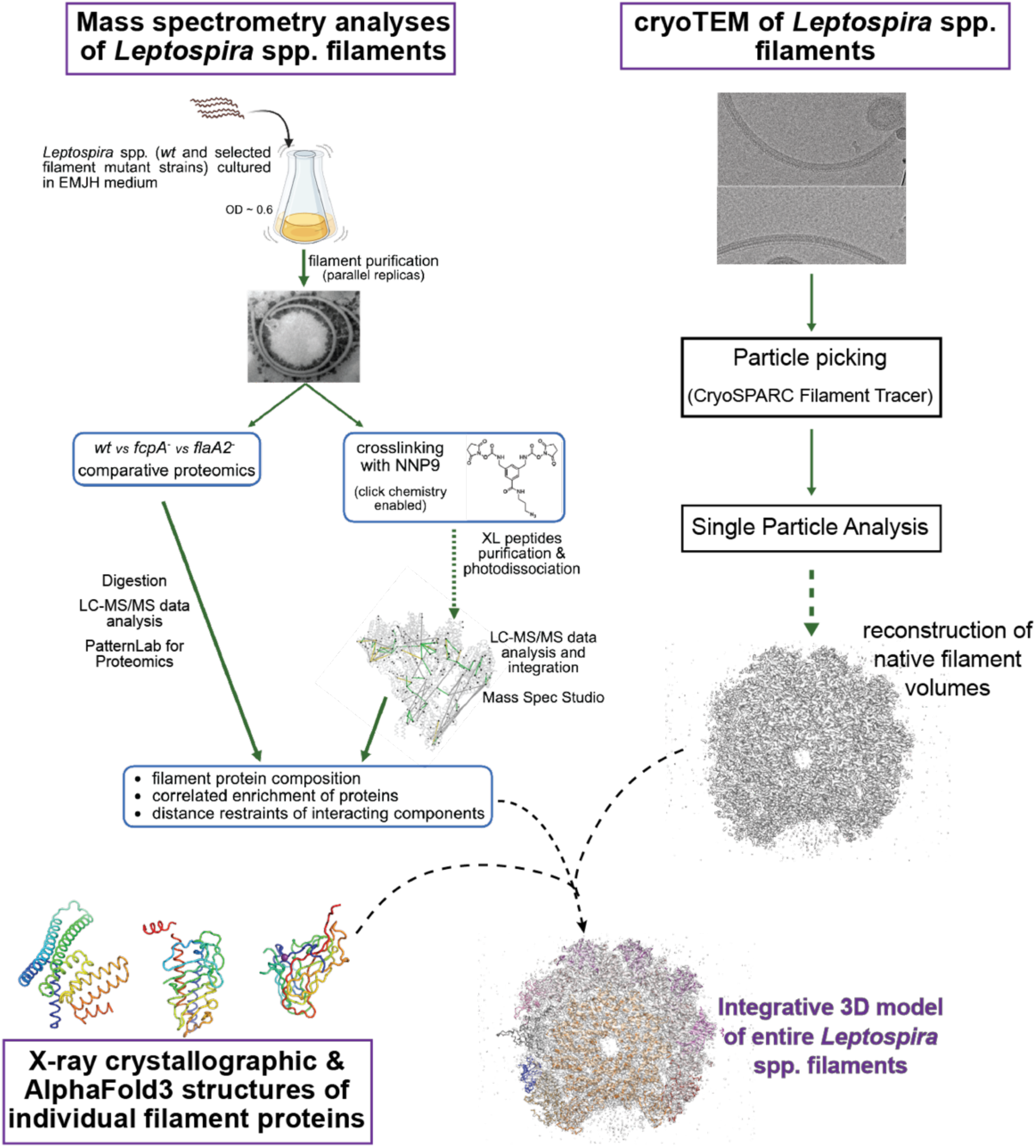
Integrative structural biology workflow. Schematic overview of the integrative approach combining three methodological branches: (i) cryo-EM single particle analysis of purified filaments to obtain 3D reconstructions (right); (ii) mass spectrometry, including comparative proteomics of *wt vs* sheath mutants (*flaA2^-^*, *fcpA^-^*) and crosslinking MS to derive distance restraints and identify co-enriched components (left); (iii) X-ray crystallographic and AlphaFold3-predicted structures of individual filament proteins (bottom). These three branches converge to generate integrative atomic models of native *Leptospira* spp. filaments.

**Extended Data Fig. 2.**
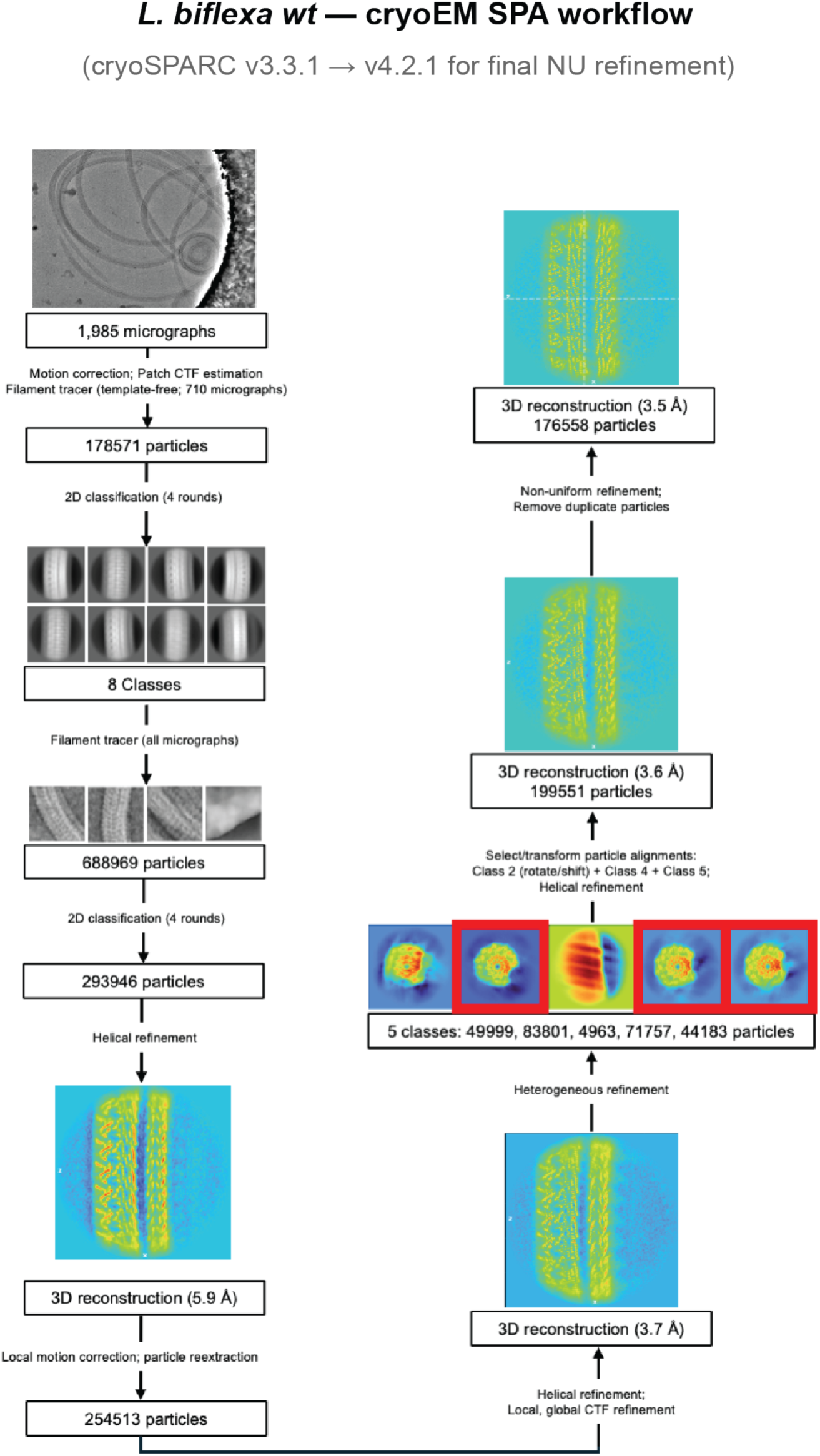
Cryo-EM single particle analysis workflow. 3D reconstruction workflow to obtain the structure of wild-type *Leptospira biflexa* filaments.

**Extended Data Fig. 3.**
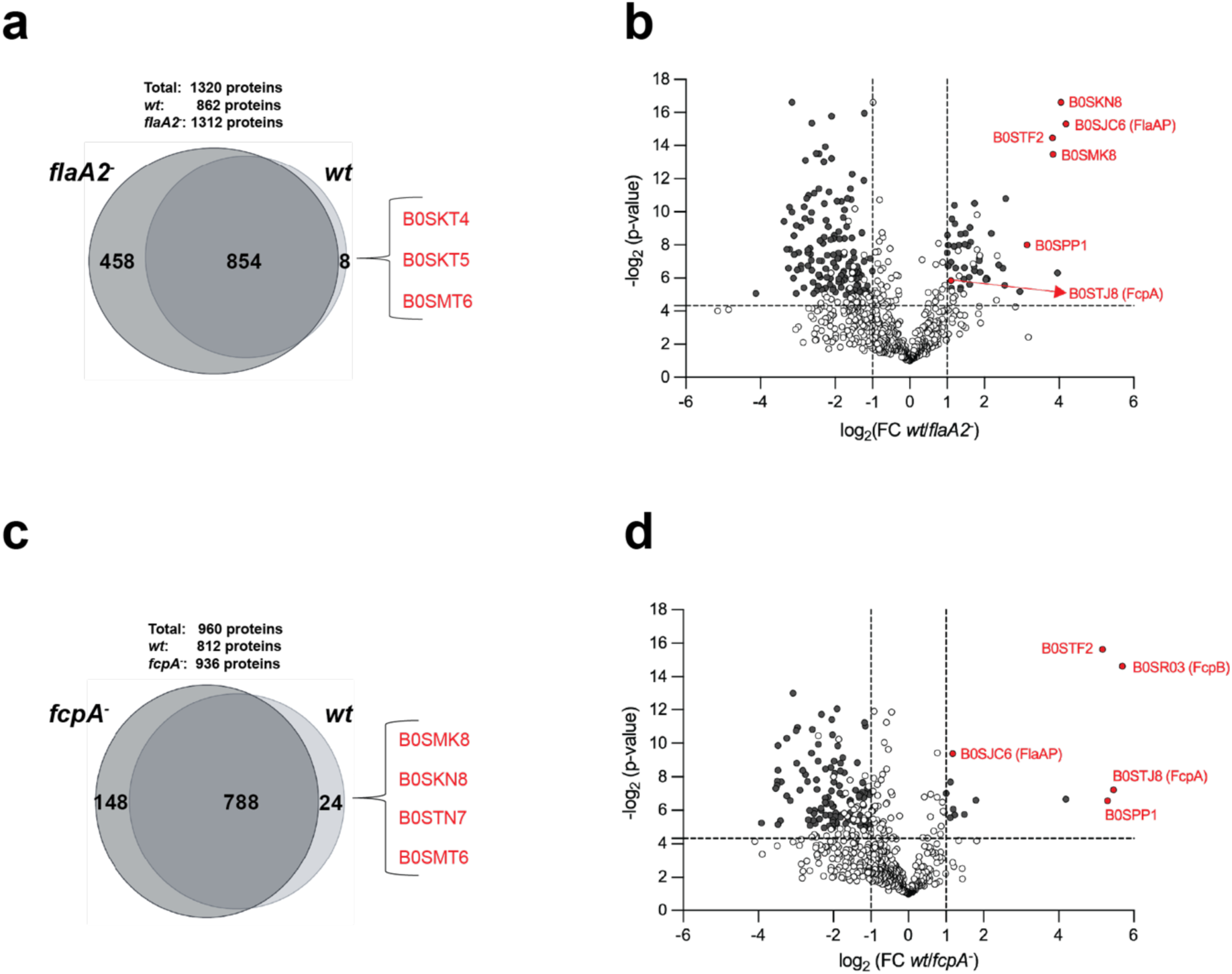
Label-free comparative proteomics of *L. biflexa* filaments from *wt* and sheath mutant strains. **(a)** Venn diagram comparing proteins identified in enriched filament preparations from *L. biflexa wt* and *flaA2^-^* strains (Bayesian probability, p < 0.05 [63]). Selected filament proteins identified in our structures and exclusively detected in *wt* are listed, with gene names in parentheses where annotated. **(b)** Volcano plot describing differential expression of proteins present in both *L. biflexa wt* and *flaA2^-^* strains. Each dot represents a protein, with darker dots being significantly differential (|log2FC| ≥ 1; p ≤ 0.05). Proteins with altered expression also identified within the cryo-EM structure of *L. biflexa* filaments, are flagged in red and labelled with UniProt IDs (gene names for those annotated). See Supplementary Table 3 for detailed information. **(c)** Venn diagram comparing *L. biflexa wt vs fcpA^-^*, as in (a). **(d)** Volcano plot following same colour/labelling scheme as in (b).

**Extended Data Fig. 4.**
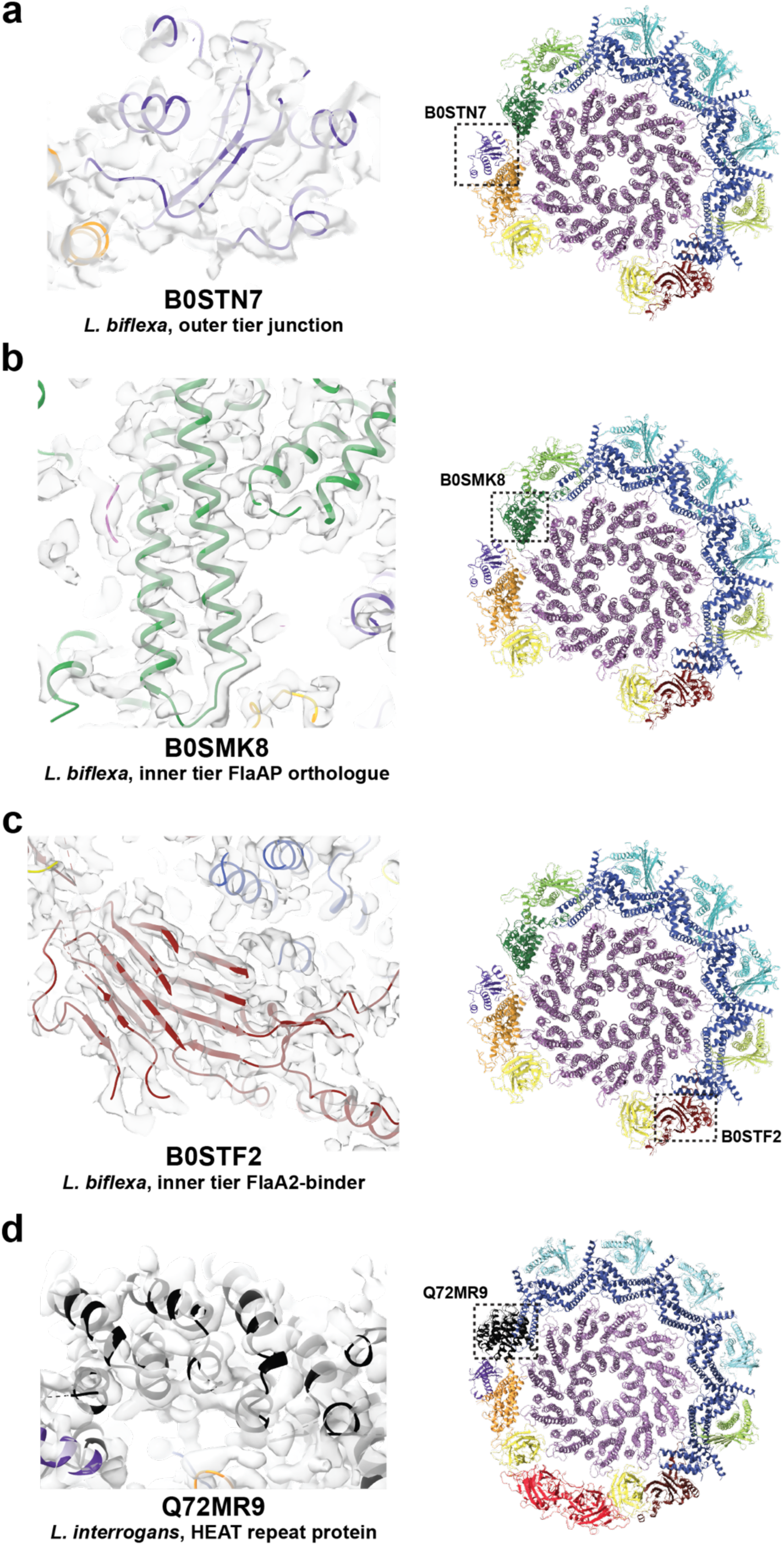
Validation of novel sheath protein assignments. Representative examples of newly identified sheath proteins fitted into cryo-EM density; all consistent with interprotein crosslinks detected by mass spectrometry (see Table 1). **(a)** B0STN7 (purple) occupies the outer tier at the junction between FlaAP and B0SMK8 in *L. biflexa*. Left: cryo-EM density (semi-transparent surface) with refined model (cartoon). Right: position within the filament cross-section (boxed) **(b)** B0SMK8 (green) adopts a FlaAP-like fold in the *L. biflexa* inner tier. In the inner tier, B0SMK8 interacts directly with the flagellin core, quasi-symmetrical to FlaAP (B0SJC6), of which it is a structural paralogue **(c)** B0STF2 (brick red) associates with one of the FlaA2 protofilaments on the concave surface in *L. biflexa*, contributing to the heterogeneous inner tier organization **(d)** Q72MR9 (black), a HEAT repeat protein unique to the *L. interrogans* sheath, directly contacts the FlaB1 core in the compositional plasticity zone. This protein is absent in FlaB4-dominated *L. biflexa wt* filaments, where the mixed B0SMK8/B0SMT6 protofilament occupies the equivalent position.

**Extended Data Fig. 5.**
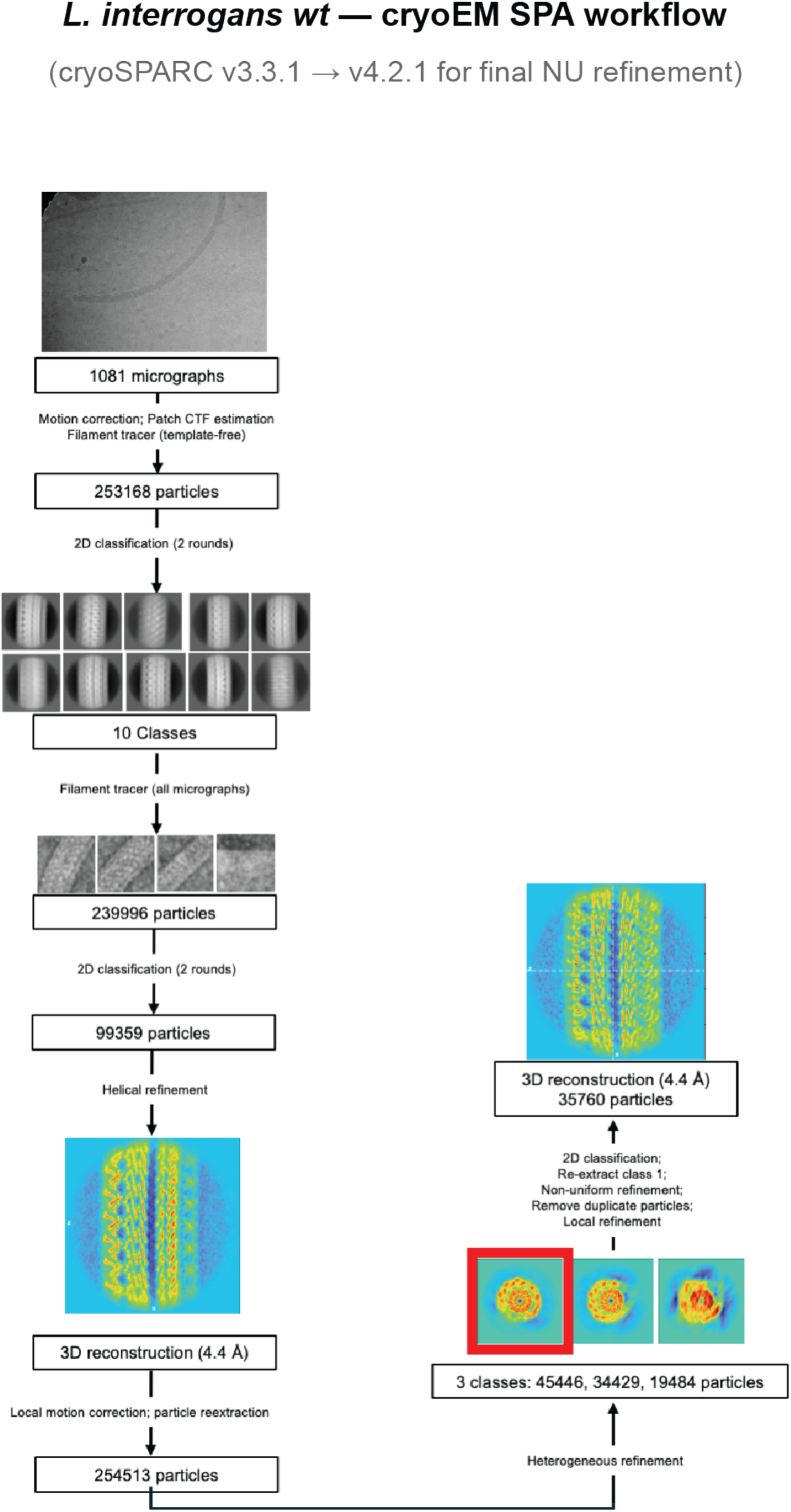
Cryo-EM single particle analysis workflow. 3D reconstruction workflow to obtain the structure of wild-type *Leptospira interrogans* filaments.

**Extended Data Fig. 6.**
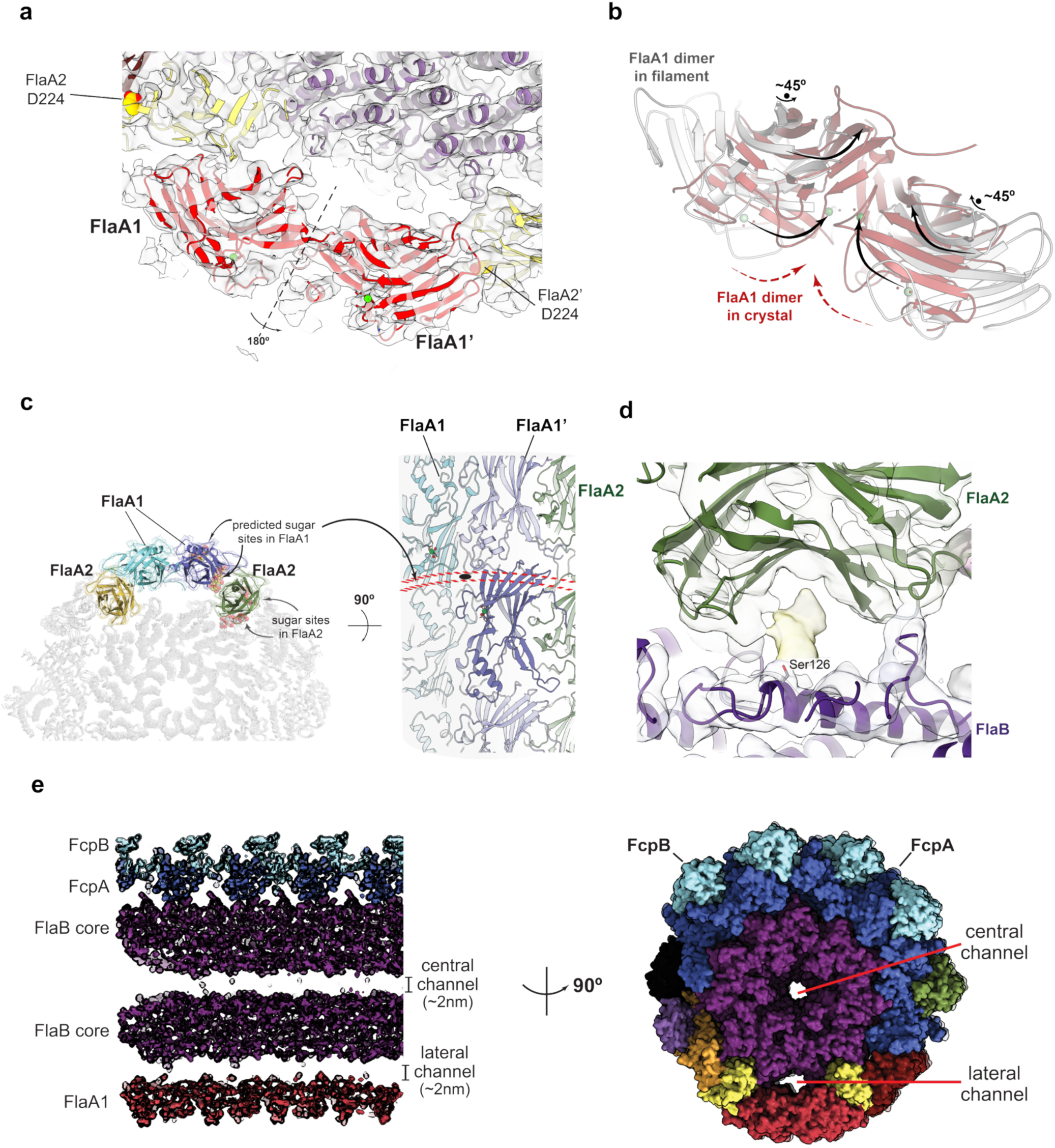
FlaA1 and FlaA2 in the assembled filament: differential orientation of carbohydrate-binding grooves. **(a)** *L. interrogans wt* cryo-EM map (semi-transparent) with fitted atomic model, viewed along the filament axis. FlaA1 (red) occupies the lateral groove as an antiparallel dimer related by a 2-fold axis perpendicular to the filament (marked). The non-equivalent FlaA1-FlaA2 nterfaces on either side are indicated by a labelled residue on FlaA2. **(b)** Comparison of FlaA1 dimer geometry in the crystal (pink) *vs* in the assembled filament (grey). A ∼45° opening motion separates he two configurations, likely induced by interactions with flanking FlaA2 protofilaments. **(c)** Left: ransverse view of *L. interrogans* cryo-EM structure showing the positions of the two FlaA2 protofilaments and the dimeric FlaA1, with putative sugar-binding sites indicated on both isoforms. Right: 90° rotated view revealing that the convex sugar-binding groove of each FlaA1 protomer faces outward; because the two protomers are related by a 2-fold axis (black ellipse), their individual grooves align to form a single continuous crevice spanning both subunits (red dotted lines; the two related FlaA1 protomers are highlighted in darker colours). **(d)** Close-up of *L. interrogans* FlaA2 engaging a FlaB1 *O*-glycan (Ser126) through its concave sugar-binding groove. The cryo-EM map (semi-ransparent) shows clear glycan density. **(e)** The lateral channel beneath FlaA1 (∼20 Å diameter), comparable in size to the central channel despite its flattened geometry.

**Extended Data Fig. 7.**
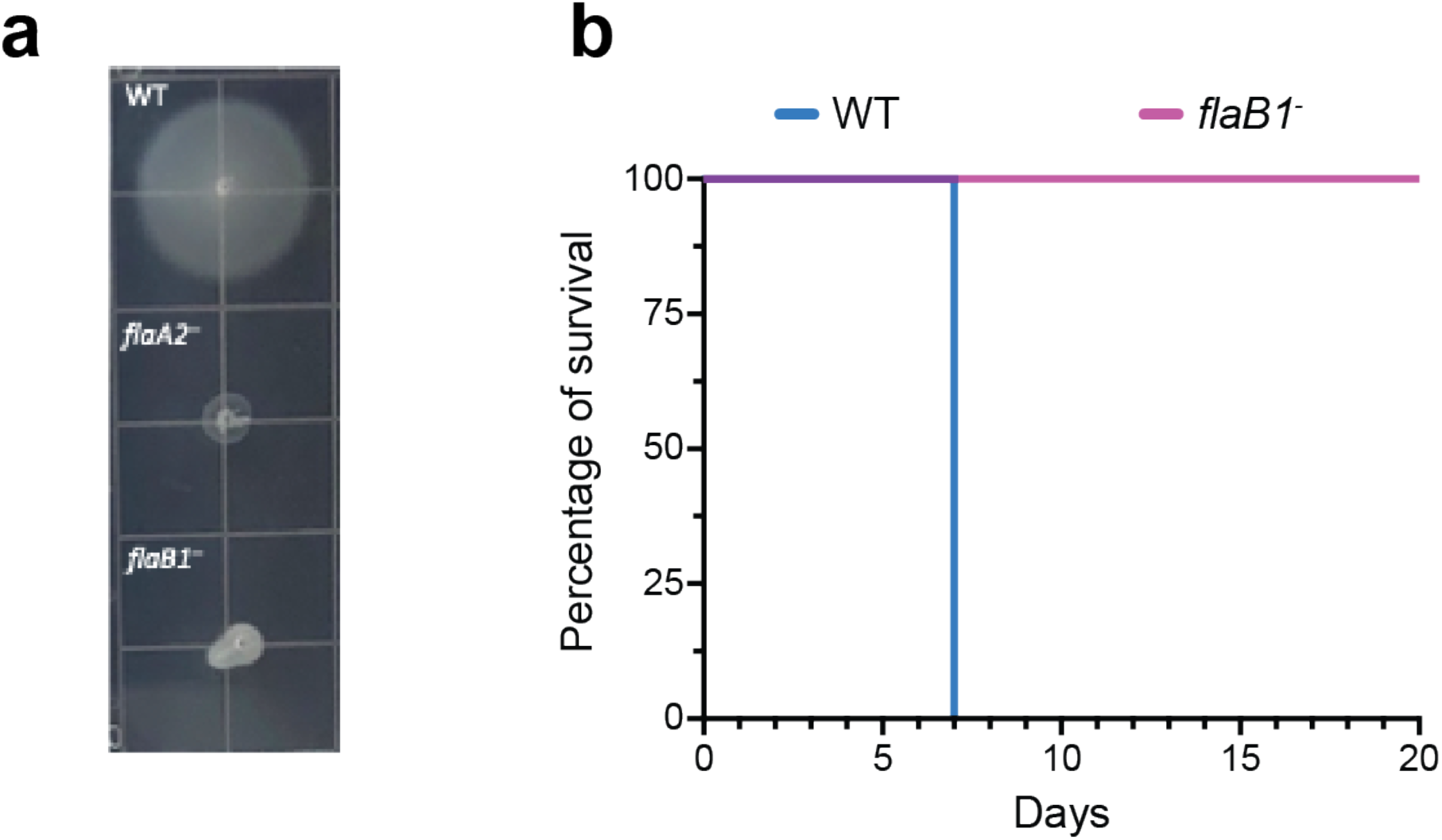
Extended phenotypic characterization of *L. interrogans flaB1^-^*mutant. **(a)** Representative images of dispersion halos in semi-solid agar for WT, *flaA2^-^* (non-motile control), and *flaB1^-^*strains after 10 days at 30°C. **(b)** Survival curves of Golden Syrian hamsters (n = 4 per group) infected intraperitoneally with 10⁶ *L. interrogans* serovar Manilae WT or *flaB1^-^*. All WT-infected animals succumbed by day 7, while *flaB1^-^*-infected animals showed 100% survival throughout the 20-day observation period, demonstrating complete virulence attenuation.

**Extended Data Fig. 8.**
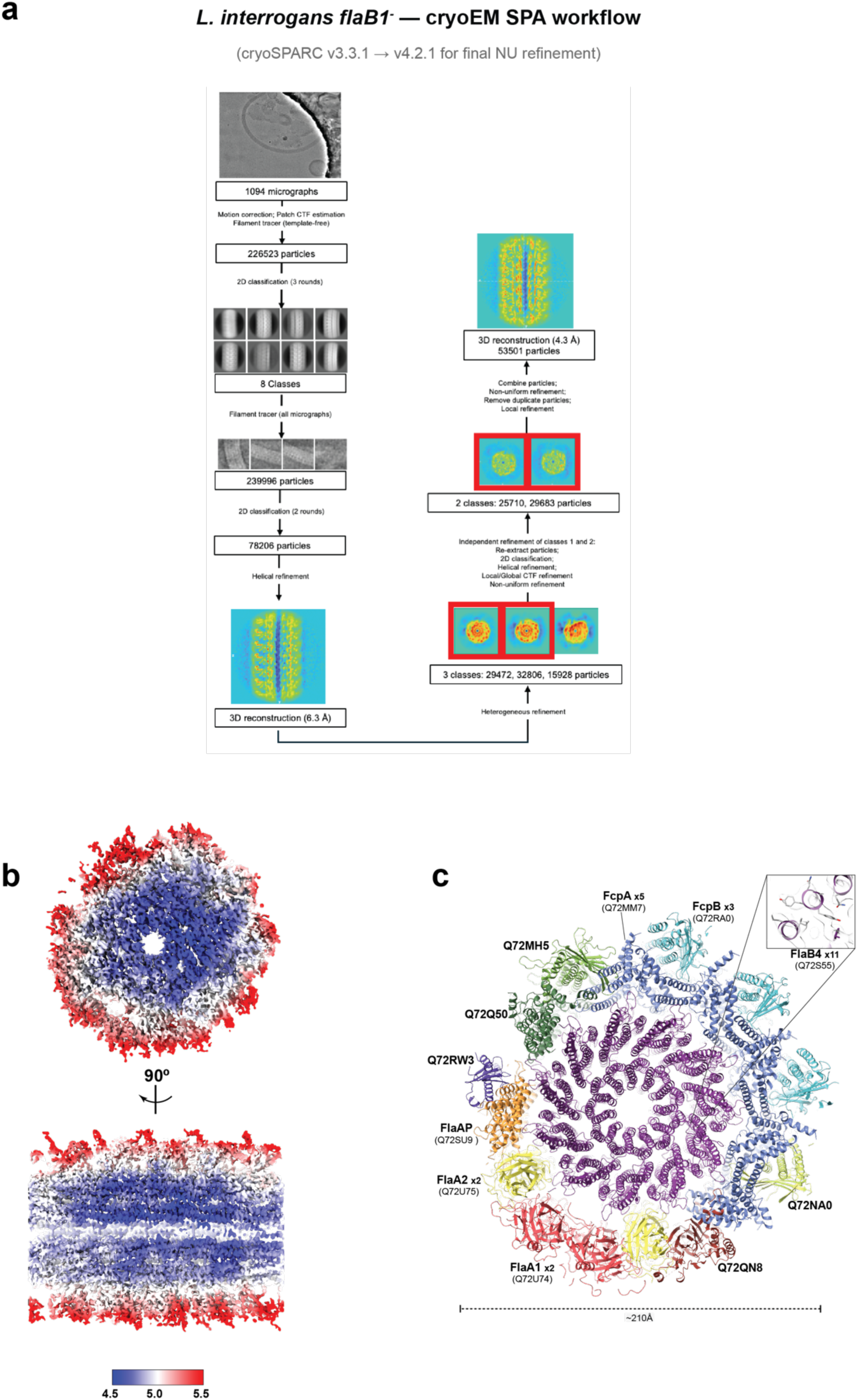
Cryo-EM structure of *Leptospira interrogans flaB1^-^* flagellar filament. **(a)** 3D reconstruction workflow to obtain the structure of *L. interrogans flaB1^-^*filaments by single particle analysis. **(b)** Cryo-EM map coloured by local resolution (scale bar: 4.5–5.5 Å), viewed from one end (top) and from the side (bottom), contoured at 0.1 threshold. **(c)** Refined atomic model shown as cartoon in transverse cross-section. Orthologous proteins are coloured as in Fig. 1. Novel *L. interrogans*-specific components are labelled distinctly. Inset: close-up of FlaB4 core density with fitted model.

**Extended Data Fig. 9.**
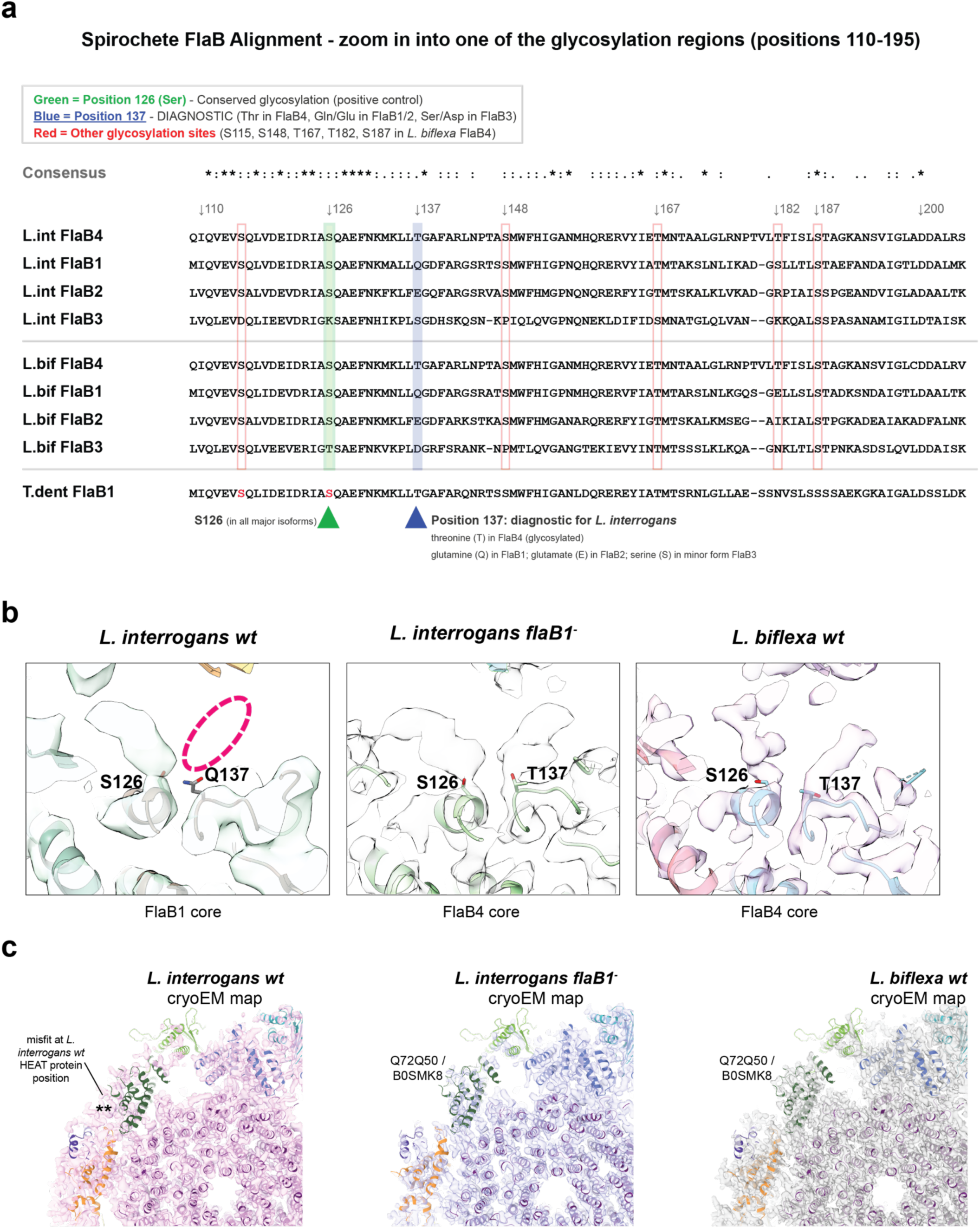
Core FlaB-induced variation of sheath architecture. **(a)** Multiple sequence alignment of *Leptospira* FlaB isoforms (L.int: *L. interrogans*; L.bif: *L. biflexa*) and *Treponema denticola* FlaB1 (T.dent), whose glycosylation sites have been determined experimentally [64]. The aligned fragment corresponds to a glycosylation-rich region (positions ∼110–200). Diagnostic sites to distinguish FlaB isoforms are detailed. Note that for some sites, *O*-glycosylation is possible only in specific FlaB isoforms: among the most expressed isoforms, position 137 can only be glycosylated in FlaB4 (FlaB3 possesses a Ser that could accept an *O*-glycan, yet FlaB3 is a significantly minor component [23]). **(b)** Comparison of cryo-EM density at position 137 across three strains. Position 126 (serine) is glycosylated in all FlaB isoforms, while position 137 distinguishes them: threonine (glycosylated) in FlaB4 versus glutamine in FlaB1 (cannot glycosylate). *Left: L. interrogans wt* (FlaB1-dominated) shows a glycan at S126 only; Q137 lacks density (highlighted). *Centre: L. interrogans flaB1^-^* mutant shows glycan density at both positions, confirming FlaB4 dominance. *Right: L. biflexa wt* (FlaB4-dominated) shows clear glycan density at both S126 and T137. Atomic models are overlaid for reference. **(c)** Comparison of cryo-EM densities at the compositionally variable sheath sector. The refined *flaB1^-^*atomic model (coloured as in Extended Data Figure 8c) is shown fitted into three maps: *L. interrogans wt* (left), *L. interrogans flaB1^-^*(centre), and *L. biflexa wt* (right), all displayed at comparable contour levels. The model fits well into both *flaB1^-^* and *L. biflexa* maps. In contrast, the *L. interrogans wt* map shows additional density (asterisks) corresponding to the HEAT repeat protein Q72MR9 absent from the *flaB1^-^* and *L. biflexa* architectures, confirming that the mutant adopts a saprophyte-like sheath organization.

**Extended Data Fig. 10.**
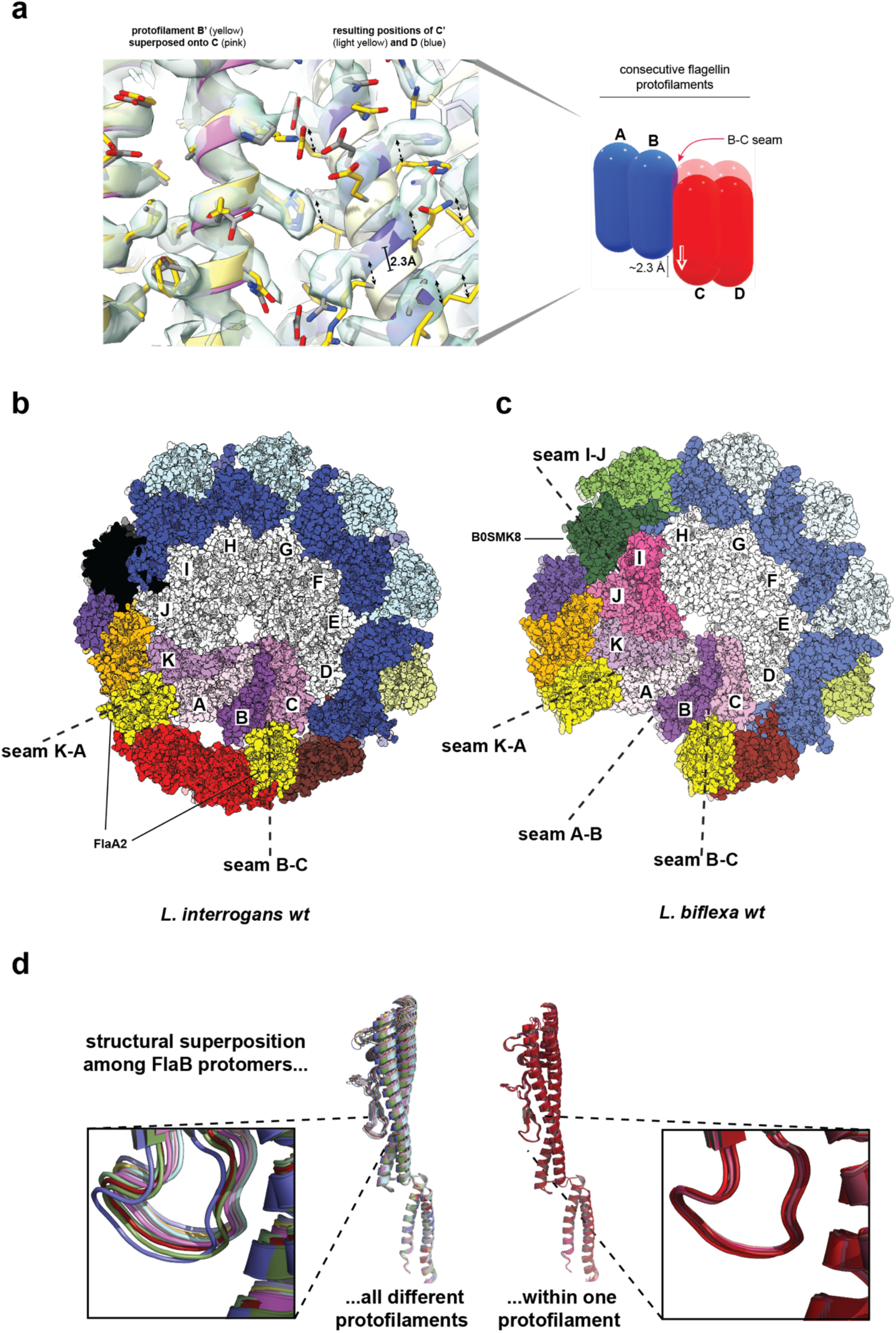
Seams and conformational flexibility in flagellin cores. **(a)** Detection of seams by structural superposition. To reveal the translational discontinuity at a seamed interface, the seamed B′–C′ pair was superposed onto the seamless C–D pair by aligning chain B′ (yellow) onto chain C (pink) (left). If both interfaces were equivalent, their partners C′ (light yellow) and D (blue) would coincide; instead, C′ is displaced ∼2.3 Å along the helical axis relative to D (right), revealing the seam. Cryo-EM density shown as semi-transparent surface. Schematic (far right) illustrates the protofilament sliding that defines seams. **(b)** *L. interrogans* filament cross-section with protofilaments labelled A-K. Seams at K-A and B-C (marked) lie beneath FlaA2 sheath moieties; seamless interfaces contact FcpA (blue). **(c)** *L. biflexa* filament cross-section showing four seams. Besides K-A and B-C, additional seams at A-B (beneath the FlaA1-absent position) and I-J (beneath B0SMK8) are marked. **(d)** Superposition of FlaB protomers reveals continuous conformational variation. Left: 11 protomers from different protofilaments at one axial level, showing substantial conformational spread. Right: protomers from a single protofilament (chains A) show much less variation. This continuous flexibility, rather than discrete L/R state switching, characterises *Leptospira* flagellin cores. The continuous conformational range among protofilaments is similar in *L. interrogans* (not shown).

**Supplementary Video 1. *L. interrogans flaB1^-^* mutant displays *wild-type* swimming behaviour in liquid medium.**

Darkfield microscopy videos comparing swimming behaviour of *L. interrogans wt* (left) and *flaB1^-^* mutant (right) in liquid EMJH medium. Both strains display characteristic spirochete motility with similar swimming patterns, demonstrating that the *flaB1^-^* mutant assembles functional flagella capable of propulsion in low-viscosity environments. Scale bar: 10 µm. Frame rate: 12 fps; playback: real-time.

*(Attached separately as a standalone mp4 file)*

**Supplementary Table 1.** Cryo-EM data collection, processing and model refinement statistics.

|  | *L. biflexa* wt<br>pdb_000010LM<br>EMD-75270 | *L. interrogans* wt<br>pdb_000010LK<br>EMD-75268 | *L. interrogans*<br>*flaB1* mutant<br>pdb_000010LL<br>EMD-75269 |
| --- | --- | --- | --- |
| **Data collection & processing** |  |  |  |
| Microscope | Titan Krios G2 | Titan Krios G2 | Titan Krios G2 |
| Detector | Gatan K3 | Gatan K3 | Gatan K3 |
| Magnification | 81,000 | 81,000 | 81,000 |
| Voltage (kV) | 300 | 300 | 300 |
| Electron exposure (e<sup>-</sup>/Å<sup>2</sup>) | 25.7 | 61.5 | 61.5 |
| Defocus range (μm) | -1.5 to -2.6 | -1.5 to -2.6 | -1.5 to -2.6 |
| Pixel size (Å) | 1.068 | 1.068 | 1.068 |
| Symmetry imposed | C1 | C1 | C1 |
| Number of Micrographs | 1,985 | 1,081 | 1,094 |
| Initial particle images (n) | 688,969 | 239,996 | 226,523 |
| Final particle images (n) | 176,558 | 35,760 | 53,501 |
| Resolution (Å) @ FSC threshold 0.143 | 3.5 | 4.4 | 4.3 |
| estimated resolution after<br>density modification (Å) | 3.2 | 4.2 | 4.2 |
| **Model refinement** |  |  |  |
| Number of atoms |  |  |  |
| All | 248,112 | 248,049 | 253,413 |
| Protein | 248,082 | 247,995 | 253,359 |
| Water | 20 | 36 | 36 |
| Ligand (Ca<sup>2+</sup>) | 10 | 18 | 18 |
| Model to map correlation |  |  |  |
| CC(mask) | 0.76 | 0.69 | 0.66 |
| CC(volume) | 0.72 | 0.65 | 0.62 |
| CC(box) | 0.62 | 0.53 | 0.51 |
| RMSD |  |  |  |
| Bond lengths (Å) | 0.006 | 0.005 | 0.007 |
| Bond angles (°) | 1.11 | 1.12 | 1.14 |
| Ramachandran statistics |  |  |  |
| Favored (%) | 97.54 | 95.14 | 95.01 |
| Allowed (%) | 2.42 | 5.31 | 3.11 |
| Outliers (%) | 0.05 | 0.32 | 0.06 |
| Rama-Z score | 1.20 | -0.76 | -0.27 |
| Rotamer outliers (%) | 3.30 | 0.36 | 0.06 |
| C-beta outliers (%) | 0 | 0 | 0.01 |
| Clashscore | 7.26 | 6.67 | 8.95 |
| MolProbity overall score | 1.88 | 1.71 | 1.82 |

**Supplementary Table 2.** Crosslinking mass spectrometry analysis of purified filaments from *Leptospira biflexa*.

*(Attached separately as a standalone xlsx file)*

**Supplementary Table 3.** Label-free comparative proteomics of *Leptospira biflexa* filaments from *wt vs flaA2^-^* or *fcpA^-^*mutant strains.

*(Attached separately as a standalone xlsx file)*

**Supplementary Table 4.** Protein components (UniProt ID codes) constituting the endoflagellar filament in *Leptospira*. Marked in bold red are the proteins identified and modelled within the cryo-EM map for each *Leptospira* species. Orthologues are included for completeness.

| Protein | MW (kDa) | *L. biflexa* ID | *L. interrogans* ID | Copies/cell <sup>1</sup> | Predicted function |
| --- | --- | --- | --- | --- | --- |
| FlaB1 | 31 | B0SSZ5 † | **Q72R58** | 24,000 | Core flagellin |
| FlaB4 | 31 | **B0SQZ5** | Q72S55 † | 7,160 | Core flagellin |
| FlaB2 | 31 | B0SSZ4 | Q72R59 | 2,800 | Core flagellin |
| FlaB3 | 31 | B0SS86 | Q72S54 | 200 | Core flagellin |
| FcpA | 36 | **B0STJ8** | **Q72MM7** | 22,100 | Sheath (convex) |
| FcpB | 32 | **B0SR03** | **Q72RA0** | 9,260 | Sheath (convex) |
| FlaA1 | 36 | B0SKT4** | **Q72U74** | 5,590 | Sheath (concave) |
| FlaA2 | 27 | **B0SKT5** | **Q72U75** | 6,020 | Sheath (concave) |
| FlaAP | 28 | **B0SJC6** | **Q72SU9** | 870 | FlaA2-associated |
| B0STF2 | 25 | **B0STF2** | **Q72QN8** | 1,190 | FlaA2-associated |
| B0SPP1 | 24 | **B0SPP1** | **Q72NA0** | 300 | Lateral region |
| B0STN7 | 22 | **B0STN7** | Q75FY6 †<br>(replaced by<br>paralogue<br>**Q72RW3**) | 150 (Q75FY6)<br>500 (Q72RW3) | Lateral region |
| B0SMK8 | 25 | **B0SMK8**<br>(interspersed with<br>paralogue **B0SMT6**) | Q72Q50 †/<br>Q72U89 † | 1,100 | Lateral region |
| B0SKN8 | 25 | **B0SKN8** | Q72MH5 † | 340 | Lateral region |
| HEAT repeat | 33 | B0SNJ6 † | **Q72MR9** | 1,400 | Lateral region |

<sup>1</sup> Measured in *L. interrogans* by Schmidt et al [23]
† Not detected in the cryo-EM structures solved in this study.
\*\* Present in the MS data, yet systematically lost from the filament during sample preparation and absent from refined atomic model

**Supplementary Table 5.** X-ray data collection, processing and model refinement statistics.

| FlaA1 |  |
| --- | --- |
| **Data collection** |  |
| Beamline | Diamond Light Source I04 |
| Wavelength (Å) | 0.97951 |
| Space group | P2<sub>1</sub>2<sub>1</sub>2<sub>1</sub> |
| Cell dimensions |  |
| *a*, *b*, *c* (Å) | 81.4, 105.8, 108.7 |
| α, β, γ (°) | 90, 90, 90 |
| Resolution (Å) | 75.82–1.816 (1.848–1.816)* |
| Number unique reflections | 85228 (4238)* |
| R<sub>merge</sub> | 0.106 (5.769)* |
| R<sub>pim</sub> | 0.030 (1.596)* |
| Mean *I* / σ*I* | 47.5 (0.5)* |
| CC(1/2) | 0.998 (0.375)* |
| Completeness (%) | 100.0 (100.0)* |
| Redundancy | 13.8 (14.0)* |
| **Refinement** |  |
| Resolution (Å) | 75.82–1.82 (1.84–1.82)* |
| Number reflections used in refinement | 85179 (2640)* |
| Number reflections in the free set | 4230 (146)* |
| R<sub>work</sub> / R<sub>free</sub> | 0.184 / 0.215 |
| Number of atoms (non-H) | 5528 |
| Protein | 5073 |
| Ions | 38 (3 Ca<sup>2+</sup> / 35 SO<sub>4</sub><sup>2-</sup>) |
| Water | 417 |
| Standard coordinate uncertainty (Å) | 0.242 |
| Average *B*-factor | 50.15 |
| R.m.s. deviations |  |
| Bond lengths (Å) | 0.017 |
| Bond angles (°) | 1.32 |
| Ramachandran |  |
| Favored (%) | 96.85 |
| Allowed (%) | 3.15 |
| Outliers (%) | 0.00 |
| Rama-Z score | 0.47 |
| Clashscore | 2.92 |
| PDB ID | pdb_00009cyn |

\* Highest resolution shell

**Supplementary Table 6.** Orthologues/paralogues of *Leptospira* flagellar filament proteins. Proteins are grouped according to structural similarity identified by FoldSeek. Orthologous proteins (rows) are shown for 5 representative species. Paralogues, when identified, are listed within each Table cell. Proteins identified in *Leptospira* filaments are marked in bold. Low reliability hits are in italics (their structural similarities explain FoldSeek scores, but overall correspondence is not unambiguous as the other cases)

| Protein family | Orthologues (in five representative species) & paralogues (within each cell)<br>UniProt IDs |  |  |  |  |
| --- | --- | --- | --- | --- | --- |
|  | *Leptospira biflexa* | *Leptospira interrogans* | *Treponema pallidum* | *Borrelia burgdorferi* | *Brachyspira hyodysenteriae* |
| FcpA-like | **B0STJ8**<br>B0SS28 | **Q72MM7**<br>Q72NU5 | *O83457\*\** | ND‡ | ND‡ |
| FcpB-like | **B0SR03**<br>**B0SPP1**<br>**B0SKN8**<br>B0SSU6<br>- | **Q72RA0**<br>**Q72NA0**<br>Q72MH5<br>-<br>Q72UI5 | ND‡ | ND‡ | ND‡ |
| B0STN7-like | **B0STN7**<br>B0SR41 | Q75FY6<br>**Q72RW3** | ND‡ | ND‡ | ND‡ |
| B0SNJ6-like | B0SNJ6 | **Q72MR9** | ND‡ | ND‡ | ND‡ |
| FlaA-like | **B0SKT4**<br>(FlaA1)<br>**B0SKT5**<br>(FlaA2)<br>B0SPB9<br>B0SMM5 | **Q72U74** (FlaA1)<br>**Q72U75** (FlaA2)<br>Q72Q36<br>Q72MZ0<br>Q72UQ4 | P18193 (FlaA)<br>O83670<br>(FlaA2) | P70856 | A0A3B6W463<br>A0A3B6VR02<br>A0A3B6VWM6 |
| FlaAP-like<br>(associated to<br>one FlaA2<br>protofilament in<br>*Leptospira*) | **B0SJC6**<br>(FlaAP)<br>**B0SMK8**<br>**B0SMT6**<br>B0SQV9<br>B0SPD8<br>B0STI0<br>B0SMK9 | **Q72SU9**<br>(FlaAP)<br>Q72Q50<br>Q72U89<br>Q72VN4<br>Q72WC0<br>Q72TL5 | *P07643\*\** | ND‡ | A0A3B6VTA9 |
| B0STF2-like<br>(associated to<br>the other FlaA2,<br>opposite to<br>FlaAP ) | **B0STF2** | **Q72QN8** | O83478<br>(residues 60-169) | *O51431\*\** | *A0A3B6VQC5\*\** |

**\*\* low reliability prediction**
‡ ND not detected

**Supplementary Data 1.** Home-made bash script to obtain filament coordinates from Relion star files (processed cryo-EM micrographs data).

*(attached as a standalone plain text file)*

**Supplementary Data 2.** Home-made bash script to apply the Crenshaw method to calculate the filaments’ curvature (kappa) and torsion (tau) parameters from their coordinates in micrographs.

*(attached as a standalone plain text file)*

**Supplementary Data 3.** Home-made bash script to download AlphaFold-predicted proteins from entire proteomes from selected Spirochete representative species.

*(attached as a standalone plain text file)*

## References

1. Moustafa, M.A.M., S. Schlachter, and N. Parveen, Innovative Strategies to Study the Pathogenesis of Elusive Spirochetes and Difficulties Managing the Chronic Infections They Cause. Annu Rev Microbiol, 2024. 78(1): p. 337–360.

2. Worrall, L.J., D.D. Majewski, and N.C.J. Strynadka, Structural Insights into Type III Secretion Systems of the Bacterial Flagellum and Injectisome. Annu Rev Microbiol, 2023. 77: p. 669–698.

3. Charon, N.W., et al., The unique paradigm of spirochete motility and chemotaxis. Annu Rev Microbiol, 2012. 66: p. 349–70.

4. Wolgemuth, C.W., et al., The flagellar cytoskeleton of the spirochetes. J Mol Microbiol Biotechnol, 2006. 11(3-5): p. 221–7.

5. Wolgemuth, C.W., Flagellar motility of the pathogenic spirochetes. Semin Cell Dev Biol, 2015. 46: p. 104–12.

6. Costa, F., et al., Global Morbidity and Mortality of Leptospirosis: A Systematic Review. PLoS Negl Trop Dis, 2015. 9(9): p. e0003898.

7. Ko, A.I., C. Goarant, and M. Picardeau, Leptospira: the dawn of the molecular genetics era for an emerging zoonotic pathogen. Nat Rev Microbiol, 2009. 7(10): p. 736–47.

8. Picardeau, M., Virulence of the zoonotic agent of leptospirosis: still terra incognita? Nat Rev Microbiol, 2017. 15(5): p. 297–307.

9. Lambert, A., et al., FlaA proteins in Leptospira interrogans are essential for motility and virulence but are not required for formation of the flagellum sheath. Infect Immun, 2012. 80(6): p. 2019–25.

10. Wunder, E.A., Jr., et al., A novel flagellar sheath protein, FcpA, determines filament coiling, translational motility and virulence for the Leptospira spirochete. Mol Microbiol, 2016. 101(3): p. 457–70.

11. Carroll, B.L. and J. Liu, Structural Conservation and Adaptation of the Bacterial Flagella Motor. Biomolecules, 2020. 10(11).

12. San Martin, F., et al., Diving into the complexity of the spirochetal endoflagellum. Trends Microbiol, 2023. 31(3): p. 294–307.

13. Wunder, E.A., Jr., et al., FcpB Is a Surface Filament Protein of the Endoflagellum Required for the Motility of the Spirochete Leptospira. Front Cell Infect Microbiol, 2018. 8: p. 130.

14. Gibson, K.H., et al., An asymmetric sheath controls flagellar supercoiling and motility in the leptospira spirochete. Elife, 2020. 9.

15. Brady, M.R., et al., Structural basis of flagellar filament asymmetry and supercoil templating by Leptospira spirochete sheath proteins. bioRxiv, 2022: p. 2022.03.03.482903.

16. Punjani, A., H. Zhang, and D.J. Fleet, Non-uniform refinement: adaptive regularization improves single-particle cryo-EM reconstruction. Nat Methods, 2020. 17(12): p. 1214–1221.

17. Terwilliger, T.C., et al., Improvement of cryo-EM maps by density modification. Nat Methods, 2020. 17(9): p. 923–927.

18. Graziadei, A. and J. Rappsilber, Leveraging crosslinking mass spectrometry in structural and cell biology. Structure, 2022. 30(1): p. 37–54.

19. Maki-Yonekura, S., K. Yonekura, and K. Namba, Conformational change of flagellin for polymorphic supercoiling of the flagellar filament. Nat Struct Mol Biol, 2010. 17(4): p. 417–22.

20. Malmstrom, J., et al., Proteome-wide cellular protein concentrations of the human pathogen Leptospira interrogans. Nature, 2009. 460(7256): p. 762–5.

21. Boraston, A.B., et al., Carbohydrate-binding modules: fine-tuning polysaccharide recognition. The Biochemical journal, 2004. 382(Pt 3): p. 769–781.

22. Zhang, H., et al., dbCAN2: a meta server for automated carbohydrate-active enzyme annotation. Nucleic Acids Research, 2018. 46(W1): p. W95–W101.

23. Schmidt, A., et al., Absolute quantification of microbial proteomes at different states by directed mass spectrometry. Mol Syst Biol, 2011. 7: p. 510.

24. Kobe, B., et al., Turn up the HEAT. Structure, 1999. 7(5): p. R91–7.

25. Crenshaw, H.C., C.N. Ciampaglio, and M. McHenry, Analysis of the three-dimensional trajectories of organisms: estimates of velocity, curvature and torsion from positional information. J Exp Biol, 2000. 203(Pt 6): p. 961–82.

26. Hasegawa, K., I. Yamashita, and K. Namba, Quasi- and nonequivalence in the structure of bacterial flagellar filament. Biophys J, 1998. 74(1): p. 569–75.

27. Kreutzberger, M.A.B., et al., Flagellin outer domain dimerization modulates motility in pathogenic and soil bacteria from viscous environments. Nat Commun, 2022. 13(1): p. 1422.

28. Yonekura, K., S. Maki-Yonekura, and K. Namba, Complete atomic model of the bacterial flagellar filament by electron cryomicroscopy. Nature, 2003. 424(6949): p. 643–50.

29. Calladine, C.R., B.F. Luisi, and J.V. Pratap, A “mechanistic” explanation of the multiple helical forms adopted by bacterial flagellar filaments. J Mol Biol, 2013. 425(5): p. 914–28.

30. Kreutzberger, M.A.B., et al., Convergent evolution in the supercoiling of prokaryotic flagellar filaments. Cell, 2022. 185(19): p. 3487–3500 e14.

31. Zhang, X., et al., Differential Bending Stiffness of the Bacterial Flagellar Hook under Counterclockwise and Clockwise Rotations. Phys Rev Lett, 2023. 130(13): p. 138401.

32. Timoshenko, S., Analysis of Bi-Metal Thermostats J Opt Soc Am, 1925. 11(3): p. 233–255.

33. McCullough, B.R., et al., Cofilin increases the bending flexibility of actin filaments: implications for severing and cell mechanics. J Mol Biol, 2008. 381(3): p. 550–8.

34. Gan, L., S. Chen, and G.J. Jensen, Molecular organization of Gram-negative peptidoglycan. Proc Natl Acad Sci U S A, 2008. 105(48): p. 18953–7.

35. Johnson, R.C. and V.G. Harris, Differentiation of pathogenic and saprophytic letospires. I. Growth at low temperatures. J Bacteriol, 1967. 94(1): p. 27–31.

36. Murray, G.L., et al., Genome-wide transposon mutagenesis in pathogenic Leptospira species. Infect Immun, 2009. 77(2): p. 810–6.

37. Louvel, H. and M. Picardeau, Genetic manipulation of Leptospira biflexa. Curr Protoc Microbiol, 2007. **Chapter** 12: p. Unit 12E 4.

38. Wunder, E.A., Jr., Cell Monolayer Translocation Assay. Methods Mol Biol, 2020. 2134: p. 161–170.

39. Mastronarde, D.N., Automated electron microscope tomography using robust prediction of specimen movements. J Struct Biol, 2005. 152(1): p. 36–51.

40. Liebschner, D., et al., Macromolecular structure determination using X-rays, neutrons and electrons: recent developments in Phenix. Acta Crystallogr D Struct Biol, 2019. 75(Pt 10): p. 861–877.

41. Scheres, S.H., RELION: implementation of a Bayesian approach to cryo-EM structure determination. J Struct Biol, 2012. 180(3): p. 519–30.

42. Abramson, J., et al., Accurate structure prediction of biomolecular interactions with AlphaFold 3. Nature, 2024. 630(8016): p. 493–500.

43. Meng, E.C., et al., UCSF ChimeraX: Tools for structure building and analysis. Protein Sci, 2023. 32(11): p. e4792.

44. Casanal, A., B. Lohkamp, and P. Emsley, Current developments in Coot for macromolecular model building of Electron Cryo-microscopy and Crystallographic Data. Protein Sci, 2020. 29(4): p. 1069–1078.

45. Afonine, P.V., et al., New tools for the analysis and validation of cryo-EM maps and atomic models. Acta Crystallogr D Struct Biol, 2018. 74(Pt 9): p. 814–840.

46. Yamashita, K., et al., Cryo-EM single-particle structure refinement and map calculation using Servalcat. Acta Crystallogr D Struct Biol, 2021. 77(Pt 10): p. 1282–1291.

47. Nicholls, R.A., et al., Conformation-independent structural comparison of macromolecules with ProSMART. Acta Crystallogr D Biol Crystallogr, 2014. 70(Pt 9): p. 2487–99.

48. Williams, C.J., et al., MolProbity: More and better reference data for improved all-atom structure validation. Protein Sci, 2018. 27(1): p. 293–315.

49. Chojnowski, G., Sequence-assignment validation in cryo-EM models with checkMySequence. Acta Crystallogr D Struct Biol, 2022. 78(Pt 7): p. 806–816.

50. Gil, M., et al., New substrates and interactors of the mycobacterial Serine/Threonine protein kinase PknG identified by a tailored interactomic approach. J Proteomics, 2019. 192: p. 321–333.

51. Carvalho, P.C., et al., Integrated analysis of shotgun proteomic data with PatternLab for proteomics 4.0. Nat Protoc, 2016. 11(1): p. 102–17.

52. Rey, M., et al., eXL-MS: An Enhanced Cross-Linking Mass Spectrometry Workflow To Study Protein Complexes. Anal Chem, 2018. 90(18): p. 10707–10714.

53. Sarpe, V., et al., High Sensitivity Crosslink Detection Coupled With Integrative Structure Modeling in the Mass Spec Studio. Mol Cell Proteomics, 2016. 15(9): p. 3071–80.

54. Combe, C.W., et al., xiVIEW: Visualisation of Crosslinking Mass Spectrometry Data. J Mol Biol, 2024. 436(17): p. 168656.

55. Correa, A., et al., Generation of a vector suite for protein solubility screening. Front Microbiol, 2014. 5: p. 67.

56. Kabsch, W., Xds. Acta Crystallogr D Biol Crystallogr, 2010. 66(Pt 2): p. 125–32.

57. Evans, P.R. and G.N. Murshudov, How good are my data and what is the resolution? Acta Crystallographica Section D: Biological Crystallography, 2013. 69(Pt 7): p. 1204–1214.

58. McCoy, A.J., et al., Phaser crystallographic software. Journal of Applied Crystallography, 2007. 40(4): p. 658–674.

59. Afonine, P.V., et al., Towards automated crystallographic structure refinement with phenix.refine. Acta Crystallogr D Biol Crystallogr, 2012. 68(Pt 4): p. 352–67.

60. van Kempen, M., et al., Fast and accurate protein structure search with Foldseek. Nat Biotechnol, 2024. 42(2): p. 243–246.

61. Sievers, F. and D.G. Higgins, Clustal omega. Curr Protoc Bioinformatics, 2014. 48: p. 3 13 1–3 13 16.

62. Perez-Riverol, Y., et al., The PRIDE database at 20 years: 2025 update. Nucleic Acids Res, 2025. 53(D1): p. D543–D553.

63. Carvalho, P.C., et al., Analyzing marginal cases in differential shotgun proteomics. Bioinformatics, 2011. 27(2): p. 275–6.

64. Kurniyati, K., et al., A novel glycan modifies the flagellar filament proteins of the oral bacterium Treponema denticola. Mol Microbiol, 2017. 103(1): p. 67–85.

