## Supplementary Table 2 for "Core–sheath coupling controls flagellar curvature and motility in *Leptospira*"

**Table S2. Cross-linking mass spectrometry analysis of purified filaments from Leptospira biflexa.**

PepSeq1 and PepSeq2 = amino acid sequence of peptides 1 and 2, respectively;
PepPos1 = position of peptide 1 in protein 1;
PepPos2 = position of peptide 2 in protein 2;
LinkPos1 = position of crosslinked residue for peptide 1;
LinkPos2 = position of crosslinked residue for peptide 2;
Protein1 and Protein2 = UniProt entry of cross-linked proteins 1 and 2, respectively.

| PepSeq1 | PepSeq2 | PepPos1 | PepPos2 | LinkPos1 | LinkPos2 | Protein1 | Protein2 |
|---|---|---|---|---|---|---|---|
| NFRGEDPEGKPSELNAPEGKEGFR | DYSKESAESIKK | 47 | 85 | 10 | 4 | B0SJC6 | B0STN7 |
| ELEPIVKNFRGEDPEGKPSELNAPEGK | VYELDSNNMVFMKYDQER | 40 | 330 | 7 | 13 | B0SJC6 | B0SJC6 |
| ELEPIVKNFR | YLEAKR | 40 | 74 | 7 | 5 | B0SJC6 | B0SJC6 |
| KALNVPPKPAQGSTTTPQQ | TEASKDETGASVEGQPVK | 349 | 270 | 1 | 5 | B0SJC6 | B0SJC6 |
| ESVSSKSYQK | HQKLQPSHR | 21 | 219 | 6 | 3 | B0SJC6 | B0SJC6 |
| IDGVTYDFTK | NPYIKYDNR | 288 | 298 | 5 | 5 | B0SJC6 | B0SJC6 |
| VYELDSNNMVFMKYDQER | ELEPIVKNFR | 330 | 40 | 13 | 7 | B0SJC6 | B0SJC6 |
| TEASKDETGASVEGQPVKIDGVTYDFT | TEASKDETGASVEGQPVKIDGVTYDFTK | 270 | 270 | 5 | 18 | B0SJC6 | B0SJC6 |
| NFRGEDPEGKPSELNAPEGKEGFRMK | NFRGEDPEGKPSELNAPEGKEGFRMKK | 47 | 47 | 10 | 26 | B0SJC6 | B0SJC6 |
| NFRGEDPEGKPSELNAPEGKEGFR | NFRGEDPEGKPSELNAPEGKEGFR | 47 | 47 | 10 | 20 | B0SJC6 | B0SJC6 |
| ELEPIVKNFRGEDPEGKPSELNAPEGK | ELEPIVKNFRGEDPEGKPSELNAPEGK | 40 | 40 | 7 | 17 | B0SJC6 | B0SJC6 |
| NFRGEDPEGKPSELNAPEGKEGFR | NFRGEDPEGKPSELNAPEGKEGFR | 47 | 47 | 10 | 20 | B0SJC6 | B0SJC6 |
| GEDPEGKPSELNAPEGKEGFR | GEDPEGKPSELNAPEGKEGFR | 50 | 50 | 7 | 17 | B0SJC6 | B0SJC6 |
| IDGVTYDFTKNPYIKYDNR | IDGVTYDFTKNPYIKYDNR | 288 | 288 | 15 | 10 | B0SJC6 | B0SJC6 |
| ALNVPPKPAQGSTTTPQQ | ALNVPPKPAQGSTTTPQQ | 350 | 350 | 7 | 7 | B0SJC6 | B0SJC6 |
| ALKVER | NLEKHQK | 209 | 215 | 3 | 4 | B0SJC6 | B0SJC6 |
| TEASKDETGASVEGQPVK | IDGVTYDFTK | 270 | 288 | 5 | 5 | B0SJC6 | B0SJC6 |
| TEASKDETGASVEGQPVK | TEASKDETGASVEGQPVK | 270 | 270 | 5 | 11 | B0SJC6 | B0SJC6 |
| NFRGEDPEGKPSELNAPEGK | NFRGEDPEGKPSELNAPEGK | 47 | 47 | 10 | 12 | B0SJC6 | B0SJC6 |
| NLEKHQKLQPSHR | NLEKHQKLQPSHR | 215 | 215 | 4 | 7 | B0SJC6 | B0SJC6 |
| HQKLQPSHR | HQKLQPSHR | 219 | 219 | 3 | 7 | B0SJC6 | B0SJC6 |
| LKYPYDNYYLNNSQAK | LKYPYDNYYLNNSQAK | 254 | 254 | 2 | 13 | B0SJC6 | B0SJC6 |
| IDGVTYDFTKNPYIKYDNR | IDGVTYDFTKNPYIKYDNR | 288 | 288 | 10 | 15 | B0SJC6 | B0SJC6 |
| VYELDSNNMVFMKYDQERKK | VYELDSNNMVFMKYDQERKK | 330 | 330 | 13 | 19 | B0SJC6 | B0SJC6 |
| KKALNVPPKPAQGSTTTPQQ | KKALNVPPKPAQGSTTTPQQ | 348 | 348 | 1 | 9 | B0SJC6 | B0SJC6 |
| ESVSSKSYQKR | ESVSSKSYQKR | 21 | 21 | 6 | 10 | B0SJC6 | B0SJC6 |
| EAHYSYNKYDIEK | EAHYSYNKYDIEK | 178 | 178 | 5 | 8 | B0SJC6 | B0SJC6 |
| NMELGYKHLGLAK | NMELGYKHLGLAK | 191 | 191 | 7 | 13 | B0SJC6 | B0SJC6 |
| AVNIYKLK | AVNIYKLK | 248 | 248 | 6 | 8 | B0SJC6 | B0SJC6 |
| NPYIKYDNR | NPYIKYDNR | 298 | 298 | 5 | 5 | B0SJC6 | B0SJC6 |
| VYELDSNNMVFMKYDQER | VYELDSNNMVFMKYDQER | 330 | 330 | 6 | 13 | B0SJC6 | B0SJC6 |
| TGASVEGQPVKIDGVTYDFTK | TGASVEGQPVKIDGVTYDFTK | 277 | 277 | 4 | 11 | B0SJC6 | B0SJC6 |
| VYELDSNNMVFMKYDQERKK | VYELDSNNMVFMKYDQERKK | 330 | 330 | 6 | 19 | B0SJC6 | B0SJC6 |
| ESQKQIDLQR | KNLDFDYFK | 32 | 67 | 4 | 1 | B0SKN8 | B0SMK8 |
| EKPASATLK | NYEKVR | 21 | 230 | 2 | 4 | B0SKN8 | B0SMK8 |
| KIAKQTAFK | NYEKVR | 240 | 230 | 1 | 4 | B0SKN8 | B0SMK8 |
| IAKQTAFK | NYEKVR | 241 | 230 | 3 | 4 | B0SKN8 | B0SMK8 |
| EKPASATLK | VRDTLINFIENKR | 21 | 234 | 2 | 12 | B0SKN8 | B0SMK8 |
| EKPASATLK | TVVPPGNPDAKPLDLLEQHDDNYGFITS | 21 | 250 | 2 | 11 | B0SKN8 | B0SMK8 |
| EKPASATLK | KNLDFDYFK | 21 | 67 | 2 | 1 | B0SKN8 | B0SMK8 |
| EKPASATLK | DTLINFIENKR | 21 | 236 | 2 | 10 | B0SKN8 | B0SMK8 |
| SEGTEKDEKGEAR | EKPASATLK | 166 | 21 | 9 | 2 | B0SKN8 | B0SKN8 |
| SEGTEKDEK | EKPASATLK | 166 | 21 | 6 | 2 | B0SKN8 | B0SKN8 |
| EKPASATLK | IAKQTAFK | 21 | 241 | 2 | 3 | B0SKN8 | B0SKN8 |
| SVELDQKQMITK | EKPASATLK | 251 | 21 | 7 | 2 | B0SKN8 | B0SKN8 |
| KLQDLGVDKTAAQYLAQTPQEHK | EKPASATLK | 66 | 21 | 1 | 2 | B0SKN8 | B0SKN8 |
| KLQDLGVDK | EKPASATLK | 66 | 21 | 1 | 2 | B0SKN8 | B0SKN8 |
| ESQKQIDLQR | EKPASATLK | 32 | 21 | 4 | 2 | B0SKN8 | B0SKN8 |
| KNGFGDNEIDTLHANIGNLR | EKPASATLK | 42 | 21 | 1 | 2 | B0SKN8 | B0SKN8 |
| SVELDQKQMITK | KLQDLGVDK | 251 | 66 | 7 | 1 | B0SKN8 | B0SKN8 |
| KLQDLGVDKTAAQYLAQTPQEHK | KDKDGKPYLEIK | 66 | 93 | 1 | 1 | B0SKN8 | B0SKN8 |
| LQDLGVDKTAAQYLAQTPQEHK | KDKDGKPYLEIK | 67 | 93 | 8 | 1 | B0SKN8 | B0SKN8 |
| EKPASATLK | EKPASATLK | 21 | 21 | 2 | 7 | B0SKN8 | B0SKN8 |
| KNGFGDNEIDTLHANIGNLRK | KNGFGDNEIDTLHANIGNLRK | 42 | 42 | 1 | 11 | B0SKN8 | B0SKN8 |
| LQDLGVDKTAAQYLAQTPQEHK | LQDLGVDKTAAQYLAQTPQEHK | 67 | 67 | 8 | 17 | B0SKN8 | B0SKN8 |
| KNGFGDNEIDTLHANIGNLR | KNGFGDNEIDTLHANIGNLR | 42 | 42 | 1 | 11 | B0SKN8 | B0SKN8 |
| KNGFGDNEIDTLHANIGNLR | ESQKQIDLQR | 42 | 31 | 1 | 5 | B0SKN8 | B0SKN8 |
| KLQDLGVDKTAAQYLAQTPQEHK | KLQDLGVDKTAAQYLAQTPQEHK | 66 | 66 | 1 | 9 | B0SKN8 | B0SKN8 |
| LQDLGVDKTAAQYLAQTPQEHK | LQDLGVDKTAAQYLAQTPQEHK | 67 | 67 | 8 | 17 | B0SKN8 | B0SKN8 |
| SVELDQKQMITK | SVELDQKQMITK | 251 | 251 | 1 | 7 | B0SKN8 | B0SKN8 |
| LRSVELDQKQMITK | LRSVELDQKQMITK | 249 | 249 | 3 | 9 | B0SKN8 | B0SKN8 |
| TAAQYLAQTPQEHKELYK | TAAQYLAQTPQEHKELYK | 75 | 75 | 14 | 14 | B0SKN8 | B0SKN8 |
| EKPASATLKER | EKPASATLKER | 21 | 21 | 2 | 9 | B0SKN8 | B0SKN8 |
| QKHIMMSYK | QKHIMMSYK | 227 | 227 | 2 | 7 | B0SKN8 | B0SKN8 |
| KIAKQTAFK | KIAKQTAFK | 240 | 240 | 1 | 4 | B0SKN8 | B0SKN8 |
| IAKQTAFKLR | IAKQTAFKLR | 241 | 241 | 3 | 8 | B0SKN8 | B0SKN8 |
| IKKLQDLGVDK | IKKLQDLGVDK | 64 | 64 | 2 | 3 | B0SKN8 | B0SKN8 |
| KLQDLGVDKTAAQYLAQTPQEHK | KLQDLGVDKTAAQYLAQTPQEHK | 66 | 66 | 1 | 9 | B0SKN8 | B0SKN8 |
| TAAQYLAQTPQEHKELYKK | TAAQYLAQTPQEHKELYKK | 75 | 75 | 14 | 18 | B0SKN8 | B0SKN8 |
| KDKDGKPYLEIK | KDKDGKPYLEIK | 93 | 93 | 1 | 3 | B0SKN8 | B0SKN8 |
| ADGTIHVKEMR | ADGTIHVKEMR | 147 | 147 | 4 | 8 | B0SKN8 | B0SKN8 |
| SNDSNAKALLR | FKSLHIQGK | 139 | 68 | 7 | 2 | B0SMK8 | B0SMT6 |
| SNDSNAKALLR | YIEDAR | 139 | 121 | 7 | 1 | B0SMK8 | B0SMK8 |
| KLDLLLEANAQIKETEGAR | KLDLLLEANAQIKETEGAR | 279 | 279 | 1 | 13 | B0SMK8 | B0SMK8 |
| LDLLLEANAQIKETEGARR | LDLLLEANAQIKETEGARR | 280 | 280 | 12 | 14 | B0SMK8 | B0SMK8 |
| KNLDFDYFKLLK | KNLDFDYFKLLK | 67 | 67 | 1 | 9 | B0SMK8 | B0SMK8 |
| LLKAANQSDFSGNMWYLQSNYVYGFR | LLKAANQSDFSGNMWYLQSNYVYGFR | 76 | 76 | 3 | 8 | B0SMK8 | B0SMK8 |
| LTLYKEGILTLR | LTLYKEGILTLR | 177 | 177 | 2 | 5 | B0SMK8 | B0SMK8 |
| LDLLLEANAQIKETEGAR | LDLLLEANAQIKETEGAR | 280 | 280 | 12 | 14 | B0SMK8 | B0SMK8 |
| ADEKNESVSQSDATK | VKFLSFEK | 20 | 72 | 4 | 2 | B0SPP1 | B0SPP1 |
| IGKEVDDYHAMINEK | ADEKNESVSQSDATK | 57 | 20 | 3 | 4 | B0SPP1 | B0SPP1 |
| YIQAKGNLETK | FLSFEKK | 211 | 74 | 5 | 6 | B0SPP1 | B0SPP1 |
| LNPDGSIALIPYNAKQQQK | ADEKNESVSQSDATK | 35 | 20 | 15 | 4 | B0SPP1 | B0SPP1 |
| ADEKNESVSQSDATK | NNTTGGVKGLVLSVR | 20 | 153 | 4 | 8 | B0SPP1 | B0SPP1 |
| IGKEVDDYHAMINEK | NNTTGGVKGLVLSVR | 57 | 153 | 3 | 8 | B0SPP1 | B0SPP1 |
| ADEKNESVSQSDATKLNPDGSIALIPYN | ADEKNESVSQSDATKLNPDGSIALIPYNA | 20 | 20 | 4 | 15 | B0SPP1 | B0SPP1 |
| ADEKNESVSQSDATK | ADEKNESVSQSDATK | 20 | 20 | 4 | 11 | B0SPP1 | B0SPP1 |
| SLTEKGKDPVTTYR | SLTEKGKDPVTTYR | 136 | 136 | 5 | 7 | B0SPP1 | B0SPP1 |
| ELKNNTTGGVKGLVLSVR | ELKNNTTGGVKGLVLSVR | 150 | 150 | 3 | 11 | B0SPP1 | B0SPP1 |
| ALPFTLFGETGPSKEK | KLTATVPGFIPQSTR | 184 | 174 | 14 | 1 | B0STF2 | B0SKT5 |
| ALPFTLFGETGPSKEK | GLIKDVFDEK | 184 | 79 | 14 | 4 | B0STF2 | B0SKT5 |
| ALPFTLFGETGPSKEK | TVPEDGGDKIEK | 184 | 89 | 14 | 9 | B0STF2 | B0SKT5 |
| ALPFTLFGETGPSKEK | DVFDEKTVPEDGGDKIEK | 184 | 83 | 14 | 6 | B0STF2 | B0SKT5 |
| ALPFTLFGETGPSKEK | QVIYNYVGQTADEK | 184 | 200 | 14 | 10 | B0STF2 | B0STF2 |
| TLYFSNMPTPFTAKEMVGEETYAK | TLYFSNMPTPFTAKEMVGEETYAK | 124 | 124 | 5 | 15 | B0STF2 | B0STF2 |
| EKQVIYNYVGQTADEKK | EKQVIYNYVGQTADEKK | 198 | 198 | 2 | 16 | B0STF2 | B0STF2 |
| KPTNSLLFR | VAILVFDTK | 264 | 255 | 1 | 8 | B0STF2 | B0STF2 |
| KFIDISDLK | KPTNSLLFR | 273 | 264 | 1 | 1 | B0STF2 | B0STF2 |
| HADTGKGEFLDVQVEIESR | QVHETLPKHTYTIYNNK | 62 | 216 | 6 | 8 | B0STF2 | B0STF2 |
| DNRDLYPLSMYDAR | KFIDISDLK | 37 | 273 | 9 | 1 | B0STF2 | B0STF2 |
| KFIDISDLK | LKNVTFVR | 273 | 53 | 1 | 2 | B0STF2 | B0STF2 |
| QESGNPQGTQVSAASSKDNR | QESGNPQGTQVSAASSKDNR | 20 | 20 | 3 | 17 | B0STF2 | B0STF2 |
| NNAKALPFTLFGETGPSK | NNAKALPFTLFGETGPSK | 180 | 180 | 5 | 17 | B0STF2 | B0STF2 |
| EKQVIYNYVGQTADEK | EKQVIYNYVGQTADEK | 198 | 198 | 2 | 12 | B0STF2 | B0STF2 |
| HTYTIYNNK | RQVHETLPK | 224 | 215 | 2 | 6 | B0STF2 | B0STF2 |
| KKEEMEKR | KKEEMEKR | 149 | 149 | 1 | 2 | B0STF2 | B0STF2 |
| ALPFTLFGETGPSKEKQVIYNYVGQTA | ALPFTLFGETGPSKEKQVIYNYVGQTADE | 184 | 184 | 14 | 16 | B0STF2 | B0STF2 |
| QVIYNYVGQTADEKKR | QVIYNYVGQTADEKKR | 200 | 200 | 14 | 15 | B0STF2 | B0STF2 |
| YKTTITSHHYTQYRPNFLSFNK | YKTTITSHHYTQYRPNFLSFNK | 233 | 233 | 2 | 7 | B0STF2 | B0STF2 |
| VAILVFDTKKPTNSLLFR | VAILVFDTKKPTNSLLFR | 255 | 255 | 9 | 10 | B0STF2 | B0STF2 |
| KFIDISDLK | KFIDISDLK | 273 | 273 | 1 | 6 | B0STF2 | B0STF2 |
| KADPEKDER | KADPEKDER | 115 | 115 | 1 | 6 | B0STF2 | B0STF2 |
| ALPFTLFGETGPSKEK | ALPFTLFGETGPSKEK | 184 | 184 | 13 | 14 | B0STF2 | B0STF2 |
| QVIYNYVGQTADEKK | QVIYNYVGQTADEKK | 200 | 200 | 10 | 14 | B0STF2 | B0STF2 |
| QVHETLPKHTYTIYNNK | QVHETLPKHTYTIYNNK | 216 | 216 | 5 | 8 | B0STF2 | B0STF2 |
| HTYTIYNNKYK | HTYTIYNNKYK | 224 | 224 | 2 | 9 | B0STF2 | B0STF2 |
| QVIYNYVGQTADEK | QVIYNYVGQTADEK | 200 | 200 | 4 | 10 | B0STF2 | B0STF2 |
| DNRDLYPLSMYDAR | DNRDLYPLSMYDAR | 37 | 37 | 6 | 9 | B0STF2 | B0STF2 |
| AKLEYNLIKTLK | ESAESIKK | 67 | 89 | 2 | 7 | B0STN7 | B0STN7 |
| KIDATNIQYER | DYSKESAESIK | 96 | 85 | 1 | 4 | B0STN7 | B0STN7 |
| QSHLVFSGKDKFEITIPKP | QSHLVFSGKDKFEITIPKP | 157 | 157 | 9 | 11 | B0STN7 | B0STN7 |
| AKLEYNLIKTLKK | AKLEYNLIKTLKK | 67 | 67 | 2 | 9 | B0STN7 | B0STN7 |
| LEYNLIKTLKK | LEYNLIKTLKK | 69 | 69 | 7 | 10 | B0STN7 | B0STN7 |
| DYSKESAESIKK | DYSKESAESIKK | 85 | 85 | 4 | 11 | B0STN7 | B0STN7 |
| ESAESIKKIDATNIQYER | ESAESIKKIDATNIQYER | 89 | 89 | 7 | 8 | B0STN7 | B0STN7 |
| KIDATNIQYER | KIDATNIQYER | 96 | 96 | 1 | 5 | B0STN7 | B0STN7 |
| QDTKPQSTSTDVTQENHDAK | QDTKPQSTSTDVTQENHDAK | 24 | 24 | 1 | 4 | B0STN7 | B0STN7 |
| QSHLVFSGKDK | QSHLVFSGKDK | 157 | 157 | 2 | 9 | B0STN7 | B0STN7 |
| LALSNYQSGVNTIMKNR | IAKQTAFK | 102 | 241 | 15 | 3 | B0STJ8 | B0SKN8 |
| LALSNYQSGVNTIMKNR | QTAFKLR | 102 | 244 | 15 | 5 | B0STJ8 | B0SKN8 |
| QDALNYFSKAINHLDEIKNPDLR | SNEDLKEAR | 162 | 215 | 9 | 6 | B0STJ8 | B0SMK8 |
| EKMK | GKDPVTTYR | 156 | 141 | 2 | 2 | B0STJ8 | B0SPP1 |
| AKQDALNYFSK | GKDPVTTYR | 160 | 141 | 2 | 2 | B0STJ8 | B0SPP1 |
| TKGPDDMLFKYR | KSLTEK | 250 | 135 | 2 | 1 | B0STJ8 | B0SPP1 |
| GPDDMLFKYR | KSLTEK | 252 | 135 | 8 | 1 | B0STJ8 | B0SPP1 |
| NPDFKGYNK | GKDPVTTYR | 75 | 141 | 5 | 2 | B0STJ8 | B0SPP1 |
| AINHLDEIKNPDLR | KADPEKDER | 171 | 115 | 9 | 1 | B0STJ8 | B0STF2 |
| TKGPDDMLFKYR | KFIDISDLK | 250 | 273 | 2 | 1 | B0STJ8 | B0STF2 |
| GPDDMLFKYR | KFIDISDLK | 252 | 273 | 8 | 1 | B0STJ8 | B0STF2 |
| AKQDALNYFSKAINHLDEIKNPDLR | SSKNLGAK | 160 | 120 | 2 | 3 | B0STJ8 | B0SR03 |
| QDALNYFSKAINHLDEIKNPDLR | SSKNLGAK | 162 | 120 | 9 | 3 | B0STJ8 | B0SR03 |
| DQVDELLKGELVPENDDAELTEDQKK | KMELFFEGSNNADPDPR | 36 | 130 | 8 | 1 | B0STJ8 | B0SR03 |
| ERPEFKR | KQFYFK | 185 | 235 | 6 | 1 | B0STJ8 | B0SR03 |
| TKGPDDMLFKYR | NLGAKFKK | 250 | 123 | 2 | 5 | B0STJ8 | B0SR03 |
| TKGPDDMLFKYR | GSLKY | 250 | 273 | 2 | 4 | B0STJ8 | B0SR03 |
| GPDDMLFKYR | NLGAKFK | 252 | 123 | 8 | 5 | B0STJ8 | B0SR03 |
| GPDDMLFKYR | GSLKY | 252 | 273 | 8 | 4 | B0STJ8 | B0SR03 |
| YGKDSPEYKHIVNVINRDEVISVAQ | YILSNVENTKSNPIR | 276 | 216 | 3 | 10 | B0STJ8 | B0SR03 |
| DSPEYKHIVNVINRDEVISVAQ | YILSNVENTKSNPIR | 279 | 216 | 6 | 10 | B0STJ8 | B0SR03 |
| KEIMEQESLWKNPDFK | IFDYNDRDSNKHYK | 64 | 252 | 1 | 11 | B0STJ8 | B0SR03 |
| KEIMEQESLWKNPDFK | KNVEALK | 64 | 266 | 1 | 1 | B0STJ8 | B0SR03 |
| NPDFKGYNK | SSKNLGAK | 75 | 120 | 5 | 3 | B0STJ8 | B0SR03 |
| GYNKTFQELHQLSK | SSKNLGAK | 80 | 120 | 4 | 3 | B0STJ8 | B0SR03 |
| TFQELHQLSKTFANNQFR | KMELFFEGSNNADPDPR | 84 | 130 | 10 | 1 | B0STJ8 | B0SR03 |
| TFQELHQLSKTFANNQFR | YILSNVENTKSNPIR | 84 | 216 | 10 | 10 | B0STJ8 | B0SR03 |
| KEEAEKK | KNVEALK | 126 | 266 | 1 | 1 | B0STJ8 | B0SR03 |
| KEEAEK | NLGAKFK | 126 | 123 | 1 | 5 | B0STJ8 | B0SR03 |
| DSPEYKHIVNVINR | YILSNVENTKSNPIR | 279 | 216 | 6 | 10 | B0STJ8 | B0SR03 |
| KEEAEK | IFDYNDRDSNKHYK | 126 | 252 | 1 | 11 | B0STJ8 | B0SR03 |
| EKMK | KNVEALK | 156 | 266 | 2 | 1 | B0STJ8 | B0SR03 |
| KEEAEK | KNVEALK | 126 | 266 | 1 | 1 | B0STJ8 | B0SR03 |
| KEEAEKK | NVEALKGSLKY | 126 | 267 | 1 | 6 | B0STJ8 | B0SR03 |
| KEEAEKK | NVEALKGSLK | 126 | 267 | 1 | 6 | B0STJ8 | B0SR03 |
| KEEAEK | NVEALKGSLK | 126 | 267 | 1 | 6 | B0STJ8 | B0SR03 |
| YGKDSPEYK | SMADTEKELDDNISEVNKR | 276 | 40 | 3 | 7 | B0STJ8 | B0SR03 |
| DQVDELLKGELVPENDDAELTEDQK | KMELFFEGSNNADPDPR | 36 | 130 | 8 | 1 | B0STJ8 | B0SR03 |
| TKGPDDMLFKYR | NLGAKFK | 250 | 123 | 10 | 5 | B0STJ8 | B0SR03 |
| GPDDMLFKYR | NLGAKFKK | 252 | 123 | 8 | 5 | B0STJ8 | B0SR03 |
| AKQDALNYFSK | NPDFKGYNK | 160 | 75 | 2 | 5 | B0STJ8 | B0STJ8 |
| AINHLDEIKNPDLR | KEEAEK | 171 | 126 | 9 | 1 | B0STJ8 | B0STJ8 |
| YLASAYSFEENMIKK | GYNKTFQELHQLSK | 235 | 80 | 14 | 4 | B0STJ8 | B0STJ8 |
| KEIMEQESLWKNPDFK | ATELKYGK | 64 | 271 | 1 | 5 | B0STJ8 | B0STJ8 |
| TFQELHQLSKTFANNQFR | ATELKYGK | 84 | 271 | 10 | 5 | B0STJ8 | B0STJ8 |
| TFANNQFR | ATELKYGK | 94 | 271 | 1 | 5 | B0STJ8 | B0STJ8 |
| EYPAHKYLASAYSFEENMIKK | ATELKYGK | 229 | 271 | 6 | 5 | B0STJ8 | B0STJ8 |
| ATELKYGK | KNVHLLR | 271 | 264 | 5 | 1 | B0STJ8 | B0STJ8 |
| GYNKTFQELHQLSK | AINHLDEIKNPDLR | 80 | 171 | 4 | 9 | B0STJ8 | B0STJ8 |
| EYPAHKYLASAYSFEENMIK | ATELKYGK | 229 | 271 | 6 | 5 | B0STJ8 | B0STJ8 |
| DQVDELLKGELVPENDDAELTEDQKKI | DQVDELLKGELVPENDDAELTEDQKKKI | 36 | 36 | 8 | 27 | B0STJ8 | B0STJ8 |
| DQVDELLKGELVPENDDAELTEDQKKI | DQVDELLKGELVPENDDAELTEDQKKK | 36 | 36 | 8 | 27 | B0STJ8 | B0STJ8 |
| DQVDELLKGELVPENDDAELTEDQKKI | DQVDELLKGELVPENDDAELTEDQKK | 36 | 36 | 8 | 27 | B0STJ8 | B0STJ8 |
| DQVDELLKGELVPENDDAELTEDQK | DQVDELLKGELVPENDDAELTEDQK | 36 | 36 | 8 | 27 | B0STJ8 | B0STJ8 |
| KRLDEKWYWQK | KRLDEKWYWQK | 132 | 132 | 1 | 6 | B0STJ8 | B0STJ8 |
| DQVDELLKGELVPENDDAELTEDQKKI | DQVDELLKGELVPENDDAELTEDQKKK | 36 | 36 | 8 | 25 | B0STJ8 | B0STJ8 |
| KEIMEQESLWKNPDFKGYNK | KEIMEQESLWKNPDFKGYNK | 64 | 64 | 1 | 16 | B0STJ8 | B0STJ8 |
| GYNKTFQELHQLSKTFANNQFR | GYNKTFQELHQLSKTFANNQFR | 80 | 80 | 4 | 14 | B0STJ8 | B0STJ8 |
| GYNKTFQELHQLSK | GYNKTFQELHQLSK | 80 | 80 | 4 | 13 | B0STJ8 | B0STJ8 |
| YLASAYSFEENMIKK | YLASAYSFEENMIKK | 235 | 235 | 4 | 14 | B0STJ8 | B0STJ8 |
| TFQELHQLSKTFANNQFR | TFQELHQLSKTFANNQFR | 84 | 84 | 9 | 10 | B0STJ8 | B0STJ8 |
| LALSNYQSGVNTIMKNR | LALSNYQSGVNTIMKNR | 102 | 102 | 4 | 15 | B0STJ8 | B0STJ8 |
| DQVDELLKGELVPENDDAELTEDQKK | DQVDELLKGELVPENDDAELTEDQKK | 36 | 36 | 8 | 25 | B0STJ8 | B0STJ8 |
| AKQDALNYFSKAINHLDEIKNPDLR | AKQDALNYFSKAINHLDEIKNPDLR | 160 | 160 | 2 | 11 | B0STJ8 | B0STJ8 |
| AKQDALNYFSK | AKQDALNYFSK | 160 | 160 | 2 | 10 | B0STJ8 | B0STJ8 |
| QDALNYFSKAINHLDEIKNPDLR | QDALNYFSKAINHLDEIKNPDLR | 162 | 162 | 9 | 18 | B0STJ8 | B0STJ8 |
| EYPAHKYLASAYSFEENMIK | EYPAHKYLASAYSFEENMIK | 229 | 229 | 6 | 10 | B0STJ8 | B0STJ8 |
| KTKGPDDMLFK | KTKGPDDMLFK | 249 | 249 | 1 | 3 | B0STJ8 | B0STJ8 |
| TKGPDDMLFKYR | TKGPDDMLFKYR | 250 | 250 | 2 | 10 | B0STJ8 | B0STJ8 |
| GPDDMLFKYRYKK | GPDDMLFKYRYKK | 252 | 252 | 8 | 12 | B0STJ8 | B0STJ8 |
| YKKNVHLLR | YKKNVHLLR | 262 | 262 | 2 | 3 | B0STJ8 | B0STJ8 |
| ATELKYGKDSPEYK | ATELKYGKDSPEYK | 271 | 271 | 5 | 8 | B0STJ8 | B0STJ8 |
| YGKDSPEYKHIVNVINRDEVISVAQ | YGKDSPEYKHIVNVINRDEVISVAQ | 276 | 276 | 3 | 9 | B0STJ8 | B0STJ8 |
| KKEIMEQESLWK | KKEIMEQESLWK | 63 | 63 | 1 | 2 | B0STJ8 | B0STJ8 |
| KEIMEQESLWKNPDFK | KEIMEQESLWKNPDFK | 64 | 64 | 1 | 8 | B0STJ8 | B0STJ8 |
| NPDFKGYNKTFQELHQLSK | NPDFKGYNKTFQELHQLSK | 75 | 75 | 5 | 9 | B0STJ8 | B0STJ8 |
| NPDFKGYNK | NPDFKGYNK | 75 | 75 | 5 | 7 | B0STJ8 | B0STJ8 |
| KEEAEKK | KEEAEKK | 126 | 126 | 1 | 6 | B0STJ8 | B0STJ8 |
| QDALNYFSKAINHLDEIK | QDALNYFSKAINHLDEIK | 162 | 162 | 8 | 9 | B0STJ8 | B0STJ8 |
| DSPEYKHIVNVINRDEVISVAQ | DSPEYKHIVNVINRDEVISVAQ | 279 | 279 | 2 | 6 | B0STJ8 | B0STJ8 |
| WYWQKVDRK | WYWQKVDRK | 138 | 138 | 2 | 5 | B0STJ8 | B0STJ8 |
| EYPAHKYLASAYSFEENMIKK | EYPAHKYLASAYSFEENMIKK | 229 | 229 | 6 | 10 | B0STJ8 | B0STJ8 |
| YGKDSPEYKHIVNVINR | YGKDSPEYKHIVNVINR | 276 | 276 | 3 | 9 | B0STJ8 | B0STJ8 |
| DSPEYKHIVNVINR | DSPEYKHIVNVINR | 279 | 279 | 2 | 6 | B0STJ8 | B0STJ8 |
| KTKGPDDMLFKYR | KTKGPDDMLFKYR | 249 | 249 | 1 | 3 | B0STJ8 | B0STJ8 |
| AKQDALNYFSK | EKMK | 160 | 156 | 2 | 2 | B0STJ8 | B0STJ8 |
| KMELFFEGSNNADPDPR | ADEKNESVSQSDATK | 130 | 20 | 1 | 4 | B0SR03 | B0SPP1 |
| NLGAKFK | GSLKY | 123 | 273 | 5 | 4 | B0SR03 | B0SR03 |
| QFYFKNLDYFDK | NVEALKGSLK | 236 | 267 | 5 | 6 | B0SR03 | B0SR03 |
| KNVEALK | NLGAKFK | 266 | 123 | 1 | 5 | B0SR03 | B0SR03 |
| KNVEALK | GSLKY | 266 | 273 | 1 | 4 | B0SR03 | B0SR03 |
| NVEALKGSLKY | NLGAKFK | 267 | 123 | 6 | 5 | B0SR03 | B0SR03 |
| NVEALKGSLKY | KQFYFK | 267 | 235 | 6 | 1 | B0SR03 | B0SR03 |
| GVGKYILSNVENTK | NLTKIR | 212 | 152 | 4 | 4 | B0SR03 | B0SR03 |
| YILSNVENTKSNPIR | KQFYFK | 216 | 235 | 10 | 1 | B0SR03 | B0SR03 |
| IFDYNDRDSNKHYKK | NVEALKGSLK | 252 | 267 | 11 | 6 | B0SR03 | B0SR03 |
| NVEALKGSLK | KQFYFK | 267 | 235 | 6 | 1 | B0SR03 | B0SR03 |
| IFDYNDR | KNVEALK | 252 | 266 | 4 | 1 | B0SR03 | B0SR03 |
| IFDYNDRDSNKHYK | KNVEALK | 252 | 266 | 11 | 1 | B0SR03 | B0SR03 |
| DSNKHYK | KNVEALK | 259 | 266 | 4 | 1 | B0SR03 | B0SR03 |
| IFDYNDRDSNK | KNVEALK | 252 | 266 | 9 | 1 | B0SR03 | B0SR03 |
| IFDYNDR | DSNKHYK | 252 | 259 | 4 | 4 | B0SR03 | B0SR03 |
| SSKNLGAKFK | SSKNLGAKFK | 120 | 120 | 3 | 8 | B0SR03 | B0SR03 |
| KMELFFEGSNNADPDPRK | KMELFFEGSNNADPDPRK | 130 | 130 | 1 | 9 | B0SR03 | B0SR03 |
| KMELFFEGSNNADPDPR | KMELFFEGSNNADPDPR | 130 | 130 | 1 | 9 | B0SR03 | B0SR03 |
| GVGKYILSNVENTK | GVGKYILSNVENTK | 212 | 212 | 4 | 8 | B0SR03 | B0SR03 |
| KQFYFKNLDYFDK | KQFYFKNLDYFDK | 235 | 235 | 1 | 6 | B0SR03 | B0SR03 |
| QFYFKNLDYFDKLFTK | QFYFKNLDYFDKLFTK | 236 | 236 | 5 | 12 | B0SR03 | B0SR03 |
| LFTKIFDYNDRDSNK | LFTKIFDYNDRDSNK | 248 | 248 | 4 | 13 | B0SR03 | B0SR03 |
| IFDYNDRDSNKHYKKNVEALK | IFDYNDRDSNKHYKKNVEALK | 252 | 252 | 11 | 14 | B0SR03 | B0SR03 |
| IFDYNDRDSNKHYKK | IFDYNDRDSNKHYKK | 252 | 252 | 11 | 14 | B0SR03 | B0SR03 |
| DSNKHYKKNVEALK | DSNKHYKKNVEALK | 259 | 259 | 4 | 7 | B0SR03 | B0SR03 |
| DSNKHYKK | DSNKHYKK | 259 | 259 | 4 | 7 | B0SR03 | B0SR03 |
| HYKKNVEALK | HYKKNVEALK | 263 | 263 | 3 | 4 | B0SR03 | B0SR03 |
| KNVEALKGSLKY | KNVEALKGSLKY | 266 | 266 | 1 | 7 | B0SR03 | B0SR03 |
| KNVEALKGSLK | KNVEALKGSLK | 266 | 266 | 1 | 7 | B0SR03 | B0SR03 |
| NVEALKGSLKY | NVEALKGSLKY | 267 | 267 | 6 | 10 | B0SR03 | B0SR03 |
| SMADTEKELDDNISEVNKR | SMADTEKELDDNISEVNKR | 40 | 40 | 1 | 7 | B0SR03 | B0SR03 |
| IFDYNDRDSNKHYK | IFDYNDRDSNKHYK | 252 | 252 | 9 | 11 | B0SR03 | B0SR03 |
| IELFYQHDDYPVWGTPETPSEK | IELFYQHDDYPVWGTPETPSEK | 190 | 190 | 15 | 20 | B0SR03 | B0SR03 |
| TLPHKTVLYK | TLPHKTVLYK | 72 | 72 | 1 | 5 | B0SR03 | B0SR03 |
| SGAISTSAKVR | VKLEQMLQK | 306 | 297 | 9 | 2 | B0SKT4 | B0SKT4 |
| SGAISTSAKVR | LVPGKPGDVK | 306 | 109 | 9 | 5 | B0SKT4 | B0SKT4 |
| FDENNKYSPVLNDPAKSPLVSR | SYIDADNKK | 84 | 159 | 6 | 8 | B0SKT4 | B0SKT4 |
| YSPVLNDPAKSPLVSR | SYIDADNKK | 90 | 159 | 10 | 8 | B0SKT4 | B0SKT4 |
| LVPGKPGDVKNVDAGNAK | SGAISTSAKVR | 109 | 306 | 5 | 9 | B0SKT4 | B0SKT4 |
| VRDCGGSNGSSQNK | NVDAGNAKVLAVK | 315 | 119 | 7 | 8 | B0SKT4 | B0SKT4 |
| VRDCGGSNGSSQNK | LVPGKPGDVK | 315 | 109 | 7 | 5 | B0SKT4 | B0SKT4 |
| LVPGKPGDVKNVDAGNAK | LVPGKPGDVKNVDAGNAK | 109 | 109 | 5 | 10 | B0SKT4 | B0SKT4 |
| LVPGKPGDVKNVDAGNAK | VRDCGGSNGSSQNK | 109 | 315 | 5 | 7 | B0SKT4 | B0SKT4 |
| NVDAGNAKVLAVK | NLIFKQFK | 119 | 244 | 8 | 5 | B0SKT4 | B0SKT4 |
| YSPVLNDPAKSPLVSR | DGSGDMKFDENNK | 90 | 77 | 10 | 7 | B0SKT4 | B0SKT4 |
| EVKLVPGKPGDVK | NVDAGNAKVLAVK | 106 | 119 | 3 | 8 | B0SKT4 | B0SKT4 |
| FDENNKYSPVLNDPAKSPLVSR | FDENNKYSPVLNDPAKSPLVSR | 84 | 84 | 6 | 16 | B0SKT4 | B0SKT4 |
| KMEKVYGIEAPGNVK | KMEKVYGIEAPGNVK | 168 | 168 | 1 | 4 | B0SKT4 | B0SKT4 |
| MEKVYGIEAPGNVK | MEKVYGIEAPGNVK | 169 | 169 | 3 | 5 | B0SKT4 | B0SKT4 |
| NLIFKQFKIR | NLIFKQFKIR | 244 | 244 | 5 | 8 | B0SKT4 | B0SKT4 |
| VKLEQMLQKSGAISTSAKVR | VKLEQMLQKSGAISTSAKVR | 297 | 297 | 2 | 9 | B0SKT4 | B0SKT4 |
| VKLEQMLQKSGAISTSAK | VKLEQMLQKSGAISTSAK | 297 | 297 | 2 | 10 | B0SKT4 | B0SKT4 |
| LEQMLQKSGAISTSAKVR | LEQMLQKSGAISTSAKVR | 299 | 299 | 7 | 16 | B0SKT4 | B0SKT4 |
| LEQMLQKSGAISTSAK | LEQMLQKSGAISTSAK | 299 | 299 | 7 | 8 | B0SKT4 | B0SKT4 |
| DGSGDMKFDENNKYSPVLNDPAK | DGSGDMKFDENNKYSPVLNDPAK | 77 | 77 | 7 | 13 | B0SKT4 | B0SKT4 |
| FDENNKYSPVLNDPAK | FDENNKYSPVLNDPAK | 84 | 84 | 6 | 8 | B0SKT4 | B0SKT4 |
| EVKLVPGKPGDVKNVDAGNAK | EVKLVPGKPGDVKNVDAGNAK | 106 | 106 | 3 | 8 | B0SKT4 | B0SKT4 |
| EVKLVPGKPGDVK | EVKLVPGKPGDVK | 106 | 106 | 3 | 8 | B0SKT4 | B0SKT4 |
| LVPGKPGDVKNVDAGNAKVLAVK | LVPGKPGDVKNVDAGNAKVLAVK | 109 | 109 | 5 | 10 | B0SKT4 | B0SKT4 |
| QYKVSNGVATPVGNDIGALELK | QYKVSNGVATPVGNDIGALELK | 26 | 26 | 3 | 3 | B0SKT4 | B0SKT4 |
| YSPVLNDPAKSPLVSR | YSPVLNDPAKSPLVSR | 90 | 90 | 2 | 10 | B0SKT4 | B0SKT4 |
| NVDAGNAKVLAVK | LVPGKPGDVK | 119 | 109 | 8 | 5 | B0SKT4 | B0SKT4 |
| MEKVYGIEAPGNVK | SYIDADNKK | 169 | 159 | 3 | 8 | B0SKT4 | B0SKT4 |
| KMEKVYGIEAPGNVK | SYIDADNKK | 168 | 159 | 4 | 8 | B0SKT4 | B0SKT4 |
| LVPGKPGDVKNVDAGNAK | LVPGKPGDVKNVDAGNAK | 109 | 109 | 5 | 10 | B0SKT4 | B0SKT4 |
| VELYPPHEYIIKGK | NPYIKYDNR | 118 | 298 | 12 | 5 | B0SKT5 | B0SJC6 |
| VELYPPHEYIIKGK | DGVNKDGSGDMK | 118 | 72 | 12 | 5 | B0SKT5 | B0SKT4 |
| SEAKYPGSEIK | GLIKDVFDEK | 228 | 79 | 4 | 4 | B0SKT5 | B0SKT5 |
| VELYPPHEYIIKGK | SEAKYPGSEIK | 118 | 228 | 12 | 4 | B0SKT5 | B0SKT5 |
| VELYPPHEYIIKGK | YPGSEIKDNW | 118 | 232 | 12 | 7 | B0SKT5 | B0SKT5 |
| KLTATVPGFIPQSTR | KIVLDDFEEAEDWR | 174 | 47 | 1 | 1 | B0SKT5 | B0SKT5 |
| GLIKDVFDEKTVPEDGGDKIEK | KLTATVPGFIPQSTR | 79 | 174 | 4 | 1 | B0SKT5 | B0SKT5 |
| DVFDEKTVPEDGGDKIEK | KLTATVPGFIPQSTR | 83 | 174 | 6 | 1 | B0SKT5 | B0SKT5 |
| TVPEDGGDKIEK | GLIKDVFDEK | 89 | 79 | 9 | 4 | B0SKT5 | B0SKT5 |
| IEKNHILGVK | TVPEDGGDK | 98 | 89 | 3 | 1 | B0SKT5 | B0SKT5 |
| ATTPLGETR | TLKLVQR | 63 | 72 | 2 | 3 | B0SKT5 | B0SKT5 |
| DYKDVTHNIR | DYKDVTHNIR | 154 | 154 | 3 | 6 | B0SKT5 | B0SKT5 |
| KLTATVPGFIPQSTR | KLTATVPGFIPQSTR | 174 | 174 | 1 | 5 | B0SKT5 | B0SKT5 |
| TDRSEAKYPGSEIKDNW | TDRSEAKYPGSEIKDNW | 225 | 225 | 7 | 14 | B0SKT5 | B0SKT5 |
| GLIKDVFDEKTVPEDGGDKIEK | GLIKDVFDEKTVPEDGGDKIEK | 79 | 79 | 4 | 10 | B0SKT5 | B0SKT5 |
| DVFDEKTVPEDGGDKIEK | DVFDEKTVPEDGGDKIEK | 83 | 83 | 6 | 15 | B0SKT5 | B0SKT5 |
| TVPEDGGDKIEKNHILGVK | TVPEDGGDKIEKNHILGVK | 89 | 89 | 9 | 12 | B0SKT5 | B0SKT5 |
| NHILGVKTHFAAK | NHILGVKTHFAAK | 101 | 101 | 7 | 8 | B0SKT5 | B0SKT5 |
| HTLFAKFR | HTLFAKFR | 146 | 146 | 2 | 6 | B0SKT5 | B0SKT5 |
| THFAAKGLDR | THFAAKGLDR | 108 | 108 | 1 | 6 | B0SKT5 | B0SKT5 |
| GLDRVELYPPHEYIIKGK | GLDRVELYPPHEYIIKGK | 114 | 114 | 8 | 16 | B0SKT5 | B0SKT5 |
| GLDRVELYPPHEYIIK | GLDRVELYPPHEYIIK | 114 | 114 | 8 | 13 | B0SKT5 | B0SKT5 |
| MIINHNLAAINSHR | ALKFTQWDVDK | 1 | 15 | 1 | 3 | B0SSZ5 | B0SSZ4 |
| VLKFQNEEVSK | MIINHNMSAIQSHR | 15 | 1 | 3 | 1 | B0SSZ5 | B0SSZ4 |
| AGDDASGLAVSEKMR | MIINHNVSAIFAHR | 39 | 1 | 13 | 1 | B0SSZ5 | B0SQZ5 |
| MIINHNLAAINSHR | TLKSNDANLSK | 1 | 15 | 1 | 3 | B0SSZ5 | B0SQZ5 |
| VLKFQNEEVSK | MIINHNVSAIFAHR | 15 | 1 | 3 | 1 | B0SSZ5 | B0SQZ5 |
| NQILVQSGTAMLAQANVRPQGVLSLLR | IRDADMAEETVAFTK | 255 | 240 | 7 | 10 | B0SSZ5 | B0SSZ5 |
| MIINHNLAAINSHR | VLKFQNEEVSK | 1 | 15 | 1 | 3 | B0SSZ5 | B0SSZ5 |
| VLKFQNEEVSK | VLKFQNEEVSK | 15 | 15 | 3 | 10 | B0SSZ5 | B0SSZ5 |
| SNDAIGTLDAALTR | ISKQR | 191 | 205 | 1 | 3 | B0SSZ5 | B0SSZ5 |
| VLKFQNEEVSK | VLKFQNEEVSK | 15 | 15 | 3 | 10 | B0SSZ5 | B0SSZ5 |
| LEHAAKGLMNAYENTQASESR | LEHAAKGLMNAYENTQASESR | 219 | 219 | 6 | 18 | B0SSZ5 | B0SSZ5 |
| MIINHNLAAINSHRVLKFQNEEVSK | MIINHNLAAINSHRVLKFQNEEVSK | 1 | 1 | 1 | 17 | B0SSZ5 | B0SSZ5 |
| MIINHNLAAINSHR | MIINHNLAAINSHR | 1 | 1 | 1 | 12 | B0SSZ5 | B0SSZ5 |
| GLMNAYENTQASESR | GLMNAYENTQASESR | 225 | 225 | 12 | 14 | B0SSZ5 | B0SSZ5 |
| NQILVQSGTAMLAQANVRPQGVLSLLR | NQILVQSGTAMLAQANVRPQGVLSLLR | 255 | 255 | 7 | 24 | B0SSZ5 | B0SSZ5 |
| ADEAIAKADFALNK | ALKMSEGAIK | 190 | 172 | 7 | 3 | B0SSZ4 | B0SSZ4 |
| ADFALNKIMK | ALKMSEGAIK | 197 | 172 | 7 | 3 | B0SSZ4 | B0SSZ4 |
| QILVQSGTAMLAQASLRPNSVLR | IRDADMAEEMVALTTK | 255 | 239 | 6 | 14 | B0SSZ4 | B0SSZ4 |
| MIINHNMSAIQSHR | ALKFTQWDVDK | 1 | 15 | 1 | 3 | B0SSZ4 | B0SSZ4 |
| MIINHNMSAIQSHRALKFTQWDVDK | MIINHNMSAIQSHRALKFTQWDVDK | 1 | 1 | 1 | 17 | B0SSZ4 | B0SSZ4 |
| MIINHNMSAIQSHR | MIINHNMSAIQSHR | 1 | 1 | 1 | 8 | B0SSZ4 | B0SSZ4 |
| IGTSAEFNKVKPLDGR | IGTSAEFNKVKPLDGR | 126 | 126 | 4 | 11 | B0SS86 | B0SS86 |
| INKAGDDASGLAVSEK | MIINHNVSAIFAHR | 36 | 1 | 3 | 1 | B0SQZ5 | B0SQZ5 |
| MIINHNVSAIFAHRTLKSNDANLSK | TLKSNDANLSK | 1 | 15 | 1 | 3 | B0SQZ5 | B0SQZ5 |
| MIINHNVSAIFAHR | TLKSNDANLSK | 1 | 15 | 1 | 3 | B0SQZ5 | B0SQZ5 |
| MIINHNVSAIFAHR | SNDANLSKDIEK | 1 | 18 | 1 | 8 | B0SQZ5 | B0SQZ5 |

| | | | | | | | |
|---|---|---|---|---|---|---|---|
| **MIINHNVSAIFAHRTLK** | SNDANLSK | 1 | 18 | 1 | 1 | B0SQZ5 | B0SQZ5 |
| **TLKSNDANLSKDIEK** | TLKSNDANLSKDIEK | 15 | 15 | 3 | 11 | B0SQZ5 | B0SQZ5 |
| **SNDANLSKDIEKLSSGMR** | SNDANLSKDIEKLSSGMR | 18 | 18 | 8 | 12 | B0SQZ5 | B0SQZ5 |
| **DIEKLSSGMR** | DIEKLSSGMR | 26 | 26 | 4 | 6 | B0SQZ5 | B0SQZ5 |
| **INKAGDDASGLAVSEK** | INKAGDDASGLAVSEK | 36 | 36 | 3 | 9 | B0SQZ5 | B0SQZ5 |
| **MIINHNVSAIFAHR** | MIINHNVSAIFAHR | 1 | 1 | 1 | 8 | B0SQZ5 | B0SQZ5 |
| **MIINHNVSAIFAHRTLKSNDANLSKDIEK** | MIINHNVSAIFAHRTLKSNDANLSKDIEK | 1 | 1 | 17 | 25 | B0SQZ5 | B0SQZ5 |
