## Supplementary Table 3 for "Core–sheath coupling controls flagellar curvature and motility in *Leptospira*"

**Table S3.** Label-free comparative proteomics of *L. biflexa* filaments from *wt vs* mutant strains.

**Label-free comparative proteomics of *L. biflexa* filaments from *wt* and *flaA2-* mutant strains. Analysis by Spectral Counts (SC)**

Proteins identified in the cryoEM structure of *L. biflexa* filaments are marked in red

Core flagellins (FlaBs) and paralogues of *Leptospira* flagellar filament proteins are shown in **bold** (see Supplementary Table 4)

### Venn Diagram

**Proteins exclusively identified in *L. biflexa wt***

| | Number of Spectra | | | | | |
|---|---|---|---|---|---|---|
| **Locus** | **WT_R1** | **WT_R2** | **WT_R3** | **Total Signal** | **Replicate Count** | **Description** |
| tr\|B0SKT5\|B0SKT5_LEPBP | 283 | 135 | 199 | 617 | 3 | Flagellar filament outer layer protein (Sheath protein) putative signal peptide OS=Leptospira biflexa serovar Patoc (strain Patoc 1 / ATCC 23582 / Paris) OX=456481 GN=flaA2 PE=4 SV=1 |
| tr\|B0SKT4\|B0SKT4_LEPBP | 101 | 12 | 75 | 188 | 3 | Flagellar filament outer layer protein A OS=Leptospira biflexa serovar Patoc (strain Patoc 1 / ATCC 23582 / Paris) OX=456481 GN=flaA1 PE=4 SV=1 |
| **tr\|B0SNJ6\|B0SNJ6_LEPBP** | 26 | 12 | 43 | 81 | 3 | Uncharacterized protein OS=Leptospira biflexa serovar Patoc (strain Patoc 1 / ATCC 23582 / Paris) OX=456481 GN=LEPBI_I1161 PE=4 SV=1 |
| **tr\|B0SQV9\|B0SQV9_LEPBP** | 20 | 20 | 21 | 61 | 3 | Uncharacterized protein OS=Leptospira biflexa serovar Patoc (strain Patoc 1 / ATCC 23582 / Paris) OX=456481 GN=LEPBI_I3291 PE=4 SV=1 |
| tr\|B0SQJ6\|B0SQJ6_LEPBP | 7 | 3 | 4 | 14 | 3 | Putative short-chain dehydrogenase/reductase SDR OS=Leptospira biflexa serovar Patoc (strain Patoc 1 / ATCC 23582 / Paris) OX=456481 GN=LEPBI_I1547 PE=4 SV=1 |
| tr\|B0SMT6\|B0SMT6_LEPBP | 4 | 2 | 4 | 10 | 3 | Uncharacterized protein OS=Leptospira biflexa serovar Patoc (strain Patoc 1 / ATCC 23582 / Paris) OX=456481 GN=LEPBI_I2732 PE=4 SV=1 |
| tr\|B0SPA5\|B0SPA5_LEPBP | | 3 | 3 | 6 | 2 | tRNA N6-adenosine threonylcarbamoyltransferase OS=Leptospira biflexa serovar Patoc (strain Patoc 1 / ATCC 23582 / Paris) OX=456481 GN=tsaD PE=3 SV=1 |
| tr\|B0SU70\|B0SU70_LEPBP | 2 | 3 | | 5 | 2 | Ankyrin_rpt-contain_dom domain-containing protein OS=Leptospira biflexa serovar Patoc (strain Patoc 1 / ATCC 23582 / Paris) OX=456481 GN=LEPBI_II0221 PE=4 SV=1 |

**Proteins exclusively identified in *L. biflexa flaA2-***

| | Number of Spectra | | | | | |
|---|---|---|---|---|---|---|
| **Locus** | **FlaA_R1** | **FlaA_R2** | **FlaA_R3** | **Total Signal** | **Replicate Count** | **Description** |
| tr\|B0SRK9\|B0SRK9_LEPBP | 53 | 176 | 213 | 442 | 3 | Succinate dehydrogenase flavoprotein subunit SdhA (Fumarate reductase) OS=Leptospira biflexa serovar Patoc (strain Patoc 1 / ATCC 23582 / Paris) OX=456481 GN=sdhA1 PE=4 SV=1 |
| tr\|B0STP2\|B0STP2_LEPBP | 70 | 54 | 47 | 171 | 3 | Putative alkyldihydroxyacetonephosphate synthase OS=Leptospira biflexa serovar Patoc (strain Patoc 1 / ATCC 23582 / Paris) OX=456481 GN=eapA PE=4 SV=1 |
| tr\|B0SMZ3\|B0SMZ3_LEPBP | 85 | 56 | 21 | 162 | 3 | Dihydrolipoyl dehydrogenase OS=Leptospira biflexa serovar Patoc (strain Patoc 1 / ATCC 23582 / Paris) OX=456481 GN=lpdA1 PE=3 SV=1 |
| tr\|B0SUE2\|B0SUE2_LEPBP | 53 | 48 | 32 | 133 | 3 | Hemin degradation protein HemS OS=Leptospira biflexa serovar Patoc (strain Patoc 1 / ATCC 23582 / Paris) OX=456481 GN=hemS PE=4 SV=1 |
| tr\|B0SPN5\|B0SPN5_LEPBP | 48 | 48 | 10 | 106 | 3 | Penicillin-binding protein 2 OS=Leptospira biflexa serovar Patoc (strain Patoc 1 / ATCC 23582 / Paris) OX=456481 GN=mrdA PE=4 SV=1 |
| tr\|B0SRR4\|B0SRR4_LEPBP | 51 | 31 | 15 | 97 | 3 | UDP-N-acetylmuramoyl-tripeptide--D-alanyl-D-alanine ligase OS=Leptospira biflexa serovar Patoc (strain Patoc 1 / ATCC 23582 / Paris) OX=456481 GN=murF PE=3 SV=1 |
| tr\|B0SME8\|B0SME8_LEPBP | 22 | 37 | 33 | 92 | 3 | Trigger factor OS=Leptospira biflexa serovar Patoc (strain Patoc 1 / ATCC 23582 / Paris) OX=456481 GN=tig PE=3 SV=1 |
| tr\|B0SUD2\|B0SUD2_LEPBP | 30 | 28 | 31 | 89 | 3 | Putative chromosome partitioning protein ParB OS=Leptospira biflexa serovar Patoc (strain Patoc 1 / ATCC 23582 / Paris) OX=456481 GN=LEPBI_p0002 PE=3 SV=1 |
| sp\|B0SNZ7\|GATB_LEPBP | 14 | 29 | 40 | 83 | 3 | Aspartyl/glutamyl-tRNA(Asn/Gln) amidotransferase subunit B OS=Leptospira biflexa serovar Patoc (strain Patoc 1 / ATCC 23582 / Paris) OX=456481 GN=gatB PE=3 SV=1 |
| tr\|B0SL67\|B0SL67_LEPBP | 27 | 27 | 28 | 82 | 3 | Glutamate-1-semialdehyde 2,1-aminomutase OS=Leptospira biflexa serovar Patoc (strain Patoc 1 / ATCC 23582 / Paris) OX=456481 GN=hemL2 PE=3 SV=1 |
| tr\|B0SPZ1\|B0SPZ1_LEPBP | 40 | 23 | 18 | 81 | 3 | SPOR domain-containing protein OS=Leptospira biflexa serovar Patoc (strain Patoc 1 / ATCC 23582 / Paris) OX=456481 GN=LEPBI_I1449 PE=4 SV=1 |
| tr\|B0SK30\|B0SK30_LEPBP | 32 | 36 | 10 | 78 | 3 | Beta-galactosidase OS=Leptospira biflexa serovar Patoc (strain Patoc 1 / ATCC 23582 / Paris) OX=456481 GN=LEPBI_I0024 PE=3 SV=1 |
| tr\|B0SRI4\|B0SRI4_LEPBP | 14 | 22 | 32 | 68 | 3 | Putative oxidoreductase, GMC family OS=Leptospira biflexa serovar Patoc (strain Patoc 1 / ATCC 23582 / Paris) OX=456481 GN=LEPBI_I3414 PE=4 SV=1 |
| tr\|B0SSM7\|B0SSM7_LEPBP | 34 | 22 | 11 | 67 | 3 | Glyco_trans_2-like domain-containing protein OS=Leptospira biflexa serovar Patoc (strain Patoc 1 / ATCC 23582 / Paris) OX=456481 GN=LEPBI_I2015 PE=4 SV=1 |
| tr\|B0SSM9\|B0SSM9_LEPBP | 30 | 23 | 14 | 67 | 3 | ABC-type transport system, ATP binding protein OS=Leptospira biflexa serovar Patoc (strain Patoc 1 / ATCC 23582 / Paris) OX=456481 GN=LEPBI_I2017 PE=3 SV=1 |
| tr\|B0SNJ0\|B0SNJ0_LEPBP | 29 | 20 | 17 | 66 | 3 | Uncharacterized protein OS=Leptospira biflexa serovar Patoc (strain Patoc 1 / ATCC 23582 / Paris) OX=456481 GN=LEPBI_I1155 PE=4 SV=1 |
| tr\|B0SPE8\|B0SPE8_LEPBP | 23 | 24 | 18 | 65 | 3 | Putative alkaline phosphatase putative membrane protein OS=Leptospira biflexa serovar Patoc (strain Patoc 1 / ATCC 23582 / Paris) OX=456481 GN=LEPBI_I1362 PE=4 SV=1 |
| tr\|B0SPC0\|B0SPC0_LEPBP | 25 | 12 | 27 | 64 | 3 | Uncharacterized protein OS=Leptospira biflexa serovar Patoc (strain Patoc 1 / ATCC 23582 / Paris) OX=456481 GN=LEPBI_I1334 PE=4 SV=1 |
| tr\|B0SL64\|B0SL64_LEPBP | 25 | 23 | 11 | 59 | 3 | Flagellar M-ring protein OS=Leptospira biflexa serovar Patoc (strain Patoc 1 / ATCC 23582 / Paris) OX=456481 GN=fliF PE=3 SV=1 |
| tr\|B0SMC3\|B0SMC3_LEPBP | 21 | 12 | 20 | 53 | 3 | Flagella-associated GTP-binding protein OS=Leptospira biflexa serovar Patoc (strain Patoc 1 / ATCC 23582 / Paris) OX=456481 GN=flhF PE=3 SV=1 |
| tr\|B0SL95\|B0SL95_LEPBP | 22 | 23 | 7 | 52 | 3 | Alpha-glucosidase OS=Leptospira biflexa serovar Patoc (strain Patoc 1 / ATCC 23582 / Paris) OX=456481 GN=LEPBI_I0772 PE=4 SV=1 |
| sp\|B0SNG1\|SECA_LEPBP | 13 | 29 | 10 | 52 | 3 | Protein translocase subunit SecA OS=Leptospira biflexa serovar Patoc (strain Patoc 1 / ATCC 23582 / Paris) OX=456481 GN=secA PE=3 SV=1 |
| tr\|B0SQH9\|B0SQH9_LEPBP | 15 | 13 | 23 | 51 | 3 | Putative Zn-dependent peptidase OS=Leptospira biflexa serovar Patoc (strain Patoc 1 / ATCC 23582 / Paris) OX=456481 GN=LEPBI_I1530 PE=4 SV=1 |
| tr\|B0SRJ5\|B0SRJ5_LEPBP | 21 | 20 | 8 | 49 | 3 | Cell division protein FtsA OS=Leptospira biflexa serovar Patoc (strain Patoc 1 / ATCC 23582 / Paris) OX=456481 GN=ftsA2 PE=4 SV=1 |
| tr\|B0SS54\|B0SS54_LEPBP | 26 | 20 | 2 | 48 | 3 | Putative petidase S49, protease IV family OS=Leptospira biflexa serovar Patoc (strain Patoc 1 / ATCC 23582 / Paris) OX=456481 GN=LEPBI_I1838 PE=4 SV=1 |
| tr\|B0SQ78\|B0SQ78_LEPBP | 7 | 27 | 13 | 47 | 3 | MaoC_dehydrat_N domain-containing protein OS=Leptospira biflexa serovar Patoc (strain Patoc 1 / ATCC 23582 / Paris) OX=456481 GN=LEPBI_I3165 PE=4 SV=1 |
| tr\|B0SSZ6\|B0SSZ6_LEPBP | 22 | 13 | 11 | 46 | 3 | Uncharacterized protein OS=Leptospira biflexa serovar Patoc (strain Patoc 1 / ATCC 23582 / Paris) OX=456481 GN=LEPBI_I2134 PE=4 SV=1 |
| tr\|B0SLS4\|B0SLS4_LEPBP | 21 | 18 | 5 | 44 | 3 | ABC-type transport system, ATP binding protein and permease putative membrane protein putative signal peptide OS=Leptospira biflexa serovar Patoc (strain Patoc 1 / ATCC 23582 / Paris) OX=456481 GN=LEPBI_I0849 PE=4 SV=1 |
| tr\|B0SLF2\|B0SLF2_LEPBP | 8 | 28 | 7 | 43 | 3 | Chaperone ClpB OS=Leptospira biflexa serovar Patoc (strain Patoc 1 / ATCC 23582 / Paris) OX=456481 GN=clpB PE=3 SV=1 |
| tr\|B0STP5\|B0STP5_LEPBP | 22 | 14 | 7 | 43 | 3 | Glycerol-3-phosphate oxidase OS=Leptospira biflexa serovar Patoc (strain Patoc 1 / ATCC 23582 / Paris) OX=456481 GN=glpD PE=3 SV=1 |
| tr\|B0SSB7\|B0SSB7_LEPBP | 18 | 12 | 13 | 43 | 3 | tRNA uridine(34) hydroxylase OS=Leptospira biflexa serovar Patoc (strain Patoc 1 / ATCC 23582 / Paris) OX=456481 GN=trhO PE=3 SV=1 |
| tr\|B0SQ49\|B0SQ49_LEPBP | 3 | 16 | 23 | 42 | 3 | Succinate dehydrogenase flavoprotein subunit SdhA (Fumarate reductase) OS=Leptospira biflexa serovar Patoc (strain Patoc 1 / ATCC 23582 / Paris) OX=456481 GN=sdhA2 PE=4 SV=1 |
| tr\|B0SJH9\|B0SJH9_LEPBP | 15 | 23 | 4 | 42 | 3 | Uncharacterized protein OS=Leptospira biflexa serovar Patoc (strain Patoc 1 / ATCC 23582 / Paris) OX=456481 GN=LEPBI_I0389 PE=4 SV=1 |
| tr\|B0SUC3\|B0SUC3_LEPBP | 17 | 22 | 3 | 42 | 3 | Uncharacterized protein OS=Leptospira biflexa serovar Patoc (strain Patoc 1 / ATCC 23582 / Paris) OX=456481 GN=LEPBI_II0274 PE=4 SV=1 |
| tr\|B0SP04\|B0SP04_LEPBP | 20 | 20 | 2 | 42 | 3 | Putative two-component sensor protein putative membrane protein putative signal peptide OS=Leptospira biflexa serovar Patoc (strain Patoc 1 / ATCC 23582 / Paris) OX=456481 GN=LEPBI_I2948 PE=4 SV=1 |
| tr\|B0SS05\|B0SS05_LEPBP | 20 | 14 | 6 | 40 | 3 | Putative ADP-heptose:LPS heptosyltransferase II OS=Leptospira biflexa serovar Patoc (strain Patoc 1 / ATCC 23582 / Paris) OX=456481 GN=LEPBI_I1789 PE=4 SV=1 |
| tr\|B0SP92\|B0SP92_LEPBP | 13 | 17 | 8 | 38 | 3 | NADH-quinone oxidoreductase, chain M OS=Leptospira biflexa serovar Patoc (strain Patoc 1 / ATCC 23582 / Paris) OX=456481 GN=nuoM PE=3 SV=1 |
| tr\|B0SRZ6\|B0SRZ6_LEPBP | 21 | 12 | 4 | 37 | 3 | ABC-type transport system, ATP binding protein putative membrane protein putative signal peptide OS=Leptospira biflexa serovar Patoc (strain Patoc 1 / ATCC 23582 / Paris) OX=456481 GN=LEPBI_I3472 PE=4 SV=1 |
| tr\|B0SLV4\|B0SLV4_LEPBP | 23 | 9 | 5 | 37 | 3 | Putative ubiquinone biosynthesis protein OS=Leptospira biflexa serovar Patoc (strain Patoc 1 / ATCC 23582 / Paris) OX=456481 GN=LEPBI_I0881 PE=4 SV=1 |

### Volcano plot

**Proteins enriched in *L. biflexa wt* vs *flaA2-* (|FC| ≥ 2, p-value ≤ 0.05)**

| Locus | Fold Change | pValue | Signal(Bottom) | Signal(Top) | Description |
|---|---|---|---|---|---|
| tr\|B0SJC6\|B0SJC6_LEPBP | 18.15 | 2.47E-05 | 2.24E-03 | 4.06E-02 | Uncharacterized protein OS=Leptospira biflexa serovar Patoc (strain Patoc 1 / ATCC 23582 / Paris) OX=456481 GN=LEPBI_I0551 PE=4 SV=1 |
| tr\|B0SKN8\|B0SKN8_LEPBP | 16.58 | 1.00E-05 | 9.37E-04 | 1.55E-02 | Uncharacterized protein OS=Leptospira biflexa serovar Patoc (strain Patoc 1 / ATCC 23582 / Paris) OX=456481 GN=LEPBI_I0662 PE=4 SV=1 |
| tr\|B0SM18\|B0SM18_LEPBP | 15.49 | 1.27E-02 | 4.93E-05 | 7.63E-04 | O-succinylhomoserine sulfhydrylase OS=Leptospira biflexa serovar Patoc (strain Patoc 1 / ATCC 23582 / Paris) OX=456481 GN=metZ PE=3 SV=1 |
| tr\|B0SMK8\|B0SMK8_LEPBP | 14.27 | 8.75E-05 | 1.60E-03 | 2.29E-02 | Uncharacterized protein OS=Leptospira biflexa serovar Patoc (strain Patoc 1 / ATCC 23582 / Paris) OX=456481 GN=LEPBI_I1029 PE=4 SV=1 |
| tr\|B0STF2\|B0STF2_LEPBP | 14.17 | 4.40E-05 | 2.16E-03 | 3.07E-02 | Uncharacterized protein OS=Leptospira biflexa serovar Patoc (strain Patoc 1 / ATCC 23582 / Paris) OX=456481 GN=LEPBI_I2297 PE=4 SV=1 |
| tr\|B0SPP1\|B0SPP1_LEPBP | 8.84 | 3.91E-03 | 1.10E-03 | 9.71E-03 | Uncharacterized protein OS=Leptospira biflexa serovar Patoc (strain Patoc 1 / ATCC 23582 / Paris) OX=456481 GN=LEPBI_I3081 PE=4 SV=1 |
| tr\|B0SKF0\|B0SKF0_LEPBP | 7.74 | 2.76E-02 | 9.38E-05 | 7.25E-04 | Acyl-[acyl-carrier-protein]--UDP-N-acetylglucosamine O-acyltransferase OS=Leptospira biflexa serovar Patoc (strain Patoc 1 / ATCC 23582 / Paris) OX=456481 GN=lpxA PE=4 SV=1 |
| tr\|B0SUF9\|B0SUF9_LEPBP | 5.94 | 5.65E-04 | 9.56E-05 | 5.68E-04 | Uncharacterized protein OS=Leptospira biflexa serovar Patoc (strain Patoc 1 / ATCC 23582 / Paris) OX=456481 GN=LEPBI_p0029 PE=4 SV=1 |
| sp\|B0STJ3\|SYL_LEPBP | 5.82 | 2.12E-02 | 1.45E-04 | 8.46E-04 | Leucine--tRNA ligase OS=Leptospira biflexa serovar Patoc (strain Patoc 1 / ATCC 23582 / Paris) OX=456481 GN=leuS PE=3 SV=1 |
| tr\|B0SJA5\|B0SJA5_LEPBP | 5.63 | 1.03E-02 | 1.13E-04 | 6.37E-04 | Polyphosphate kinase OS=Leptospira biflexa serovar Patoc (strain Patoc 1 / ATCC 23582 / Paris) OX=456481 GN=ppk PE=3 SV=1 |
| tr\|B0STS1\|B0STS1_LEPBP | 5.22 | 9.02E-03 | 5.00E-04 | 2.61E-03 | Glutamate synthase large chain OS=Leptospira biflexa serovar Patoc (strain Patoc 1 / ATCC 23582 / Paris) OX=456481 GN=gltB PE=3 SV=1 |
| tr\|B0SK22\|B0SK22_LEPBP | 4.54 | 2.41E-03 | 2.63E-04 | 1.20E-03 | 3-oxoacyl-[acyl-carrier-protein] reductase OS=Leptospira biflexa serovar Patoc (strain Patoc 1 / ATCC 23582 / Paris) OX=456481 GN=fabG4 PE=4 SV=1 |
| tr\|B0SMM0\|B0SMM0_LEPBP | 4.23 | 1.64E-02 | 1.36E-04 | 5.76E-04 | Uncharacterized protein OS=Leptospira biflexa serovar Patoc (strain Patoc 1 / ATCC 23582 / Paris) OX=456481 GN=LEPBI_I2666 PE=4 SV=1 |
| tr\|B0SQY0\|B0SQY0_LEPBP | 4.14 | 1.57E-02 | 2.19E-04 | 9.08E-04 | Beta-lactamase domain-containing protein OS=Leptospira biflexa serovar Patoc (strain Patoc 1 / ATCC 23582 / Paris) OX=456481 GN=LEPBI_I3312 PE=4 SV=1 |
| tr\|B0SKE6\|B0SKE6_LEPBP | 4.10 | 1.67E-02 | 1.53E-04 | 6.27E-04 | Putative integral outer membrane protein TolC, efflux pump component putative signal peptide OS=Leptospira biflexa serovar Patoc (strain Patoc 1 / ATCC 23582 / Paris) OX=456481 GN=LEPBI_I0360 PE=3 SV=1 |
| tr\|B0SLY4\|B0SLY4_LEPBP | 3.69 | 1.22E-02 | 2.37E-04 | 8.73E-04 | Putative PLP-dependent aminotransferase OS=Leptospira biflexa serovar Patoc (strain Patoc 1 / ATCC 23582 / Paris) OX=456481 GN=LEPBI_I0912 PE=4 SV=1 |
| tr\|B0SQM4\|B0SQM4_LEPBP | 3.64 | 6.83E-03 | 1.80E-04 | 6.57E-04 | Putative alpha-glucosidase II OS=Leptospira biflexa serovar Patoc (strain Patoc 1 / ATCC 23582 / Paris) OX=456481 GN=LEPBI_I3206 PE=3 SV=1 |
| tr\|B0SJX7\|B0SJX7_LEPBP | 3.64 | 7.60E-03 | 3.50E-04 | 1.27E-03 | Uncharacterized protein OS=Leptospira biflexa serovar Patoc (strain Patoc 1 / ATCC 23582 / Paris) OX=456481 GN=LEPBI_I0145 PE=4 SV=1 |
| tr\|B0SKQ9\|B0SKQ9_LEPBP | 3.33 | 6.90E-04 | 2.42E-04 | 8.07E-04 | Uncharacterized protein OS=Leptospira biflexa serovar Patoc (strain Patoc 1 / ATCC 23582 / Paris) OX=456481 GN=LEPBI_I0684 PE=4 SV=1 |
| tr\|B0SJ54\|B0SJ54_LEPBP | 3.33 | 1.15E-02 | 1.79E-04 | 5.96E-04 | Uncharacterized protein OS=Leptospira biflexa serovar Patoc (strain Patoc 1 / ATCC 23582 / Paris) OX=456481 GN=LEPBI_I0512 PE=4 SV=1 |
| tr\|B0SM40\|B0SM40_LEPBP | 3.31 | 9.57E-03 | 3.91E-04 | 1.29E-03 | 3-oxoacyl-[acyl-carrier-protein] reductase OS=Leptospira biflexa serovar Patoc (strain Patoc 1 / ATCC 23582 / Paris) OX=456481 GN=fabG PE=3 SV=1 |
| tr\|B0SS97\|B0SS97_LEPBP | 3.10 | 3.49E-03 | 3.90E-03 | 1.21E-02 | Iron(III) dicitrate TonB-dependent receptor putative signal peptide OS=Leptospira biflexa serovar Patoc (strain Patoc 1 / ATCC 23582 / Paris) OX=456481 GN=fecA PE=3 SV=1 |
| tr\|B0STW0\|B0STW0_LEPBP | 3.05 | 1.89E-03 | 5.68E-04 | 1.73E-03 | Uncharacterized protein OS=Leptospira biflexa serovar Patoc (strain Patoc 1 / ATCC 23582 / Paris) OX=456481 GN=LEPBI_II0108 PE=4 SV=1 |
| tr\|B0SLY1\|B0SLY1_LEPBP | 3.04 | 2.03E-02 | 3.31E-04 | 1.01E-03 | Putative periplasmic component of the Tol biopolymer transport system OS=Leptospira biflexa serovar Patoc (strain Patoc 1 / ATCC 23582 / Paris) OX=456481 GN=LEPBI_I0909 PE=4 SV=1 |
| tr\|B0SK54\|B0SK54_LEPBP | 3.01 | 1.51E-02 | 3.58E-04 | 1.08E-03 | 3-oxoacyl-[acyl-carrier-protein] reductase OS=Leptospira biflexa serovar Patoc (strain Patoc 1 / ATCC 23582 / Paris) OX=456481 GN=fabG3b PE=4 SV=1 |
| tr\|B0SPM8\|B0SPM8_LEPBP | 2.96 | 4.06E-03 | 2.64E-04 | 7.83E-04 | Putative ankyrin-like protein OS=Leptospira biflexa serovar Patoc (strain Patoc 1 / ATCC 23582 / Paris) OX=456481 GN=LEPBI_I3068 PE=4 SV=1 |
| tr\|B0STW1\|B0STW1_LEPBP | 2.80 | 2.43E-03 | 3.96E-04 | 1.11E-03 | Uncharacterized protein OS=Leptospira biflexa serovar Patoc (strain Patoc 1 / ATCC 23582 / Paris) OX=456481 GN=LEPBI_II0109 PE=4 SV=1 |
| tr\|B0SQ25\|B0SQ25_LEPBP | 2.78 | 3.82E-03 | 9.67E-04 | 2.69E-03 | Uncharacterized protein OS=Leptospira biflexa serovar Patoc (strain Patoc 1 / ATCC 23582 / Paris) OX=456481 GN=LEPBI_I3111 PE=4 SV=1 |
| tr\|B0SPD3\|B0SPD3_LEPBP | 2.76 | 1.01E-02 | 5.25E-04 | 1.45E-03 | Uncharacterized protein OS=Leptospira biflexa serovar Patoc (strain Patoc 1 / ATCC 23582 / Paris) OX=456481 GN=LEPBI_I1347 PE=4 SV=1 |
| tr\|B0SR80\|B0SR80_LEPBP | 2.73 | 1.64E-02 | 3.80E-04 | 1.04E-03 | General secretion pathway protein C putative signal peptide OS=Leptospira biflexa serovar Patoc (strain Patoc 1 / ATCC 23582 / Paris) OX=456481 GN=gspC PE=4 SV=1 |
| tr\|B0SMX3\|B0SMX3_LEPBP | 2.57 | 2.50E-03 | 2.09E-03 | 5.37E-03 | Uncharacterized protein OS=Leptospira biflexa serovar Patoc (strain Patoc 1 / ATCC 23582 / Paris) OX=456481 GN=LEPBI_I1037 PE=4 SV=1 |
| tr\|B0SKQ7\|B0SKQ7_LEPBP | 2.57 | 2.33E-02 | 3.71E-04 | 9.55E-04 | Uncharacterized protein OS=Leptospira biflexa serovar Patoc (strain Patoc 1 / ATCC 23582 / Paris) OX=456481 GN=LEPBI_I0682 PE=4 SV=1 |
| tr\|B0SQ77\|B0SQ77_LEPBP | 2.57 | 2.53E-02 | 5.23E-04 | 1.34E-03 | Putative acylglycerol lipase OS=Leptospira biflexa serovar Patoc (strain Patoc 1 / ATCC 23582 / Paris) OX=456481 GN=LEPBI_I3164 PE=4 SV=1 |
| tr\|B0SLN1\|B0SLN1_LEPBP | 2.49 | 3.96E-03 | 3.58E-04 | 8.93E-04 | Uncharacterized protein OS=Leptospira biflexa serovar Patoc (strain Patoc 1 / ATCC 23582 / Paris) OX=456481 GN=LEPBI_I2536 PE=4 SV=1 |
| tr\|B0SSW5\|B0SSW5_LEPBP | 2.35 | 1.02E-02 | 4.31E-04 | 1.01E-03 | Uncharacterized protein OS=Leptospira biflexa serovar Patoc (strain Patoc 1 / ATCC 23582 / Paris) OX=456481 GN=LEPBI_I2103 PE=4 SV=1 |
| tr\|B0SKP2\|B0SKP2_LEPBP | 2.31 | 1.61E-03 | 9.27E-04 | 2.15E-03 | Uncharacterized protein OS=Leptospira biflexa serovar Patoc (strain Patoc 1 / ATCC 23582 / Paris) OX=456481 GN=LEPBI_I0666 PE=4 SV=1 |
| tr\|B0SQQ8\|B0SQQ8_LEPBP | 2.29 | 7.45E-04 | 1.17E-03 | 2.68E-03 | Putative TPR-repeat-containing protein putative signal peptide OS=Leptospira biflexa serovar Patoc (strain Patoc 1 / ATCC 23582 / Paris) OX=456481 GN=LEPBI_I3240 PE=4 SV=1 |
| sp\|B0SJR2\|ASSY_LEPBP | 2.27 | 4.13E-03 | 2.39E-04 | 5.43E-04 | Argininosuccinate synthase OS=Leptospira biflexa serovar Patoc (strain Patoc 1 / ATCC 23582 / Paris) OX=456481 GN=argG PE=3 SV=1 |
| tr\|B0SQC6\|B0SQC6_LEPBP | 2.19 | 1.32E-03 | 3.21E-03 | 7.02E-03 | Uncharacterized protein OS=Leptospira biflexa serovar Patoc (strain Patoc 1 / ATCC 23582 / Paris) OX=456481 GN=LEPBI_I1477 PE=4 SV=1 |
| tr\|B0SKQ8\|B0SKQ8_LEPBP | 2.17 | 2.41E-02 | 6.58E-04 | 1.43E-03 | Putative lipoprotein putative signal peptide OS=Leptospira biflexa serovar Patoc (strain Patoc 1 / ATCC 23582 / Paris) OX=456481 GN=LEPBI_I0683 PE=4 SV=1 |
| tr\|B0STR9\|B0STR9_LEPBP | 2.15 | 8.06E-03 | 8.43E-04 | 1.81E-03 | DUF5683 domain-containing protein OS=Leptospira biflexa serovar Patoc (strain Patoc 1 / ATCC 23582 / Paris) OX=456481 GN=LEPBI_II0065 PE=4 SV=1 |
| tr\|B0STJ8\|B0STJ8_LEPBP | 2.14 | 1.74E-02 | 4.90E-02 | 1.05E-01 | Uncharacterized protein OS=Leptospira biflexa serovar Patoc (strain Patoc 1 / ATCC 23582 / Paris) OX=456481 GN=LEPBI_I0267 PE=1 SV=1 |
| **tr\|B0SSZ5\|B0SSZ5_LEPBP** | 2.04 | 9.48E-03 | 2.01E-02 | 4.11E-02 | Flagellin OS=Leptospira biflexa serovar Patoc (strain Patoc 1 / ATCC 23582 / Paris) OX=456481 GN=flaB2 PE=1 SV=1 |
| tr\|B0SS87\|B0SS87_LEPBP | 2.04 | 3.94E-03 | 1.38E-03 | 2.82E-03 | Putative outer membrane protein, OmpA domain putative membrane protein OS=Leptospira biflexa serovar Patoc (strain Patoc 1 / ATCC 23582 / Paris) OX=456481 GN=LEPBI_I1873 PE=4 SV=1 |
| tr\|B0SSX9\|B0SSX9_LEPBP | 2.02 | 2.59E-03 | 3.80E-03 | 7.66E-03 | Uncharacterized protein OS=Leptospira biflexa serovar Patoc (strain Patoc 1 / ATCC 23582 / Paris) OX=456481 GN=LEPBI_I2117 PE=4 SV=1 |
| sp\|B0SSI8\|RL11_LEPBP | 2.00 | 5.42E-03 | 9.59E-04 | 1.92E-03 | 50S ribosomal protein L11 OS=Leptospira biflexa serovar Patoc (strain Patoc 1 / ATCC 23582 / Paris) OX=456481 GN=rplK PE=3 SV=1 |

**Proteins enriched in *L. biflexa flaA2-* vs *wt* (|FC| ≥ 2, p-value ≤ 0.05)**

| Locus | Fold Change | pValue | Signal(Bottom) | Signal(Top) | Description |
|---|---|---|---|---|---|
| tr\|B0STI6\|B0STI6_LEPBP | -17.45 | 2.98E-02 | 1.29E-03 | 7.37E-05 | OmpA-like domain-containing protein OS=Leptospira biflexa serovar Patoc (strain Patoc 1 / ATCC 23582 / Paris) OX=456481 GN=LEPBI_I0255 PE=4 SV=1 |
| sp\|B0SL33\|ILVC_LEPBP | -10.33 | 1.47E-03 | 2.16E-03 | 2.09E-04 | Ketol-acid reductoisomerase (NADP(+)) OS=Leptospira biflexa serovar Patoc (strain Patoc 1 / ATCC 23582 / Paris) OX=456481 GN=ilvC PE=3 SV=1 |

| | | | | | | |
|---|---|---|---|---|---|---|
| tr\|B0SLS3\|B0SLS3_LEPBP | 9 | 12 | 15 | 36 | 3 | ABC-type transport system, periplasmic component OS=Leptospira biflexa serovar Patoc (strain Patoc 1 / ATCC 23582 / Paris) OX=456481 GN=LEPBI_I0848 PE=4 SV=1 |
| tr\|B0SL14\|B0SL14_LEPBP | 12 | 11 | 13 | 36 | 3 | Hydrogenase-4 subunit G OS=Leptospira biflexa serovar Patoc (strain Patoc 1 / ATCC 23582 / Paris) OX=456481 GN=hycG PE=3 SV=1 |
| tr\|B0SLY3\|B0SLY3_LEPBP | 14 | 8 | 14 | 36 | 3 | Putative periplasmic trypsin-like serine protease OS=Leptospira biflexa serovar Patoc (strain Patoc 1 / ATCC 23582 / Paris) OX=456481 GN=LEPBI_I0911 PE=4 SV=1 |
| tr\|B0SJG0\|B0SJG0_LEPBP | 18 | 12 | 6 | 36 | 3 | Uncharacterized protein OS=Leptospira biflexa serovar Patoc (strain Patoc 1 / ATCC 23582 / Paris) OX=456481 GN=LEPBI_I0542 PE=4 SV=1 |
| tr\|B0SLI3\|B0SLI3_LEPBP | 11 | 15 | 8 | 34 | 3 | 2,4-dienoyl-CoA reductase (NADPH) OS=Leptospira biflexa serovar Patoc (strain Patoc 1 / ATCC 23582 / Paris) OX=456481 GN=fadH PE=3 SV=1 |
| sp\|B0SMJ9\|RL28_LEPBP | 12 | 17 | 5 | 34 | 3 | 50S ribosomal protein L28 OS=Leptospira biflexa serovar Patoc (strain Patoc 1 / ATCC 23582 / Paris) OX=456481 GN=rpmB PE=3 SV=1 |
| tr\|B0SU07\|B0SU07_LEPBP | 11 | 17 | 6 | 34 | 3 | DUF4842 domain-containing protein OS=Leptospira biflexa serovar Patoc (strain Patoc 1 / [illegible] |
| tr\|B0SJ68\|B0SJ68_LEPBP | 15 | 13 | 6 | 34 | 3 | Putative TonB-dependent outer membrane receptor putative signal peptide OS=Leptospira biflexa serovar Patoc (strain Patoc 1 / ATCC 23582 / Paris) OX=456481 GN=LEPBI_I0500 PE=4 SV=1 |
| sp\|B0SKZ6\|RL35_LEPBP | 10 | 11 | 12 | 33 | 3 | 50S ribosomal protein L35 OS=Leptospira biflexa serovar Patoc (strain Patoc 1 / ATCC 23582 / Paris) OX=456481 GN=rpmI PE=3 SV=1 |
| tr\|B0SNH8\|B0SNH8_LEPBP | 4 | 8 | 21 | 33 | 3 | GGDEF domain-containing protein OS=Leptospira biflexa serovar Patoc (strain Patoc 1 / ATCC 23582 / Paris) OX=456481 GN=LEPBI_I2876 PE=4 SV=1 |
| tr\|B0SN43\|B0SN43_LEPBP | 12 | 19 | 2 | 33 | 3 | Long-chain acyl-CoA synthetase, AMP-forming (Acyl-CoA synthetase) OS=Leptospira biflexa serovar Patoc (strain Patoc 1 / ATCC 23582 / Paris) OX=456481 GN=fadD2 PE=4 SV=1 |
| tr\|B0SRU5\|B0SRU5_LEPBP | 12 | 15 | 6 | 33 | 3 | Putative outer membrane efflux protein putative signal peptide OS=Leptospira biflexa serovar Patoc (strain Patoc 1 / ATCC 23582 / Paris) OX=456481 GN=LEPBI_I1784 PE=3 SV=1 |
| tr\|B0SQK9\|B0SQK9_LEPBP | 1 | 6 | 26 | 33 | 3 | Uncharacterized protein OS=Leptospira biflexa serovar Patoc (strain Patoc 1 / ATCC 23582 / Paris) OX=456481 GN=LEPBI_I1560 PE=4 SV=1 |
| tr\|B0SMN6\|B0SMN6_LEPBP | 13 | 14 | 6 | 33 | 3 | Cytochrome c oxidase polypeptide III (Cytochrome aa3 subunit 3 Oxidase aa(3) subunit 3) OS=Leptospira biflexa serovar Patoc (strain Patoc 1 / ATCC 23582 / Paris) OX=456481 GN=ctaE PE=3 SV=1 |
| tr\|B0SSW7\|B0SSW7_LEPBP | 15 | 8 | 9 | 32 | 3 | Uncharacterized protein OS=Leptospira biflexa serovar Patoc (strain Patoc 1 / ATCC 23582 / Paris) OX=456481 GN=LEPBI_I2105 PE=4 SV=1 |
| tr\|B0SSQ5\|B0SSQ5_LEPBP | 12 | 7 | 13 | 32 | 3 | Putative acyltransferase OS=Leptospira biflexa serovar Patoc (strain Patoc 1 / ATCC 23582 / Paris) OX=456481 GN=LEPBI_I2043 PE=4 SV=1 |
| tr\|B0SKS5\|B0SKS5_LEPBP | 11 | 16 | 5 | 32 | 3 | GST N-terminal domain-containing protein OS=Leptospira biflexa serovar Patoc (strain Patoc 1 / ATCC 23582 / Paris) OX=456481 GN=LEPBI_I0700 PE=4 SV=1 |
| sp\|B0SRL5\|ENO_LEPBP | 9 | 5 | 18 | 32 | 3 | Enolase OS=Leptospira biflexa serovar Patoc (strain Patoc 1 / ATCC 23582 / Paris) OX=456481 GN=eno PE=3 SV=1 |
| tr\|B0SRF8\|B0SRF8_LEPBP | 16 | 8 | 8 | 32 | 3 | Putative alpha-glucosidase OS=Leptospira biflexa serovar Patoc (strain Patoc 1 / ATCC 23582 / Paris) OX=456481 GN=LEPBI_I3386 PE=4 SV=1 |
| tr\|B0SMW6\|B0SMW6_LEPBP | 10 | 11 | 11 | 32 | 3 | Putative iron-regulated membrane protein putative membrane protein OS=Leptospira biflexa serovar Patoc (strain Patoc 1 / ATCC 23582 / Paris) OX=456481 GN=LEPBI_I2762 PE=4 SV=1 |
| tr\|B0SM55\|B0SM55_LEPBP | 12 | 18 | 1 | 31 | 3 | Putative adenylate/guanylate cyclase putative membrane protein OS=Leptospira biflexa serovar Patoc (strain Patoc 1 / ATCC 23582 / Paris) OX=456481 GN=LEPBI_I2606 PE=4 SV=1 |
| tr\|B0SSV5\|B0SSV5_LEPBP | 10 | 20 | 1 | 31 | 3 | Putative metal-dependent phosphohydrolase, HD subdomain putative membrane protein OS=Leptospira biflexa serovar Patoc (strain Patoc 1 / ATCC 23582 / Paris) OX=456481 GN=LEPBI_I2093 PE=4 SV=1 |
| tr\|B0SQN4\|B0SQN4_LEPBP | 13 | 14 | 4 | 31 | 3 | Uncharacterized protein OS=Leptospira biflexa serovar Patoc (strain Patoc 1 / ATCC 23582 / Paris) OX=456481 GN=LEPBI_I3216 PE=4 SV=1 |
| tr\|B0SPJ2\|B0SPJ2_LEPBP | 10 | 13 | 7 | 30 | 3 | Putative sulfate transporter family protein OS=Leptospira biflexa serovar Patoc (strain Patoc 1 / ATCC 23582 / Paris) OX=456481 GN=LEPBI_I3032 PE=4 SV=1 |
| tr\|B0SLT1\|B0SLT1_LEPBP | 10 | 8 | 12 | 30 | 3 | Uncharacterized protein OS=Leptospira biflexa serovar Patoc (strain Patoc 1 / ATCC 23582 / Paris) OX=456481 GN=LEPBI_I0856 PE=3 SV=1 |
| tr\|B0SQY3\|B0SQY3_LEPBP | 16 | 10 | 4 | 30 | 3 | Acetyl-CoA carboxylase alpha subunit OS=Leptospira biflexa serovar Patoc (strain Patoc 1 / ATCC 23582 / Paris) OX=456481 GN=accA PE=4 SV=1 |
| tr\|B0SQE6\|B0SQE6_LEPBP | 13 | 13 | 4 | 30 | 3 | ABC-type transport system, ATP binding protein OS=Leptospira biflexa serovar Patoc (strain Patoc 1 / ATCC 23582 / Paris) OX=456481 GN=LEPBI_I1497 PE=4 SV=1 |
| tr\|B0SQT2\|B0SQT2_LEPBP | 8 | 7 | 15 | 30 | 3 | Putative glycosyltransferase OS=Leptospira biflexa serovar Patoc (strain Patoc 1 / ATCC 23582 / Paris) OX=456481 GN=LEPBI_I3264 PE=4 SV=1 |
| tr\|B0SU06\|B0SU06_LEPBP | 14 | 14 | 2 | 30 | 3 | Demethylmenaquinone methyltransferase OS=Leptospira biflexa serovar Patoc (strain Patoc 1 / ATCC 23582 / Paris) OX=456481 GN=ubiE PE=3 SV=1 |
| tr\|B0STI5\|B0STI5_LEPBP | 9 | 8 | 12 | 29 | 3 | Sigma-54 factor interaction domain-containing protein OS=Leptospira biflexa serovar Patoc (strain Patoc 1 / ATCC 23582 / Paris) OX=456481 GN=LEPBI_I0254 PE=4 SV=1 |
| tr\|B0SQH1\|B0SQH1_LEPBP | 11 | 5 | 13 | 29 | 3 | Uncharacterized protein OS=Leptospira biflexa serovar Patoc (strain Patoc 1 / ATCC 23582 / Paris) OX=456481 GN=LEPBI_I1522 PE=4 SV=1 |
| tr\|B0SJP3\|B0SJP3_LEPBP | 17 | 8 | 4 | 29 | 3 | ABC transporter, atp-binding protein putative membrane protein OS=Leptospira biflexa serovar Patoc (strain Patoc 1 / ATCC 23582 / Paris) OX=456481 GN=LEPBI_I0059 PE=4 SV=1 |
| tr\|B0STR7\|B0STR7_LEPBP | 12 | 12 | 5 | 29 | 3 | Putative cytochrome oxidase subunit II OS=Leptospira biflexa serovar Patoc (strain Patoc 1 / ATCC 23582 / Paris) OX=456481 GN=LEPBI_II0063 PE=4 SV=1 |
| tr\|B0SQ04\|B0SQ04_LEPBP | 6 | 9 | 13 | 28 | 3 | Putative ribosomal subunit pseudouridine synthase OS=Leptospira biflexa serovar Patoc (strain Patoc 1 / ATCC 23582 / Paris) OX=456481 GN=LEPBI_I1463 PE=4 SV=1 |
| tr\|B0SPS5\|B0SPS5_LEPBP | 6 | 13 | 9 | 28 | 3 | Enoyl-CoA hydratase OS=Leptospira biflexa serovar Patoc (strain Patoc 1 / ATCC 23582 / Paris) OX=456481 GN=fadB1 PE=3 SV=1 |
| tr\|B0SML1\|B0SML1_LEPBP | 9 | 11 | 8 | 28 | 3 | Putative betaine aldehyde dehydrogenase OS=Leptospira biflexa serovar Patoc (strain Patoc 1 / ATCC 23582 / Paris) OX=456481 GN=LEPBI_I2656 PE=3 SV=1 |
| tr\|B0SSV8\|B0SSV8_LEPBP | 10 | 5 | 13 | 28 | 3 | Replicative DNA helicase OS=Leptospira biflexa serovar Patoc (strain Patoc 1 / ATCC 23582 / Paris) OX=456481 GN=dnaB PE=3 SV=1 |
| tr\|B0SRW7\|B0SRW7_LEPBP | 4 | 7 | 17 | 28 | 3 | Putative proton glutamate symport protein putative membrane protein OS=Leptospira biflexa serovar Patoc (strain Patoc 1 / ATCC 23582 / Paris) OX=456481 GN=gltP PE=4 SV=1 |
| tr\|B0SSJ4\|B0SSJ4_LEPBP | 11 | 5 | 12 | 28 | 3 | ADP-L-glycero-D-manno-heptose-6-epimerase OS=Leptospira biflexa serovar Patoc (strain Patoc 1 / ATCC 23582 / Paris) OX=456481 GN=hldD PE=3 SV=1 |
| tr\|B0SSU7\|B0SSU7_LEPBP | 13 | 6 | 8 | 27 | 3 | Histidine kinase sensor protein OS=Leptospira biflexa serovar Patoc (strain Patoc 1 / ATCC 23582 / Paris) OX=456481 GN=LEPBI_Ia2079 PE=4 SV=1 |
| tr\|B0SKT7\|B0SKT7_LEPBP | 5 | 20 | 2 | 27 | 3 | Putative cAMP-binding protein OS=Leptospira biflexa serovar Patoc (strain Patoc 1 / ATCC 23582 / Paris) OX=456481 GN=LEPBI_I2338 PE=4 SV=1 |
| tr\|B0SK98\|B0SK98_LEPBP | 14 | 11 | 2 | 27 | 3 | Putative sodium/hydrogen exchanger putative membrane protein OS=Leptospira biflexa serovar Patoc (strain Patoc 1 / ATCC 23582 / Paris) OX=456481 GN=LEPBI_I0200 PE=4 SV=1 |
| tr\|B0SJG2\|B0SJG2_LEPBP | 10 | 8 | 8 | 26 | 3 | Holliday junction ATP-dependent DNA helicase RuvB OS=Leptospira biflexa serovar Patoc (strain Patoc 1 / ATCC 23582 / Paris) OX=456481 GN=ruvB PE=3 SV=1 |
| tr\|B0SQN1\|B0SQN1_LEPBP | 11 | 8 | 6 | 25 | 3 | Flagellar hook-associated protein 1 OS=Leptospira biflexa serovar Patoc (strain Patoc 1 / ATCC 23582 / Paris) OX=456481 GN=flgK PE=3 SV=1 |
| tr\|B0SSI9\|B0SSI9_LEPBP | 7 | 17 | 1 | 25 | 3 | Transcription termination/antitermination protein NusG OS=Leptospira biflexa serovar Patoc (strain Patoc 1 / ATCC 23582 / Paris) OX=456481 GN=nusG PE=3 SV=1 |
| tr\|B0SKS1\|B0SKS1_LEPBP | 2 | 6 | 17 | 25 | 3 | Putative aminotransferase, DegT/DnrJ/EryC1/StrS family OS=Leptospira biflexa serovar Patoc (strain Patoc 1 / ATCC 23582 / Paris) OX=456481 GN=LEPBI_I0696 PE=3 SV=1 |
| tr\|B0SJI2\|B0SJI2_LEPBP | 11 | 10 | 4 | 25 | 3 | Putative protease (U32 family) OS=Leptospira biflexa serovar Patoc (strain Patoc 1 / ATCC 23582 / Paris) OX=456481 GN=LEPBI_I0392 PE=4 SV=1 |
| tr\|B0SQL7\|B0SQL7_LEPBP | 7 | 5 | 13 | 25 | 3 | Uncharacterized protein OS=Leptospira biflexa serovar Patoc (strain Patoc 1 / ATCC 23582 / Paris) OX=456481 GN=LEPBI_I1568 PE=4 SV=1 |
| tr\|B0SQW9\|B0SQW9_LEPBP | 10 | 12 | 3 | 25 | 3 | AB hydrolase-1 domain-containing protein OS=Leptospira biflexa serovar Patoc (strain Patoc 1 / ATCC 23582 / Paris) OX=456481 GN=LEPBI_I3301 PE=4 SV=1 |
| tr\|B0SJN5\|B0SJN5_LEPBP | 2 | 6 | 17 | 25 | 3 | Peptidase_M23 domain-containing protein OS=Leptospira biflexa serovar Patoc (strain Patoc 1 / ATCC 23582 / Paris) OX=456481 GN=LEPBI_I0051 PE=4 SV=1 |
| tr\|B0SSD6\|B0SSD6_LEPBP | 7 | 12 | 5 | 24 | 3 | Phosphoribosylformylglycinamidine synthase subunit PurQ OS=Leptospira biflexa serovar Patoc (strain Patoc 1 / ATCC 23582 / Paris) OX=456481 GN=purQ PE=3 SV=1 |
| sp\|B0SR12\|ARLY_LEPBP | 11 | 10 | 3 | 24 | 3 | Argininosuccinate lyase OS=Leptospira biflexa serovar Patoc (strain Patoc 1 / ATCC 23582 / Paris) OX=456481 GN=argH PE=3 SV=1 |
| tr\|B0SLR4\|B0SLR4_LEPBP | 13 | 10 | 1 | 24 | 3 | Putative glycosyl transferase OS=Leptospira biflexa serovar Patoc (strain Patoc 1 / ATCC 23582 / Paris) OX=456481 GN=LEPBI_I0839 PE=4 SV=1 |
| tr\|B0SJK1\|B0SJK1_LEPBP | 3 | 7 | 14 | 24 | 3 | ABC-type transport system, Phosphate-transporting ATPase OS=Leptospira biflexa serovar Patoc (strain Patoc 1 / ATCC 23582 / Paris) OX=456481 GN=dppD PE=4 SV=1 |
| tr\|B0SKG6\|B0SKG6_LEPBP | 2 | 15 | 7 | 24 | 3 | Bifunctional GlmU protein: UDP-N-acetylglucosamine pyrophosphorylase (N-acetylglucosamine-1-phosphate uridyltransferase)/Glucosamine-1-phosphate N-acetyltransferase OS=Leptospira biflexa serovar Patoc (strain Patoc 1 / ATCC 23582 / Paris) OX=456481 GN=glmU PE=4 SV=1 |

| | | | | | |
|---|---|---|---|---|---|
| tr\|B0SKU7\|B0SKU7_LEPBP | -9.84 | 4.70E-03 | 2.11E-03 | 2.14E-04 | Iron-sulfur cluster carrier protein OS=Leptospira biflexa serovar Patoc (strain Patoc 1 / ATCC 23582 / Paris) OX=456481 GN=mrp PE=3 SV=1 |
| tr\|B0SRH3\|B0SRH3_LEPBP | -9.53 | 1.04E-02 | 2.58E-03 | 2.71E-04 | Thiolase I, 3-ketoacyl-CoA thiolase beta subunit of the fatty acid oxidation multienzyme complex OS=Leptospira biflexa serovar Patoc (strain Patoc 1 / ATCC 23582 / Paris) OX=456481 GN=fadA1 PE=3 SV=1 |
| sp\|B0SLC6\|ATPA_LEPBP | -9.33 | 8.02E-04 | 1.10E-02 | 1.17E-03 | ATP synthase subunit alpha OS=Leptospira biflexa serovar Patoc (strain Patoc 1 / ATCC 23582 / Paris) OX=456481 GN=atpA PE=3 SV=1 |
| tr\|B0SNK0\|B0SNK0_LEPBP | -9.29 | 4.68E-03 | 2.60E-03 | 2.80E-04 | Protoporphyrinogen oxidase (PPO) OS=Leptospira biflexa serovar Patoc (strain Patoc 1 / ATCC 23582 / Paris) OX=456481 GN=hemG PE=4 SV=1 |
| sp\|B0SLC8\|ATPB_LEPBP | -8.91 | 1.00E-05 | 9.39E-03 | 1.05E-03 | ATP synthase subunit beta OS=Leptospira biflexa serovar Patoc (strain Patoc 1 / ATCC 23582 / Paris) OX=456481 GN=atpD PE=3 SV=1 |
| tr\|B0SSD4\|B0SSD4_LEPBP | -8.90 | 9.91E-04 | 6.56E-04 | 7.37E-05 | Putative diguanylate phosphodiesterase OS=Leptospira biflexa serovar Patoc (strain Patoc 1 / ATCC 23582 / Paris) OX=456481 GN=LEPBI_I1921 PE=4 SV=1 |
| sp\|B0SRX5\|ACSA_LEPBP | -8.67 | 1.57E-02 | 1.35E-03 | 1.55E-04 | Acetyl-coenzyme A synthetase OS=Leptospira biflexa serovar Patoc (strain Patoc 1 / ATCC [illegible] |
| tr\|B0SKU8\|B0SKU8_LEPBP | -8.60 | 2.65E-03 | 1.36E-03 | 1.57E-04 | Putative metallo-dependent phosphatase OS=Leptospira biflexa serovar Patoc (strain Patoc 1 / ATCC 23582 / Paris) OX=456481 GN=LEPBI_I2350 PE=4 SV=1 |
| tr\|B0SN73\|B0SN73_LEPBP | -8.56 | 5.85E-03 | 1.64E-03 | 1.92E-04 | Thioredoxin domain-containing protein OS=Leptospira biflexa serovar Patoc (strain Patoc 1 / ATCC 23582 / Paris) OX=456481 GN=LEPBI_I1143 PE=4 SV=1 |
| tr\|B0SJ48\|B0SJ48_LEPBP | -8.24 | 3.00E-02 | 1.58E-03 | 1.92E-04 | ABC_transp_aux domain-containing protein OS=Leptospira biflexa serovar Patoc (strain Patoc 1 / ATCC 23582 / Paris) OX=456481 GN=LEPBI_I0506 PE=4 SV=1 |
| sp\|B0SRF0\|DNAJ_LEPBP | -8.21 | 1.74E-03 | 9.08E-04 | 1.11E-04 | Chaperone protein DnaJ OS=Leptospira biflexa serovar Patoc (strain Patoc 1 / ATCC 23582 / Paris) OX=456481 GN=dnaJ PE=3 SV=1 |
| **tr\|B0STI0\|B0STI0_LEPBP** | -8.10 | 1.11E-02 | 4.18E-03 | 5.16E-04 | Putative ATPase OS=Leptospira biflexa serovar Patoc (strain Patoc 1 / ATCC 23582 / Paris) OX=456481 GN=LEPBI_I2327 PE=4 SV=1 |
| tr\|B0SSX3\|B0SSX3_LEPBP | -7.88 | 5.38E-03 | 1.49E-03 | 1.90E-04 | Ribosome-binding ATPase YchF OS=Leptospira biflexa serovar Patoc (strain Patoc 1 / ATCC 23582 / Paris) OX=456481 GN=ychF PE=3 SV=1 |
| tr\|B0SPR2\|B0SPR2_LEPBP | -7.57 | 1.07E-02 | 2.61E-03 | 3.45E-04 | Putative molybdopterin oxidoreductase, iron-sulfur binding subunit OS=Leptospira biflexa serovar Patoc (strain Patoc 1 / ATCC 23582 / Paris) OX=456481 GN=LEPBI_I1369 PE=4 SV=1 |
| tr\|B0SP59\|B0SP59_LEPBP | -7.53 | 9.25E-03 | 2.26E-03 | 3.01E-04 | Inosine-5'-monophosphate dehydrogenase OS=Leptospira biflexa serovar Patoc (strain Patoc 1 / ATCC 23582 / Paris) OX=456481 GN=guaB PE=3 SV=1 |
| tr\|B0SN68\|B0SN68_LEPBP | -7.53 | 1.87E-03 | 6.39E-04 | 8.49E-05 | Putative acyl-CoA dehydrogenase, short-chain specific (SCAD Butyryl-CoA dehydrogenase) OS=Leptospira biflexa serovar Patoc (strain Patoc 1 / ATCC 23582 / Paris) OX=456481 GN=LEPBI_I1138 PE=3 SV=1 |
| tr\|B0SRL0\|B0SRL0_LEPBP | -7.13 | 2.35E-02 | 2.14E-03 | 3.00E-04 | Succinate dehydrogenase iron-sulfur subunit SdhB OS=Leptospira biflexa serovar Patoc (strain Patoc 1 / ATCC 23582 / Paris) OX=456481 GN=sdhB PE=3 SV=1 |
| sp\|B0SM57\|TRPB_LEPBP | -7.11 | 7.92E-04 | 3.11E-03 | 4.38E-04 | Tryptophan synthase beta chain OS=Leptospira biflexa serovar Patoc (strain Patoc 1 / ATCC 23582 / Paris) OX=456481 GN=trpB PE=3 SV=1 |
| tr\|B0STK6\|B0STK6_LEPBP | -7.02 | 1.29E-03 | 1.11E-03 | 1.57E-04 | Putative chromosome partitioning protein ParB OS=Leptospira biflexa serovar Patoc (strain Patoc 1 / ATCC 23582 / Paris) OX=456481 GN=LEPBI_II0002 PE=3 SV=1 |
| tr\|B0SKM2\|B0SKM2_LEPBP | -6.96 | 1.14E-04 | 1.99E-03 | 2.85E-04 | Octoprenyl-diphosphate synthase (Octaprenyl pyrophosphate synthetase OPP synthetase) OS=Leptospira biflexa serovar Patoc (strain Patoc 1 / ATCC 23582 / Paris) OX=456481 GN=ispA2 PE=3 SV=1 |
| tr\|B0SLB3\|B0SLB3_LEPBP | -6.77 | 5.57E-04 | 9.01E-04 | 1.33E-04 | Signal peptidase I OS=Leptospira biflexa serovar Patoc (strain Patoc 1 / ATCC 23582 / Paris) OX=456481 GN=lepB2 PE=3 SV=1 |
| sp\|B0SRF1\|DNAK_LEPBP | -6.63 | 4.78E-03 | 2.66E-03 | 4.01E-04 | Chaperone protein DnaK OS=Leptospira biflexa serovar Patoc (strain Patoc 1 / ATCC 23582 / Paris) OX=456481 GN=dnaK PE=3 SV=1 |
| tr\|B0SR96\|B0SR96_LEPBP | -6.63 | 9.31E-03 | 1.63E-03 | 2.46E-04 | Putative arylesterase OS=Leptospira biflexa serovar Patoc (strain Patoc 1 / ATCC 23582 / Paris) OX=456481 GN=LEPBI_I3323 PE=4 SV=1 |
| tr\|B0SNZ9\|B0SNZ9_LEPBP | -6.62 | 5.41E-03 | 2.84E-03 | 4.29E-04 | Aldehyde dehydrogenase OS=Leptospira biflexa serovar Patoc (strain Patoc 1 / ATCC 23582 / Paris) OX=456481 GN=LEPBI_I2943 PE=3 SV=1 |
| tr\|B0STX1\|B0STX1_LEPBP | -6.56 | 4.93E-04 | 1.79E-03 | 2.73E-04 | Putative UTP--glucose-1-phosphate uridylyltransferase OS=Leptospira biflexa serovar Patoc (strain Patoc 1 / ATCC 23582 / Paris) OX=456481 GN=galU2 PE=4 SV=1 |
| tr\|B0SN64\|B0SN64_LEPBP | -6.39 | 4.53E-03 | 3.07E-03 | 4.80E-04 | Electron transfer flavoprotein beta-subunit (Beta-ETF Electron transfer flavoprotein small subunit) (ETFSS) OS=Leptospira biflexa serovar Patoc (strain Patoc 1 / ATCC 23582 / Paris) OX=456481 GN=etfB PE=4 SV=1 |
| sp\|B0SMF0\|CLPX_LEPBP | -6.32 | 5.73E-03 | 1.50E-03 | 2.37E-04 | ATP-dependent Clp protease ATP-binding subunit ClpX OS=Leptospira biflexa serovar Patoc (strain Patoc 1 / ATCC 23582 / Paris) OX=456481 GN=clpX PE=3 SV=1 |
| tr\|B0SMT5\|B0SMT5_LEPBP | -6.31 | 1.44E-03 | 1.04E-03 | 1.65E-04 | ABC-type transport system, ATP binding protein OS=Leptospira biflexa serovar Patoc (strain Patoc 1 / ATCC 23582 / Paris) OX=456481 GN=LEPBI_I2731 PE=4 SV=1 |
| tr\|B0STL3\|B0STL3_LEPBP | -6.22 | 7.46E-03 | 2.17E-03 | 3.49E-04 | Heat shock protein HtpG OS=Leptospira biflexa serovar Patoc (strain Patoc 1 / ATCC 23582 / Paris) OX=456481 GN=htpG PE=3 SV=1 |
| tr\|B0SML4\|B0SML4_LEPBP | -6.20 | 1.01E-03 | 2.52E-03 | 4.06E-04 | Uncharacterized protein OS=Leptospira biflexa serovar Patoc (strain Patoc 1 / ATCC 23582 / Paris) OX=456481 GN=LEPBI_I2660 PE=4 SV=1 |
| tr\|B0SKX3\|B0SKX3_LEPBP | -6.18 | 1.12E-02 | 2.42E-03 | 3.92E-04 | Hemolysin putative signal peptide OS=Leptospira biflexa serovar Patoc (strain Patoc 1 / ATCC 23582 / Paris) OX=456481 GN=LEPBI_I2375 PE=4 SV=1 |
| tr\|B0SPJ5\|B0SPJ5_LEPBP | -6.17 | 2.38E-05 | 1.62E-03 | 2.63E-04 | Alginate_exp domain-containing protein OS=Leptospira biflexa serovar Patoc (strain Patoc 1 / ATCC 23582 / Paris) OX=456481 GN=LEPBI_I3035 PE=4 SV=1 |
| sp\|B0SSG2\|RL18_LEPBP | -6.01 | 1.87E-02 | 6.36E-04 | 1.06E-04 | 50S ribosomal protein L18 OS=Leptospira biflexa serovar Patoc (strain Patoc 1 / ATCC 23582 / Paris) OX=456481 GN=rplR PE=3 SV=1 |
| sp\|B0SSF1\|RL17_LEPBP | -5.96 | 2.99E-02 | 2.65E-03 | 4.45E-04 | 50S ribosomal protein L17 OS=Leptospira biflexa serovar Patoc (strain Patoc 1 / ATCC 23582 / Paris) OX=456481 GN=rplQ PE=3 SV=1 |
| tr\|B0STM5\|B0STM5_LEPBP | -5.91 | 4.47E-04 | 1.38E-03 | 2.34E-04 | NADH dehydrogenase OS=Leptospira biflexa serovar Patoc (strain Patoc 1 / ATCC 23582 / Paris) OX=456481 GN=ndh PE=4 SV=1 |
| tr\|B0SS56\|B0SS56_LEPBP | -5.69 | 8.53E-05 | 2.02E-03 | 3.54E-04 | Putative glycosidase OS=Leptospira biflexa serovar Patoc (strain Patoc 1 / ATCC 23582 / Paris) OX=456481 GN=LEPBI_I1840 PE=3 SV=1 |
| tr\|B0SKQ4\|B0SKQ4_LEPBP | -5.67 | 2.67E-02 | 1.25E-03 | 2.21E-04 | DNA topoisomerase (ATP-hydrolyzing) OS=Leptospira biflexa serovar Patoc (strain Patoc 1 / ATCC 23582 / Paris) OX=456481 GN=gyrA2 PE=4 SV=1 |
| tr\|B0SSZ8\|B0SSZ8_LEPBP | -5.48 | 2.20E-02 | 2.13E-03 | 3.88E-04 | ATP-dependent RNA helicase OS=Leptospira biflexa serovar Patoc (strain Patoc 1 / ATCC 23582 / Paris) OX=456481 GN=LEPBI_I2136 PE=3 SV=1 |
| tr\|B0SQ75\|B0SQ75_LEPBP | -5.44 | 1.37E-02 | 8.87E-04 | 1.63E-04 | Uncharacterized protein OS=Leptospira biflexa serovar Patoc (strain Patoc 1 / ATCC 23582 / Paris) OX=456481 GN=LEPBI_I3162 PE=4 SV=1 |
| tr\|B0SJF6\|B0SJF6_LEPBP | -5.42 | 3.73E-04 | 5.41E-03 | 9.99E-04 | Putative FAD-linked oxidase OS=Leptospira biflexa serovar Patoc (strain Patoc 1 / ATCC 23582 / Paris) OX=456481 GN=LEPBI_I0538 PE=4 SV=1 |
| tr\|B0SR90\|B0SR90_LEPBP | -5.40 | 8.62E-05 | 2.98E-03 | 5.51E-04 | Acyl-CoA dehydrogenase OS=Leptospira biflexa serovar Patoc (strain Patoc 1 / ATCC 23582 / Paris) OX=456481 GN=acdA3 PE=3 SV=1 |
| tr\|B0SSS3\|B0SSS3_LEPBP | -5.37 | 1.38E-02 | 9.48E-04 | 1.77E-04 | Uncharacterized protein OS=Leptospira biflexa serovar Patoc (strain Patoc 1 / ATCC 23582 / Paris) OX=456481 GN=LEPBI_I2061 PE=4 SV=1 |
| tr\|B0SND9\|B0SND9_LEPBP | -5.36 | 7.62E-03 | 1.30E-03 | 2.43E-04 | Uncharacterized protein OS=Leptospira biflexa serovar Patoc (strain Patoc 1 / ATCC 23582 / Paris) OX=456481 GN=LEPBI_I2835 PE=4 SV=1 |
| tr\|B0SQ39\|B0SQ39_LEPBP | -5.29 | 3.27E-03 | 3.09E-03 | 5.83E-04 | Putative sodium-dependent phosphate transport protein OS=Leptospira biflexa serovar Patoc (strain Patoc 1 / ATCC 23582 / Paris) OX=456481 GN=LEPBI_I3125 PE=4 SV=1 |
| tr\|B0SQ34\|B0SQ34_LEPBP | -5.29 | 1.51E-02 | 1.14E-03 | 2.16E-04 | Putative cytochrome c peroxidase putative signal peptide OS=Leptospira biflexa serovar Patoc (strain Patoc 1 / ATCC 23582 / Paris) OX=456481 GN=LEPBI_I3120 PE=4 SV=1 |
| sp\|B0SRP1\|GATA_LEPBP | -5.12 | 7.68E-03 | 7.54E-04 | 1.47E-04 | Glutamyl-tRNA(Gln) amidotransferase subunit A OS=Leptospira biflexa serovar Patoc (strain Patoc 1 / ATCC 23582 / Paris) OX=456481 GN=gatA PE=3 SV=1 |
| sp\|B0SSI1\|RS7_LEPBP | -5.11 | 2.56E-02 | 4.99E-03 | 9.76E-04 | 30S ribosomal protein S7 OS=Leptospira biflexa serovar Patoc (strain Patoc 1 / ATCC 23582 / Paris) OX=456481 GN=rpsG PE=3 SV=1 |
| tr\|B0SR09\|B0SR09_LEPBP | -5.10 | 2.53E-02 | 1.19E-03 | 2.34E-04 | PHB domain-containing protein OS=Leptospira biflexa serovar Patoc (strain Patoc 1 / ATCC 23582 / Paris) OX=456481 GN=LEPBI_I1603 PE=3 SV=1 |
| tr\|B0SMI0\|B0SMI0_LEPBP | -5.00 | 3.65E-03 | 1.50E-03 | 3.00E-04 | Glycosyltransferase OS=Leptospira biflexa serovar Patoc (strain Patoc 1 / ATCC 23582 / Paris) OX=456481 GN=LEPBI_I1000 PE=4 SV=1 |
| tr\|B0SRM6\|B0SRM6_LEPBP | -4.93 | 1.19E-04 | 3.57E-03 | 7.24E-04 | Uncharacterized protein OS=Leptospira biflexa serovar Patoc (strain Patoc 1 / ATCC 23582 / Paris) OX=456481 GN=LEPBI_I1715 PE=4 SV=1 |
| tr\|B0SSM2\|B0SSM2_LEPBP | -4.91 | 2.84E-03 | 1.16E-03 | 2.36E-04 | Putative glycosyltransferase OS=Leptospira biflexa serovar Patoc (strain Patoc 1 / ATCC 23582 / Paris) OX=456481 GN=LEPBI_I2010 PE=4 SV=1 |
| tr\|B0SP47\|B0SP47_LEPBP | -4.89 | 6.97E-04 | 1.42E-03 | 2.91E-04 | Acyl-CoA dehydrogenase family protein OS=Leptospira biflexa serovar Patoc (strain Patoc 1 / ATCC 23582 / Paris) OX=456481 GN=LEPBI_I1259 PE=3 SV=1 |
| tr\|B0STX2\|B0STX2_LEPBP | -4.80 | 6.45E-05 | 1.64E-03 | 3.42E-04 | Putative UDP-glucose 4-epimerase OS=Leptospira biflexa serovar Patoc (strain Patoc 1 / ATCC 23582 / Paris) OX=456481 GN=LEPBI_II0120 PE=4 SV=1 |
| tr\|B0SLW3\|B0SLW3_LEPBP | -4.79 | 2.37E-02 | 1.68E-03 | 3.52E-04 | Glycine--tRNA ligase OS=Leptospira biflexa serovar Patoc (strain Patoc 1 / ATCC 23582 / Paris) OX=456481 GN=glyQS PE=3 SV=1 |
| tr\|B0SMQ9\|B0SMQ9_LEPBP | -4.75 | 1.06E-02 | 6.32E-04 | 1.33E-04 | Uncharacterized protein OS=Leptospira biflexa serovar Patoc (strain Patoc 1 / ATCC 23582 / Paris) OX=456481 GN=LEPBI_I2705 PE=4 SV=1 |

| | | | | | | |
|---|---|---|---|---|---|---|
| tr\|B0STX4\|B0STX4_LEPBP | 10 | 6 | 8 | 24 | 3 | Uncharacterized protein OS=Leptospira biflexa serovar Patoc (strain Patoc 1 / ATCC 23582 / Paris) OX=456481 GN=LEPBI_II0122 PE=4 SV=1 |
| tr\|B0SQE8\|B0SQE8_LEPBP | 9 | 5 | 9 | 23 | 3 | Mrr_cat domain-containing protein OS=Leptospira biflexa serovar Patoc (strain Patoc 1 / ATCC 23582 / Paris) OX=456481 GN=LEPBI_I1499 PE=4 SV=1 |
| tr\|B0SS24\|B0SS24_LEPBP | 6 | 12 | 5 | 23 | 3 | Uncharacterized protein OS=Leptospira biflexa serovar Patoc (strain Patoc 1 / ATCC 23582 / Paris) OX=456481 GN=LEPBI_I1808 PE=4 SV=1 |
| tr\|B0SQV2\|B0SQV2_LEPBP | 9 | 5 | 9 | 23 | 3 | Uncharacterized protein OS=Leptospira biflexa serovar Patoc (strain Patoc 1 / ATCC 23582 / Paris) OX=456481 GN=LEPBI_I3284 PE=4 SV=1 |
| tr\|B0SQL9\|B0SQL9_LEPBP | 6 | 10 | 7 | 23 | 3 | Tyrosine--tRNA ligase OS=Leptospira biflexa serovar Patoc (strain Patoc 1 / ATCC 23582 / Paris) OX=456481 GN=tyrS PE=3 SV=1 |
| tr\|B0SP91\|B0SP91_LEPBP | 5 | 10 | 7 | 22 | 3 | NADH:ubiquinone reductase (H(+)-translocating) OS=Leptospira biflexa serovar Patoc (strain Patoc 1 / ATCC 23582 / Paris) OX=456481 GN=nuoL PE=3 SV=1 |
| tr\|B0SPU3\|B0SPU3_LEPBP | 13 | 8 | 1 | 22 | 3 | Uncharacterized protein OS=Leptospira biflexa serovar Patoc (strain Patoc 1 / ATCC 23582 / Paris) OX=456481 GN=LEPBI_I1400 PE=4 SV=1 |
| tr\|B0SKW7\|B0SKW7_LEPBP | 7 | 7 | 8 | 22 | 3 | Uncharacterized protein OS=Leptospira biflexa serovar Patoc (strain Patoc 1 / ATCC 23582 / Paris) OX=456481 GN=LEPBI_I2369 PE=4 SV=1 |
| tr\|B0ST04\|B0ST04_LEPBP | 9 | 11 | 2 | 22 | 3 | Putative SET domain-containing protein OS=Leptospira biflexa serovar Patoc (strain Patoc 1 / ATCC 23582 / Paris) OX=456481 GN=LEPBI_I2142 PE=4 SV=1 |
| tr\|B0SL42\|B0SL42_LEPBP | 10 | 9 | 3 | 22 | 3 | Putative hemolysin-related membrane protein with CBS regulatory domain OS=Leptospira biflexa serovar Patoc (strain Patoc 1 / ATCC 23582 / Paris) OX=456481 GN=LEPBI_I0717 PE=3 SV=1 |
| tr\|B0STR5\|B0STR5_LEPBP | 10 | 7 | 4 | 21 | 3 | Putative cytochrome c oxidase, subunit III OS=Leptospira biflexa serovar Patoc (strain Patoc 1 / ATCC 23582 / Paris) OX=456481 GN=LEPBI_II0061 PE=4 SV=1 |
| sp\|B0SPC6\|MNMA_LEPBP | 6 | 5 | 10 | 21 | 3 | tRNA-specific 2-thiouridylase MnmA OS=Leptospira biflexa serovar Patoc (strain Patoc 1 / ATCC 23582 / Paris) OX=456481 GN=mnmA PE=3 SV=1 |
| tr\|B0SJ86\|B0SJ86_LEPBP | 3 | 8 | 10 | 21 | 3 | Flagellar hook-associated protein 2 OS=Leptospira biflexa serovar Patoc (strain Patoc 1 / ATCC 23582 / Paris) OX=456481 GN=fliD PE=3 SV=1 |
| tr\|B0STD2\|B0STD2_LEPBP | 9 | 8 | 4 | 21 | 3 | Short-chain dehydrogenase OS=Leptospira biflexa serovar Patoc (strain Patoc 1 / ATCC 23582 / Paris) OX=456481 GN=LEPBI_I2277 PE=4 SV=1 |
| tr\|B0SPS8\|B0SPS8_LEPBP | 6 | 6 | 9 | 21 | 3 | Putative membrane associated metalloendopeptidase OS=Leptospira biflexa serovar Patoc (strain Patoc 1 / ATCC 23582 / Paris) OX=456481 GN=LEPBI_I1385 PE=4 SV=1 |
| tr\|B0SRU9\|B0SRU9_LEPBP | 10 | 4 | 7 | 21 | 3 | Flagellar motor switch protein FliG OS=Leptospira biflexa serovar Patoc (strain Patoc 1 / ATCC 23582 / Paris) OX=456481 GN=fliG2 PE=3 SV=1 |
| tr\|B0SUD0\|B0SUD0_LEPBP | 9 | 10 | 2 | 21 | 3 | Putative transcriptional regulator OS=Leptospira biflexa serovar Patoc (strain Patoc 1 / ATCC 23582 / Paris) OX=456481 GN=LEPBI_II0281 PE=4 SV=1 |
| tr\|B0SSJ3\|B0SSJ3_LEPBP | 4 | 14 | 3 | 21 | 3 | Putative transcriptional regulator, ArsR family OS=Leptospira biflexa serovar Patoc (strain Patoc 1 / ATCC 23582 / Paris) OX=456481 GN=LEPBI_I1981 PE=4 SV=1 |
| tr\|B0SJ65\|B0SJ65_LEPBP | 11 | 6 | 3 | 20 | 3 | Putative penicillin-binding protein, transpeptidase OS=Leptospira biflexa serovar Patoc (strain Patoc 1 / ATCC 23582 / Paris) OX=456481 GN=LEPBI_I0497 PE=4 SV=1 |
| sp\|B0SU50\|EFP_LEPBP | 6 | 7 | 7 | 20 | 3 | Elongation factor P OS=Leptospira biflexa serovar Patoc (strain Patoc 1 / ATCC 23582 / Paris) OX=456481 GN=efp PE=3 SV=1 |
| tr\|B0SPB8\|B0SPB8_LEPBP | 9 | 5 | 6 | 20 | 3 | 3-deoxy-D-manno-octulosonic acid transferase OS=Leptospira biflexa serovar Patoc (strain Patoc 1 / ATCC 23582 / Paris) OX=456481 GN=LEPBI_I1332 PE=3 SV=1 |
| tr\|B0SPX2\|B0SPX2_LEPBP | 12 | 6 | 2 | 20 | 3 | Putative metalloendopeptidase OS=Leptospira biflexa serovar Patoc (strain Patoc 1 / ATCC 23582 / Paris) OX=456481 GN=LEPBI_I1430 PE=4 SV=1 |
| tr\|B0SP11\|B0SP11_LEPBP | 8 | 5 | 7 | 20 | 3 | Nitric oxide reductase, cytochrome c-containing subunit OS=Leptospira biflexa serovar Patoc (strain Patoc 1 / ATCC 23582 / Paris) OX=456481 GN=norC PE=4 SV=1 |
| tr\|B0SUH3\|B0SUH3_LEPBP | 10 | 5 | 4 | 19 | 3 | Uncharacterized protein OS=Leptospira biflexa serovar Patoc (strain Patoc 1 / ATCC 23582 / Paris) OX=456481 GN=LEPBI_p0045 PE=4 SV=1 |
| tr\|B0SQF8\|B0SQF8_LEPBP | 7 | 10 | 2 | 19 | 3 | Putative methyl-accepting chemotaxis protein putative membrane protein OS=Leptospira biflexa serovar Patoc (strain Patoc 1 / ATCC 23582 / Paris) OX=456481 GN=LEPBI_I1509 PE=4 SV=1 |
| tr\|B0SUA1\|B0SUA1_LEPBP | 3 | 5 | 11 | 19 | 3 | S-adenosylmethionine:tRNA ribosyltransferase-isomerase OS=Leptospira biflexa serovar Patoc (strain Patoc 1 / ATCC 23582 / Paris) OX=456481 GN=queA PE=3 SV=1 |
| tr\|B0SSV6\|B0SSV6_LEPBP | 9 | 5 | 5 | 19 | 3 | Phosphate starvation-inducible protein (PhoH-like) OS=Leptospira biflexa serovar Patoc (strain Patoc 1 / ATCC 23582 / Paris) OX=456481 GN=phoH PE=4 SV=1 |
| tr\|B0SK85\|B0SK85_LEPBP | 5 | 9 | 5 | 19 | 3 | Aspartokinase OS=Leptospira biflexa serovar Patoc (strain Patoc 1 / ATCC 23582 / Paris) OX=456481 GN=lysC PE=3 SV=1 |
| tr\|B0SKI7\|B0SKI7_LEPBP | 3 | 11 | 5 | 19 | 3 | Uncharacterized protein OS=Leptospira biflexa serovar Patoc (strain Patoc 1 / ATCC 23582 / Paris) OX=456481 GN=LEPBI_I0608 PE=4 SV=1 |
| tr\|B0SK46\|B0SK46_LEPBP | 7 | 7 | 5 | 19 | 3 | Uncharacterized protein OS=Leptospira biflexa serovar Patoc (strain Patoc 1 / ATCC 23582 / Paris) OX=456481 GN=LEPBI_I0025 PE=4 SV=1 |
| tr\|B0SM21\|B0SM21_LEPBP | 9 | 7 | 3 | 19 | 3 | HlyD_D23 domain-containing protein OS=Leptospira biflexa serovar Patoc (strain Patoc 1 / ATCC 23582 / Paris) OX=456481 GN=LEPBI_I2570 PE=4 SV=1 |
| tr\|B0SL97\|B0SL97_LEPBP | 7 | 6 | 5 | 18 | 3 | DNA repair protein RadA OS=Leptospira biflexa serovar Patoc (strain Patoc 1 / ATCC 23582 / Paris) OX=456481 GN=radA PE=3 SV=1 |
| sp\|B0SQV0\|RF1_LEPBP | 2 | 4 | 12 | 18 | 3 | Peptide chain release factor 1 OS=Leptospira biflexa serovar Patoc (strain Patoc 1 / ATCC 23582 / Paris) OX=456481 GN=prfA PE=3 SV=1 |
| tr\|B0SKJ0\|B0SKJ0_LEPBP | 8 | 3 | 7 | 18 | 3 | Uncharacterized protein OS=Leptospira biflexa serovar Patoc (strain Patoc 1 / ATCC 23582 / Paris) OX=456481 GN=LEPBI_I0612 PE=4 SV=1 |
| tr\|B0SPQ6\|B0SPQ6_LEPBP | 7 | 5 | 6 | 18 | 3 | Putative membrane bound 0-acyl transferase, putative alginate biosynthesis protein putative membrane protein OS=Leptospira biflexa serovar Patoc (strain Patoc 1 / ATCC 23582 / Paris) OX=456481 GN=LEPBI_I3096 PE=3 SV=1 |
| tr\|B0SP34\|B0SP34_LEPBP | 7 | 5 | 6 | 18 | 3 | Glu_synthase domain-containing protein OS=Leptospira biflexa serovar Patoc (strain Patoc 1 / ATCC 23582 / Paris) OX=456481 GN=LEPBI_I2978 PE=3 SV=1 |
| tr\|B0SMS7\|B0SMS7_LEPBP | 5 | 5 | 8 | 18 | 3 | PilZ domain-containing protein OS=Leptospira biflexa serovar Patoc (strain Patoc 1 / ATCC 23582 / Paris) OX=456481 GN=LEPBI_I2723 PE=4 SV=1 |
| tr\|B0SQ33\|B0SQ33_LEPBP | 1 | 8 | 9 | 18 | 3 | Uncharacterized protein OS=Leptospira biflexa serovar Patoc (strain Patoc 1 / ATCC 23582 / Paris) OX=456481 GN=LEPBI_I3119 PE=4 SV=1 |
| tr\|B0SJ80\|B0SJ80_LEPBP | 2 | 2 | 14 | 18 | 3 | Uncharacterized protein OS=Leptospira biflexa serovar Patoc (strain Patoc 1 / ATCC 23582 / Paris) OX=456481 GN=LEPBI_I0492 PE=4 SV=1 |
| sp\|B0SRH5\|NUOI_LEPBP | 8 | 9 | 1 | 18 | 3 | NADH-quinone oxidoreductase subunit I OS=Leptospira biflexa serovar Patoc (strain Patoc 1 / ATCC 23582 / Paris) OX=456481 GN=nuoI PE=3 SV=1 |
| sp\|B0SPT9\|RIMO_LEPBP | 7 | 6 | 5 | 18 | 3 | Ribosomal protein S12 methylthiotransferase RimO OS=Leptospira biflexa serovar Patoc (strain Patoc 1 / ATCC 23582 / Paris) OX=456481 GN=rimO PE=3 SV=1 |
| tr\|B0SS30\|B0SS30_LEPBP | 5 | 6 | 7 | 18 | 3 | Putative permease putative membrane protein OS=Leptospira biflexa serovar Patoc (strain Patoc 1 / ATCC 23582 / Paris) OX=456481 GN=LEPBI_I1814 PE=4 SV=1 |
| tr\|B0SKR1\|B0SKR1_LEPBP | 5 | 4 | 9 | 18 | 3 | Uncharacterized protein OS=Leptospira biflexa serovar Patoc (strain Patoc 1 / ATCC 23582 / Paris) OX=456481 GN=LEPBI_I0686 PE=4 SV=1 |
| tr\|B0SU45\|B0SU45_LEPBP | 7 | 6 | 4 | 17 | 3 | Putative divalent cation transport-related protein OS=Leptospira biflexa serovar Patoc (strain Patoc 1 / ATCC 23582 / Paris) OX=456481 GN=LEPBI_II0196 PE=3 SV=1 |
| tr\|B0SR83\|B0SR83_LEPBP | 6 | 4 | 7 | 17 | 3 | General secretion pathway protein F OS=Leptospira biflexa serovar Patoc (strain Patoc 1 / ATCC 23582 / Paris) OX=456481 GN=gspF PE=3 SV=1 |
| tr\|B0SRX3\|B0SRX3_LEPBP | 7 | 7 | 3 | 17 | 3 | Putative K(+)-stimulated pyrophosphate-energized sodium pump OS=Leptospira biflexa serovar Patoc (strain Patoc 1 / ATCC 23582 / Paris) OX=456481 GN=hppA PE=3 SV=1 |
| tr\|B0SS17\|B0SS17_LEPBP | 5 | 6 | 6 | 17 | 3 | Radical SAM core domain-containing protein OS=Leptospira biflexa serovar Patoc (strain Patoc 1 / ATCC 23582 / Paris) OX=456481 GN=LEPBI_I1801 PE=4 SV=1 |
| tr\|B0SLW4\|B0SLW4_LEPBP | 9 | 7 | 1 | 17 | 3 | Uncharacterized protein OS=Leptospira biflexa serovar Patoc (strain Patoc 1 / ATCC 23582 / Paris) OX=456481 GN=LEPBI_I0891 PE=4 SV=1 |
| tr\|B0SMA6\|B0SMA6_LEPBP | 9 | 6 | 2 | 17 | 3 | Putative phosphoserine phosphatase RsbU (Sigma factor SigB regulation protein RsbU) OS=Leptospira biflexa serovar Patoc (strain Patoc 1 / ATCC 23582 / Paris) OX=456481 GN=LEPBI_I0926 PE=4 SV=1 |
| tr\|B0SS09\|B0SS09_LEPBP | 4 | 8 | 5 | 17 | 3 | Putative acetylglutamate kinase OS=Leptospira biflexa serovar Patoc (strain Patoc 1 / ATCC 23582 / Paris) OX=456481 GN=LEPBI_I1793 PE=4 SV=1 |
| tr\|B0SNN1\|B0SNN1_LEPBP | 3 | 2 | 12 | 17 | 3 | UDP-N-acetylmuramoylalanine--D-glutamate ligase OS=Leptospira biflexa serovar Patoc (strain Patoc 1 / ATCC 23582 / Paris) OX=456481 GN=murD PE=3 SV=1 |
| tr\|B0SKP1\|B0SKP1_LEPBP | 5 | 2 | 10 | 17 | 3 | WYL domain-containing protein OS=Leptospira biflexa serovar Patoc (strain Patoc 1 / ATCC 23582 / Paris) OX=456481 GN=LEPBI_I0665 PE=4 SV=1 |
| tr\|B0SMT9\|B0SMT9_LEPBP | 4 | 8 | 5 | 17 | 3 | Formate-dependent phosphoribosylglycinamide formyltransferase OS=Leptospira biflexa serovar Patoc (strain Patoc 1 / ATCC 23582 / Paris) OX=456481 GN=purT PE=3 SV=1 |
| sp\|B0SRT7\|OBG_LEPBP | 4 | 5 | 8 | 17 | 3 | GTPase Obg OS=Leptospira biflexa serovar Patoc (strain Patoc 1 / ATCC 23582 / Paris) OX=456481 GN=obg PE=3 SV=1 |
| sp\|B0SSH5\|RL23_LEPBP | 7 | 8 | 2 | 17 | 3 | 50S ribosomal protein L23 OS=Leptospira biflexa serovar Patoc (strain Patoc 1 / ATCC 23582 / Paris) OX=456481 GN=rplW PE=3 SV=1 |
| tr\|B0STI2\|B0STI2_LEPBP | 4 | 9 | 4 | 17 | 3 | Phosphoribosylamine--glycine ligase OS=Leptospira biflexa serovar Patoc (strain Patoc 1 / ATCC 23582 / Paris) OX=456481 GN=purD PE=3 SV=1 |
| sp\|B0SQH2\|RIMP_LEPBP | 3 | 12 | 2 | 17 | 3 | Ribosome maturation factor RimP OS=Leptospira biflexa serovar Patoc (strain Patoc 1 / ATCC 23582 / Paris) OX=456481 GN=rimP PE=3 SV=1 |
| tr\|B0SN09\|B0SN09_LEPBP | 4 | 6 | 6 | 16 | 3 | Probable malate:quinone oxidoreductase OS=Leptospira biflexa serovar Patoc (strain Patoc 1 / ATCC 23582 / Paris) OX=456481 GN=mqo PE=3 SV=1 |
| tr\|B0SME0\|B0SME0_LEPBP | 6 | 4 | 6 | 16 | 3 | Uncharacterized protein OS=Leptospira biflexa serovar Patoc (strain Patoc 1 / ATCC 23582 / Paris) OX=456481 GN=LEPBI_I0960 PE=4 SV=1 |

| | | | | | |
|---|---|---|---|---|---|
| tr\|B0SSS0\|B0SSS0_LEPBP | -4.74 | 1.39E-03 | 1.16E-03 | 2.44E-04 | Putative outer membrane efflux protein putative signal peptide OS=Leptospira biflexa serovar Patoc (strain Patoc 1 / ATCC 23582 / Paris) OX=456481 GN=LEPBI_I2058 PE=3 SV=1 |
| tr\|B0SM44\|B0SM44_LEPBP | -4.67 | 1.53E-02 | 3.53E-03 | 7.55E-04 | 30S ribosomal protein S1 OS=Leptospira biflexa serovar Patoc (strain Patoc 1 / ATCC 23582 / Paris) OX=456481 GN=rpsA PE=3 SV=1 |
| sp\|B0SMR8\|THIM_LEPBP | -4.62 | 6.00E-03 | 7.18E-04 | 1.55E-04 | Hydroxyethylthiazole kinase OS=Leptospira biflexa serovar Patoc (strain Patoc 1 / ATCC 23582 / Paris) OX=456481 GN=thiM PE=3 SV=1 |
| tr\|B0SML9\|B0SML9_LEPBP | -4.54 | 1.82E-03 | 2.79E-03 | 6.15E-04 | ABC-type transport system, ATP-binding component OS=Leptospira biflexa serovar Patoc (strain Patoc 1 / ATCC 23582 / Paris) OX=456481 GN=ybiT PE=4 SV=1 |
| tr\|B0SK90\|B0SK90_LEPBP | -4.48 | 4.24E-04 | 1.79E-03 | 3.99E-04 | NAD(P) transhydrogenase subunit beta OS=Leptospira biflexa serovar Patoc (strain Patoc 1 / ATCC 23582 / Paris) OX=456481 GN=pntB PE=3 SV=1 |
| tr\|B0SL39\|B0SL39_LEPBP | -4.40 | 2.03E-02 | 6.83E-04 | 1.55E-04 | ABC-type transport system, ATP binding protein OS=Leptospira biflexa serovar Patoc (strain Patoc 1 / ATCC 23582 / Paris) OX=456481 GN=LEPBI_I0714 PE=4 SV=1 |
| tr\|B0SLS1\|B0SLS1_LEPBP | -4.35 | 1.59E-02 | 1.41E-03 | 3.25E-04 | Lon protease OS=Leptospira biflexa serovar Patoc (strain Patoc 1 / ATCC 23582 / Paris) OX=456481 GN=lon PE=2 SV=1 |
| tr\|B0SNV5\|B0SNV5_LEPBP | -4.33 | 3.14E-03 | 8.10E-04 | 1.87E-04 | PEGA domain-containing protein OS=Leptospira biflexa serovar Patoc (strain Patoc 1 / ATCC 23582 / Paris) OX=456481 GN=LEPBI_I2898 PE=4 SV=1 |
| tr\|B0SJR6\|B0SJR6_LEPBP | -4.29 | 1.78E-05 | 9.18E-04 | 2.14E-04 | Putative tRNA (Uracil-5-)-methyltransferase OS=Leptospira biflexa serovar Patoc (strain Patoc 1 / ATCC 23582 / Paris) OX=456481 GN=LEPBI_I0091 PE=3 SV=1 |
| tr\|B0SPS9\|B0SPS9_LEPBP | -4.27 | 1.05E-04 | 4.22E-03 | 9.89E-04 | Poly(A) polymerase I OS=Leptospira biflexa serovar Patoc (strain Patoc 1 / ATCC 23582 / Paris) OX=456481 GN=pcnB PE=3 SV=1 |
| tr\|B0SML0\|B0SML0_LEPBP | -4.18 | 2.56E-02 | 8.95E-04 | 2.14E-04 | Putative transcriptional regulator OS=Leptospira biflexa serovar Patoc (strain Patoc 1 / ATCC 23582 / Paris) OX=456481 GN=LEPBI_I1031 PE=4 SV=1 |
| tr\|B0STV0\|B0STV0_LEPBP | -4.15 | 2.36E-02 | 1.56E-03 | 3.77E-04 | Rubrerythrin domain-containing protein OS=Leptospira biflexa serovar Patoc (strain Patoc 1 / ATCC 23582 / Paris) OX=456481 GN=LEPBI_II0098 PE=4 SV=1 |
| tr\|B0SJP1\|B0SJP1_LEPBP | -4.13 | 5.88E-03 | 1.21E-03 | 2.93E-04 | Acyl-CoA dehydrogenase OS=Leptospira biflexa serovar Patoc (strain Patoc 1 / ATCC 23582 / Paris) OX=456481 GN=acdA2 PE=3 SV=1 |
| tr\|B0SR08\|B0SR08_LEPBP | -4.12 | 8.27E-03 | 1.33E-03 | 3.22E-04 | PHB domain-containing protein OS=Leptospira biflexa serovar Patoc (strain Patoc 1 / ATCC 23582 / Paris) OX=456481 GN=LEPBI_I1602 PE=3 SV=1 |
| tr\|B0SR81\|B0SR81_LEPBP | -4.12 | 3.00E-03 | 2.10E-03 | 5.10E-04 | General secretory pathway protein D OS=Leptospira biflexa serovar Patoc (strain Patoc 1 / ATCC 23582 / Paris) OX=456481 GN=gspD PE=3 SV=1 |
| tr\|B0SMV8\|B0SMV8_LEPBP | -4.06 | 2.62E-03 | 1.24E-03 | 3.05E-04 | Uncharacterized protein OS=Leptospira biflexa serovar Patoc (strain Patoc 1 / ATCC 23582 / Paris) OX=456481 GN=LEPBI_I2754 PE=4 SV=1 |
| sp\|B0SRS3\|RSMH_LEPBP | -3.97 | 2.82E-02 | 6.61E-04 | 1.66E-04 | Ribosomal RNA small subunit methyltransferase H OS=Leptospira biflexa serovar Patoc (strain Patoc 1 / ATCC 23582 / Paris) OX=456481 GN=rsmH PE=3 SV=1 |
| tr\|B0SSN2\|B0SSN2_LEPBP | -3.93 | 9.89E-03 | 1.15E-03 | 2.93E-04 | Uncharacterized protein OS=Leptospira biflexa serovar Patoc (strain Patoc 1 / ATCC 23582 / Paris) OX=456481 GN=LEPBI_I2020 PE=4 SV=1 |
| tr\|B0SM29\|B0SM29_LEPBP | -3.86 | 6.28E-04 | 9.20E-04 | 2.39E-04 | Uncharacterized protein OS=Leptospira biflexa serovar Patoc (strain Patoc 1 / ATCC 23582 / Paris) OX=456481 GN=LEPBI_I2578 PE=4 SV=1 |
| tr\|B0SQL8\|B0SQL8_LEPBP | -3.84 | 6.85E-03 | 1.58E-03 | 4.11E-04 | Aerobic glycerol-3-phosphate dehydrogenase OS=Leptospira biflexa serovar Patoc (strain Patoc 1 / ATCC 23582 / Paris) OX=456481 GN=glpD PE=3 SV=1 |
| tr\|B0SK74\|B0SK74_LEPBP | -3.76 | 6.87E-03 | 2.10E-03 | 5.59E-04 | Putative ribonuclease D (RNase D) OS=Leptospira biflexa serovar Patoc (strain Patoc 1 / ATCC 23582 / Paris) OX=456481 GN=LEPBI_I0172 PE=4 SV=1 |
| tr\|B0SJ44\|B0SJ44_LEPBP | -3.75 | 5.52E-03 | 2.22E-03 | 5.91E-04 | Putative peptidase, S41 family putative signal peptide OS=Leptospira biflexa serovar Patoc (strain Patoc 1 / ATCC 23582 / Paris) OX=456481 GN=LEPBI_I0474 PE=3 SV=1 |
| tr\|B0SR64\|B0SR64_LEPBP | -3.75 | 1.62E-02 | 9.22E-04 | 2.46E-04 | Uncharacterized protein OS=Leptospira biflexa serovar Patoc (strain Patoc 1 / ATCC 23582 / Paris) OX=456481 GN=LEPBI_I1658 PE=4 SV=1 |
| tr\|B0SQL6\|B0SQL6_LEPBP | -3.74 | 1.83E-02 | 2.96E-03 | 7.90E-04 | RNA polymerase sigma factor SigA OS=Leptospira biflexa serovar Patoc (strain Patoc 1 / ATCC 23582 / Paris) OX=456481 GN=rpoD PE=3 SV=1 |
| tr\|B0SU22\|B0SU22_LEPBP | -3.74 | 2.25E-03 | 1.18E-03 | 3.14E-04 | Sigma factor SigB regulation protein RsbU putative phosphoserine phosphatase OS=Leptospira biflexa serovar Patoc (strain Patoc 1 / ATCC 23582 / Paris) OX=456481 GN=LEPBI_II0171 PE=4 SV=1 |
| tr\|B0SPR0\|B0SPR0_LEPBP | -3.69 | 3.05E-03 | 1.01E-03 | 2.73E-04 | Putative dolichyl-phosphate-mannose--protein mannosyltransferase protein, family 39 putative membrane protein OS=Leptospira biflexa serovar Patoc (strain Patoc 1 / ATCC 23582 / Paris) OX=456481 GN=LEPBI_I1367 PE=4 SV=1 |
| tr\|B0SQL3\|B0SQL3_LEPBP | -3.57 | 1.41E-02 | 6.05E-04 | 1.69E-04 | 30S ribosomal protein S21 OS=Leptospira biflexa serovar Patoc (strain Patoc 1 / ATCC 23582 / Paris) OX=456481 GN=rpsU PE=3 SV=1 |
| tr\|B0SM59\|B0SM59_LEPBP | -3.57 | 2.33E-02 | 1.14E-03 | 3.18E-04 | Putative membrane-associated Zn-dependent metalloprotease, M50B family putative membrane protein OS=Leptospira biflexa serovar Patoc (strain Patoc 1 / ATCC 23582 / Paris) OX=456481 GN=LEPBI_I2610 PE=3 SV=1 |
| tr\|B0SRQ8\|B0SRQ8_LEPBP | -3.57 | 1.82E-02 | 8.45E-04 | 2.37E-04 | AAA domain-containing protein OS=Leptospira biflexa serovar Patoc (strain Patoc 1 / ATCC 23582 / Paris) OX=456481 GN=LEPBI_I1747 PE=4 SV=1 |
| tr\|B0SM43\|B0SM43_LEPBP | -3.55 | 6.34E-04 | 1.24E-03 | 3.50E-04 | Uncharacterized protein OS=Leptospira biflexa serovar Patoc (strain Patoc 1 / ATCC 23582 / Paris) OX=456481 GN=LEPBI_I2594 PE=4 SV=1 |
| tr\|B0STH0\|B0STH0_LEPBP | -3.46 | 1.27E-02 | 3.15E-03 | 9.10E-04 | Phosphomannomutase (Phosphomannose mutase) OS=Leptospira biflexa serovar Patoc (strain Patoc 1 / ATCC 23582 / Paris) OX=456481 GN=manB PE=3 SV=1 |
| sp\|B0SSH9\|EFTU_LEPBP | -3.45 | 1.24E-03 | 1.35E-02 | 3.90E-03 | Elongation factor Tu OS=Leptospira biflexa serovar Patoc (strain Patoc 1 / ATCC 23582 / Paris) OX=456481 GN=tuf PE=3 SV=1 |
| tr\|B0SRG2\|B0SRG2_LEPBP | -3.41 | 1.17E-02 | 1.99E-03 | 5.82E-04 | Putative ABC-type transport sytem, outer membrane protein OS=Leptospira biflexa serovar Patoc (strain Patoc 1 / ATCC 23582 / Paris) OX=456481 GN=LEPBI_I3390 PE=3 SV=1 |
| tr\|B0SQI9\|B0SQI9_LEPBP | -3.40 | 5.92E-03 | 1.46E-03 | 4.30E-04 | ATP-grasp domain-containing protein OS=Leptospira biflexa serovar Patoc (strain Patoc 1 / ATCC 23582 / Paris) OX=456481 GN=LEPBI_I1540 PE=4 SV=1 |
| tr\|B0STT2\|B0STT2_LEPBP | -3.37 | 2.97E-03 | 2.18E-03 | 6.47E-04 | Uncharacterized protein OS=Leptospira biflexa serovar Patoc (strain Patoc 1 / ATCC 23582 / Paris) OX=456481 GN=LEPBI_II0078 PE=4 SV=1 |
| tr\|B0SQT0\|B0SQT0_LEPBP | -3.36 | 9.04E-04 | 1.32E-03 | 3.94E-04 | Uncharacterized protein OS=Leptospira biflexa serovar Patoc (strain Patoc 1 / ATCC 23582 / Paris) OX=456481 GN=LEPBI_I3262 PE=4 SV=1 |
| tr\|B0SR15\|B0SR15_LEPBP | -3.36 | 1.71E-02 | 7.35E-04 | 2.19E-04 | Uncharacterized protein OS=Leptospira biflexa serovar Patoc (strain Patoc 1 / ATCC 23582 / Paris) OX=456481 GN=LEPBI_I1609 PE=4 SV=1 |
| tr\|B0SSU9\|B0SSU9_LEPBP | -3.35 | 1.52E-02 | 1.25E-03 | 3.74E-04 | Uncharacterized protein OS=Leptospira biflexa serovar Patoc (strain Patoc 1 / ATCC 23582 / Paris) OX=456481 GN=LEPBI_I2087 PE=4 SV=1 |
| tr\|B0SLN9\|B0SLN9_LEPBP | -3.28 | 3.18E-02 | 1.97E-03 | 6.01E-04 | DNA helicase OS=Leptospira biflexa serovar Patoc (strain Patoc 1 / ATCC 23582 / Paris) OX=456481 GN=LEPBI_I2544 PE=3 SV=1 |
| tr\|B0STF7\|B0STF7_LEPBP | -3.23 | 1.82E-03 | 7.05E-04 | 2.19E-04 | Acyl coenzyme A dehydrogenase OS=Leptospira biflexa serovar Patoc (strain Patoc 1 / ATCC 23582 / Paris) OX=456481 GN=LEPBI_I2302 PE=3 SV=1 |
| tr\|B0SJ90\|B0SJ90_LEPBP | -3.18 | 5.60E-04 | 1.13E-03 | 3.55E-04 | Putative S-adenosyl-L-methionine-dependent methyltransferase OS=Leptospira biflexa serovar Patoc (strain Patoc 1 / ATCC 23582 / Paris) OX=456481 GN=LEPBI_I0584 PE=4 SV=1 |
| tr\|B0SJ13\|B0SJ13_LEPBP | -3.13 | 2.50E-02 | 3.65E-03 | 1.17E-03 | Membrane protein insertase YidC OS=Leptospira biflexa serovar Patoc (strain Patoc 1 / ATCC 23582 / Paris) OX=456481 GN=yidC PE=3 SV=1 |
| tr\|B0STL1\|B0STL1_LEPBP | -3.11 | 9.54E-03 | 1.25E-03 | 4.01E-04 | Uncharacterized protein OS=Leptospira biflexa serovar Patoc (strain Patoc 1 / ATCC 23582 / Paris) OX=456481 GN=LEPBI_II0007 PE=4 SV=1 |
| tr\|B0SJ63\|B0SJ63_LEPBP | -3.09 | 8.62E-03 | 3.83E-03 | 1.24E-03 | Glucose-1-phosphate thymidylyltransferase OS=Leptospira biflexa serovar Patoc (strain Patoc 1 / ATCC 23582 / Paris) OX=456481 GN=rfbA PE=4 SV=1 |
| tr\|B0SRZ3\|B0SRZ3_LEPBP | -3.00 | 3.73E-04 | 8.87E-04 | 2.95E-04 | Peptidase_M23 domain-containing protein OS=Leptospira biflexa serovar Patoc (strain Patoc 1 / ATCC 23582 / Paris) OX=456481 GN=LEPBI_I3469 PE=4 SV=1 |
| sp\|B0SLC7\|ATPG_LEPBP | -2.99 | 7.96E-03 | 5.30E-03 | 1.77E-03 | ATP synthase gamma chain OS=Leptospira biflexa serovar Patoc (strain Patoc 1 / ATCC 23582 / Paris) OX=456481 GN=atpG PE=3 SV=1 |
| tr\|B0SSB2\|B0SSB2_LEPBP | -2.98 | 5.90E-04 | 2.98E-03 | 1.00E-03 | 2-oxoglutarate reductase OS=Leptospira biflexa serovar Patoc (strain Patoc 1 / ATCC 23582 / Paris) OX=456481 GN=serA2 PE=3 SV=1 |
| tr\|B0SMF6\|B0SMF6_LEPBP | -2.94 | 2.89E-02 | 7.14E-04 | 2.43E-04 | Hydrolase_4 domain-containing protein OS=Leptospira biflexa serovar Patoc (strain Patoc 1 / ATCC 23582 / Paris) OX=456481 GN=LEPBI_I0976 PE=4 SV=1 |
| sp\|B0SSW0\|RS18_LEPBP | -2.92 | 2.92E-03 | 1.34E-03 | 4.58E-04 | 30S ribosomal protein S18 OS=Leptospira biflexa serovar Patoc (strain Patoc 1 / ATCC 23582 / Paris) OX=456481 GN=rpsR PE=3 SV=1 |
| tr\|B0SMM9\|B0SMM9_LEPBP | -2.92 | 2.01E-04 | 2.51E-03 | 8.59E-04 | TRAM domain-containing protein OS=Leptospira biflexa serovar Patoc (strain Patoc 1 / ATCC 23582 / Paris) OX=456481 GN=LEPBI_I2675 PE=3 SV=1 |
| tr\|B0STP0\|B0STP0_LEPBP | -2.90 | 1.17E-02 | 8.70E-04 | 3.00E-04 | Putative DNA-binding protein (HU-like protein) OS=Leptospira biflexa serovar Patoc (strain Patoc 1 / ATCC 23582 / Paris) OX=456481 GN=LEPBI_II0036 PE=3 SV=1 |
| tr\|B0SUA7\|B0SUA7_LEPBP | -2.88 | 1.30E-02 | 1.01E-03 | 3.49E-04 | Uncharacterized protein OS=Leptospira biflexa serovar Patoc (strain Patoc 1 / ATCC 23582 / Paris) OX=456481 GN=LEPBI_II0258 PE=4 SV=1 |
| tr\|B0STN0\|B0STN0_LEPBP | -2.87 | 3.04E-02 | 1.31E-03 | 4.55E-04 | Putative chromosome partitioning protein ParB OS=Leptospira biflexa serovar Patoc (strain Patoc 1 / ATCC 23582 / Paris) OX=456481 GN=LEPBI_II0026 PE=3 SV=1 |
| sp\|B0SLE9\|RECA_LEPBP | -2.87 | 1.55E-02 | 4.43E-03 | 1.54E-03 | Protein RecA OS=Leptospira biflexa serovar Patoc (strain Patoc 1 / ATCC 23582 / Paris) OX=456481 GN=recA PE=3 SV=1 |
| tr\|B0SRR8\|B0SRR8_LEPBP | -2.84 | 5.29E-03 | 9.86E-04 | 3.47E-04 | UDP-N-acetylglucosamine--N-acetylmuramyl-(pentapeptide) pyrophosphoryl-undecaprenol N-acetylglucosamine transferase OS=Leptospira biflexa serovar Patoc (strain Patoc 1 / ATCC 23582 / Paris) OX=456481 GN=murG PE=3 SV=1 |
| tr\|B0SKG0\|B0SKG0_LEPBP | -2.81 | 6.67E-03 | 2.24E-03 | 7.98E-04 | ATP-dependent zinc metalloprotease FtsH OS=Leptospira biflexa serovar Patoc (strain Patoc 1 / ATCC 23582 / Paris) OX=456481 GN=ftsH PE=3 SV=1 |
| sp\|B0SNL6\|CYSI_LEPBP | -2.79 | 1.20E-02 | 8.44E-04 | 3.03E-04 | Sulfite reductase [NADPH] hemoprotein beta-component OS=Leptospira biflexa serovar Patoc (strain Patoc 1 / ATCC 23582 / Paris) OX=456481 GN=cysI PE=3 SV=1 |
| tr\|B0SKW0\|B0SKW0_LEPBP | -2.78 | 2.60E-02 | 1.57E-03 | 5.66E-04 | Acetolactate synthase OS=Leptospira biflexa serovar Patoc (strain Patoc 1 / ATCC 23582 / Paris) OX=456481 GN=ilvB2 PE=3 SV=1 |
| tr\|B0STR8\|B0STR8_LEPBP | -2.70 | 2.18E-02 | 9.56E-04 | 3.55E-04 | Putative cytochrome c oxidase, subunit I (Heme B/copper cytochrome c oxidase subunit) putative membrane protein OS=Leptospira biflexa serovar Patoc (strain Patoc 1 / ATCC 23582 / Paris) OX=456481 GN=LEPBI_II0064 PE=3 SV=1 |

| | | | | | | |
|---|---|---|---|---|---|---|
| tr\|B0SIW6\|B0SIW6_LEPBP | 6 | 7 | 3 | 16 | 3 | Uncharacterized protein OS=Leptospira biflexa serovar Patoc (strain Patoc 1 / ATCC 23582 / Paris) OX=456481 GN=LEPBI_I0417 PE=4 SV=1 |
| tr\|B0SJQ1\|B0SJQ1_LEPBP | 5 | 2 | 9 | 16 | 3 | FGE-sulfatase domain-containing protein OS=Leptospira biflexa serovar Patoc (strain Patoc 1 / ATCC 23582 / Paris) OX=456481 GN=LEPBI_I0067 PE=4 SV=1 |
| tr\|B0SNG9\|B0SNG9_LEPBP | 3 | 12 | 1 | 16 | 3 | Uncharacterized protein OS=Leptospira biflexa serovar Patoc (strain Patoc 1 / ATCC 23582 / Paris) OX=456481 GN=LEPBI_I2867 PE=4 SV=1 |
| tr\|B0SU96\|B0SU96_LEPBP | 5 | 6 | 5 | 16 | 3 | Methyltransf_25 domain-containing protein OS=Leptospira biflexa serovar Patoc (strain Patoc 1 / ATCC 23582 / Paris) OX=456481 GN=LEPBI_II0247 PE=4 SV=1 |
| tr\|B0SMR0\|B0SMR0_LEPBP | 4 | 4 | 8 | 16 | 3 | PlsC domain-containing protein OS=Leptospira biflexa serovar Patoc (strain Patoc 1 / ATCC 23582 / Paris) OX=456481 GN=LEPBI_I2706 PE=4 SV=1 |
| tr\|B0SKZ3\|B0SKZ3_LEPBP | 9 | 3 | 4 | 16 | 3 | Uncharacterized protein OS=Leptospira biflexa serovar Patoc (strain Patoc 1 / ATCC 23582 / Paris) OX=456481 GN=LEPBI_I2396 PE=4 SV=1 |
| tr\|B0SQK6\|B0SQK6_LEPBP | 11 | 4 | 1 | 16 | 3 | Acetyltransferase component of pyruvate dehydrogenase complex OS=Leptospira biflexa serovar Patoc (strain Patoc 1 / ATCC 23582 / Paris) OX=456481 GN=acoC PE=3 SV=1 |
| tr\|B0SLY5\|B0SLY5_LEPBP | 8 | 7 | 1 | 16 | 3 | Glutamate racemase OS=Leptospira biflexa serovar Patoc (strain Patoc 1 / ATCC 23582 / Paris) OX=456481 GN=murI PE=3 SV=1 |
| tr\|B0SUB4\|B0SUB4_LEPBP | 8 | 6 | 2 | 16 | 3 | Hypothetical methyl-accepting chemotaxis protein putative membrane protein OS=Leptospira biflexa serovar Patoc (strain Patoc 1 / ATCC 23582 / Paris) OX=456481 GN=LEPBI_II0265 PE=4 SV=1 |
| tr\|B0SJ91\|B0SJ91_LEPBP | 7 | 8 | 1 | 16 | 3 | Diaminopimelate epimerase OS=Leptospira biflexa serovar Patoc (strain Patoc 1 / ATCC 23582 / Paris) OX=456481 GN=dapF PE=3 SV=1 |
| tr\|B0SPN8\|B0SPN8_LEPBP | 2 | 6 | 8 | 16 | 3 | Riboflavin biosynthesis protein OS=Leptospira biflexa serovar Patoc (strain Patoc 1 / ATCC 23582 / Paris) OX=456481 GN=ribF PE=3 SV=1 |
| tr\|B0SQG5\|B0SQG5_LEPBP | 3 | 6 | 7 | 16 | 3 | PPK2 domain-containing protein OS=Leptospira biflexa serovar Patoc (strain Patoc 1 / ATCC 23582 / Paris) OX=456481 GN=LEPBI_I1516 PE=4 SV=1 |
| tr\|B0SPE5\|B0SPE5_LEPBP | 4 | 8 | 3 | 15 | 3 | Putative ABC-type transport system, permease, iron-regulated OS=Leptospira biflexa serovar Patoc (strain Patoc 1 / ATCC 23582 / Paris) OX=456481 GN=LEPBI_I1359 PE=3 SV=1 |
| tr\|B0SSD9\|B0SSD9_LEPBP | 5 | 5 | 5 | 15 | 3 | Putative two-component regulator OS=Leptospira biflexa serovar Patoc (strain Patoc 1 / ATCC 23582 / Paris) OX=456481 GN=LEPBI_I1926 PE=4 SV=1 |
| tr\|B0SUC7\|B0SUC7_LEPBP | 4 | 9 | 2 | 15 | 3 | Uncharacterized protein OS=Leptospira biflexa serovar Patoc (strain Patoc 1 / ATCC 23582 / Paris) OX=456481 GN=LEPBI_II0278 PE=4 SV=1 |
| tr\|B0SJQ4\|B0SJQ4_LEPBP | 10 | 4 | 1 | 15 | 3 | DUF4139 domain-containing protein OS=Leptospira biflexa serovar Patoc (strain Patoc 1 / ATCC 23582 / Paris) OX=456481 GN=LEPBI_I0070 PE=4 SV=1 |
| tr\|B0SLN3\|B0SLN3_LEPBP | 6 | 5 | 4 | 15 | 3 | Putative hydrolase, alpha/beta superfamily putative signal peptide OS=Leptospira biflexa serovar Patoc (strain Patoc 1 / ATCC 23582 / Paris) OX=456481 GN=LEPBI_I2538 PE=4 SV=1 |
| tr\|B0SQA8\|B0SQA8_LEPBP | 8 | 5 | 2 | 15 | 3 | Uncharacterized protein OS=Leptospira biflexa serovar Patoc (strain Patoc 1 / ATCC 23582 / Paris) OX=456481 GN=LEPBI_I3195 PE=4 SV=1 |
| tr\|B0SMI3\|B0SMI3_LEPBP | 5 | 6 | 4 | 15 | 3 | Putative lipopolysaccharide core biosynthesis glycosyl transferase RfaQ OS=Leptospira biflexa serovar Patoc (strain Patoc 1 / ATCC 23582 / Paris) OX=456481 GN=LEPBI_I1003 PE=4 SV=1 |
| tr\|B0SPR1\|B0SPR1_LEPBP | 2 | 5 | 7 | 14 | 3 | Putative cytochrome c3 putative signal peptide OS=Leptospira biflexa serovar Patoc (strain Patoc 1 / ATCC 23582 / Paris) OX=456481 GN=LEPBI_I1368 PE=4 SV=1 |
| tr\|B0SP52\|B0SP52_LEPBP | 5 | 6 | 3 | 14 | 3 | ABC-type transport system, ATPase OS=Leptospira biflexa serovar Patoc (strain Patoc 1 / ATCC 23582 / Paris) OX=456481 GN=LEPBI_I1265 PE=4 SV=1 |
| tr\|B0SKI6\|B0SKI6_LEPBP | 5 | 3 | 6 | 14 | 3 | Uncharacterized protein OS=Leptospira biflexa serovar Patoc (strain Patoc 1 / ATCC 23582 / Paris) OX=456481 GN=LEPBI_I0607 PE=4 SV=1 |
| sp\|B0SJC4\|TRUA_LEPBP | 6 | 6 | 2 | 14 | 3 | tRNA pseudouridine synthase A OS=Leptospira biflexa serovar Patoc (strain Patoc 1 / ATCC 23582 / Paris) OX=456481 GN=truA PE=3 SV=1 |
| tr\|B0SJX2\|B0SJX2_LEPBP | 4 | 3 | 7 | 14 | 3 | Uncharacterized protein OS=Leptospira biflexa serovar Patoc (strain Patoc 1 / ATCC 23582 / Paris) OX=456481 GN=LEPBI_I0134 PE=4 SV=1 |
| tr\|B0SNH0\|B0SNH0_LEPBP | 5 | 4 | 5 | 14 | 3 | Uncharacterized protein OS=Leptospira biflexa serovar Patoc (strain Patoc 1 / ATCC 23582 / Paris) OX=456481 GN=LEPBI_I2868 PE=4 SV=1 |
| tr\|B0SS42\|B0SS42_LEPBP | 3 | 2 | 9 | 14 | 3 | Uncharacterized protein OS=Leptospira biflexa serovar Patoc (strain Patoc 1 / ATCC 23582 / Paris) OX=456481 GN=LEPBI_I1826 PE=4 SV=1 |
| tr\|B0SS13\|B0SS13_LEPBP | 6 | 4 | 4 | 14 | 3 | L-lysine 2,3-aminomutase OS=Leptospira biflexa serovar Patoc (strain Patoc 1 / ATCC 23582 / Paris) OX=456481 GN=kamA PE=4 SV=1 |
| sp\|B0SQH7\|RS15_LEPBP | 4 | 6 | 4 | 14 | 3 | 30S ribosomal protein S15 OS=Leptospira biflexa serovar Patoc (strain Patoc 1 / ATCC 23582 / Paris) OX=456481 GN=rpsO PE=3 SV=1 |
| tr\|B0SUC9\|B0SUC9_LEPBP | 4 | 3 | 6 | 13 | 3 | Uncharacterized protein OS=Leptospira biflexa serovar Patoc (strain Patoc 1 / ATCC 23582 / Paris) OX=456481 GN=LEPBI_II0280 PE=4 SV=1 |
| tr\|B0SRC4\|B0SRC4_LEPBP | 6 | 6 | 1 | 13 | 3 | Dihydroxy-acid dehydratase OS=Leptospira biflexa serovar Patoc (strain Patoc 1 / ATCC 23582 / Paris) OX=456481 GN=ilvD PE=3 SV=1 |
| tr\|B0SRS5\|B0SRS5_LEPBP | 5 | 5 | 3 | 13 | 3 | Protein-glutamate O-methyltransferase OS=Leptospira biflexa serovar Patoc (strain Patoc 1 / ATCC 23582 / Paris) OX=456481 GN=cheR PE=4 SV=1 |
| tr\|B0SQI7\|B0SQI7_LEPBP | 4 | 4 | 5 | 13 | 3 | Putative S-adenosylmethionine-dependent methyltransferase OS=Leptospira biflexa serovar Patoc (strain Patoc 1 / ATCC 23582 / Paris) OX=456481 GN=LEPBI_I1538 PE=4 SV=1 |
| tr\|B0SKW3\|B0SKW3_LEPBP | 6 | 6 | 1 | 13 | 3 | Uncharacterized protein OS=Leptospira biflexa serovar Patoc (strain Patoc 1 / ATCC 23582 / Paris) OX=456481 GN=LEPBI_I2365 PE=4 SV=1 |
| tr\|B0SRY3\|B0SRY3_LEPBP | 6 | 3 | 4 | 13 | 3 | Putative RNA methyltransferase, TrmA family OS=Leptospira biflexa serovar Patoc (strain Patoc 1 / ATCC 23582 / Paris) OX=456481 GN=LEPBI_I3458 PE=3 SV=1 |
| tr\|B0SMY6\|B0SMY6_LEPBP | 8 | 4 | 1 | 13 | 3 | Putative pyridoxal-phosphate-dependent aminotransferase OS=Leptospira biflexa serovar Patoc (strain Patoc 1 / ATCC 23582 / Paris) OX=456481 GN=LEPBI_I1050 PE=3 SV=1 |
| tr\|B0SU97\|B0SU97_LEPBP | 3 | 3 | 7 | 13 | 3 | Uncharacterized protein OS=Leptospira biflexa serovar Patoc (strain Patoc 1 / ATCC 23582 / Paris) OX=456481 GN=LEPBI_II0248 PE=4 SV=1 |
| tr\|B0SNY0\|B0SNY0_LEPBP | 5 | 4 | 4 | 13 | 3 | Uncharacterized protein OS=Leptospira biflexa serovar Patoc (strain Patoc 1 / ATCC 23582 / Paris) OX=456481 GN=LEPBI_I2923 PE=4 SV=1 |
| tr\|B0SLY2\|B0SLY2_LEPBP | 5 | 5 | 3 | 13 | 3 | Uncharacterized protein OS=Leptospira biflexa serovar Patoc (strain Patoc 1 / ATCC 23582 / Paris) OX=456481 GN=LEPBI_I0910 PE=4 SV=1 |
| tr\|B0SQE3\|B0SQE3_LEPBP | 7 | 5 | 1 | 13 | 3 | Uncharacterized protein OS=Leptospira biflexa serovar Patoc (strain Patoc 1 / ATCC 23582 / Paris) OX=456481 GN=LEPBI_I1494 PE=4 SV=1 |
| tr\|B0SJR0\|B0SJR0_LEPBP | 7 | 5 | 1 | 13 | 3 | Glutathione-regulated potassium-efflux system protein KefB putative membrane protein OS=Leptospira biflexa serovar Patoc (strain Patoc 1 / ATCC 23582 / Paris) OX=456481 GN=kefB PE=4 SV=1 |
| tr\|B0SRH6\|B0SRH6_LEPBP | 4 | 4 | 5 | 13 | 3 | NADH-quinone oxidoreductase chain G OS=Leptospira biflexa serovar Patoc (strain Patoc 1 / ATCC 23582 / Paris) OX=456481 GN=nuoG PE=4 SV=1 |
| tr\|B0SPL1\|B0SPL1_LEPBP | 2 | 6 | 4 | 12 | 3 | Putative ankyrin-like protein OS=Leptospira biflexa serovar Patoc (strain Patoc 1 / ATCC 23582 / Paris) OX=456481 GN=LEPBI_I3051 PE=4 SV=1 |
| tr\|B0SQP7\|B0SQP7_LEPBP | 6 | 4 | 2 | 12 | 3 | Uncharacterized protein OS=Leptospira biflexa serovar Patoc (strain Patoc 1 / ATCC 23582 / Paris) OX=456481 GN=LEPBI_I3229 PE=4 SV=1 |
| tr\|B0SMY1\|B0SMY1_LEPBP | 2 | 1 | 9 | 12 | 3 | Uncharacterized protein OS=Leptospira biflexa serovar Patoc (strain Patoc 1 / ATCC 23582 / Paris) OX=456481 GN=LEPBI_I1045 PE=4 SV=1 |
| tr\|B0SMF2\|B0SMF2_LEPBP | 5 | 1 | 6 | 12 | 3 | Uncharacterized protein OS=Leptospira biflexa serovar Patoc (strain Patoc 1 / ATCC 23582 / Paris) OX=456481 GN=LEPBI_I0972 PE=4 SV=1 |
| tr\|B0SKR8\|B0SKR8_LEPBP | 3 | 7 | 2 | 12 | 3 | Lysine decarboxylase OS=Leptospira biflexa serovar Patoc (strain Patoc 1 / ATCC 23582 / Paris) OX=456481 GN=LEPBI_I0693 PE=3 SV=1 |
| tr\|B0SRF4\|B0SRF4_LEPBP | 7 | 2 | 3 | 12 | 3 | Uncharacterized protein OS=Leptospira biflexa serovar Patoc (strain Patoc 1 / ATCC 23582 / Paris) OX=456481 GN=LEPBI_I3382 PE=4 SV=1 |
| tr\|B0SJI3\|B0SJI3_LEPBP | 5 | 5 | 2 | 12 | 3 | Uncharacterized protein OS=Leptospira biflexa serovar Patoc (strain Patoc 1 / ATCC 23582 / Paris) OX=456481 GN=LEPBI_I0393 PE=4 SV=1 |
| tr\|B0STK4\|B0STK4_LEPBP | 1 | 7 | 4 | 12 | 3 | ABC-type transport system, ATP binding protein Putative heme-transporting ATPase OS=Leptospira biflexa serovar Patoc (strain Patoc 1 / ATCC 23582 / Paris) OX=456481 GN=LEPBI_I0274 PE=4 SV=1 |
| sp\|B0SS82\|LPXC_LEPBP | 3 | 3 | 6 | 12 | 3 | UDP-3-O-acyl-N-acetylglucosamine deacetylase OS=Leptospira biflexa serovar Patoc (strain Patoc 1 / ATCC 23582 / Paris) OX=456481 GN=lpxC PE=3 SV=1 |
| tr\|B0SRJ7\|B0SRJ7_LEPBP | 4 | 2 | 6 | 12 | 3 | Uncharacterized protein OS=Leptospira biflexa serovar Patoc (strain Patoc 1 / ATCC 23582 / Paris) OX=456481 GN=LEPBI_I1685 PE=4 SV=1 |
| sp\|B0SLQ3\|BIOB_LEPBP | 5 | 5 | 2 | 12 | 3 | Biotin synthase OS=Leptospira biflexa serovar Patoc (strain Patoc 1 / ATCC 23582 / Paris) OX=456481 GN=bioB PE=3 SV=1 |
| tr\|B0SSY3\|B0SSY3_LEPBP | 1 | 6 | 5 | 12 | 3 | Peptidyl-prolyl cis-trans isomerase OS=Leptospira biflexa serovar Patoc (strain Patoc 1 / ATCC 23582 / Paris) OX=456481 GN=ppiA PE=3 SV=1 |
| tr\|B0STW5\|B0STW5_LEPBP | 1 | 3 | 8 | 12 | 3 | Cyclopropane-fatty-acyl-phospholipid synthase OS=Leptospira biflexa serovar Patoc (strain Patoc 1 / ATCC 23582 / Paris) OX=456481 GN=cfa PE=3 SV=1 |
| tr\|B0SRR7\|B0SRR7_LEPBP | 2 | 2 | 8 | 12 | 3 | UDP-N-acetylmuramate--L-alanine ligase OS=Leptospira biflexa serovar Patoc (strain Patoc 1 / ATCC 23582 / Paris) OX=456481 GN=murC PE=4 SV=1 |
| tr\|B0SLQ6\|B0SLQ6_LEPBP | 5 | 5 | 1 | 11 | 3 | Putative hydrolase putative signal peptide OS=Leptospira biflexa serovar Patoc (strain Patoc 1 / ATCC 23582 / Paris) OX=456481 GN=LEPBI_I0831 PE=4 SV=1 |
| tr\|B0SU59\|B0SU59_LEPBP | 4 | 6 | 1 | 11 | 3 | S5A_REDUCTASE domain-containing protein OS=Leptospira biflexa serovar Patoc (strain Patoc 1 / ATCC 23582 / Paris) OX=456481 GN=LEPBI_II0210 PE=4 SV=1 |
| tr\|B0SJE1\|B0SJE1_LEPBP | 4 | 4 | 3 | 11 | 3 | Cell division protein FtsZ OS=Leptospira biflexa serovar Patoc (strain Patoc 1 / ATCC 23582 / Paris) OX=456481 GN=ftsZ PE=3 SV=1 |
| tr\|B0SPS3\|B0SPS3_LEPBP | 5 | 4 | 2 | 11 | 3 | TPR-repeat-containing protein OS=Leptospira biflexa serovar Patoc (strain Patoc 1 / ATCC 23582 / Paris) OX=456481 GN=LEPBI_I1380 PE=4 SV=1 |
| tr\|B0SSW9\|B0SSW9_LEPBP | 3 | 4 | 4 | 11 | 3 | CHAT domain-containing protein OS=Leptospira biflexa serovar Patoc (strain Patoc 1 / ATCC 23582 / Paris) OX=456481 GN=LEPBI_I2107 PE=4 SV=1 |

| | | | | | |
|---|---|---|---|---|---|
| tr\|B0SJP9\|B0SJP9_LEPBP | -2.69 | 1.60E-02 | 7.95E-04 | 2.95E-04 | Uncharacterized protein OS=Leptospira biflexa serovar Patoc (strain Patoc 1 / ATCC 23582 / Paris) OX=456481 GN=LEPBI_I0065 PE=4 SV=1 |
| tr\|B0SPN3\|B0SPN3_LEPBP | -2.66 | 1.25E-02 | 1.13E-03 | 4.25E-04 | Cell shape protein MreC OS=Leptospira biflexa serovar Patoc (strain Patoc 1 / ATCC 23582 / Paris) OX=456481 GN=mreC PE=3 SV=1 |
| sp\|B0SSH6\|RL4_LEPBP | -2.66 | 9.67E-03 | 2.69E-03 | 1.01E-03 | 50S ribosomal protein L4 OS=Leptospira biflexa serovar Patoc (strain Patoc 1 / ATCC 23582 / Paris) OX=456481 GN=rplD PE=3 SV=1 |
| tr\|B0SKQ0\|B0SKQ0_LEPBP | -2.64 | 2.07E-02 | 1.18E-03 | 4.49E-04 | Glycosyl transferase OS=Leptospira biflexa serovar Patoc (strain Patoc 1 / ATCC 23582 / Paris) OX=456481 GN=LEPBI_I0675 PE=4 SV=1 |
| tr\|B0SR32\|B0SR32_LEPBP | -2.61 | 2.30E-02 | 9.90E-04 | 3.79E-04 | 3-isopropylmalate dehydratase small subunit OS=Leptospira biflexa serovar Patoc (strain Patoc 1 / ATCC 23582 / Paris) OX=456481 GN=leuD PE=3 SV=1 |
| tr\|B0SLT5\|B0SLT5_LEPBP | -2.59 | 1.14E-02 | 5.55E-04 | 2.14E-04 | Uncharacterized protein OS=Leptospira biflexa serovar Patoc (strain Patoc 1 / ATCC 23582 / Paris) OX=456481 GN=LEPBI_I0860 PE=4 SV=1 |
| tr\|B0SN63\|B0SN63_LEPBP | -2.57 | 1.24E-02 | 8.63E-03 | 3.36E-03 | Electron transfer flavoprotein alpha-subunit (Alpha-ETF Electron transfer flavoprotein large subunit) (ETFLS) OS=Leptospira biflexa serovar Patoc (strain Patoc 1 / ATCC 23582 / Paris) OX=456481 GN=etfA PE=3 SV=1 |
| tr\|B0SQ11\|B0SQ11_LEPBP | -2.54 | 2.74E-02 | 6.82E-04 | 2.68E-04 | Uncharacterized protein OS=Leptospira biflexa serovar Patoc (strain Patoc 1 / ATCC 23582 / Paris) OX=456481 GN=LEPBI_I3097 PE=4 SV=1 |
| tr\|B0SM86\|B0SM86_LEPBP | -2.53 | 1.52E-02 | 1.99E-03 | 7.85E-04 | Putative strictosidine synthase putative signal peptide OS=Leptospira biflexa serovar Patoc (strain Patoc 1 / ATCC 23582 / Paris) OX=456481 GN=LEPBI_I2637 PE=4 SV=1 |
| tr\|B0SJK2\|B0SJK2_LEPBP | -2.48 | 2.84E-02 | 7.61E-04 | 3.07E-04 | Oligopetide ABC-type transport system, ATPase OS=Leptospira biflexa serovar Patoc (strain Patoc 1 / ATCC 23582 / Paris) OX=456481 GN=dppF PE=4 SV=1 |
| tr\|B0SPU4\|B0SPU4_LEPBP | -2.44 | 1.41E-02 | 3.26E-03 | 1.34E-03 | Protein translocase subunit SecD OS=Leptospira biflexa serovar Patoc (strain Patoc 1 / ATCC 23582 / Paris) OX=456481 GN=secD PE=3 SV=1 |
| tr\|B0SR55\|B0SR55_LEPBP | -2.44 | 2.34E-03 | 1.53E-03 | 6.27E-04 | ABC-type transport system, ATPase OS=Leptospira biflexa serovar Patoc (strain Patoc 1 / ATCC 23582 / Paris) OX=456481 GN=LEPBI_I1649 PE=4 SV=1 |
| tr\|B0SKT2\|B0SKT2_LEPBP | -2.44 | 2.37E-03 | 1.11E-03 | 4.55E-04 | Putative metalloendopeptidase putative signal peptide OS=Leptospira biflexa serovar Patoc (strain Patoc 1 / ATCC 23582 / Paris) OX=456481 GN=LEPBI_I2332 PE=4 SV=1 |
| tr\|B0SU57\|B0SU57_LEPBP | -2.35 | 2.59E-02 | 2.07E-03 | 8.81E-04 | ATP-dependent RNA helicase, DEAD-box family (DeaD) OS=Leptospira biflexa serovar Patoc (strain Patoc 1 / ATCC 23582 / Paris) OX=456481 GN=deaD PE=3 SV=1 |
| sp\|B0SSI6\|RL10_LEPBP | -2.34 | 2.60E-04 | 9.98E-04 | 4.26E-04 | 50S ribosomal protein L10 OS=Leptospira biflexa serovar Patoc (strain Patoc 1 / ATCC 23582 / Paris) OX=456481 GN=rplJ PE=3 SV=1 |
| tr\|B0SLV0\|B0SLV0_LEPBP | -2.34 | 7.85E-03 | 7.71E-04 | 3.30E-04 | Putative DSBA oxidoreductase putative membrane protein putative signal peptide OS=Leptospira biflexa serovar Patoc (strain Patoc 1 / ATCC 23582 / Paris) OX=456481 GN=LEPBI_I0877 PE=3 SV=1 |
| tr\|B0SP24\|B0SP24_LEPBP | -2.32 | 1.58E-05 | 2.00E-03 | 8.61E-04 | S-adenosylmethionine synthase OS=Leptospira biflexa serovar Patoc (strain Patoc 1 / ATCC 23582 / Paris) OX=456481 GN=metK PE=3 SV=1 |
| tr\|B0SJI0\|B0SJI0_LEPBP | -2.28 | 2.75E-02 | 6.04E-04 | 2.65E-04 | Putative long-chain fatty acid transport protein putative signal peptide OS=Leptospira biflexa serovar Patoc (strain Patoc 1 / ATCC 23582 / Paris) OX=456481 GN=LEPBI_I0390 PE=4 SV=1 |
| sp\|B0SMC0\|LPXK_LEPBP | -2.20 | 2.17E-02 | 6.02E-04 | 2.74E-04 | Tetraacyldisaccharide 4'-kinase OS=Leptospira biflexa serovar Patoc (strain Patoc 1 / ATCC 23582 / Paris) OX=456481 GN=lpxK PE=3 SV=1 |
| sp\|B0SSF5\|RS13_LEPBP | -2.20 | 2.81E-02 | 1.83E-03 | 8.34E-04 | 30S ribosomal protein S13 OS=Leptospira biflexa serovar Patoc (strain Patoc 1 / ATCC 23582 / Paris) OX=456481 GN=rpsM PE=3 SV=1 |
| tr\|B0SMN8\|B0SMN8_LEPBP | -2.08 | 6.91E-03 | 1.53E-03 | 7.36E-04 | Cytochrome c oxidase subunit 2 OS=Leptospira biflexa serovar Patoc (strain Patoc 1 / ATCC 23582 / Paris) OX=456481 GN=ctaC PE=3 SV=1 |
| tr\|B0SJ51\|B0SJ51_LEPBP | -2.00 | 1.17E-02 | 2.37E-03 | 1.18E-03 | UDP-n-acetylmuramate:l-alanyl-gamma-d-glutamyl-meso-diaminopimelate ligase OS=Leptospira biflexa serovar Patoc (strain Patoc 1 / ATCC 23582 / Paris) OX=456481 GN=mpl PE=4 SV=1 |

| | | | | | | |
|---|---|---|---|---|---|---|
| tr\|B0SLE8\|B0SLE8_LEPBP | 4 | 4 | 3 | 11 | 3 | Tyrosine recombinase XerC OS=Leptospira biflexa serovar Patoc (strain Patoc 1 / ATCC 23582 / Paris) OX=456481 GN=xerC PE=3 SV=1 |
| tr\|B0SQM5\|B0SQM5_LEPBP | 5 | 3 | 3 | 11 | 3 | Metallophos domain-containing protein OS=Leptospira biflexa serovar Patoc (strain Patoc 1 / ATCC 23582 / Paris) OX=456481 GN=LEPBI_I3207 PE=4 SV=1 |
| tr\|B0SLR1\|B0SLR1_LEPBP | 5 | 2 | 4 | 11 | 3 | FolC bifunctional protein: Folylpolyglutamate synthase/Tetrahydrofolate ynthase OS=Leptospira biflexa serovar Patoc (strain Patoc 1 / ATCC 23582 / Paris) OX=456481 GN=folC PE=4 SV=1 |
| tr\|B0SQT5\|B0SQT5_LEPBP | 5 | 3 | 2 | 10 | 3 | Uncharacterized protein OS=Leptospira biflexa serovar Patoc (strain Patoc 1 / ATCC 23582 / Paris) OX=456481 GN=LEPBI_I3267 PE=4 SV=1 |
| tr\|B0SK42\|B0SK42_LEPBP | 3 | 2 | 5 | 10 | 3 | DNA_pol3_delta domain-containing protein OS=Leptospira biflexa serovar Patoc (strain Patoc 1 / ATCC 23582 / Paris) OX=456481 GN=LEPBI_I0012 PE=4 SV=1 |
| tr\|B0SQZ2\|B0SQZ2_LEPBP | 4 | 4 | 2 | 10 | 3 | Putative transcriptional regulatory protein ZraR OS=Leptospira biflexa serovar Patoc (strain Patoc 1 / ATCC 23582 / Paris) OX=456481 GN=LEPBI_I1586 PE=4 SV=1 |
| tr\|B0SL96\|B0SL96_LEPBP | 5 | 4 | 1 | 10 | 3 | Ribonuclease D OS=Leptospira biflexa serovar Patoc (strain Patoc 1 / ATCC 23582 / Paris) OX=456481 GN=rnd PE=4 SV=1 |
| tr\|B0SQT9\|B0SQT9_LEPBP | 1 | 5 | 4 | 10 | 3 | PMT_2 domain-containing protein OS=Leptospira biflexa serovar Patoc (strain Patoc 1 / ATCC 23582 / Paris) OX=456481 GN=LEPBI_I3271 PE=4 SV=1 |
| tr\|B0SQG6\|B0SQG6_LEPBP | 4 | 4 | 2 | 10 | 3 | PPK2 domain-containing protein OS=Leptospira biflexa serovar Patoc (strain Patoc 1 / ATCC 23582 / Paris) OX=456481 GN=LEPBI_I1517 PE=4 SV=1 |
| tr\|B0SKH5\|B0SKH5_LEPBP | 3 | 5 | 2 | 10 | 3 | Putative aerotaxis sensor receptor putative membrane protein OS=Leptospira biflexa serovar Patoc (strain Patoc 1 / ATCC 23582 / Paris) OX=456481 GN=LEPBI_I0595 PE=4 SV=1 |
| tr\|B0SSK5\|B0SSK5_LEPBP | 4 | 4 | 2 | 10 | 3 | Uncharacterized protein OS=Leptospira biflexa serovar Patoc (strain Patoc 1 / ATCC 23582 / Paris) OX=456481 GN=LEPBI_I1993 PE=4 SV=1 |
| tr\|B0SK83\|B0SK83_LEPBP | 1 | 3 | 6 | 10 | 3 | Radical SAM core domain-containing protein OS=Leptospira biflexa serovar Patoc (strain Patoc 1 / ATCC 23582 / Paris) OX=456481 GN=LEPBI_I0185 PE=4 SV=1 |
| tr\|B0SQT3\|B0SQT3_LEPBP | 4 | 2 | 4 | 10 | 3 | Putative glycosyltransferase OS=Leptospira biflexa serovar Patoc (strain Patoc 1 / ATCC 23582 / Paris) OX=456481 GN=LEPBI_I3265 PE=4 SV=1 |
| tr\|B0SUA4\|B0SUA4_LEPBP | 3 | 5 | 2 | 10 | 3 | Uncharacterized protein OS=Leptospira biflexa serovar Patoc (strain Patoc 1 / ATCC 23582 / Paris) OX=456481 GN=LEPBI_II0255 PE=4 SV=1 |
| tr\|B0SL10\|B0SL10_LEPBP | 3 | 3 | 4 | 10 | 3 | Putative RNA methyltransferase OS=Leptospira biflexa serovar Patoc (strain Patoc 1 / ATCC 23582 / Paris) OX=456481 GN=LEPBI_I2413 PE=3 SV=1 |
| tr\|B0SL01\|B0SL01_LEPBP | 4 | 4 | 2 | 10 | 3 | Putative alpha/beta hydrolase putative signal peptide OS=Leptospira biflexa serovar Patoc (strain Patoc 1 / ATCC 23582 / Paris) OX=456481 GN=LEPBI_I2404 PE=4 SV=1 |
| tr\|B0SMQ1\|B0SMQ1_LEPBP | 3 | 4 | 2 | 9 | 3 | Putative ATPase OS=Leptospira biflexa serovar Patoc (strain Patoc 1 / ATCC 23582 / Paris) OX=456481 GN=LEPBI_I2697 PE=4 SV=1 |
| sp\|B0SQ03\|LIPA_LEPBP | 3 | 3 | 3 | 9 | 3 | Lipoyl synthase OS=Leptospira biflexa serovar Patoc (strain Patoc 1 / ATCC 23582 / Paris) OX=456481 GN=lipA PE=3 SV=1 |
| tr\|B0SNZ8\|B0SNZ8_LEPBP | 3 | 2 | 4 | 9 | 3 | Putative hemerythrin HHE cation binding protein OS=Leptospira biflexa serovar Patoc (strain Patoc 1 / ATCC 23582 / Paris) OX=456481 GN=LEPBI_I2942 PE=3 SV=1 |
| tr\|B0SKE9\|B0SKE9_LEPBP | 2 | 4 | 3 | 9 | 3 | UDP-glucose 4-epimerase OS=Leptospira biflexa serovar Patoc (strain Patoc 1 / ATCC 23582 / Paris) OX=456481 GN=galE PE=3 SV=1 |
| tr\|B0SSL5\|B0SSL5_LEPBP | 4 | 2 | 3 | 9 | 3 | Putative glycosyltransferase OS=Leptospira biflexa serovar Patoc (strain Patoc 1 / ATCC 23582 / Paris) OX=456481 GN=LEPBI_I2003 PE=4 SV=1 |
| tr\|B0SKZ4\|B0SKZ4_LEPBP | 4 | 2 | 3 | 9 | 3 | Uncharacterized protein OS=Leptospira biflexa serovar Patoc (strain Patoc 1 / ATCC 23582 / Paris) OX=456481 GN=LEPBI_I2397 PE=4 SV=1 |
| tr\|B0SN41\|B0SN41_LEPBP | 3 | 3 | 3 | 9 | 3 | Putative sodium:solute symporter family protein OS=Leptospira biflexa serovar Patoc (strain Patoc 1 / ATCC 23582 / Paris) OX=456481 GN=LEPBI_I1109 PE=3 SV=1 |
| tr\|B0SNI4\|B0SNI4_LEPBP | 2 | 5 | 2 | 9 | 3 | Putative cAMP-dependent protein kinase OS=Leptospira biflexa serovar Patoc (strain Patoc 1 / ATCC 23582 / Paris) OX=456481 GN=LEPBI_I1149 PE=4 SV=1 |
| tr\|B0SS69\|B0SS69_LEPBP | 3 | 2 | 4 | 9 | 3 | Uncharacterized protein OS=Leptospira biflexa serovar Patoc (strain Patoc 1 / ATCC 23582 / Paris) OX=456481 GN=LEPBI_I1854 PE=4 SV=1 |
| tr\|B0SJ45\|B0SJ45_LEPBP | 2 | 3 | 4 | 9 | 3 | Uncharacterized protein OS=Leptospira biflexa serovar Patoc (strain Patoc 1 / ATCC 23582 / Paris) OX=456481 GN=LEPBI_I0475 PE=4 SV=1 |
| tr\|B0SNV7\|B0SNV7_LEPBP | 1 | 3 | 5 | 9 | 3 | Small ribosomal subunit biogenesis GTPase RsgA OS=Leptospira biflexa serovar Patoc (strain Patoc 1 / ATCC 23582 / Paris) OX=456481 GN=rsgA PE=3 SV=1 |
| tr\|B0SKR6\|B0SKR6_LEPBP | 2 | 5 | 1 | 8 | 3 | Putative cation exchanger putative membrane protein OS=Leptospira biflexa serovar Patoc (strain Patoc 1 / ATCC 23582 / Paris) OX=456481 GN=LEPBI_I0691 PE=4 SV=1 |
| tr\|B0SR35\|B0SR35_LEPBP | 1 | 3 | 4 | 8 | 3 | Putative cyclic nucleotide-binding protein OS=Leptospira biflexa serovar Patoc (strain Patoc 1 / ATCC 23582 / Paris) OX=456481 GN=LEPBI_I1629 PE=3 SV=1 |
| tr\|B0SM30\|B0SM30_LEPBP | 4 | 3 | 1 | 8 | 3 | Uncharacterized protein OS=Leptospira biflexa serovar Patoc (strain Patoc 1 / ATCC 23582 / Paris) OX=456481 GN=LEPBI_I2579 PE=4 SV=1 |
| tr\|B0SU36\|B0SU36_LEPBP | 2 | 4 | 2 | 8 | 3 | Putative methyl-accepting chemotaxis protein tlpA, putative transmembrane protein OS=Leptospira biflexa serovar Patoc (strain Patoc 1 / ATCC 23582 / Paris) OX=456481 GN=LEPBI_II0187 PE=4 SV=1 |
| tr\|B0SM54\|B0SM54_LEPBP | 3 | 2 | 3 | 8 | 3 | 1-deoxy-D-xylulose-5-phosphate synthase OS=Leptospira biflexa serovar Patoc (strain Patoc 1 / ATCC 23582 / Paris) OX=456481 GN=dxs PE=3 SV=1 |
| tr\|B0SJU6\|B0SJU6_LEPBP | 3 | 2 | 3 | 8 | 3 | Putative methyl-accepting chemotaxis protein putative membrane protein OS=Leptospira biflexa serovar Patoc (strain Patoc 1 / ATCC 23582 / Paris) OX=456481 GN=LEPBI_I0116 PE=4 SV=1 |
| tr\|B0SQP5\|B0SQP5_LEPBP | 3 | 3 | 2 | 8 | 3 | Histidine kinase OS=Leptospira biflexa serovar Patoc (strain Patoc 1 / ATCC 23582 / Paris) OX=456481 GN=LEPBI_I3227 PE=4 SV=1 |
| tr\|B0STI1\|B0STI1_LEPBP | 1 | 5 | 2 | 8 | 3 | Anti-sigma factor antagonist OS=Leptospira biflexa serovar Patoc (strain Patoc 1 / ATCC 23582 / Paris) OX=456481 GN=LEPBI_I2328 PE=3 SV=1 |
| tr\|B0SLH0\|B0SLH0_LEPBP | 3 | 2 | 3 | 8 | 3 | Putative type I phosphodiesterase/nucleotide pyrophosphatase OS=Leptospira biflexa serovar Patoc (strain Patoc 1 / ATCC 23582 / Paris) OX=456481 GN=LEPBI_I2467 PE=4 SV=1 |
| tr\|B0SNI9\|B0SNI9_LEPBP | 3 | 1 | 4 | 8 | 3 | Putative cation transporter putative membrane protein putative signal peptide OS=Leptospira biflexa serovar Patoc (strain Patoc 1 / ATCC 23582 / Paris) OX=456481 GN=LEPBI_I1154 PE=4 SV=1 |
| tr\|B0STE6\|B0STE6_LEPBP | 2 | 4 | 2 | 8 | 3 | Probable lipid II flippase MurJ OS=Leptospira biflexa serovar Patoc (strain Patoc 1 / ATCC 23582 / Paris) OX=456481 GN=murJ PE=3 SV=1 |
| tr\|B0SKM7\|B0SKM7_LEPBP | 5 | 2 | 1 | 8 | 3 | Uncharacterized protein OS=Leptospira biflexa serovar Patoc (strain Patoc 1 / ATCC 23582 / Paris) OX=456481 GN=LEPBI_I0651 PE=4 SV=1 |
| sp\|B0SM63\|RRF_LEPBP | 2 | 3 | 3 | 8 | 3 | Ribosome-recycling factor OS=Leptospira biflexa serovar Patoc (strain Patoc 1 / ATCC 23582 / Paris) OX=456481 GN=frr PE=3 SV=1 |
| tr\|B0SL92\|B0SL92_LEPBP | 2 | 4 | 2 | 8 | 3 | Uncharacterized protein OS=Leptospira biflexa serovar Patoc (strain Patoc 1 / ATCC 23582 / Paris) OX=456481 GN=LEPBI_I0768 PE=4 SV=1 |
| tr\|B0SK55\|B0SK55_LEPBP | 3 | 3 | 1 | 7 | 3 | Putative inositol monophosphatase OS=Leptospira biflexa serovar Patoc (strain Patoc 1 / ATCC 23582 / Paris) OX=456481 GN=LEPBI_I0026 PE=4 SV=1 |
| tr\|B0SNI3\|B0SNI3_LEPBP | 4 | 2 | 1 | 7 | 3 | Peptide chain release factor N(5)-glutamine methyltransferase OS=Leptospira biflexa serovar Patoc (strain Patoc 1 / ATCC 23582 / Paris) OX=456481 GN=hemK PE=4 SV=1 |
| tr\|B0SQM0\|B0SQM0_LEPBP | 2 | 2 | 3 | 7 | 3 | NodB homology domain-containing protein OS=Leptospira biflexa serovar Patoc (strain Patoc 1 / ATCC 23582 / Paris) OX=456481 GN=LEPBI_I1571 PE=4 SV=1 |
| tr\|B0SJE4\|B0SJE4_LEPBP | 2 | 2 | 3 | 7 | 3 | Cation efflux protein putative membrane protein OS=Leptospira biflexa serovar Patoc (strain Patoc 1 / ATCC 23582 / Paris) OX=456481 GN=LEPBI_I0298 PE=4 SV=1 |
| tr\|B0SLR5\|B0SLR5_LEPBP | 2 | 3 | 2 | 7 | 3 | Uncharacterized protein OS=Leptospira biflexa serovar Patoc (strain Patoc 1 / ATCC 23582 / Paris) OX=456481 GN=LEPBI_I0840 PE=4 SV=1 |
| tr\|B0SR29\|B0SR29_LEPBP | 3 | 1 | 3 | 7 | 3 | FHA domain-containing protein OS=Leptospira biflexa serovar Patoc (strain Patoc 1 / ATCC 23582 / Paris) OX=456481 GN=LEPBI_I1623 PE=4 SV=1 |
| tr\|B0SQS8\|B0SQS8_LEPBP | 3 | 1 | 3 | 7 | 3 | Putative metalloendopeptidase OS=Leptospira biflexa serovar Patoc (strain Patoc 1 / ATCC 23582 / Paris) OX=456481 GN=LEPBI_I3260 PE=3 SV=1 |
| sp\|B0SJP0\|Y056_LEPBP | 2 | 2 | 2 | 6 | 3 | Probable transcriptional regulatory protein LEPBI_I0056 OS=Leptospira biflexa serovar Patoc (strain Patoc 1 / ATCC 23582 / Paris) OX=456481 GN=LEPBI_I0056 PE=3 SV=1 |
| tr\|B0SS88\|B0SS88_LEPBP | 2 | 3 | 1 | 6 | 3 | Nucleoid-associated protein LEPBI_I1874 OS=Leptospira biflexa serovar Patoc (strain Patoc 1 / ATCC 23582 / Paris) OX=456481 GN=LEPBI_I1874 PE=3 SV=1 |
| **tr\|B0SS28\|B0SS28_LEPBP** | 2 | 2 | 2 | 6 | 3 | Uncharacterized protein OS=Leptospira biflexa serovar Patoc (strain Patoc 1 / ATCC 23582 / Paris) OX=456481 GN=LEPBI_I1812 PE=4 SV=1 |
| tr\|B0SM50\|B0SM50_LEPBP | 1 | 4 | 1 | 6 | 3 | Uncharacterized protein OS=Leptospira biflexa serovar Patoc (strain Patoc 1 / ATCC 23582 / Paris) OX=456481 GN=LEPBI_I2601 PE=4 SV=1 |
| tr\|B0SS89\|B0SS89_LEPBP | 2 | 2 | 2 | 6 | 3 | Putative cyclic-nucleotide-gated cation channel putative membrane protein OS=Leptospira biflexa serovar Patoc (strain Patoc 1 / ATCC 23582 / Paris) OX=456481 GN=LEPBI_I1875 PE=4 SV=1 |
| tr\|B0SMP0\|B0SMP0_LEPBP | 2 | 2 | 2 | 6 | 3 | Cytochrome oxidase assembly protein putative membrane protein OS=Leptospira biflexa serovar Patoc (strain Patoc 1 / ATCC 23582 / Paris) OX=456481 GN=ctaA PE=4 SV=1 |
| tr\|B0SLC2\|B0SLC2_LEPBP | 3 | 2 | 1 | 6 | 3 | ATP synthase subunit a OS=Leptospira biflexa serovar Patoc (strain Patoc 1 / ATCC 23582 / Paris) OX=456481 GN=atpB PE=3 SV=1 |
| tr\|B0SLN8\|B0SLN8_LEPBP | 1 | 2 | 3 | 6 | 3 | Aminotransferase OS=Leptospira biflexa serovar Patoc (strain Patoc 1 / ATCC 23582 / Paris) OX=456481 GN=aatA PE=3 SV=1 |
| tr\|B0STU3\|B0STU3_LEPBP | 2 | 2 | 2 | 6 | 3 | Putative response regulatory protein OS=Leptospira biflexa serovar Patoc (strain Patoc 1 / ATCC 23582 / Paris) OX=456481 GN=LEPBI_II0090 PE=4 SV=1 |
| tr\|B0SNJ8\|B0SNJ8_LEPBP | 2 | 3 | 1 | 6 | 3 | DNA topoisomerase OS=Leptospira biflexa serovar Patoc (strain Patoc 1 / ATCC 23582 / Paris) OX=456481 GN=topA PE=3 SV=1 |
| tr\|B0SNX8\|B0SNX8_LEPBP | 2 | 1 | 3 | 6 | 3 | Uncharacterized protein OS=Leptospira biflexa serovar Patoc (strain Patoc 1 / ATCC 23582 / Paris) OX=456481 GN=LEPBI_I2921 PE=4 SV=1 |
| tr\|B0SQL5\|B0SQL5_LEPBP | 2 | 3 | 1 | 6 | 3 | DNA primase OS=Leptospira biflexa serovar Patoc (strain Patoc 1 / ATCC 23582 / Paris) OX=456481 GN=dnaG PE=3 SV=1 |

| | | | | | | |
|---|---|---|---|---|---|---|
| tr\|B0SLH8\|B0SLH8_LEPBP | 1 | 3 | 1 | 5 | 3 | Putative alcohol dehydrogenase, zinc-containing OS=Leptospira biflexa serovar Patoc (strain Patoc 1 / ATCC 23582 / Paris) OX=456481 GN=LEPBI_I2475 PE=4 SV=1 |
| tr\|B0STE7\|B0STE7_LEPBP | 1 | 3 | 1 | 5 | 3 | STAS domain-containing protein OS=Leptospira biflexa serovar Patoc (strain Patoc 1 / ATCC 23582 / Paris) OX=456481 GN=LEPBI_I2292 PE=4 SV=1 |
| sp\|B0SK33\|RECF_LEPBP | 1 | 2 | 2 | 5 | 3 | DNA replication and repair protein RecF OS=Leptospira biflexa serovar Patoc (strain Patoc 1 / ATCC 23582 / Paris) OX=456481 GN=recF PE=3 SV=1 |
| tr\|B0SMZ7\|B0SMZ7_LEPBP | 1 | 2 | 1 | 4 | 3 | Uncharacterized protein OS=Leptospira biflexa serovar Patoc (strain Patoc 1 / ATCC 23582 / Paris) OX=456481 GN=LEPBI_I1061 PE=4 SV=1 |
| tr\|B0SNM6\|B0SNM6_LEPBP | 1 | 2 | 1 | 4 | 3 | Flagellar hook protein FlgE OS=Leptospira biflexa serovar Patoc (strain Patoc 1 / ATCC 23582 / Paris) OX=456481 GN=flgE PE=3 SV=1 |
| tr\|B0SLV9\|B0SLV9_LEPBP | 2 | 1 | 1 | 4 | 3 | Lactonizing lipase (Triacylglycerol lipase) OS=Leptospira biflexa serovar Patoc (strain Patoc 1 / ATCC 23582 / Paris) OX=456481 GN=lip PE=4 SV=1 |
| tr\|B0SU18\|B0SU18_LEPBP | 1 | 1 | 2 | 4 | 3 | Alanine racemase OS=Leptospira biflexa serovar Patoc (strain Patoc 1 / ATCC 23582 / Paris) OX=456481 GN=alr PE=3 SV=1 |
| tr\|B0SUI0\|B0SUI0_LEPBP | 1 | 1 | 2 | 4 | 3 | Stress response kinase A OS=Leptospira biflexa serovar Patoc (strain Patoc 1 / ATCC 23582 / Paris) OX=456481 GN=srkA PE=3 SV=1 |
| tr\|B0SQD2\|B0SQD2_LEPBP | 2 | 1 | 1 | 4 | 3 | N-acetyl-gamma-glutamyl-phosphate reductase OS=Leptospira biflexa serovar Patoc (strain Patoc 1 / ATCC 23582 / Paris) OX=456481 GN=argC PE=3 SV=1 |
| tr\|B0SKF6\|B0SKF6_LEPBP | 1 | 1 | 2 | 4 | 3 | Dimethylaniline monooxygenase [N-oxide-forming] 5 OS=Leptospira biflexa serovar Patoc (strain Patoc 1 / ATCC 23582 / Paris) OX=456481 GN=LEPBI_I0245 PE=3 SV=1 |
| tr\|B0SML2\|B0SML2_LEPBP | 2 | 1 | 1 | 4 | 3 | Uncharacterized protein OS=Leptospira biflexa serovar Patoc (strain Patoc 1 / ATCC 23582 / Paris) OX=456481 GN=LEPBI_I2657 PE=4 SV=1 |
| sp\|B0SLC5\|ATPD_LEPBP | 19 | 41 | | 60 | 2 | ATP synthase subunit delta OS=Leptospira biflexa serovar Patoc (strain Patoc 1 / ATCC 23582 / Paris) OX=456481 GN=atpH PE=3 SV=1 |
| tr\|B0SMR3\|B0SMR3_LEPBP | | 34 | 4 | 38 | 2 | Uncharacterized protein OS=Leptospira biflexa serovar Patoc (strain Patoc 1 / ATCC 23582 / Paris) OX=456481 GN=LEPBI_I2709 PE=4 SV=1 |
| tr\|B0STL2\|B0STL2_LEPBP | 14 | 23 | | 37 | 2 | Histidine kinase sensor protein OS=Leptospira biflexa serovar Patoc (strain Patoc 1 / ATCC 23582 / Paris) OX=456481 GN=LEPBI_II0008 PE=4 SV=1 |
| tr\|B0SJK6\|B0SJK6_LEPBP | 23 | 12 | | 35 | 2 | Putative DNA mismatch repair protein putative membrane protein OS=Leptospira biflexa serovar Patoc (strain Patoc 1 / ATCC 23582 / Paris) OX=456481 GN=LEPBI_I0334 PE=4 SV=1 |
| tr\|B0SQY8\|B0SQY8_LEPBP | 12 | 22 | | 34 | 2 | Putative methyl-accepting chemotaxis protein OS=Leptospira biflexa serovar Patoc (strain Patoc 1 / ATCC 23582 / Paris) OX=456481 GN=LEPBI_I1582 PE=4 SV=1 |
| tr\|B0SKI5\|B0SKI5_LEPBP | 21 | 10 | | 31 | 2 | Putative FAD binding dehydrogenase OS=Leptospira biflexa serovar Patoc (strain Patoc 1 / ATCC 23582 / Paris) OX=456481 GN=LEPBI_I0606 PE=4 SV=1 |
| tr\|B0SUE3\|B0SUE3_LEPBP | 19 | 12 | | 31 | 2 | ABC-type hemin transport system, ATPase OS=Leptospira biflexa serovar Patoc (strain Patoc 1 / ATCC 23582 / Paris) OX=456481 GN=hemV PE=4 SV=1 |
| tr\|B0SJK5\|B0SJK5_LEPBP | | 4 | 26 | 30 | 2 | SH3b domain-containing protein OS=Leptospira biflexa serovar Patoc (strain Patoc 1 / ATCC 23582 / Paris) OX=456481 GN=LEPBI_I0332 PE=4 SV=1 |
| tr\|B0STZ2\|B0STZ2_LEPBP | 13 | 15 | | 28 | 2 | Uncharacterized protein OS=Leptospira biflexa serovar Patoc (strain Patoc 1 / ATCC 23582 / Paris) OX=456481 GN=LEPBI_II0141 PE=4 SV=1 |
| tr\|B0SP25\|B0SP25_LEPBP | 17 | 10 | | 27 | 2 | DUF374 domain-containing protein OS=Leptospira biflexa serovar Patoc (strain Patoc 1 / ATCC 23582 / Paris) OX=456481 GN=LEPBI_I2969 PE=4 SV=1 |
| tr\|B0SJI7\|B0SJI7_LEPBP | 16 | 10 | | 26 | 2 | Putative inner membrane protein involved in colicin E2 resistance OS=Leptospira biflexa serovar Patoc (strain Patoc 1 / ATCC 23582 / Paris) OX=456481 GN=creD PE=4 SV=1 |
| tr\|B0SQV7\|B0SQV7_LEPBP | | 6 | 19 | 25 | 2 | Putative polyol dehydrogenase OS=Leptospira biflexa serovar Patoc (strain Patoc 1 / ATCC 23582 / Paris) OX=456481 GN=LEPBI_I3289 PE=4 SV=1 |
| tr\|B0SNL5\|B0SNL5_LEPBP | 15 | 10 | | 25 | 2 | Sulfite reductase [NADPH] flavoprotein alpha-component (SIR-FP) OS=Leptospira biflexa serovar Patoc (strain Patoc 1 / ATCC 23582 / Paris) OX=456481 GN=cysJ PE=4 SV=1 |
| tr\|B0SS00\|B0SS00_LEPBP | | 6 | 17 | 23 | 2 | Uncharacterized protein OS=Leptospira biflexa serovar Patoc (strain Patoc 1 / ATCC 23582 / Paris) OX=456481 GN=LEPBI_I3476 PE=4 SV=1 |
| tr\|B0SRV8\|B0SRV8_LEPBP | 8 | 15 | | 23 | 2 | TonB-dependent receptor protein putative signal peptide OS=Leptospira biflexa serovar Patoc (strain Patoc 1 / ATCC 23582 / Paris) OX=456481 GN=LEPBI_I3432 PE=3 SV=1 |
| tr\|B0SSU0\|B0SSU0_LEPBP | 5 | 18 | | 23 | 2 | Signal peptidase I OS=Leptospira biflexa serovar Patoc (strain Patoc 1 / ATCC 23582 / Paris) OX=456481 GN=LEPBI_I2078 PE=3 SV=1 |
| tr\|B0SRE1\|B0SRE1_LEPBP | 5 | 18 | | 23 | 2 | Cation efflux system protein, acrB/acrD/acrF family putative membrane protein OS=Leptospira biflexa serovar Patoc (strain Patoc 1 / ATCC 23582 / Paris) OX=456481 GN=LEPBI_I3369 PE=3 SV=1 |
| sp\|B0SQA1\|GCST_LEPBP | | 5 | 17 | 22 | 2 | Aminomethyltransferase OS=Leptospira biflexa serovar Patoc (strain Patoc 1 / ATCC 23582 / Paris) OX=456481 GN=gcvT PE=3 SV=1 |
| tr\|B0SSN3\|B0SSN3_LEPBP | 15 | 7 | | 22 | 2 | Uncharacterized protein OS=Leptospira biflexa serovar Patoc (strain Patoc 1 / ATCC 23582 / Paris) OX=456481 GN=LEPBI_I2021 PE=4 SV=1 |
| tr\|B0SKX4\|B0SKX4_LEPBP | 13 | 9 | | 22 | 2 | Uncharacterized protein OS=Leptospira biflexa serovar Patoc (strain Patoc 1 / ATCC 23582 / Paris) OX=456481 GN=LEPBI_I2376 PE=4 SV=1 |
| tr\|B0SSX0\|B0SSX0_LEPBP | | 2 | 18 | 20 | 2 | Uncharacterized protein OS=Leptospira biflexa serovar Patoc (strain Patoc 1 / ATCC 23582 / Paris) OX=456481 GN=LEPBI_I2108 PE=4 SV=1 |
| tr\|B0SRQ4\|B0SRQ4_LEPBP | 14 | 6 | | 20 | 2 | Putative sigma factor sigB regulation protein RsbU putative membrane protein putative signal peptide OS=Leptospira biflexa serovar Patoc (strain Patoc 1 / ATCC 23582 / Paris) OX=456481 GN=LEPBI_I1743 PE=4 SV=1 |
| tr\|B0SPH9\|B0SPH9_LEPBP | 9 | 11 | | 20 | 2 | Biopolymer transport ExbB protein putative membrane protein OS=Leptospira biflexa serovar Patoc (strain Patoc 1 / ATCC 23582 / Paris) OX=456481 GN=exbB2 PE=3 SV=1 |
| tr\|B0STX3\|B0STX3_LEPBP | 11 | 9 | | 20 | 2 | Putative Dolichyl-phosphate beta-D-mannosyltransferase OS=Leptospira biflexa serovar Patoc (strain Patoc 1 / ATCC 23582 / Paris) OX=456481 GN=LEPBI_II0121 PE=4 SV=1 |
| tr\|B0SJF3\|B0SJF3_LEPBP | 7 | 13 | | 20 | 2 | Uncharacterized protein OS=Leptospira biflexa serovar Patoc (strain Patoc 1 / ATCC 23582 / Paris) OX=456481 GN=LEPBI_I0307 PE=4 SV=1 |
| tr\|B0SJZ8\|B0SJZ8_LEPBP | 10 | 10 | | 20 | 2 | Putative cAMP-binding protein OS=Leptospira biflexa serovar Patoc (strain Patoc 1 / ATCC 23582 / Paris) OX=456481 GN=LEPBI_I0156 PE=3 SV=1 |
| tr\|B0SLR6\|B0SLR6_LEPBP | 3 | | 16 | 19 | 2 | Uncharacterized protein OS=Leptospira biflexa serovar Patoc (strain Patoc 1 / ATCC 23582 / Paris) OX=456481 GN=LEPBI_I0841 PE=4 SV=1 |
| tr\|B0SRP9\|B0SRP9_LEPBP | | 8 | 11 | 19 | 2 | Putative peptidase, M48 family OS=Leptospira biflexa serovar Patoc (strain Patoc 1 / ATCC 23582 / Paris) OX=456481 GN=LEPBI_I1738 PE=3 SV=1 |
| sp\|B0STK1\|PANB_LEPBP | 9 | 10 | | 19 | 2 | 3-methyl-2-oxobutanoate hydroxymethyltransferase OS=Leptospira biflexa serovar Patoc (strain Patoc 1 / ATCC 23582 / Paris) OX=456481 GN=panB PE=3 SV=1 |
| tr\|B0SN79\|B0SN79_LEPBP | 13 | 6 | | 19 | 2 | Cyclic pyranopterin monophosphate synthase OS=Leptospira biflexa serovar Patoc (strain Patoc 1 / ATCC 23582 / Paris) OX=456481 GN=moaCB PE=4 SV=1 |
| tr\|B0SJD1\|B0SJD1_LEPBP | 7 | 12 | | 19 | 2 | Putative phosphoglycerol transferase, alkaline phosphatase superfamily putative membrane protein OS=Leptospira biflexa serovar Patoc (strain Patoc 1 / ATCC 23582 / Paris) OX=456481 GN=LEPBI_I0556 PE=4 SV=1 |
| tr\|B0SST2\|B0SST2_LEPBP | | 5 | 13 | 18 | 2 | Uncharacterized protein OS=Leptospira biflexa serovar Patoc (strain Patoc 1 / ATCC 23582 / Paris) OX=456481 GN=LEPBI_I2070 PE=4 SV=1 |
| tr\|B0SJ20\|B0SJ20_LEPBP | 11 | 7 | | 18 | 2 | Cholesterol oxidase OS=Leptospira biflexa serovar Patoc (strain Patoc 1 / ATCC 23582 / Paris) OX=456481 GN=choB PE=4 SV=1 |
| tr\|B0SQV1\|B0SQV1_LEPBP | 5 | 13 | | 18 | 2 | Uncharacterized protein OS=Leptospira biflexa serovar Patoc (strain Patoc 1 / ATCC 23582 / Paris) OX=456481 GN=LEPBI_I3283 PE=4 SV=1 |
| tr\|B0SP00\|B0SP00_LEPBP | 7 | 11 | | 18 | 2 | Putative adenylate/guanylate cyclase putative membrane protein OS=Leptospira biflexa serovar Patoc (strain Patoc 1 / ATCC 23582 / Paris) OX=456481 GN=LEPBI_I2944 PE=4 SV=1 |
| tr\|B0SR38\|B0SR38_LEPBP | 6 | 11 | | 17 | 2 | Biopolymer transport ExbD protein OS=Leptospira biflexa serovar Patoc (strain Patoc 1 / ATCC 23582 / Paris) OX=456481 GN=exbD4 PE=3 SV=1 |
| tr\|B0SSS9\|B0SSS9_LEPBP | 7 | 10 | | 17 | 2 | Putative glutathione S-transferase related protein OS=Leptospira biflexa serovar Patoc (strain Patoc 1 / ATCC 23582 / Paris) OX=456481 GN=LEPBI_I2067 PE=4 SV=1 |
| tr\|B0SLC9\|B0SLC9_LEPBP | 5 | 12 | | 17 | 2 | ATP synthase epsilon chain OS=Leptospira biflexa serovar Patoc (strain Patoc 1 / ATCC 23582 / Paris) OX=456481 GN=atpC PE=3 SV=1 |
| tr\|B0SL09\|B0SL09_LEPBP | 7 | 10 | | 17 | 2 | Putative adenylate or guanylate cyclase putative membrane protein OS=Leptospira biflexa serovar Patoc (strain Patoc 1 / ATCC 23582 / Paris) OX=456481 GN=LEPBI_I2412 PE=4 SV=1 |
| tr\|B0SK40\|B0SK40_LEPBP | | 4 | 12 | 16 | 2 | Uncharacterized protein OS=Leptospira biflexa serovar Patoc (strain Patoc 1 / ATCC 23582 / Paris) OX=456481 GN=LEPBI_I0010 PE=4 SV=1 |
| tr\|B0SK96\|B0SK96_LEPBP | 6 | 10 | | 16 | 2 | Flagellar protein FliL OS=Leptospira biflexa serovar Patoc (strain Patoc 1 / ATCC 23582 / Paris) OX=456481 GN=fliL PE=3 SV=1 |
| tr\|B0SSA9\|B0SSA9_LEPBP | 8 | 8 | | 16 | 2 | Uncharacterized protein OS=Leptospira biflexa serovar Patoc (strain Patoc 1 / ATCC 23582 / Paris) OX=456481 GN=LEPBI_I1896 PE=4 SV=1 |
| tr\|B0SKY1\|B0SKY1_LEPBP | 8 | 8 | | 16 | 2 | Uncharacterized protein OS=Leptospira biflexa serovar Patoc (strain Patoc 1 / ATCC 23582 / Paris) OX=456481 GN=LEPBI_I2384 PE=3 SV=1 |
| tr\|B0SLQ8\|B0SLQ8_LEPBP | 6 | 10 | | 16 | 2 | DNA mismatch repair protein MutS OS=Leptospira biflexa serovar Patoc (strain Patoc 1 / ATCC 23582 / Paris) OX=456481 GN=mutS PE=3 SV=1 |
| tr\|B0SLB5\|B0SLB5_LEPBP | 9 | 7 | | 16 | 2 | Acetolactate synthase OS=Leptospira biflexa serovar Patoc (strain Patoc 1 / ATCC 23582 / Paris) OX=456481 GN=ilvB1 PE=3 SV=1 |
| tr\|B0SRU7\|B0SRU7_LEPBP | 8 | 8 | | 16 | 2 | Putative enoyl-CoA hydratase OS=Leptospira biflexa serovar Patoc (strain Patoc 1 / ATCC 23582 / Paris) OX=456481 GN=LEPBI_I1786 PE=4 SV=1 |
| tr\|B0SMM4\|B0SMM4_LEPBP | | 1 | 14 | 15 | 2 | Uncharacterized protein OS=Leptospira biflexa serovar Patoc (strain Patoc 1 / ATCC 23582 / Paris) OX=456481 GN=LEPBI_I2670 PE=4 SV=1 |
| tr\|B0SSU8\|B0SSU8_LEPBP | | 7 | 8 | 15 | 2 | Putative two-component response regulator OS=Leptospira biflexa serovar Patoc (strain Patoc 1 / ATCC 23582 / Paris) OX=456481 GN=LEPBI_I2086 PE=4 SV=1 |
| tr\|B0SLQ7\|B0SLQ7_LEPBP | | 6 | 9 | 15 | 2 | O-phosphoserine phosphohydrolase OS=Leptospira biflexa serovar Patoc (strain Patoc 1 / ATCC 23582 / Paris) OX=456481 GN=serB PE=3 SV=1 |

| | | | | | | |
|---|---|---|---|---|---|---|
| tr\|B0SRY5\|B0SRY5_LEPBP | 8 | 7 | | 15 | 2 | DNA polymerase III subunit gamma/tau OS=Leptospira biflexa serovar Patoc (strain Patoc 1 / ATCC 23582 / Paris) OX=456481 GN=dnaX1 PE=3 SV=1 |
| tr\|B0SQY5\|B0SQY5_LEPBP | | 2 | 12 | 14 | 2 | Protein-glutamate methylesterase/protein-glutamine glutaminase OS=Leptospira biflexa serovar Patoc (strain Patoc 1 / ATCC 23582 / Paris) OX=456481 GN=cheB4 PE=3 SV=1 |
| tr\|B0SN60\|B0SN60_LEPBP | | 3 | 11 | 14 | 2 | Tryptophan--tRNA ligase OS=Leptospira biflexa serovar Patoc (strain Patoc 1 / ATCC 23582 / Paris) OX=456481 GN=trpS PE=3 SV=1 |
| tr\|B0STH7\|B0STH7_LEPBP | | 6 | 8 | 14 | 2 | ABC-type transport system, periplasmic binding protein OS=Leptospira biflexa serovar Patoc (strain Patoc 1 / ATCC 23582 / Paris) OX=456481 GN=LEPBI_I2322 PE=4 SV=1 |
| tr\|B0STJ9\|B0STJ9_LEPBP | | 2 | 12 | 14 | 2 | DNA replication protein DnaC OS=Leptospira biflexa serovar Patoc (strain Patoc 1 / ATCC 23582 / Paris) OX=456481 GN=dnaC PE=4 SV=1 |
| tr\|B0STJ6\|B0STJ6_LEPBP | 10 | 4 | | 14 | 2 | ABC-type transport system, ATPase OS=Leptospira biflexa serovar Patoc (strain Patoc 1 / ATCC 23582 / Paris) OX=456481 GN=lolD PE=4 SV=1 |
| sp\|B0SJ57\|RS20_LEPBP | 4 | 10 | | 14 | 2 | 30S ribosomal protein S20 OS=Leptospira biflexa serovar Patoc (strain Patoc 1 / ATCC 23582 / Paris) OX=456481 GN=rpsT PE=3 SV=1 |
| tr\|B0STN1\|B0STN1_LEPBP | 7 | 6 | | 13 | 2 | Uncharacterized protein OS=Leptospira biflexa serovar Patoc (strain Patoc 1 / ATCC 23582 / Paris) OX=456481 GN=LEPBI_II0027 PE=4 SV=1 |
| tr\|B0SSP0\|B0SSP0_LEPBP | 6 | 7 | | 13 | 2 | Putative methionyl-tRNA formyltransferase OS=Leptospira biflexa serovar Patoc (strain Patoc 1 / ATCC 23582 / Paris) OX=456481 GN=LEPBI_I2028 PE=4 SV=1 |
| tr\|B0SPX3\|B0SPX3_LEPBP | 6 | 7 | | 13 | 2 | Uncharacterized protein OS=Leptospira biflexa serovar Patoc (strain Patoc 1 / ATCC 23582 / Paris) OX=456481 GN=LEPBI_I1431 PE=4 SV=1 |
| tr\|B0SM22\|B0SM22_LEPBP | | 5 | 7 | 12 | 2 | Uncharacterized protein OS=Leptospira biflexa serovar Patoc (strain Patoc 1 / ATCC 23582 / Paris) OX=456481 GN=LEPBI_I2571 PE=4 SV=1 |
| tr\|B0SKN9\|B0SKN9_LEPBP | | 3 | 9 | 12 | 2 | Putative triacylglycerol lipase OS=Leptospira biflexa serovar Patoc (strain Patoc 1 / ATCC 23582 / Paris) OX=456481 GN=LEPBI_I0663 PE=4 SV=1 |
| tr\|B0SN35\|B0SN35_LEPBP | | 4 | 8 | 12 | 2 | Putative metallo-dependent phosphatase OS=Leptospira biflexa serovar Patoc (strain Patoc 1 / ATCC 23582 / Paris) OX=456481 GN=LEPBI_I1103 PE=4 SV=1 |
| tr\|B0SQU3\|B0SQU3_LEPBP | 7 | 5 | | 12 | 2 | Methylenetetrahydrofolate reductase OS=Leptospira biflexa serovar Patoc (strain Patoc 1 / ATCC 23582 / Paris) OX=456481 GN=metF PE=3 SV=1 |
| tr\|B0SSL1\|B0SSL1_LEPBP | 9 | 3 | | 12 | 2 | Glyco_trans_2-like domain-containing protein OS=Leptospira biflexa serovar Patoc (strain Patoc 1 / ATCC 23582 / Paris) OX=456481 GN=LEPBI_I1999 PE=4 SV=1 |
| tr\|B0SS48\|B0SS48_LEPBP | 7 | 5 | | 12 | 2 | Magnesium transporter MgtE OS=Leptospira biflexa serovar Patoc (strain Patoc 1 / ATCC 23582 / Paris) OX=456481 GN=LEPBI_I1832 PE=3 SV=1 |
| tr\|B0SP66\|B0SP66_LEPBP | 5 | 7 | | 12 | 2 | Putative sulfatase putative membrane protein OS=Leptospira biflexa serovar Patoc (strain Patoc 1 / ATCC 23582 / Paris) OX=456481 GN=LEPBI_I1280 PE=4 SV=1 |
| tr\|B0SJN1\|B0SJN1_LEPBP | 4 | 8 | | 12 | 2 | PPM-type phosphatase domain-containing protein OS=Leptospira biflexa serovar Patoc (strain Patoc 1 / ATCC 23582 / Paris) OX=456481 GN=LEPBI_I0448 PE=4 SV=1 |
| tr\|B0SRJ8\|B0SRJ8_LEPBP | 2 | 10 | | 12 | 2 | Vitamin B12-dependent ribonucleotide reductase OS=Leptospira biflexa serovar Patoc (strain Patoc 1 / ATCC 23582 / Paris) OX=456481 GN=nrdA PE=3 SV=1 |
| tr\|B0SN52\|B0SN52_LEPBP | | 3 | 8 | 11 | 2 | Ribosomal protein L11 methyltransferase (L11 Mtase) OS=Leptospira biflexa serovar Patoc (strain Patoc 1 / ATCC 23582 / Paris) OX=456481 GN=prmA PE=4 SV=1 |
| tr\|B0SSN5\|B0SSN5_LEPBP | | 4 | 7 | 11 | 2 | Uncharacterized protein OS=Leptospira biflexa serovar Patoc (strain Patoc 1 / ATCC 23582 / Paris) OX=456481 GN=LEPBI_I2023 PE=4 SV=1 |
| tr\|B0SRN0\|B0SRN0_LEPBP | | 9 | 2 | 11 | 2 | Putative ATP-dependent helicase OS=Leptospira biflexa serovar Patoc (strain Patoc 1 / ATCC 23582 / Paris) OX=456481 GN=LEPBI_I1719 PE=4 SV=1 |
| sp\|Q5XPH7\|FEOB_LEPBP | 7 | 4 | | 11 | 2 | Fe(2+) transporter FeoB OS=Leptospira biflexa serovar Patoc (strain Patoc 1 / ATCC 23582 / Paris) OX=456481 GN=feoB PE=3 SV=1 |
| tr\|B0SP77\|B0SP77_LEPBP | 6 | 5 | | 11 | 2 | 2-isopropylmalate synthase (Alpha-isopropylmalate synthase Alpha-IPM synthetase 1) OS=Leptospira biflexa serovar Patoc (strain Patoc 1 / ATCC 23582 / Paris) OX=456481 GN=leuA1 PE=3 SV=1 |
| tr\|B0SKF7\|B0SKF7_LEPBP | 7 | 4 | | 11 | 2 | Metallophos domain-containing protein OS=Leptospira biflexa serovar Patoc (strain Patoc 1 / ATCC 23582 / Paris) OX=456481 GN=LEPBI_I0246 PE=4 SV=1 |
| tr\|B0SU23\|B0SU23_LEPBP | 7 | 4 | | 11 | 2 | Uncharacterized protein OS=Leptospira biflexa serovar Patoc (strain Patoc 1 / ATCC 23582 / Paris) OX=456481 GN=LEPBI_II0172 PE=4 SV=1 |
| tr\|B0SQV6\|B0SQV6_LEPBP | 2 | 9 | | 11 | 2 | Uncharacterized protein OS=Leptospira biflexa serovar Patoc (strain Patoc 1 / ATCC 23582 / Paris) OX=456481 GN=LEPBI_I3288 PE=4 SV=1 |
| tr\|B0SNC0\|B0SNC0_LEPBP | 5 | | 5 | 10 | 2 | Uncharacterized protein OS=Leptospira biflexa serovar Patoc (strain Patoc 1 / ATCC 23582 / Paris) OX=456481 GN=LEPBI_I2812 PE=4 SV=1 |
| tr\|B0SRW6\|B0SRW6_LEPBP | 6 | 4 | | 10 | 2 | ATP-dependent RNA helicase OS=Leptospira biflexa serovar Patoc (strain Patoc 1 / ATCC 23582 / Paris) OX=456481 GN=hepA PE=4 SV=1 |
| tr\|B0SMX6\|B0SMX6_LEPBP | 4 | 6 | | 10 | 2 | Putative rRNA methyltransferase OS=Leptospira biflexa serovar Patoc (strain Patoc 1 / ATCC 23582 / Paris) OX=456481 GN=LEPBI_I1040 PE=4 SV=1 |
| tr\|B0SME9\|B0SME9_LEPBP | 3 | 7 | | 10 | 2 | ATP-dependent Clp protease proteolytic subunit OS=Leptospira biflexa serovar Patoc (strain Patoc 1 / ATCC 23582 / Paris) OX=456481 GN=clpP2 PE=3 SV=1 |
| tr\|B0SQ00\|B0SQ00_LEPBP | 4 | 6 | | 10 | 2 | Uncharacterized protein OS=Leptospira biflexa serovar Patoc (strain Patoc 1 / ATCC 23582 / Paris) OX=456481 GN=LEPBI_I1459 PE=4 SV=1 |
| tr\|B0SSS6\|B0SSS6_LEPBP | 4 | 6 | | 10 | 2 | Uncharacterized protein OS=Leptospira biflexa serovar Patoc (strain Patoc 1 / ATCC 23582 / Paris) OX=456481 GN=LEPBI_I2064 PE=4 SV=1 |
| sp\|B0SPF0\|PCKA_LEPBP | 4 | 6 | | 10 | 2 | Phosphoenolpyruvate carboxykinase (ATP) OS=Leptospira biflexa serovar Patoc (strain Patoc 1 / ATCC 23582 / Paris) OX=456481 GN=pckA PE=3 SV=1 |
| tr\|B0SNH1\|B0SNH1_LEPBP | 5 | 5 | | 10 | 2 | Putative nucleoside-diphosphate-sugar epimerase OS=Leptospira biflexa serovar Patoc (strain Patoc 1 / ATCC 23582 / Paris) OX=456481 GN=LEPBI_I2869 PE=4 SV=1 |
| tr\|B0SM31\|B0SM31_LEPBP | 5 | 5 | | 10 | 2 | Uncharacterized protein OS=Leptospira biflexa serovar Patoc (strain Patoc 1 / ATCC 23582 / Paris) OX=456481 GN=LEPBI_I2580 PE=4 SV=1 |
| sp\|B0SSU4\|TPIS_LEPBP | 4 | 6 | | 10 | 2 | Triosephosphate isomerase OS=Leptospira biflexa serovar Patoc (strain Patoc 1 / ATCC 23582 / Paris) OX=456481 GN=tpiA PE=3 SV=1 |
| tr\|B0STW4\|B0STW4_LEPBP | 4 | 6 | | 10 | 2 | Uncharacterized protein OS=Leptospira biflexa serovar Patoc (strain Patoc 1 / ATCC 23582 / Paris) OX=456481 GN=LEPBI_II0112 PE=4 SV=1 |
| tr\|B0SU60\|B0SU60_LEPBP | 6 | | 3 | 9 | 2 | Putative 3-oxoacyl-[acyl-carrier protein] reductase OS=Leptospira biflexa serovar Patoc (strain Patoc 1 / ATCC 23582 / Paris) OX=456481 GN=LEPBI_II0211 PE=3 SV=1 |
| tr\|B0STU5\|B0STU5_LEPBP | | 5 | 4 | 9 | 2 | Putative transport system kinase putative ArgK protein OS=Leptospira biflexa serovar Patoc (strain Patoc 1 / ATCC 23582 / Paris) OX=456481 GN=LEPBI_II0092 PE=3 SV=1 |
| tr\|B0SNL9\|B0SNL9_LEPBP | 4 | 5 | | 9 | 2 | Adenylyl-sulfate reductase (thioredoxin) OS=Leptospira biflexa serovar Patoc (strain Patoc 1 / ATCC 23582 / Paris) OX=456481 GN=cysH PE=3 SV=1 |
| tr\|B0SKZ1\|B0SKZ1_LEPBP | 2 | 7 | | 9 | 2 | 5-formyltetrahydrofolate cyclo-ligase OS=Leptospira biflexa serovar Patoc (strain Patoc 1 / ATCC 23582 / Paris) OX=456481 GN=LEPBI_I2394 PE=3 SV=1 |
| tr\|B0SLR9\|B0SLR9_LEPBP | 2 | 7 | | 9 | 2 | HATPase_c domain-containing protein OS=Leptospira biflexa serovar Patoc (strain Patoc 1 / ATCC 23582 / Paris) OX=456481 GN=LEPBI_I0844 PE=4 SV=1 |
| tr\|B0SMW2\|B0SMW2_LEPBP | 3 | 6 | | 9 | 2 | Uncharacterized protein OS=Leptospira biflexa serovar Patoc (strain Patoc 1 / ATCC 23582 / Paris) OX=456481 GN=LEPBI_I2758 PE=4 SV=1 |
| tr\|B0SRK5\|B0SRK5_LEPBP | 5 | 4 | | 9 | 2 | Uncharacterized protein OS=Leptospira biflexa serovar Patoc (strain Patoc 1 / ATCC 23582 / Paris) OX=456481 GN=LEPBI_I1694 PE=4 SV=1 |
| tr\|B0SKL3\|B0SKL3_LEPBP | 4 | 5 | | 9 | 2 | Penicillin-insensitive transglycosylase OS=Leptospira biflexa serovar Patoc (strain Patoc 1 / ATCC 23582 / Paris) OX=456481 GN=mrcA PE=4 SV=1 |
| tr\|B0SMP4\|B0SMP4_LEPBP | 3 | 6 | | 9 | 2 | Putative phosphoserine phosphatase putative membrane protein OS=Leptospira biflexa serovar Patoc (strain Patoc 1 / ATCC 23582 / Paris) OX=456481 GN=LEPBI_I2690 PE=4 SV=1 |
| tr\|B0ST42\|B0ST42_LEPBP | 4 | 5 | | 9 | 2 | Uncharacterized protein OS=Leptospira biflexa serovar Patoc (strain Patoc 1 / ATCC 23582 / Paris) OX=456481 GN=LEPBI_I2183 PE=4 SV=1 |
| tr\|B0SJR8\|B0SJR8_LEPBP | 6 | 3 | | 9 | 2 | Putative sensor protein OS=Leptospira biflexa serovar Patoc (strain Patoc 1 / ATCC 23582 / Paris) OX=456481 GN=LEPBI_I0093 PE=4 SV=1 |
| tr\|B0STU0\|B0STU0_LEPBP | 3 | 6 | | 9 | 2 | Putative acriflavine resistance protein D putative transmembrane protein OS=Leptospira biflexa serovar Patoc (strain Patoc 1 / ATCC 23582 / Paris) OX=456481 GN=LEPBI_II0087 PE=4 SV=1 |
| tr\|B0SJ66\|B0SJ66_LEPBP | | 2 | 6 | 8 | 2 | Phosphoglycerate mutase (2,3-diphosphoglycerate-independent) OS=Leptospira biflexa serovar Patoc (strain Patoc 1 / ATCC 23582 / Paris) OX=456481 GN=gpmI PE=3 SV=1 |
| tr\|B0SR21\|B0SR21_LEPBP | | 6 | 2 | 8 | 2 | GMP synthase (glutamine-hydrolyzing) OS=Leptospira biflexa serovar Patoc (strain Patoc 1 / ATCC 23582 / Paris) OX=456481 GN=guaA PE=4 SV=1 |
| tr\|B0SL91\|B0SL91_LEPBP | 2 | | 6 | 8 | 2 | Putative two-component response regulator OS=Leptospira biflexa serovar Patoc (strain Patoc 1 / ATCC 23582 / Paris) OX=456481 GN=LEPBI_I0767 PE=4 SV=1 |
| tr\|B0SSM1\|B0SSM1_LEPBP | 6 | | 2 | 8 | 2 | Putative glycosyltransferase OS=Leptospira biflexa serovar Patoc (strain Patoc 1 / ATCC 23582 / Paris) OX=456481 GN=LEPBI_I2009 PE=4 SV=1 |
| tr\|B0SJ77\|B0SJ77_LEPBP | | 4 | 4 | 8 | 2 | Uncharacterized protein OS=Leptospira biflexa serovar Patoc (strain Patoc 1 / ATCC 23582 / Paris) OX=456481 GN=LEPBI_I0489 PE=4 SV=1 |
| tr\|B0STI4\|B0STI4_LEPBP | | 2 | 6 | 8 | 2 | Peptide chain release factor 2 OS=Leptospira biflexa serovar Patoc (strain Patoc 1 / ATCC 23582 / Paris) OX=456481 GN=prfB PE=3 SV=1 |
| tr\|B0SU47\|B0SU47_LEPBP | 3 | 5 | | 8 | 2 | Putative short-chain dehydrogenase/reductase SDR OS=Leptospira biflexa serovar Patoc (strain Patoc 1 / ATCC 23582 / Paris) OX=456481 GN=LEPBI_II0198 PE=4 SV=1 |
| tr\|B0SRQ7\|B0SRQ7_LEPBP | 4 | 4 | | 8 | 2 | Uncharacterized protein OS=Leptospira biflexa serovar Patoc (strain Patoc 1 / ATCC 23582 / Paris) OX=456481 GN=LEPBI_I1746 PE=4 SV=1 |
| tr\|B0SQZ3\|B0SQZ3_LEPBP | 6 | 2 | | 8 | 2 | Putative sensor protein OS=Leptospira biflexa serovar Patoc (strain Patoc 1 / ATCC 23582 / Paris) OX=456481 GN=LEPBI_I1587 PE=4 SV=1 |
| tr\|B0SL79\|B0SL79_LEPBP | 6 | 2 | | 8 | 2 | Putative sterol desaturase family protein putative membrane protein OS=Leptospira biflexa serovar Patoc (strain Patoc 1 / ATCC 23582 / Paris) OX=456481 GN=LEPBI_I0754 PE=4 SV=1 |
| tr\|B0SP67\|B0SP67_LEPBP | 3 | 5 | | 8 | 2 | Carbamoyl-phosphate synthase large chain OS=Leptospira biflexa serovar Patoc (strain Patoc 1 / ATCC 23582 / Paris) OX=456481 GN=carB PE=3 SV=1 |
| tr\|B0SJF5\|B0SJF5_LEPBP | 5 | 3 | | 8 | 2 | Uncharacterized protein OS=Leptospira biflexa serovar Patoc (strain Patoc 1 / ATCC 23582 / Paris) OX=456481 GN=LEPBI_I0537 PE=4 SV=1 |

| | | | | | | |
|---|---|---|---|---|---|---|
| tr\|B0SP41\|B0SP41_LEPBP | 4 | 4 | | 8 | 2 | TPM_phosphatase domain-containing protein OS=Leptospira biflexa serovar Patoc (strain Patoc 1 / ATCC 23582 / Paris) OX=456481 GN=LEPBI_I2985 PE=4 SV=1 |
| tr\|B0SKY0\|B0SKY0_LEPBP | 2 | 6 | | 8 | 2 | Putative alginate o-acetyltransferase putative membrane protein OS=Leptospira biflexa serovar Patoc (strain Patoc 1 / ATCC 23582 / Paris) OX=456481 GN=LEPBI_I2383 PE=3 SV=1 |
| tr\|B0SSR0\|B0SSR0_LEPBP | 2 | 6 | | 8 | 2 | Putative sensor protein OS=Leptospira biflexa serovar Patoc (strain Patoc 1 / ATCC 23582 / Paris) OX=456481 GN=LEPBI_I2048 PE=4 SV=1 |
| tr\|B0SRJ6\|B0SRJ6_LEPBP | 3 | 5 | | 8 | 2 | Uncharacterized protein OS=Leptospira biflexa serovar Patoc (strain Patoc 1 / ATCC 23582 / Paris) OX=456481 GN=LEPBI_I1684 PE=4 SV=1 |
| tr\|B0SM20\|B0SM20_LEPBP | 4 | 4 | | 8 | 2 | ABC-type transport system, permease putative membrane protein OS=Leptospira biflexa serovar Patoc (strain Patoc 1 / ATCC 23582 / Paris) OX=456481 GN=LEPBI_I2569 PE=4 SV=1 |
| tr\|B0SPR5\|B0SPR5_LEPBP | 5 | 3 | | 8 | 2 | Putative cytochrome c_like protein OS=Leptospira biflexa serovar Patoc (strain Patoc 1 / ATCC 23582 / Paris) OX=456481 GN=LEPBI_I1372 PE=4 SV=1 |
| sp\|B0ST13\|MUTL_LEPBP | 5 | 3 | | 8 | 2 | DNA mismatch repair protein MutL OS=Leptospira biflexa serovar Patoc (strain Patoc 1 / ATCC 23582 / Paris) OX=456481 GN=mutL PE=3 SV=1 |
| tr\|B0SN69\|B0SN69_LEPBP | 4 | 4 | | 8 | 2 | Amino_oxidase domain-containing protein OS=Leptospira biflexa serovar Patoc (strain Patoc 1 / ATCC 23582 / Paris) OX=456481 GN=LEPBI_I1139 PE=4 SV=1 |
| tr\|B0SNH4\|B0SNH4_LEPBP | | 3 | 4 | 7 | 2 | Arsenate reductase (Arsenical pump modifier) OS=Leptospira biflexa serovar Patoc (strain Patoc 1 / ATCC 23582 / Paris) OX=456481 GN=arsC PE=4 SV=1 |
| tr\|B0SR17\|B0SR17_LEPBP | | 3 | 4 | 7 | 2 | Putative hydrolase putative signal peptide OS=Leptospira biflexa serovar Patoc (strain Patoc 1 / ATCC 23582 / Paris) OX=456481 GN=LEPBI_I1611 PE=4 SV=1 |
| tr\|B0SL80\|B0SL80_LEPBP | | 3 | 4 | 7 | 2 | Putative transcriptional regulator, AraC family putative membrane protein OS=Leptospira biflexa serovar Patoc (strain Patoc 1 / ATCC 23582 / Paris) OX=456481 GN=LEPBI_I0755 PE=4 SV=1 |
| tr\|B0STN9\|B0STN9_LEPBP | | 3 | 4 | 7 | 2 | STAS domain-containing protein OS=Leptospira biflexa serovar Patoc (strain Patoc 1 / ATCC 23582 / Paris) OX=456481 GN=LEPBI_II0035 PE=4 SV=1 |
| tr\|B0SU67\|B0SU67_LEPBP | 3 | | 4 | 7 | 2 | Uncharacterized protein OS=Leptospira biflexa serovar Patoc (strain Patoc 1 / ATCC 23582 / Paris) OX=456481 GN=LEPBI_II0218 PE=4 SV=1 |
| tr\|B0SKI1\|B0SKI1_LEPBP | | 5 | 2 | 7 | 2 | Uncharacterized protein OS=Leptospira biflexa serovar Patoc (strain Patoc 1 / ATCC 23582 / Paris) OX=456481 GN=LEPBI_I0601 PE=4 SV=1 |
| tr\|B0SJS6\|B0SJS6_LEPBP | 2 | | 5 | 7 | 2 | APH domain-containing protein OS=Leptospira biflexa serovar Patoc (strain Patoc 1 / ATCC 23582 / Paris) OX=456481 GN=LEPBI_I0076 PE=4 SV=1 |
| tr\|B0SK95\|B0SK95_LEPBP | 5 | | 2 | 7 | 2 | Chemotaxis MotB protein OS=Leptospira biflexa serovar Patoc (strain Patoc 1 / ATCC 23582 / Paris) OX=456481 GN=motB PE=3 SV=1 |
| tr\|B0SMY3\|B0SMY3_LEPBP | 2 | 5 | | 7 | 2 | Uncharacterized protein OS=Leptospira biflexa serovar Patoc (strain Patoc 1 / ATCC 23582 / Paris) OX=456481 GN=LEPBI_I1047 PE=4 SV=1 |
| tr\|B0SJD4\|B0SJD4_LEPBP | 3 | 4 | | 7 | 2 | DNA helicase OS=Leptospira biflexa serovar Patoc (strain Patoc 1 / ATCC 23582 / Paris) OX=456481 GN=recQ1 PE=3 SV=1 |
| tr\|B0SRK6\|B0SRK6_LEPBP | 3 | 4 | | 7 | 2 | Putative permease, MFS superfamily putative sulfate transporter putative membrane protein putative signal peptide OS=Leptospira biflexa serovar Patoc (strain Patoc 1 / ATCC 23582 / Paris) OX=456481 GN=LEPBI_I1695 PE=3 SV=1 |
| tr\|B0SS39\|B0SS39_LEPBP | 1 | 6 | | 7 | 2 | Transcription-repair-coupling factor OS=Leptospira biflexa serovar Patoc (strain Patoc 1 / ATCC 23582 / Paris) OX=456481 GN=mfd PE=3 SV=1 |
| tr\|B0SP46\|B0SP46_LEPBP | 4 | 3 | | 7 | 2 | Putative methyl-accepting chemotaxis protein putative membrane protein OS=Leptospira biflexa serovar Patoc (strain Patoc 1 / ATCC 23582 / Paris) OX=456481 GN=LEPBI_I1258 PE=4 SV=1 |
| tr\|B0SNV3\|B0SNV3_LEPBP | 5 | 2 | | 7 | 2 | tRNA (guanine-N(7)-)-methyltransferase OS=Leptospira biflexa serovar Patoc (strain Patoc 1 / ATCC 23582 / Paris) OX=456481 GN=trmB PE=3 SV=1 |
| tr\|B0SJ89\|B0SJ89_LEPBP | 3 | 4 | | 7 | 2 | Putative dehalogenase-like hydrolase OS=Leptospira biflexa serovar Patoc (strain Patoc 1 / ATCC 23582 / Paris) OX=456481 GN=LEPBI_I0583 PE=4 SV=1 |
| tr\|B0SPV4\|B0SPV4_LEPBP | 3 | 4 | | 7 | 2 | SsrA-binding protein OS=Leptospira biflexa serovar Patoc (strain Patoc 1 / ATCC 23582 / Paris) OX=456481 GN=smpB PE=3 SV=1 |
| tr\|B0SP84\|B0SP84_LEPBP | 2 | 5 | | 7 | 2 | NADH-quinone oxidoreductase subunit C OS=Leptospira biflexa serovar Patoc (strain Patoc 1 / ATCC 23582 / Paris) OX=456481 GN=nuoC PE=3 SV=1 |
| tr\|B0SJF9\|B0SJF9_LEPBP | 4 | 3 | | 7 | 2 | Putative alginate o-acetyltransferase putative membrane protein OS=Leptospira biflexa serovar Patoc (strain Patoc 1 / ATCC 23582 / Paris) OX=456481 GN=LEPBI_I0541 PE=3 SV=1 |
| tr\|B0SNB8\|B0SNB8_LEPBP | 3 | 4 | | 7 | 2 | Uncharacterized protein OS=Leptospira biflexa serovar Patoc (strain Patoc 1 / ATCC 23582 / Paris) OX=456481 GN=LEPBI_I2809 PE=4 SV=1 |
| tr\|B0SS67\|B0SS67_LEPBP | 3 | 4 | | 7 | 2 | ATP-dependent Clp protease ATP-binding subunit ClpA OS=Leptospira biflexa serovar Patoc (strain Patoc 1 / ATCC 23582 / Paris) OX=456481 GN=clpA PE=3 SV=1 |
| tr\|B0SML3\|B0SML3_LEPBP | 5 | 2 | | 7 | 2 | Uncharacterized protein OS=Leptospira biflexa serovar Patoc (strain Patoc 1 / ATCC 23582 / Paris) OX=456481 GN=LEPBI_I2658 PE=4 SV=1 |
| tr\|B0SQW1\|B0SQW1_LEPBP | | 2 | 4 | 6 | 2 | Putative zinc-binding alcohol dehydrogenase OS=Leptospira biflexa serovar Patoc (strain Patoc 1 / ATCC 23582 / Paris) OX=456481 GN=LEPBI_I3293 PE=3 SV=1 |
| tr\|B0SMI5\|B0SMI5_LEPBP | | 3 | 3 | 6 | 2 | Uncharacterized protein OS=Leptospira biflexa serovar Patoc (strain Patoc 1 / ATCC 23582 / Paris) OX=456481 GN=LEPBI_I1005 PE=4 SV=1 |
| tr\|B0SR60\|B0SR60_LEPBP | 1 | | 5 | 6 | 2 | Putative nitrogen assimilation regulatory protein NtrX OS=Leptospira biflexa serovar Patoc (strain Patoc 1 / ATCC 23582 / Paris) OX=456481 GN=LEPBI_I1654 PE=4 SV=1 |
| tr\|B0SNR1\|B0SNR1_LEPBP | 5 | | 1 | 6 | 2 | Putative transcriptional regulator, LysR family OS=Leptospira biflexa serovar Patoc (strain Patoc 1 / ATCC 23582 / Paris) OX=456481 GN=LEPBI_I1228 PE=3 SV=1 |
| tr\|B0SR04\|B0SR04_LEPBP | 2 | | 4 | 6 | 2 | Putative ATPase, AAA family OS=Leptospira biflexa serovar Patoc (strain Patoc 1 / ATCC 23582 / Paris) OX=456481 GN=LEPBI_I1598 PE=3 SV=1 |
| sp\|B0SSW2\|RS6_LEPBP | | 3 | 3 | 6 | 2 | 30S ribosomal protein S6 OS=Leptospira biflexa serovar Patoc (strain Patoc 1 / ATCC 23582 / Paris) OX=456481 GN=rpsF PE=3 SV=1 |
| tr\|B0SLF8\|B0SLF8_LEPBP | | 1 | 5 | 6 | 2 | Dipeptidase OS=Leptospira biflexa serovar Patoc (strain Patoc 1 / ATCC 23582 / Paris) OX=456481 GN=LEPBI_I2455 PE=3 SV=1 |
| sp\|B0SMK7\|DAPAT_LEPBP | | 3 | 3 | 6 | 2 | LL-diaminopimelate aminotransferase OS=Leptospira biflexa serovar Patoc (strain Patoc 1 / ATCC 23582 / Paris) OX=456481 GN=dapL PE=3 SV=1 |
| tr\|B0SP08\|B0SP08_LEPBP | | 1 | 5 | 6 | 2 | NorD protein OS=Leptospira biflexa serovar Patoc (strain Patoc 1 / ATCC 23582 / Paris) OX=456481 GN=norD PE=4 SV=1 |
| tr\|B0SP18\|B0SP18_LEPBP | 3 | | 3 | 6 | 2 | Uncharacterized protein OS=Leptospira biflexa serovar Patoc (strain Patoc 1 / ATCC 23582 / Paris) OX=456481 GN=LEPBI_I2962 PE=4 SV=1 |
| tr\|B0SLB8\|B0SLB8_LEPBP | | 3 | 3 | 6 | 2 | Glutathione peroxidase OS=Leptospira biflexa serovar Patoc (strain Patoc 1 / ATCC 23582 / Paris) OX=456481 GN=gpo PE=3 SV=1 |
| sp\|B0SRT6\|RL27_LEPBP | 1 | 5 | | 6 | 2 | 50S ribosomal protein L27 OS=Leptospira biflexa serovar Patoc (strain Patoc 1 / ATCC 23582 / Paris) OX=456481 GN=rpmA PE=3 SV=1 |
| tr\|B0SQS2\|B0SQS2_LEPBP | 3 | 3 | | 6 | 2 | Putative sugar transport protein OS=Leptospira biflexa serovar Patoc (strain Patoc 1 / ATCC 23582 / Paris) OX=456481 GN=LEPBI_I3254 PE=4 SV=1 |
| tr\|B0STI8\|B0STI8_LEPBP | 5 | 1 | | 6 | 2 | Peptidoglycan synthetase FtsI (Penicillin-binding protein 3 PBP-3) OS=Leptospira biflexa serovar Patoc (strain Patoc 1 / ATCC 23582 / Paris) OX=456481 GN=ftsI PE=4 SV=1 |
| tr\|B0SU93\|B0SU93_LEPBP | 3 | 3 | | 6 | 2 | DUF1801 domain-containing protein OS=Leptospira biflexa serovar Patoc (strain Patoc 1 / ATCC 23582 / Paris) OX=456481 GN=LEPBI_II0244 PE=4 SV=1 |
| tr\|B0SU54\|B0SU54_LEPBP | 3 | 3 | | 6 | 2 | Uncharacterized protein OS=Leptospira biflexa serovar Patoc (strain Patoc 1 / ATCC 23582 / Paris) OX=456481 GN=LEPBI_II0205 PE=4 SV=1 |
| tr\|B0SKR9\|B0SKR9_LEPBP | 2 | 4 | | 6 | 2 | Uncharacterized protein OS=Leptospira biflexa serovar Patoc (strain Patoc 1 / ATCC 23582 / Paris) OX=456481 GN=LEPBI_I0694 PE=4 SV=1 |
| sp\|B0SSG0\|RL30_LEPBP | 2 | 4 | | 6 | 2 | 50S ribosomal protein L30 OS=Leptospira biflexa serovar Patoc (strain Patoc 1 / ATCC 23582 / Paris) OX=456481 GN=rpmD PE=3 SV=1 |
| tr\|B0STY5\|B0STY5_LEPBP | 2 | 4 | | 6 | 2 | Putative helicase, ATP-dependent OS=Leptospira biflexa serovar Patoc (strain Patoc 1 / ATCC 23582 / Paris) OX=456481 GN=LEPBI_II0133 PE=4 SV=1 |
| tr\|B0SQI6\|B0SQI6_LEPBP | 2 | 4 | | 6 | 2 | Putative ribonuclease BN putative membrane protein OS=Leptospira biflexa serovar Patoc (strain Patoc 1 / ATCC 23582 / Paris) OX=456481 GN=LEPBI_I1537 PE=4 SV=1 |
| tr\|B0SR61\|B0SR61_LEPBP | 4 | 2 | | 6 | 2 | Uncharacterized protein OS=Leptospira biflexa serovar Patoc (strain Patoc 1 / ATCC 23582 / Paris) OX=456481 GN=LEPBI_I1655 PE=4 SV=1 |
| tr\|B0ST00\|B0ST00_LEPBP | 2 | 4 | | 6 | 2 | Probable UDP-N-acetylglucosamine--peptide N-acetylglucosaminyltransferase SPINDLY OS=Leptospira biflexa serovar Patoc (strain Patoc 1 / ATCC 23582 / Paris) OX=456481 GN=LEPBI_I2138 PE=4 SV=1 |
| tr\|B0SR52\|B0SR52_LEPBP | 4 | 2 | | 6 | 2 | 2-dehydro-3-deoxyphosphooctonate aldolase OS=Leptospira biflexa serovar Patoc (strain Patoc 1 / ATCC 23582 / Paris) OX=456481 GN=kdsA PE=3 SV=1 |
| tr\|B0SN54\|B0SN54_LEPBP | 3 | 3 | | 6 | 2 | Putative uroporphyrin-III C/tetrapyrrole (Corrin/Porphyrin) methyltransferase OS=Leptospira biflexa serovar Patoc (strain Patoc 1 / ATCC 23582 / Paris) OX=456481 GN=LEPBI_I1122 PE=4 SV=1 |
| tr\|B0SMI4\|B0SMI4_LEPBP | 3 | 3 | | 6 | 2 | Uncharacterized protein OS=Leptospira biflexa serovar Patoc (strain Patoc 1 / ATCC 23582 / Paris) OX=456481 GN=LEPBI_I1004 PE=4 SV=1 |
| sp\|B0SP83\|NUOB_LEPBP | 2 | 4 | | 6 | 2 | NADH-quinone oxidoreductase subunit B OS=Leptospira biflexa serovar Patoc (strain Patoc 1 / ATCC 23582 / Paris) OX=456481 GN=nuoB PE=3 SV=1 |
| tr\|B0SUD7\|B0SUD7_LEPBP | 2 | 4 | | 6 | 2 | Putative transcriptional regulator OS=Leptospira biflexa serovar Patoc (strain Patoc 1 / ATCC 23582 / Paris) OX=456481 GN=LEPBI_p0007 PE=4 SV=1 |
| tr\|B0SN39\|B0SN39_LEPBP | 2 | 4 | | 6 | 2 | Uncharacterized protein OS=Leptospira biflexa serovar Patoc (strain Patoc 1 / ATCC 23582 / Paris) OX=456481 GN=LEPBI_I1107 PE=4 SV=1 |
| tr\|B0SK08\|B0SK08_LEPBP | 2 | 4 | | 6 | 2 | Putative methyl-accepting chemotaxis protein (MCP) putative membrane protein OS=Leptospira biflexa serovar Patoc (strain Patoc 1 / ATCC 23582 / Paris) OX=456481 GN=LEPBI_I0422 PE=4 SV=1 |
| tr\|B0STG6\|B0STG6_LEPBP | 4 | 2 | | 6 | 2 | Heme chaperone HemW OS=Leptospira biflexa serovar Patoc (strain Patoc 1 / ATCC 23582 / Paris) OX=456481 GN=LEPBI_I2311 PE=3 SV=1 |
| tr\|B0SPV7\|B0SPV7_LEPBP | 3 | 3 | | 6 | 2 | Uncharacterized protein OS=Leptospira biflexa serovar Patoc (strain Patoc 1 / ATCC 23582 / Paris) OX=456481 GN=LEPBI_I1414 PE=4 SV=1 |

| | | | | | | |
|---|---|---|---|---|---|---|
| tr\|B0SKP5\|B0SKP5_LEPBP | 3 | 3 | | 6 | 2 | Heme oxygenase OS=Leptospira biflexa serovar Patoc (strain Patoc 1 / ATCC 23582 / Paris) OX=456481 GN=hmuO PE=4 SV=1 |
| tr\|B0SRE0\|B0SRE0_LEPBP | 2 | 4 | | 6 | 2 | Putative arsenate reductase, arsC family (Arsenical pump modifier) OS=Leptospira biflexa serovar Patoc (strain Patoc 1 / ATCC 23582 / Paris) OX=456481 GN=LEPBI_I3368 PE=3 SV=1 |
| tr\|B0SJI6\|B0SJI6_LEPBP | 4 | 2 | | 6 | 2 | Sensor protein CreC OS=Leptospira biflexa serovar Patoc (strain Patoc 1 / ATCC 23582 / Paris) OX=456481 GN=creC PE=4 SV=1 |

**Table S3.** Label-free comparative proteomics of *L. biflexa* filamei

**Label-free comparative proteomics of *L. biflexa* filaments from *wt* and *j***

Proteins identified in the cryoEM structure of *L. biflexa* filaments are marked in red

Core flagellins (FlaBs) and paralogues of *Leptospira* flagellar filament proteins are sho

### Venn Diagram

**Proteins exclusively identified in *L. biflexa wt***

| | Number of Spectra | | | | |
|---|---|---|---|---|---|
| **Locus** | **WT_R1** | **WT_R2** | **WT_R3** | **Total Signal** | **Replicate Count** |
| tr\|B0SMK8\|B0SMK8_LEPBP | 288 | 265 | 293 | 846 | 3 |
| tr\|B0SKN8\|B0SKN8_LEPBP | 207 | 194 | 176 | 577 | 3 |
| **tr\|B0SNJ6\|B0SNJ6_LEPBP** | 26 | 12 | 43 | 81 | 3 |
| tr\|B0STN7\|B0STN7_LEPBP | 62 | 7 | 9 | 78 | 3 |
| tr\|B0SSE3\|B0SSE3_LEPBP | 5 | 8 | 7 | 20 | 3 |
| tr\|B0SMT6\|B0SMT6_LEPBP | 4 | 2 | 4 | 10 | 3 |
| tr\|B0SL54\|B0SL54_LEPBP | 3 | 4 | 3 | 10 | 3 |
| sp\|B0SNJ9\|HEMH_LEPBP | 3 | 3 | 3 | 9 | 3 |
| tr\|B0SQ23\|B0SQ23_LEPBP | 3 | 3 | 3 | 9 | 3 |
| tr\|B0SS20\|B0SS20_LEPBP | 3 | 2 | 3 | 8 | 3 |
| tr\|B0SMH9\|B0SMH9_LEPBP | 2 | 4 | 2 | 8 | 3 |

| | | | | | |
|---|---|---|---|---|---|
| tr\|B0SR15\|B0SR15_LEPBP | 2 | 3 | 3 | 8 | 3 |
| tr\|B0SSZ1\|B0SSZ1_LEPBP | 3 | 2 | 3 | 8 | 3 |
| sp\|B0SMW9\|RL31_LEPBP | 1 | 3 | 3 | 7 | 3 |
| tr\|B0SMW5\|B0SMW5_LEPBP | 3 | 2 | 2 | 7 | 3 |
| tr\|B0SLU0\|B0SLU0_LEPBP | 2 | 3 | 2 | 7 | 3 |
| tr\|B0SKY2\|B0SKY2_LEPBP | 1 | 4 | 2 | 7 | 3 |
| tr\|B0SNN0\|B0SNN0_LEPBP | 2 | 2 | 2 | 6 | 3 |
| tr\|B0SPN3\|B0SPN3_LEPBP | 6 | 5 | | 11 | 2 |
| tr\|B0SM39\|B0SM39_LEPBP | 4 | | 3 | 7 | 2 |
| tr\|B0SSX5\|B0SSX5_LEPBP | 5 | 2 | | 7 | 2 |
| tr\|B0SQF3\|B0SQF3_LEPBP | | 4 | 2 | 6 | 2 |
| tr\|B0SU70\|B0SU70_LEPBP | 2 | 3 | | 5 | 2 |
| tr\|B0SML0\|B0SML0_LEPBP | | 2 | 3 | 5 | 2 |

**Proteins exclusively identified in *L. biflexa fcpA-***

| | Number of Spectra |
|---|---|

| Locus | FcpA_R1 | FcpA_R2 | FcpA_R3 | Total Signal | Replicate Count |
|---|---|---|---|---|---|
| tr|B0SRK9|B0SRK9_LEPBP | 136 | 14 | 13 | 163 | 3 |
| tr|B0SRV8|B0SRV8_LEPBP | 16 | 36 | 16 | 68 | 3 |
| tr|B0SJ68|B0SJ68_LEPBP | 18 | 28 | 11 | 57 | 3 |
| tr|B0SRC6|B0SRC6_LEPBP | 24 | 21 | 11 | 56 | 3 |
| tr|B0SKI7|B0SKI7_LEPBP | 13 | 17 | 14 | 44 | 3 |
| tr|B0SPX3|B0SPX3_LEPBP | 37 | 3 | 2 | 42 | 3 |
| tr|B0SRU5|B0SRU5_LEPBP | 21 | 18 | 3 | 42 | 3 |
| tr|B0SNG9|B0SNG9_LEPBP | 9 | 20 | 11 | 40 | 3 |
| tr|B0SSV5|B0SSV5_LEPBP | 5 | 10 | 21 | 36 | 3 |
| tr|B0SPE8|B0SPE8_LEPBP | 19 | 4 | 10 | 33 | 3 |
| tr|B0SPN5|B0SPN5_LEPBP | 15 | 8 | 10 | 33 | 3 |
| sp|B0SNZ7|GATB_LEPBP | 16 | 12 | 4 | 32 | 3 |
| tr|B0SJQ4|B0SJQ4_LEPBP | 4 | 21 | 7 | 32 | 3 |
| tr|B0SLH6|B0SLH6_LEPBP | 6 | 17 | 9 | 32 | 3 |

| | | | | | |
|---|---|---|---|---|---|
| tr\|B0SKT7\|B0SKT7_LEPBP | 19 | 4 | 8 | 31 | 3 |
| tr\|B0SU99\|B0SU99_LEPBP | 16 | 7 | 7 | 30 | 3 |
| tr\|B0SM55\|B0SM55_LEPBP | 7 | 12 | 10 | 29 | 3 |
| tr\|B0SQ78\|B0SQ78_LEPBP | 12 | 6 | 11 | 29 | 3 |
| tr\|B0SQY3\|B0SQY3_LEPBP | 4 | 18 | 6 | 28 | 3 |
| tr\|B0SMZ3\|B0SMZ3_LEPBP | 20 | 5 | 3 | 28 | 3 |
| tr\|B0SRX3\|B0SRX3_LEPBP | 7 | 8 | 13 | 28 | 3 |
| tr\|B0SL64\|B0SL64_LEPBP | 18 | 3 | 6 | 27 | 3 |
| sp\|B0SPF0\|PCKA_LEPBP | 5 | 11 | 9 | 25 | 3 |
| tr\|B0SUC7\|B0SUC7_LEPBP | 16 | 5 | 4 | 25 | 3 |
| tr\|B0SU07\|B0SU07_LEPBP | 5 | 17 | 3 | 25 | 3 |
| tr\|B0SRJ5\|B0SRJ5_LEPBP | 13 | 7 | 5 | 25 | 3 |
| tr\|B0SUF4\|B0SUF4_LEPBP | 3 | 6 | 15 | 24 | 3 |
| tr\|B0SJH9\|B0SJH9_LEPBP | 15 | 2 | 6 | 23 | 3 |
| tr\|B0SR38\|B0SR38_LEPBP | 15 | 3 | 5 | 23 | 3 |

| | | | | | |
|---|---|---|---|---|---|
| sp\|B0SLC5\|ATPD_LEPBP | 12 | 7 | 3 | 22 | 3 |
| tr\|B0SMN6\|B0SMN6_LEPBP | 8 | 6 | 7 | 21 | 3 |
| tr\|B0SLT1\|B0SLT1_LEPBP | 5 | 9 | 7 | 21 | 3 |
| tr\|B0SQY8\|B0SQY8_LEPBP | 2 | 13 | 5 | 20 | 3 |
| tr\|B0SKJ8\|B0SKJ8_LEPBP | 7 | 6 | 7 | 20 | 3 |
| tr\|B0ST04\|B0ST04_LEPBP | 13 | 2 | 5 | 20 | 3 |
| sp\|B0SSH5\|RL23_LEPBP | 14 | 3 | 2 | 19 | 3 |
| tr\|B0SRK5\|B0SRK5_LEPBP | 4 | 7 | 6 | 17 | 3 |
| sp\|B0SQH7\|RS15_LEPBP | 9 | 5 | 3 | 17 | 3 |
| tr\|B0SSJ3\|B0SSJ3_LEPBP | 10 | 4 | 2 | 16 | 3 |
| tr\|B0STR7\|B0STR7_LEPBP | 6 | 4 | 6 | 16 | 3 |
| tr\|B0SRH6\|B0SRH6_LEPBP | 4 | 4 | 7 | 15 | 3 |
| tr\|B0SKS1\|B0SKS1_LEPBP | 6 | 4 | 5 | 15 | 3 |
| tr\|B0SRR4\|B0SRR4_LEPBP | 9 | 3 | 2 | 14 | 3 |
| tr\|B0SP92\|B0SP92_LEPBP | 4 | 5 | 4 | 13 | 3 |
| tr\|B0SSM9\|B0SSM9_LEPBP | 8 | 2 | 3 | 13 | 3 |

| | | | | | |
|---|---|---|---|---|---|
| tr\|B0SJA7\|B0SJA7_LEPBP | 4 | 5 | 3 | 12 | 3 |
| tr\|B0SP91\|B0SP91_LEPBP | 3 | 4 | 5 | 12 | 3 |
| tr\|B0SMW2\|B0SMW2_LEPBP | 4 | 3 | 5 | 12 | 3 |
| sp\|B0SMJ9\|RL28_LEPBP | 6 | 3 | 3 | 12 | 3 |
| tr\|B0STI1\|B0STI1_LEPBP | 4 | 3 | 4 | 11 | 3 |
| tr\|B0SPQ6\|B0SPQ6_LEPBP | 3 | 3 | 5 | 11 | 3 |
| tr\|B0SLG5\|B0SLG5_LEPBP | 4 | 6 | 1 | 11 | 3 |
| tr\|B0SP25\|B0SP25_LEPBP | 6 | 2 | 3 | 11 | 3 |
| tr\|B0SJE9\|B0SJE9_LEPBP | 1 | 4 | 5 | 10 | 3 |
| tr\|B0SRJ6\|B0SRJ6_LEPBP | 4 | 3 | 3 | 10 | 3 |
| sp\|B0SSG0\|RL30_LEPBP | 3 | 4 | 3 | 10 | 3 |
| tr\|B0SIX7\|B0SIX7_LEPBP | 3 | 4 | 3 | 10 | 3 |
| tr\|B0STN1\|B0STN1_LEPBP | 5 | 3 | 2 | 10 | 3 |
| tr\|B0SRC4\|B0SRC4_LEPBP | 2 | 3 | 4 | 9 | 3 |
| tr\|B0SK36\|B0SK36_LEPBP | 3 | 3 | 3 | 9 | 3 |

| | | | | | |
|---|---|---|---|---|---|
| tr\|B0SSB9\|B0SSB9_LEPBP | 2 | 2 | 4 | 8 | 3 |
| tr\|B0SSE0\|B0SSE0_LEPBP | 3 | 2 | 3 | 8 | 3 |
| tr\|B0SQ86\|B0SQ86_LEPBP | 2 | 3 | 2 | 7 | 3 |
| tr\|B0SQ65\|B0SQ65_LEPBP | 1 | 1 | 5 | 7 | 3 |
| tr\|B0SUD7\|B0SUD7_LEPBP | 2 | 2 | 3 | 7 | 3 |
| tr\|B0SMR3\|B0SMR3_LEPBP | 76 | | 37 | 113 | 2 |
| tr\|B0SJ86\|B0SJ86_LEPBP | 41 | | 3 | 44 | 2 |
| tr\|B0SQY4\|B0SQY4_LEPBP | | 21 | 8 | 29 | 2 |
| sp\|B0SRL5\|ENO_LEPBP | 24 | 4 | | 28 | 2 |
| tr\|B0SRJ8\|B0SRJ8_LEPBP | 22 | | 4 | 26 | 2 |
| sp\|B0SKZ6\|RL35_LEPBP | 17 | | 7 | 24 | 2 |
| sp\|B0SNG1\|SECA_LEPBP | 16 | | 7 | 23 | 2 |
| tr\|B0SSI9\|B0SSI9_LEPBP | 15 | | 6 | 21 | 2 |
| tr\|B0SJN5\|B0SJN5_LEPBP | 14 | | 6 | 20 | 2 |
| tr\|B0SUB1\|B0SUB1_LEPBP | 7 | 13 | | 20 | 2 |
| tr\|B0SK85\|B0SK85_LEPBP | 11 | | 8 | 19 | 2 |

| | | | | | |
|---|---|---|---|---|---|
| tr\|B0STI5\|B0STI5_LEPBP | 11 | | 8 | 19 | 2 |
| tr\|B0SJ22\|B0SJ22_LEPBP | 6 | 13 | | 19 | 2 |
| tr\|B0SRE1\|B0SRE1_LEPBP | 11 | | 7 | 18 | 2 |
| tr\|B0SL95\|B0SL95_LEPBP | 15 | 2 | | 17 | 2 |
| tr\|B0SJR0\|B0SJR0_LEPBP | | 6 | 11 | 17 | 2 |
| tr\|B0SPE5\|B0SPE5_LEPBP | 7 | | 9 | 16 | 2 |
| tr\|B0SQE3\|B0SQE3_LEPBP | 6 | | 10 | 16 | 2 |
| tr\|B0SQ52\|B0SQ52_LEPBP | 7 | | 9 | 16 | 2 |
| tr\|B0SSU0\|B0SSU0_LEPBP | 11 | | 4 | 15 | 2 |
| sp\|B0SRH5\|NUOI_LEPBP | 11 | | 4 | 15 | 2 |
| tr\|B0SUH3\|B0SUH3_LEPBP | 10 | 5 | | 15 | 2 |
| sp\|B0SMI6\|GLYA_LEPBP | 3 | 12 | | 15 | 2 |
| sp\|B0STE5\|SYI_LEPBP | 2 | | 12 | 14 | 2 |
| tr\|B0SQV1\|B0SQV1_LEPBP | 9 | | 5 | 14 | 2 |
| tr\|B0STP2\|B0STP2_LEPBP | 11 | 3 | | 14 | 2 |

| | | | | | |
|---|---|---|---|---|---|
| tr\|B0SLQ4\|B0SLQ4_LEPBP | | 9 | 5 | 14 | 2 |
| tr\|B0STN9\|B0STN9_LEPBP | 6 | | 7 | 13 | 2 |
| sp\|B0SPT9\|RIMO_LEPBP | 8 | | 5 | 13 | 2 |
| tr\|B0SLF2\|B0SLF2_LEPBP | 2 | | 10 | 12 | 2 |
| tr\|B0SQV7\|B0SQV7_LEPBP | 5 | | 7 | 12 | 2 |
| tr\|B0SJF3\|B0SJF3_LEPBP | 5 | | 7 | 12 | 2 |
| tr\|B0SKM6\|B0SKM6_LEPBP | 9 | | 2 | 11 | 2 |
| tr\|B0SMS7\|B0SMS7_LEPBP | 6 | | 5 | 11 | 2 |
| tr\|B0SLR5\|B0SLR5_LEPBP | 9 | 2 | | 11 | 2 |
| tr\|B0STR5\|B0STR5_LEPBP | 8 | | 3 | 11 | 2 |
| tr\|B0SLS6\|B0SLS6_LEPBP | 8 | 3 | | 11 | 2 |
| tr\|B0SNM6\|B0SNM6_LEPBP | 6 | 4 | | 10 | 2 |
| tr\|B0SJ72\|B0SJ72_LEPBP | 7 | | 3 | 10 | 2 |
| tr\|B0SSD6\|B0SSD6_LEPBP | 6 | | 4 | 10 | 2 |
| tr\|B0STZ2\|B0STZ2_LEPBP | 8 | | 2 | 10 | 2 |
| tr\|B0SME8\|B0SME8_LEPBP | 7 | | 3 | 10 | 2 |

| | | | | | |
|---|---|---|---|---|---|
| tr\|B0SSB7\|B0SSB7_LEPBP | 6 | | 4 | 10 | 2 |
| tr\|B0SQQ3\|B0SQQ3_LEPBP | | 4 | 6 | 10 | 2 |
| tr\|B0SQ04\|B0SQ04_LEPBP | 6 | | 3 | 9 | 2 |
| tr\|B0SM54\|B0SM54_LEPBP | 3 | | 6 | 9 | 2 |
| tr\|B0SUB7\|B0SUB7_LEPBP | 4 | | 5 | 9 | 2 |
| tr\|B0SRI7\|B0SRI7_LEPBP | 2 | | 7 | 9 | 2 |
| tr\|B0SKF7\|B0SKF7_LEPBP | 5 | | 4 | 9 | 2 |
| tr\|B0STJ1\|B0STJ1_LEPBP | 3 | | 5 | 8 | 2 |
| sp\|B0SSW2\|RS6_LEPBP | 6 | | 2 | 8 | 2 |
| tr\|B0SKJ9\|B0SKJ9_LEPBP | 4 | | 4 | 8 | 2 |
| tr\|B0SPZ1\|B0SPZ1_LEPBP | 6 | | 2 | 8 | 2 |
| tr\|B0SLQ6\|B0SLQ6_LEPBP | 6 | | 2 | 8 | 2 |
| tr\|B0SRW7\|B0SRW7_LEPBP | 5 | | 3 | 8 | 2 |
| tr\|B0SUD2\|B0SUD2_LEPBP | 6 | 2 | | 8 | 2 |
| tr\|B0SRN0\|B0SRN0_LEPBP | 2 | | 6 | 8 | 2 |

| | | | | | |
|---|---|---|---|---|---|
| tr\|B0SSE7\|B0SSE7_LEPBP | 4 | | 4 | 8 | 2 |
| tr\|B0SJB3\|B0SJB3_LEPBP | 4 | | 4 | 8 | 2 |
| tr\|B0SKI6\|B0SKI6_LEPBP | 4 | | 4 | 8 | 2 |
| tr\|B0SPL1\|B0SPL1_LEPBP | 4 | | 4 | 8 | 2 |
| tr\|B0SUC3\|B0SUC3_LEPBP | 3 | | 5 | 8 | 2 |
| tr\|B0SNH8\|B0SNH8_LEPBP | 4 | | 4 | 8 | 2 |
| tr\|B0SNE2\|B0SNE2_LEPBP | 5 | | 2 | 7 | 2 |
| tr\|B0SS24\|B0SS24_LEPBP | 2 | | 5 | 7 | 2 |
| tr\|B0SKJ4\|B0SKJ4_LEPBP | 3 | | 4 | 7 | 2 |
| tr\|B0SQT8\|B0SQT8_LEPBP | 3 | | 4 | 7 | 2 |
| tr\|B0SK83\|B0SK83_LEPBP | 3 | | 4 | 7 | 2 |
| tr\|B0SKL3\|B0SKL3_LEPBP | 1 | | 6 | 7 | 2 |
| tr\|B0SP11\|B0SP11_LEPBP | 6 | | 1 | 7 | 2 |
| tr\|B0SP67\|B0SP67_LEPBP | 5 | 2 | | 7 | 2 |
| tr\|B0SQ32\|B0SQ32_LEPBP | 4 | | 3 | 7 | 2 |
| sp\|B0SM63\|RRF_LEPBP | 2 | | 5 | 7 | 2 |

| | | | | | |
|---|---|---|---|---|---|
| tr\|B0STU5\|B0STU5_LEPBP | 3 | | 4 | 7 | 2 |
| tr\|B0SUA4\|B0SUA4_LEPBP | 3 | | 4 | 7 | 2 |
| tr\|B0SQS1\|B0SQS1_LEPBP | 5 | | 2 | 7 | 2 |
| tr\|B0SPD2\|B0SPD2_LEPBP | 5 | | 2 | 7 | 2 |
| tr\|B0STK4\|B0STK4_LEPBP | 4 | | 3 | 7 | 2 |
| tr\|B0STJ9\|B0STJ9_LEPBP | 3 | | 4 | 7 | 2 |
| tr\|B0SUA6\|B0SUA6_LEPBP | | 2 | 5 | 7 | 2 |
| tr\|B0SK08\|B0SK08_LEPBP | | 2 | 5 | 7 | 2 |
| tr\|B0STY1\|B0STY1_LEPBP | | 3 | 4 | 7 | 2 |
| tr\|B0SL56\|B0SL56_LEPBP | | 5 | 2 | 7 | 2 |

nts from *wt vs* mutant strains.

***fcpA-* mutant strains. Analysis by Spectral Counts (SC)**

wn in **bold** (see Supplementary Table 4)

| |
|---|
| **Description** |
| Uncharacterized protein OS=Leptospira biflexa serovar Patoc (strain Patoc 1 / ATCC 23582 / Paris) OX=456481 GN=LEPBI_I1029 PE=4 SV=1 |
| Uncharacterized protein OS=Leptospira biflexa serovar Patoc (strain Patoc 1 / ATCC 23582 / Paris) OX=456481 GN=LEPBI_I0662 PE=4 SV=1 |
| Uncharacterized protein OS=Leptospira biflexa serovar Patoc (strain Patoc 1 / ATCC 23582 / Paris) OX=456481 GN=LEPBI_I1161 PE=4 SV=1 |
| Uncharacterized protein OS=Leptospira biflexa serovar Patoc (strain Patoc 1 / ATCC 23582 / Paris) OX=456481 GN=LEPBI_II0033 PE=4 SV=1 |
| Putative TPR-repeat-containing protein putative signal peptide OS=Leptospira biflexa serovar Patoc (strain Patoc 1 / ATCC 23582 / Paris) OX=456481 GN=LEPBI_I1930 PE=4 SV=1 |
| Uncharacterized protein OS=Leptospira biflexa serovar Patoc (strain Patoc 1 / ATCC 23582 / Paris) OX=456481 GN=LEPBI_I2732 PE=4 SV=1 |
| Uncharacterized protein OS=Leptospira biflexa serovar Patoc (strain Patoc 1 / ATCC 23582 / Paris) OX=456481 GN=LEPBI_I0729 PE=4 SV=1 |
| Ferrochelatase OS=Leptospira biflexa serovar Patoc (strain Patoc 1 / ATCC 23582 / Paris) OX=456481 GN=hemH PE=3 SV=1 |
| Uncharacterized protein OS=Leptospira biflexa serovar Patoc (strain Patoc 1 / ATCC 23582 / Paris) OX=456481 GN=LEPBI_I3109 PE=4 SV=1 |
| Uncharacterized protein OS=Leptospira biflexa serovar Patoc (strain Patoc 1 / ATCC 23582 / Paris) OX=456481 GN=LEPBI_I1804 PE=4 SV=1 |
| Putative lipid A core-O-antigen ligase putative membrane protein OS=Leptospira biflexa serovar Patoc (strain Patoc 1 / ATCC 23582 / Paris) OX=456481 GN=LEPBI_I0999 PE=4 SV=1 |

| |
|---|
| Uncharacterized protein OS=Leptospira biflexa serovar Patoc (strain Patoc 1 / ATCC 23582 / Paris) OX=456481 GN=LEPBI_I1609 PE=4 SV=1 |
| Uncharacterized protein OS=Leptospira biflexa serovar Patoc (strain Patoc 1 / ATCC 23582 / Paris) OX=456481 GN=LEPBI_I2129 PE=4 SV=1 |
| 50S ribosomal protein L31 OS=Leptospira biflexa serovar Patoc (strain Patoc 1 / ATCC 23582 / Paris) OX=456481 GN=rpmE PE=3 SV=1 |
| Uncharacterized protein OS=Leptospira biflexa serovar Patoc (strain Patoc 1 / ATCC 23582 / Paris) OX=456481 GN=LEPBI_I2761 PE=4 SV=1 |
| Uncharacterized protein OS=Leptospira biflexa serovar Patoc (strain Patoc 1 / ATCC 23582 / Paris) OX=456481 GN=LEPBI_I0867 PE=4 SV=1 |
| Uncharacterized protein OS=Leptospira biflexa serovar Patoc (strain Patoc 1 / ATCC 23582 / Paris) OX=456481 GN=LEPBI_I2385 PE=4 SV=1 |
| Enoyl-[acyl-carrier-protein] reductase [NADH] OS=Leptospira biflexa serovar Patoc (strain Patoc 1 / ATCC 23582 / Paris) OX=456481 GN=LEPBI_I1195 PE=3 SV=1 |
| Cell shape protein MreC OS=Leptospira biflexa serovar Patoc (strain Patoc 1 / ATCC 23582 / Paris) OX=456481 GN=mreC PE=3 SV=1 |
| Uncharacterized protein OS=Leptospira biflexa serovar Patoc (strain Patoc 1 / ATCC 23582 / Paris) OX=456481 GN=LEPBI_I2588 PE=4 SV=1 |
| Putative RNA pseudouridine synthase (RNA-uridine isomerase RNA pseudouridylate synthase) OS=Leptospira biflexa serovar Patoc (strain Patoc 1 / ATCC 23582 / Paris) OX=456481 GN=LEPBI_I2113 PE=4 SV=1 |
| Putative esterase putative signal peptide OS=Leptospira biflexa serovar Patoc (strain Patoc 1 / ATCC 23582 / Paris) OX=456481 GN=LEPBI_I1504 PE=4 SV=1 |
| Ankyrin_rpt-contain_dom domain-containing protein OS=Leptospira biflexa serovar Patoc (strain Patoc 1 / ATCC 23582 / Paris) OX=456481 GN=LEPBI_II0221 PE=4 SV=1 |
| Putative transcriptional regulator OS=Leptospira biflexa serovar Patoc (strain Patoc 1 / ATCC 23582 / Paris) OX=456481 GN=LEPBI_I1031 PE=4 SV=1 |

| Description |
| --- |
| Succinate dehydrogenase flavoprotein subunit SdhA (Fumarate reductase) OS=Leptospira biflexa serovar Patoc (strain Patoc 1 / ATCC 23582 / Paris) OX=456481 GN=sdhA1 PE=4 SV=1 |
| TonB-dependent receptor protein putative signal peptide OS=Leptospira biflexa serovar Patoc (strain Patoc 1 / ATCC 23582 / Paris) OX=456481 GN=LEPBI_I3432 PE=3 SV=1 |
| Putative TonB-dependent outer membrane receptor putative signal peptide OS=Leptospira biflexa serovar Patoc (strain Patoc 1 / ATCC 23582 / Paris) OX=456481 GN=LEPBI_I0500 PE=4 SV=1 |
| Plug domain-containing protein OS=Leptospira biflexa serovar Patoc (strain Patoc 1 / ATCC 23582 / Paris) OX=456481 GN=LEPBI_I3354 PE=4 SV=1 |
| Uncharacterized protein OS=Leptospira biflexa serovar Patoc (strain Patoc 1 / ATCC 23582 / Paris) OX=456481 GN=LEPBI_I0608 PE=4 SV=1 |
| Uncharacterized protein OS=Leptospira biflexa serovar Patoc (strain Patoc 1 / ATCC 23582 / Paris) OX=456481 GN=LEPBI_I1431 PE=4 SV=1 |
| Putative outer membrane efflux protein putative signal peptide OS=Leptospira biflexa serovar Patoc (strain Patoc 1 / ATCC 23582 / Paris) OX=456481 GN=LEPBI_I1784 PE=3 SV=1 |
| Uncharacterized protein OS=Leptospira biflexa serovar Patoc (strain Patoc 1 / ATCC 23582 / Paris) OX=456481 GN=LEPBI_I2867 PE=4 SV=1 |
| Putative metal-dependent phosphohydrolase, HD subdomain putative membrane protein OS=Leptospira biflexa serovar Patoc (strain Patoc 1 / ATCC 23582 / Paris) OX=456481 GN=LEPBI_I2093 PE=4 SV=1 |
| Putative alkaline phosphatase putative membrane protein OS=Leptospira biflexa serovar Patoc (strain Patoc 1 / ATCC 23582 / Paris) OX=456481 GN=LEPBI_I1362 PE=4 SV=1 |
| Penicillin-binding protein 2 OS=Leptospira biflexa serovar Patoc (strain Patoc 1 / ATCC 23582 / Paris) OX=456481 GN=mrdA PE=4 SV=1 |
| Aspartyl/glutamyl-tRNA(Asn/Gln) amidotransferase subunit B OS=Leptospira biflexa serovar Patoc (strain Patoc 1 / ATCC 23582 / Paris) OX=456481 GN=gatB PE=3 SV=1 |
| DUF4139 domain-containing protein OS=Leptospira biflexa serovar Patoc (strain Patoc 1 / ATCC 23582 / Paris) OX=456481 GN=LEPBI_I0070 PE=4 SV=1 |
| Alginate_exp domain-containing protein OS=Leptospira biflexa serovar Patoc (strain Patoc 1 / ATCC 23582 / Paris) OX=456481 GN=LEPBI_I2473 PE=4 SV=1 |

| |
|---|
| Putative cAMP-binding protein OS=Leptospira biflexa serovar Patoc (strain Patoc 1 / ATCC 23582 / Paris) OX=456481 GN=LEPBI_I2338 PE=4 SV=1 |
| FlgO domain-containing protein OS=Leptospira biflexa serovar Patoc (strain Patoc 1 / ATCC 23582 / Paris) OX=456481 GN=LEPBI_II0250 PE=4 SV=1 |
| Putative adenylate/guanylate cyclase putative membrane protein OS=Leptospira biflexa serovar Patoc (strain Patoc 1 / ATCC 23582 / Paris) OX=456481 GN=LEPBI_I2606 PE=4 SV=1 |
| MaoC_dehydrat_N domain-containing protein OS=Leptospira biflexa serovar Patoc (strain Patoc 1 / ATCC 23582 / Paris) OX=456481 GN=LEPBI_I3165 PE=4 SV=1 |
| Acetyl-CoA carboxylase alpha subunit OS=Leptospira biflexa serovar Patoc (strain Patoc 1 / ATCC 23582 / Paris) OX=456481 GN=accA PE=4 SV=1 |
| Dihydrolipoyl dehydrogenase OS=Leptospira biflexa serovar Patoc (strain Patoc 1 / ATCC 23582 / Paris) OX=456481 GN=lpdA1 PE=3 SV=1 |
| Putative K(+)-stimulated pyrophosphate-energized sodium pump OS=Leptospira biflexa serovar Patoc (strain Patoc 1 / ATCC 23582 / Paris) OX=456481 GN=hppA PE=3 SV=1 |
| Flagellar M-ring protein OS=Leptospira biflexa serovar Patoc (strain Patoc 1 / ATCC 23582 / Paris) OX=456481 GN=fliF PE=3 SV=1 |
| Phosphoenolpyruvate carboxykinase (ATP) OS=Leptospira biflexa serovar Patoc (strain Patoc 1 / ATCC 23582 / Paris) OX=456481 GN=pckA PE=3 SV=1 |
| Uncharacterized protein OS=Leptospira biflexa serovar Patoc (strain Patoc 1 / ATCC 23582 / Paris) OX=456481 GN=LEPBI_II0278 PE=4 SV=1 |
| DUF4842 domain-containing protein OS=Leptospira biflexa serovar Patoc (strain Patoc 1 / ATCC 23582 / Paris) OX=456481 GN=LEPBI_II0156 PE=4 SV=1 |
| Cell division protein FtsA OS=Leptospira biflexa serovar Patoc (strain Patoc 1 / ATCC 23582 / Paris) OX=456481 GN=ftsA2 PE=4 SV=1 |
| Uncharacterized protein OS=Leptospira biflexa serovar Patoc (strain Patoc 1 / ATCC 23582 / Paris) OX=456481 GN=LEPBI_p0024 PE=4 SV=1 |
| Uncharacterized protein OS=Leptospira biflexa serovar Patoc (strain Patoc 1 / ATCC 23582 / Paris) OX=456481 GN=LEPBI_I0389 PE=4 SV=1 |
| Biopolymer transport ExbD protein OS=Leptospira biflexa serovar Patoc (strain Patoc 1 / ATCC 23582 / Paris) OX=456481 GN=exbD4 PE=3 SV=1 |

| |
|---|
| ATP synthase subunit delta OS=Leptospira biflexa serovar Patoc (strain Patoc 1 / ATCC 23582 / Paris) OX=456481 GN=atpH PE=3 SV=1 |
| Cytochrome c oxidase polypeptide III (Cytochrome aa3 subunit 3 Oxidase aa(3) subunit 3) OS=Leptospira biflexa serovar Patoc (strain Patoc 1 / ATCC [illegible] |
| Uncharacterized protein OS=Leptospira biflexa serovar Patoc (strain Patoc 1 / ATCC 23582 / Paris) OX=456481 GN=LEPBI_I0856 PE=3 SV=1 |
| Putative methyl-accepting chemotaxis protein OS=Leptospira biflexa serovar Patoc (strain Patoc 1 / ATCC 23582 / Paris) OX=456481 GN=LEPBI_I1582 PE=4 SV=1 |
| Uncharacterized protein OS=Leptospira biflexa serovar Patoc (strain Patoc 1 / ATCC 23582 / Paris) OX=456481 GN=LEPBI_I0621 PE=4 SV=1 |
| Putative SET domain-containing protein OS=Leptospira biflexa serovar Patoc (strain Patoc 1 / ATCC 23582 / Paris) OX=456481 GN=LEPBI_I2142 PE=4 SV=1 |
| 50S ribosomal protein L23 OS=Leptospira biflexa serovar Patoc (strain Patoc 1 / ATCC 23582 / Paris) OX=456481 GN=rplW PE=3 SV=1 |
| Uncharacterized protein OS=Leptospira biflexa serovar Patoc (strain Patoc 1 / ATCC 23582 / Paris) OX=456481 GN=LEPBI_I1694 PE=4 SV=1 |
| 30S ribosomal protein S15 OS=Leptospira biflexa serovar Patoc (strain Patoc 1 / ATCC 23582 / Paris) OX=456481 GN=rpsO PE=3 SV=1 |
| Putative transcriptional regulator, ArsR family OS=Leptospira biflexa serovar Patoc (strain Patoc 1 / ATCC 23582 / Paris) OX=456481 GN=LEPBI_I1981 PE=4 SV=1 |
| Putative cytochrome oxidase subunit II OS=Leptospira biflexa serovar Patoc (strain Patoc 1 / ATCC 23582 / Paris) OX=456481 GN=LEPBI_II0063 PE=4 SV=1 |
| NADH-quinone oxidoreductase chain G OS=Leptospira biflexa serovar Patoc (strain Patoc 1 / ATCC 23582 / Paris) OX=456481 GN=nuoG PE=4 SV=1 |
| Putative aminotransferase, DegT/DnrJ/EryC1/StrS family OS=Leptospira biflexa serovar Patoc (strain Patoc 1 / ATCC 23582 / Paris) OX=456481 GN=LEPBI_I0696 PE=3 SV=1 |
| UDP-N-acetylmuramoyl-tripeptide--D-alanyl-D-alanine ligase OS=Leptospira biflexa serovar Patoc (strain Patoc 1 / ATCC 23582 / Paris) OX=456481 GN=murF PE=3 SV=1 |
| NADH-quinone oxidoreductase, chain M OS=Leptospira biflexa serovar Patoc (strain Patoc 1 / ATCC 23582 / Paris) OX=456481 GN=nuoM PE=3 SV=1 |
| ABC-type transport system, ATP binding protein OS=Leptospira biflexa serovar Patoc (strain Patoc 1 / ATCC 23582 / Paris) OX=456481 GN=LEPBI_I2017 PE=3 SV=1 |

| |
|---|
| Putative lipoprotein OS=Leptospira biflexa serovar Patoc (strain Patoc 1 / ATCC 23582 / Paris) OX=456481 GN=LEPBI_I0560 PE=4 SV=1 |
| NADH:ubiquinone reductase (H(+)-translocating) OS=Leptospira biflexa serovar Patoc (strain Patoc 1 / ATCC 23582 / Paris) OX=456481 GN=nuoL PE=3 SV=1 |
| Uncharacterized protein OS=Leptospira biflexa serovar Patoc (strain Patoc 1 / ATCC 23582 / Paris) OX=456481 GN=LEPBI_I2758 PE=4 SV=1 |
| 50S ribosomal protein L28 OS=Leptospira biflexa serovar Patoc (strain Patoc 1 / ATCC 23582 / Paris) OX=456481 GN=rpmB PE=3 SV=1 |
| Anti-sigma factor antagonist OS=Leptospira biflexa serovar Patoc (strain Patoc 1 / ATCC 23582 / Paris) OX=456481 GN=LEPBI_I2328 PE=3 SV=1 |
| Putative membrane bound 0-acyl transferase, putative alginate biosynthesis protein putative membrane protein OS=Leptospira biflexa serovar Patoc (strain Patoc 1 / ATCC 23582 / Paris) OX=456481 GN=LEPBI_I3096 PE=3 SV=1 |
| Uncharacterized protein OS=Leptospira biflexa serovar Patoc (strain Patoc 1 / ATCC 23582 / Paris) OX=456481 GN=LEPBI_I2462 PE=4 SV=1 |
| DUF374 domain-containing protein OS=Leptospira biflexa serovar Patoc (strain Patoc 1 / ATCC 23582 / Paris) OX=456481 GN=LEPBI_I2969 PE=4 SV=1 |
| Putative permease of the major facilitator superfamily putative signal peptide OS=Leptospira biflexa serovar Patoc (strain Patoc 1 / ATCC 23582 / Paris) OX=456481 GN=LEPBI_I0303 PE=4 SV=1 |
| Uncharacterized protein OS=Leptospira biflexa serovar Patoc (strain Patoc 1 / ATCC 23582 / Paris) OX=456481 GN=LEPBI_I1684 PE=4 SV=1 |
| 50S ribosomal protein L30 OS=Leptospira biflexa serovar Patoc (strain Patoc 1 / ATCC 23582 / Paris) OX=456481 GN=rpmD PE=3 SV=1 |
| Uncharacterized protein OS=Leptospira biflexa serovar Patoc (strain Patoc 1 / ATCC 23582 / Paris) OX=456481 GN=LEPBI_I0362 PE=4 SV=1 |
| Uncharacterized protein OS=Leptospira biflexa serovar Patoc (strain Patoc 1 / ATCC 23582 / Paris) OX=456481 GN=LEPBI_II0027 PE=4 SV=1 |
| Dihydroxy-acid dehydratase OS=Leptospira biflexa serovar Patoc (strain Patoc 1 / ATCC 23582 / Paris) OX=456481 GN=ilvD PE=3 SV=1 |
| DNA gyrase subunit A OS=Leptospira biflexa serovar Patoc (strain Patoc 1 / ATCC 23582 / Paris) OX=456481 GN=gyrA1 PE=3 SV=1 |

| |
|---|
| Transport permease protein OS=Leptospira biflexa serovar Patoc (strain Patoc 1 / ATCC 23582 / Paris) OX=456481 GN=LEPBI_I1906 PE=3 SV=1 |
| Heme exporter protein C OS=Leptospira biflexa serovar Patoc (strain Patoc 1 / ATCC 23582 / Paris) OX=456481 GN=ccmC PE=3 SV=1 |
| Uncharacterized protein OS=Leptospira biflexa serovar Patoc (strain Patoc 1 / ATCC 23582 / Paris) OX=456481 GN=LEPBI_I3173 PE=4 SV=1 |
| Putative membrane protein, resistance to tellurium, TerC family putative membrane protein OS=Leptospira biflexa serovar Patoc (strain Patoc 1 / ATCC 23582 / Paris) OX=456481 GN=LEPBI_I3152 PE=4 SV=1 |
| Putative transcriptional regulator OS=Leptospira biflexa serovar Patoc (strain Patoc 1 / ATCC 23582 / Paris) OX=456481 GN=LEPBI_p0007 PE=4 SV=1 |
| Uncharacterized protein OS=Leptospira biflexa serovar Patoc (strain Patoc 1 / ATCC 23582 / Paris) OX=456481 GN=LEPBI_I2709 PE=4 SV=1 |
| Flagellar hook-associated protein 2 OS=Leptospira biflexa serovar Patoc (strain Patoc 1 / ATCC 23582 / Paris) OX=456481 GN=fliD PE=3 SV=1 |
| Biotin carboxylase (A subunit of acetyl-CoA carboxylase ACC) OS=Leptospira biflexa serovar Patoc (strain Patoc 1 / ATCC 23582 / Paris) OX=456481 GN=fabG7 PE=4 SV=1 |
| Enolase OS=Leptospira biflexa serovar Patoc (strain Patoc 1 / ATCC 23582 / Paris) OX=456481 GN=eno PE=3 SV=1 |
| Vitamin B12-dependent ribonucleotide reductase OS=Leptospira biflexa serovar Patoc (strain Patoc 1 / ATCC 23582 / Paris) OX=456481 GN=nrdA PE=3 SV=1 |
| 50S ribosomal protein L35 OS=Leptospira biflexa serovar Patoc (strain Patoc 1 / ATCC 23582 / Paris) OX=456481 GN=rpmI PE=3 SV=1 |
| Protein translocase subunit SecA OS=Leptospira biflexa serovar Patoc (strain Patoc 1 / ATCC 23582 / Paris) OX=456481 GN=secA PE=3 SV=1 |
| Transcription termination/antitermination protein NusG OS=Leptospira biflexa serovar Patoc (strain Patoc 1 / ATCC 23582 / Paris) OX=456481 GN=nusG PE=3 SV=1 |
| Peptidase_M23 domain-containing protein OS=Leptospira biflexa serovar Patoc (strain Patoc 1 / ATCC 23582 / Paris) OX=456481 GN=LEPBI_I0051 PE=4 SV=1 |
| Uncharacterized protein OS=Leptospira biflexa serovar Patoc (strain Patoc 1 / ATCC 23582 / Paris) OX=456481 GN=LEPBI_II0262 PE=4 SV=1 |
| Aspartokinase OS=Leptospira biflexa serovar Patoc (strain Patoc 1 / ATCC 23582 / Paris) OX=456481 GN=lysC PE=3 SV=1 |

| |
|---|
| Sigma-54 factor interaction domain-containing protein OS=Leptospira biflexa serovar Patoc (strain Patoc 1 / ATCC 23582 / Paris) OX=456481 GN=LEPBI_I0254 PE=4 SV=1 |
| Uncharacterized protein OS=Leptospira biflexa serovar Patoc (strain Patoc 1 / ATCC 23582 / Paris) OX=456481 GN=LEPBI_I0288 PE=4 SV=1 |
| Cation efflux system protein, acrB/acrD/acrF family putative membrane protein OS=Leptospira biflexa serovar Patoc (strain Patoc 1 / ATCC 23582 / Paris) OX=456481 GN=LEPBI_I3369 PE=3 SV=1 |
| Alpha-glucosidase OS=Leptospira biflexa serovar Patoc (strain Patoc 1 / ATCC 23582 / Paris) OX=456481 GN=LEPBI_I0772 PE=4 SV=1 |
| Glutathione-regulated potassium-efflux system protein KefB putative membrane protein OS=Leptospira biflexa serovar Patoc (strain Patoc 1 / ATCC 23582 / Paris) OX=456481 GN=kefB PE=4 SV=1 |
| Putative ABC-type transport system, permease, iron-regulated OS=Leptospira biflexa serovar Patoc (strain Patoc 1 / ATCC 23582 / Paris) OX=456481 GN=LEPBI_I1359 PE=3 SV=1 |
| Uncharacterized protein OS=Leptospira biflexa serovar Patoc (strain Patoc 1 / ATCC 23582 / Paris) OX=456481 GN=LEPBI_I1494 PE=4 SV=1 |
| Uncharacterized protein OS=Leptospira biflexa serovar Patoc (strain Patoc 1 / ATCC 23582 / Paris) OX=456481 GN=LEPBI_I3139 PE=4 SV=1 |
| Signal peptidase I OS=Leptospira biflexa serovar Patoc (strain Patoc 1 / ATCC 23582 / Paris) OX=456481 GN=LEPBI_I2078 PE=3 SV=1 |
| NADH-quinone oxidoreductase subunit I OS=Leptospira biflexa serovar Patoc (strain Patoc 1 / ATCC 23582 / Paris) OX=456481 GN=nuoI PE=3 SV=1 |
| Uncharacterized protein OS=Leptospira biflexa serovar Patoc (strain Patoc 1 / ATCC 23582 / Paris) OX=456481 GN=LEPBI_p0045 PE=4 SV=1 |
| Serine hydroxymethyltransferase OS=Leptospira biflexa serovar Patoc (strain Patoc 1 / ATCC 23582 / Paris) OX=456481 GN=glyA PE=3 SV=1 |
| Isoleucine--tRNA ligase OS=Leptospira biflexa serovar Patoc (strain Patoc 1 / ATCC 23582 / Paris) OX=456481 GN=ileS PE=3 SV=1 |
| Uncharacterized protein OS=Leptospira biflexa serovar Patoc (strain Patoc 1 / ATCC 23582 / Paris) OX=456481 GN=LEPBI_I3283 PE=4 SV=1 |
| Putative alkyldihydroxyacetonephosphate synthase OS=Leptospira biflexa serovar Patoc (strain Patoc 1 / ATCC 23582 / Paris) OX=456481 GN=eapA PE=4 SV=1 |

| |
|---|
| Putative sodium/galactoside symporter OS=Leptospira biflexa serovar Patoc (strain Patoc 1 / ATCC 23582 / Paris) OX=456481 GN=LEPBI_I0829 PE=4 SV=1 |
| STAS domain-containing protein OS=Leptospira biflexa serovar Patoc (strain Patoc 1 / ATCC 23582 / Paris) OX=456481 GN=LEPBI_II0035 PE=4 SV=1 |
| Ribosomal protein S12 methylthiotransferase RimO OS=Leptospira biflexa serovar Patoc (strain Patoc 1 / ATCC 23582 / Paris) OX=456481 GN=rimO PE=3 SV=1 |
| Chaperone ClpB OS=Leptospira biflexa serovar Patoc (strain Patoc 1 / ATCC 23582 / Paris) OX=456481 GN=clpB PE=3 SV=1 |
| Putative polyol dehydrogenase OS=Leptospira biflexa serovar Patoc (strain Patoc 1 / ATCC 23582 / Paris) OX=456481 GN=LEPBI_I3289 PE=4 SV=1 |
| Uncharacterized protein OS=Leptospira biflexa serovar Patoc (strain Patoc 1 / ATCC 23582 / Paris) OX=456481 GN=LEPBI_I0307 PE=4 SV=1 |
| Zn_protease domain-containing protein OS=Leptospira biflexa serovar Patoc (strain Patoc 1 / ATCC 23582 / Paris) OX=456481 GN=LEPBI_I0650 PE=4 SV=1 |
| PilZ domain-containing protein OS=Leptospira biflexa serovar Patoc (strain Patoc 1 / ATCC 23582 / Paris) OX=456481 GN=LEPBI_I2723 PE=4 SV=1 |
| Uncharacterized protein OS=Leptospira biflexa serovar Patoc (strain Patoc 1 / ATCC 23582 / Paris) OX=456481 GN=LEPBI_I0840 PE=4 SV=1 |
| Putative cytochrome c oxidase, subunit III OS=Leptospira biflexa serovar Patoc (strain Patoc 1 / ATCC 23582 / Paris) OX=456481 GN=LEPBI_II0061 PE=4 SV=1 |
| Putative acetyl-CoA hydrolase/transferase OS=Leptospira biflexa serovar Patoc (strain Patoc 1 / ATCC 23582 / Paris) OX=456481 GN=LEPBI_I0851 PE=3 SV=1 |
| Flagellar hook protein FlgE OS=Leptospira biflexa serovar Patoc (strain Patoc 1 / ATCC 23582 / Paris) OX=456481 GN=flgE PE=3 SV=1 |
| Putative methyl-accepting chemotaxis protein putative membrane protein OS=Leptospira biflexa serovar Patoc (strain Patoc 1 / ATCC 23582 / Paris) OX=456481 GN=LEPBI_I0484 PE=4 SV=1 |
| Phosphoribosylformylglycinamidine synthase subunit PurQ OS=Leptospira biflexa serovar Patoc (strain Patoc 1 / ATCC 23582 / Paris) OX=456481 GN=purQ PE=3 SV=1 |
| Uncharacterized protein OS=Leptospira biflexa serovar Patoc (strain Patoc 1 / ATCC 23582 / Paris) OX=456481 GN=LEPBI_II0141 PE=4 SV=1 |
| Trigger factor OS=Leptospira biflexa serovar Patoc (strain Patoc 1 / ATCC 23582 / Paris) OX=456481 GN=tig PE=3 SV=1 |

| |
|---|
| tRNA uridine(34) hydroxylase OS=Leptospira biflexa serovar Patoc (strain Patoc 1 / ATCC 23582 / Paris) OX=456481 GN=trhO PE=3 SV=1 |
| Putative membrane-associated phospholipid phosphatase putative membrane protein OS=Leptospira biflexa serovar Patoc (strain Patoc 1 / ATCC 23582 / Paris) OX=456481 GN=LEPBI_I3235 PE=4 SV=1 |
| Putative ribosomal subunit pseudouridine synthase OS=Leptospira biflexa serovar Patoc (strain Patoc 1 / ATCC 23582 / Paris) OX=456481 GN=LEPBI_I1463 PE=4 SV=1 |
| 1-deoxy-D-xylulose-5-phosphate synthase OS=Leptospira biflexa serovar Patoc (strain Patoc 1 / ATCC 23582 / Paris) OX=456481 GN=dxs PE=3 SV=1 |
| Putative C-terminal processing peptidase OS=Leptospira biflexa serovar Patoc (strain Patoc 1 / ATCC 23582 / Paris) OX=456481 GN=LEPBI_II0268 PE=4 SV=1 |
| Alanine aminopeptidase OS=Leptospira biflexa serovar Patoc (strain Patoc 1 / ATCC 23582 / Paris) OX=456481 GN=pepN2 PE=3 SV=1 |
| Metallophos domain-containing protein OS=Leptospira biflexa serovar Patoc (strain Patoc 1 / ATCC 23582 / Paris) OX=456481 GN=LEPBI_I0246 PE=4 SV=1 |
| Uncharacterized protein OS=Leptospira biflexa serovar Patoc (strain Patoc 1 / ATCC 23582 / Paris) OX=456481 GN=LEPBI_I0260 PE=4 SV=1 |
| 30S ribosomal protein S6 OS=Leptospira biflexa serovar Patoc (strain Patoc 1 / ATCC 23582 / Paris) OX=456481 GN=rpsF PE=3 SV=1 |
| Putative hydrolase, alpha/beta superfamily OS=Leptospira biflexa serovar Patoc (strain Patoc 1 / ATCC 23582 / Paris) OX=456481 GN=LEPBI_I0622 PE=4 SV=1 |
| SPOR domain-containing protein OS=Leptospira biflexa serovar Patoc (strain Patoc 1 / ATCC 23582 / Paris) OX=456481 GN=LEPBI_I1449 PE=4 SV=1 |
| Putative hydrolase putative signal peptide OS=Leptospira biflexa serovar Patoc (strain Patoc 1 / ATCC 23582 / Paris) OX=456481 GN=LEPBI_I0831 PE=4 SV=1 |
| Putative proton glutamate symport protein putative membrane protein OS=Leptospira biflexa serovar Patoc (strain Patoc 1 / ATCC 23582 / Paris) OX=456481 GN=gltP PE=4 SV=1 |
| Putative chromosome partitioning protein ParB OS=Leptospira biflexa serovar Patoc (strain Patoc 1 / ATCC 23582 / Paris) OX=456481 GN=LEPBI_p0002 PE=3 SV=1 |
| Putative ATP-dependent helicase OS=Leptospira biflexa serovar Patoc (strain Patoc 1 / ATCC 23582 / Paris) OX=456481 GN=LEPBI_I1719 PE=4 SV=1 |

| |
|---|
| Adenylosuccinate synthetase OS=Leptospira biflexa serovar Patoc (strain Patoc 1 / ATCC 23582 / Paris) OX=456481 GN=purA PE=3 SV=1 |
| Putative two-component sensor protein OS=Leptospira biflexa serovar Patoc (strain Patoc 1 / ATCC 23582 / Paris) OX=456481 GN=LEPBI_I0566 PE=4 SV=1 |
| Uncharacterized protein OS=Leptospira biflexa serovar Patoc (strain Patoc 1 / ATCC 23582 / Paris) OX=456481 GN=LEPBI_I0607 PE=4 SV=1 |
| Putative ankyrin-like protein OS=Leptospira biflexa serovar Patoc (strain Patoc 1 / ATCC 23582 / Paris) OX=456481 GN=LEPBI_I3051 PE=4 SV=1 |
| Uncharacterized protein OS=Leptospira biflexa serovar Patoc (strain Patoc 1 / ATCC 23582 / Paris) OX=456481 GN=LEPBI_II0274 PE=4 SV=1 |
| GGDEF domain-containing protein OS=Leptospira biflexa serovar Patoc (strain Patoc 1 / ATCC 23582 / Paris) OX=456481 GN=LEPBI_I2876 PE=4 SV=1 |
| PTS system, IIA component OS=Leptospira biflexa serovar Patoc (strain Patoc 1 / ATCC 23582 / Paris) OX=456481 GN=ptsN PE=4 SV=1 |
| Uncharacterized protein OS=Leptospira biflexa serovar Patoc (strain Patoc 1 / ATCC 23582 / Paris) OX=456481 GN=LEPBI_I1808 PE=4 SV=1 |
| Putative transcriptional regulator, PadR family OS=Leptospira biflexa serovar Patoc (strain Patoc 1 / ATCC 23582 / Paris) OX=456481 GN=LEPBI_I0617 PE=4 SV=1 |
| Anti-sigma factor antagonist OS=Leptospira biflexa serovar Patoc (strain Patoc 1 / ATCC 23582 / Paris) OX=456481 GN=LEPBI_I3270 PE=3 SV=1 |
| Radical SAM core domain-containing protein OS=Leptospira biflexa serovar Patoc (strain Patoc 1 / ATCC 23582 / Paris) OX=456481 GN=LEPBI_I0185 PE=4 SV=1 |
| Penicillin-insensitive transglycosylase OS=Leptospira biflexa serovar Patoc (strain Patoc 1 / ATCC 23582 / Paris) OX=456481 GN=mrcA PE=4 SV=1 |
| Nitric oxide reductase, cytochrome c-containing subunit OS=Leptospira biflexa serovar Patoc (strain Patoc 1 / ATCC 23582 / Paris) OX=456481 GN=norC PE=4 SV=1 |
| Carbamoyl-phosphate synthase large chain OS=Leptospira biflexa serovar Patoc (strain Patoc 1 / ATCC 23582 / Paris) OX=456481 GN=carB PE=3 SV=1 |
| Uncharacterized protein OS=Leptospira biflexa serovar Patoc (strain Patoc 1 / ATCC 23582 / Paris) OX=456481 GN=LEPBI_I3118 PE=4 SV=1 |
| Ribosome-recycling factor OS=Leptospira biflexa serovar Patoc (strain Patoc 1 / ATCC 23582 / Paris) OX=456481 GN=frr PE=3 SV=1 |

| |
|---|
| Putative transport system kinase putative ArgK protein OS=Leptospira biflexa serovar Patoc (strain Patoc 1 / ATCC 23582 / Paris) OX=456481 GN=LEPBI_II0092 PE=3 SV=1 |
| Uncharacterized protein OS=Leptospira biflexa serovar Patoc (strain Patoc 1 / ATCC 23582 / Paris) OX=456481 GN=LEPBI_II0255 PE=4 SV=1 |
| Uncharacterized protein OS=Leptospira biflexa serovar Patoc (strain Patoc 1 / ATCC 23582 / Paris) OX=456481 GN=LEPBI_I3253 PE=4 SV=1 |
| Uncharacterized protein OS=Leptospira biflexa serovar Patoc (strain Patoc 1 / ATCC 23582 / Paris) OX=456481 GN=LEPBI_I1346 PE=4 SV=1 |
| ABC-type transport system, ATP binding protein Putative heme-transporting ATPase OS=Leptospira biflexa serovar Patoc (strain Patoc 1 / ATCC 23582 / Paris) OX=456481 GN=LEPBI_I0274 PE=4 SV=1 |
| DNA replication protein DnaC OS=Leptospira biflexa serovar Patoc (strain Patoc 1 / ATCC 23582 / Paris) OX=456481 GN=dnaC PE=4 SV=1 |
| Putative alginate O-acetyltransferase OS=Leptospira biflexa serovar Patoc (strain Patoc 1 / ATCC 23582 / Paris) OX=456481 GN=LEPBI_II0257 PE=3 SV=1 |
| Putative methyl-accepting chemotaxis protein (MCP) putative membrane protein OS=Leptospira biflexa serovar Patoc (strain Patoc 1 / ATCC 23582 / Paris) OX=456481 GN=LEPBI_I0422 PE=4 SV=1 |
| Putative permease putative transmembrane protein OS=Leptospira biflexa serovar Patoc (strain Patoc 1 / ATCC 23582 / Paris) OX=456481 GN=LEPBI_II0129 PE=4 SV=1 |
| Uncharacterized protein OS=Leptospira biflexa serovar Patoc (strain Patoc 1 / ATCC 23582 / Paris) OX=456481 GN=LEPBI_I0731 PE=4 SV=1 |

### Volcano plot

**Proteins enriched in *L. biflexa wt* vs *fcpA-* (|FC| ≥ 2, p-value ≤ 0.05)**

| Locus | Fold Change | pValue | Signal(Bottom) |
|---|---|---|---|
| tr\|B0SR03\|B0SR03_LEPBP | 51.94 | 3.94E-05 | 8.14E-04 |
| tr\|B0STJ8\|B0STJ8_LEPBP | 44.14 | 6.70E-03 | 2.45E-03 |
| tr\|B0SPP1\|B0SPP1_LEPBP | 39.42 | 1.05E-02 | 2.53E-04 |
| tr\|B0STF2\|B0STF2_LEPBP | 35.92 | 1.96E-05 | 8.79E-04 |
| **tr\|B0SQV9\|B0SQV9_LEPBP** | 18.29 | 9.82E-03 | 9.31E-05 |
| tr\|B0SSW5\|B0SSW5_LEPBP | 3.47 | 1.02E-02 | 3.00E-04 |
| tr\|B0SJG1\|B0SJG1_LEPBP | 2.81 | 1.85E-02 | 5.06E-04 |
| tr\|B0SR80\|B0SR80_LEPBP | 2.35 | 1.90E-02 | 4.54E-04 |
| tr\|B0SPD3\|B0SPD3_LEPBP | 2.29 | 1.49E-02 | 6.52E-04 |
| tr\|B0SJC6\|B0SJC6_LEPBP | 2.26 | 1.50E-03 | 1.85E-02 |
| tr\|B0SKP2\|B0SKP2_LEPBP | 2.17 | 4.84E-03 | 1.02E-03 |

| | | | |
|---|---|---|---|
| tr\|B0SU19\|B0SU19_LEPBP | 2.16 | 2.13E-02 | 3.16E-04 |
| sp\|B0SPA9\|GLPK_LEPBP | 2.00 | 7.74E-03 | 4.76E-04 |

**Proteins enriched in *L. biflexa fcpA-* vs *wt* (|FC| ≥ 2, p-value ≤ 0.05)**

| Locus | Fold Change | pValue | Signal(Bottom) |
|---|---|---|---|
| tr\|B0SJC1\|B0SJC1_LEPBP | -15.08 | 2.65E-02 | 2.56E-03 |
| tr\|B0SM25\|B0SM25_LEPBP | -11.58 | 6.30E-03 | 1.86E-03 |
| tr\|B0STL3\|B0STL3_LEPBP | -11.35 | 5.53E-03 | 4.08E-03 |
| tr\|B0SND9\|B0SND9_LEPBP | -11.27 | 4.70E-03 | 2.80E-03 |
| tr\|B0SNV4\|B0SNV4_LEPBP | -11.14 | 2.97E-03 | 6.00E-03 |
| tr\|B0SSS0\|B0SSS0_LEPBP | -11.13 | 2.82E-02 | 2.81E-03 |
| tr\|B0SPJ5\|B0SPJ5_LEPBP | -11.11 | 1.07E-03 | 3.01E-03 |
| tr\|B0SRK8\|B0SRK8_LEPBP | -10.63 | 2.41E-02 | 8.89E-04 |
| tr\|B0SL39\|B0SL39_LEPBP | -10.48 | 4.78E-03 | 1.68E-03 |
| tr\|B0SPR2\|B0SPR2_LEPBP | -9.48 | 7.91E-04 | 3.37E-03 |
| tr\|B0SPH7\|B0SPH7_LEPBP | -9.45 | 8.58E-03 | 7.99E-03 |

| | | | |
|---|---|---|---|
| tr\|B0SNV5\|B0SNV5_LEPBP | -8.89 | 1.04E-02 | 1.71E-03 |
| sp\|B0SL33\|ILVC_LEPBP | -8.46 | 1.19E-04 | 1.82E-03 |
| tr\|B0SLT4\|B0SLT4_LEPBP | -7.94 | 5.81E-04 | 2.22E-03 |
| tr\|B0SSF9\|B0SSF9_LEPBP | -7.86 | 2.23E-03 | 6.57E-04 |
| sp\|B0SRF1\|DNAK_LEPBP | -7.75 | 5.06E-04 | 3.21E-03 |
| tr\|B0SSX3\|B0SSX3_LEPBP | -7.42 | 6.76E-03 | 1.45E-03 |
| tr\|B0STU9\|B0STU9_LEPBP | -7.05 | 2.99E-03 | 7.67E-04 |
| tr\|B0SJ11\|B0SJ11_LEPBP | -6.71 | 3.55E-03 | 1.90E-03 |
| tr\|B0SKQ5\|B0SKQ5_LEPBP | -6.46 | 4.77E-03 | 1.95E-03 |
| tr\|B0SJE3\|B0SJE3_LEPBP | -6.37 | 2.31E-02 | 2.33E-03 |
| tr\|B0SKH1\|B0SKH1_LEPBP | -6.23 | 1.56E-02 | 1.23E-03 |
| tr\|B0SU98\|B0SU98_LEPBP | -6.16 | 2.93E-02 | 2.84E-03 |
| tr\|B0SK35\|B0SK35_LEPBP | -5.98 | 1.46E-03 | 1.21E-03 |
| sp\|B0SSH0\|RL16_LEPBP | -5.92 | 5.51E-04 | 2.29E-03 |
| tr\|B0SMV8\|B0SMV8_LEPBP | -5.81 | 2.39E-02 | 1.82E-03 |

| | | | |
|---|---|---|---|
| sp\|B0SLC6\|ATPA_LEPBP | -5.72 | 1.77E-02 | 6.93E-03 |
| tr\|B0SMQ9\|B0SMQ9_LEPBP | -5.67 | 1.57E-02 | 7.77E-04 |
| tr\|B0SLN9\|B0SLN9_LEPBP | -5.60 | 2.69E-02 | 3.46E-03 |
| tr\|B0SLE4\|B0SLE4_LEPBP | -5.46 | 4.42E-03 | 8.75E-04 |
| tr\|B0SKB5\|B0SKB5_LEPBP | -5.42 | 6.57E-03 | 9.14E-04 |
| tr\|B0STS0\|B0STS0_LEPBP | -5.32 | 2.05E-03 | 2.07E-03 |
| tr\|B0SLT5\|B0SLT5_LEPBP | -5.29 | 1.02E-03 | 1.17E-03 |
| tr\|B0SKP3\|B0SKP3_LEPBP | -5.16 | 2.23E-02 | 1.25E-03 |
| tr\|B0SKU8\|B0SKU8_LEPBP | -5.02 | 1.42E-02 | 8.15E-04 |
| tr\|B0SU22\|B0SU22_LEPBP | -5.00 | 1.55E-02 | 1.63E-03 |
| sp\|B0SLC4\|ATPF_LEPBP | -4.99 | 2.97E-04 | 1.63E-03 |
| sp\|B0SSH2\|RL22_LEPBP | -4.75 | 3.21E-02 | 1.12E-03 |
| tr\|B0SR81\|B0SR81_LEPBP | -4.64 | 1.13E-02 | 2.44E-03 |
| tr\|B0SJ48\|B0SJ48_LEPBP | -4.64 | 9.38E-03 | 9.18E-04 |
| tr\|B0SSG9\|B0SSG9_LEPBP | -4.63 | 2.82E-03 | 7.74E-04 |

| | | | |
|---|---|---|---|
| tr\|B0SMT5\|B0SMT5_LEPBP | -4.62 | 1.91E-02 | 7.82E-04 |
| tr\|B0SML9\|B0SML9_LEPBP | -4.57 | 1.20E-02 | 2.89E-03 |
| tr\|B0SS93\|B0SS93_LEPBP | -4.43 | 9.01E-03 | 3.39E-03 |
| sp\|B0SRX5\|ACSA_LEPBP | -4.34 | 2.99E-02 | 6.95E-04 |
| **tr\|B0STI0\|B0STI0_LEPBP** | -4.27 | 1.90E-02 | 2.27E-03 |
| sp\|B0SQH4\|IF2_LEPBP | -4.21 | 2.66E-03 | 2.98E-03 |
| tr\|B0SQF4\|B0SQF4_LEPBP | -4.17 | 1.18E-02 | 2.36E-03 |
| sp\|B0SMR8\|THIM_LEPBP | -4.17 | 3.25E-02 | 6.67E-04 |
| tr\|B0SP37\|B0SP37_LEPBP | -4.14 | 6.95E-03 | 1.64E-03 |
| tr\|B0SN64\|B0SN64_LEPBP | -4.13 | 3.67E-04 | 2.04E-03 |
| tr\|B0SPR0\|B0SPR0_LEPBP | -4.06 | 6.85E-04 | 1.15E-03 |
| tr\|B0SL22\|B0SL22_LEPBP | -4.05 | 2.99E-02 | 1.49E-03 |
| tr\|B0SP10\|B0SP10_LEPBP | -4.04 | 3.26E-02 | 8.70E-04 |
| tr\|B0SNK4\|B0SNK4_LEPBP | -4.04 | 2.57E-02 | 7.77E-04 |
| tr\|B0SQ39\|B0SQ39_LEPBP | -4.03 | 1.35E-02 | 2.42E-03 |
| tr\|B0SRG2\|B0SRG2_LEPBP | -4.02 | 2.34E-02 | 2.41E-03 |

| | | | |
|---|---|---|---|
| tr\|B0SP59\|B0SP59_LEPBP | -4.01 | 1.31E-02 | 1.25E-03 |
| tr\|B0SJH1\|B0SJH1_LEPBP | -3.98 | 4.83E-03 | 8.99E-04 |
| tr\|B0STZ7\|B0STZ7_LEPBP | -3.96 | 2.04E-03 | 1.87E-03 |
| tr\|B0SPJ6\|B0SPJ6_LEPBP | -3.92 | 3.38E-03 | 1.19E-03 |
| tr\|B0SU52\|B0SU52_LEPBP | -3.86 | 2.02E-02 | 8.32E-04 |
| tr\|B0SKU7\|B0SKU7_LEPBP | -3.83 | 1.75E-02 | 8.44E-04 |
| tr\|B0SR96\|B0SR96_LEPBP | -3.80 | 2.58E-02 | 9.61E-04 |
| tr\|B0SLS1\|B0SLS1_LEPBP | -3.76 | 2.68E-02 | 1.26E-03 |
| sp\|B0SSI3\|RPOC_LEPBP | -3.74 | 2.36E-04 | 6.28E-03 |
| tr\|B0SQ54\|B0SQ54_LEPBP | -3.70 | 1.08E-03 | 1.12E-03 |
| tr\|B0SK80\|B0SK80_LEPBP | -3.59 | 5.62E-03 | 1.69E-03 |
| tr\|B0SSD3\|B0SSD3_LEPBP | -3.56 | 7.67E-03 | 8.64E-04 |
| tr\|B0SSZ8\|B0SSZ8_LEPBP | -3.52 | 3.07E-03 | 1.41E-03 |
| tr\|B0STX1\|B0STX1_LEPBP | -3.51 | 2.07E-02 | 9.88E-04 |
| tr\|B0SNY1\|B0SNY1_LEPBP | -3.51 | 2.23E-03 | 8.73E-04 |

| | | | |
|---|---|---|---|
| tr\|B0SNN9\|B0SNN9_LEPBP | -3.44 | 1.90E-02 | 1.65E-03 |
| tr\|B0SR70\|B0SR70_LEPBP | -3.38 | 8.26E-03 | 2.49E-03 |
| tr\|B0SQL8\|B0SQL8_LEPBP | -3.37 | 1.77E-03 | 1.43E-03 |
| sp\|B0SSV9\|RL9_LEPBP | -3.27 | 3.29E-03 | 1.33E-03 |
| tr\|B0SS29\|B0SS29_LEPBP | -3.20 | 2.74E-02 | 8.56E-04 |
| tr\|B0SKW0\|B0SKW0_LEPBP | -3.19 | 1.90E-02 | 1.86E-03 |
| tr\|B0SR08\|B0SR08_LEPBP | -3.05 | 2.37E-02 | 1.01E-03 |
| tr\|B0SKT3\|B0SKT3_LEPBP | -2.95 | 1.74E-02 | 5.44E-03 |
| tr\|B0SP24\|B0SP24_LEPBP | -2.94 | 1.11E-02 | 2.60E-03 |
| tr\|B0SRD4\|B0SRD4_LEPBP | -2.93 | 1.03E-02 | 2.90E-03 |
| tr\|B0SPC3\|B0SPC3_LEPBP | -2.89 | 2.22E-02 | 7.88E-04 |
| tr\|B0SPR6\|B0SPR6_LEPBP | -2.73 | 2.49E-02 | 1.08E-03 |
| tr\|B0STM9\|B0STM9_LEPBP | -2.69 | 2.29E-02 | 1.70E-03 |
| tr\|B0SJ13\|B0SJ13_LEPBP | -2.65 | 6.75E-03 | 3.19E-03 |
| tr\|B0SPX8\|B0SPX8_LEPBP | -2.58 | 2.18E-03 | 7.97E-04 |
| tr\|B0SN98\|B0SN98_LEPBP | -2.50 | 8.53E-03 | 1.27E-03 |

| | | | |
|---|---|---|---|
| tr\|B0SQL6\|B0SQL6_LEPBP | -2.48 | 9.10E-03 | 2.02E-03 |
| tr\|B0SMN0\|B0SMN0_LEPBP | -2.46 | 2.62E-02 | 3.26E-03 |
| sp\|B0SKZ5\|RL20_LEPBP | -2.44 | 1.94E-02 | 1.73E-03 |
| sp\|B0SSG7\|RL14_LEPBP | -2.43 | 8.10E-03 | 1.96E-03 |
| tr\|B0SK90\|B0SK90_LEPBP | -2.42 | 2.96E-02 | 9.95E-04 |
| tr\|B0SRP4\|B0SRP4_LEPBP | -2.34 | 7.36E-03 | 1.00E-02 |
| tr\|B0SM86\|B0SM86_LEPBP | -2.33 | 4.97E-03 | 1.88E-03 |
| sp\|B0SSW0\|RS18_LEPBP | -2.33 | 4.93E-03 | 1.10E-03 |
| tr\|B0SPU4\|B0SPU4_LEPBP | -2.29 | 1.29E-02 | 3.16E-03 |
| tr\|B0SQD9\|B0SQD9_LEPBP | -2.28 | 9.26E-03 | 1.16E-03 |
| tr\|B0SJP5\|B0SJP5_LEPBP | -2.26 | 3.15E-02 | 3.52E-03 |
| tr\|B0SPB5\|B0SPB5_LEPBP | -2.25 | 4.15E-04 | 9.94E-04 |
| sp\|B0SSI4\|RPOB_LEPBP | -2.25 | 7.65E-03 | 4.22E-03 |
| tr\|B0SUA7\|B0SUA7_LEPBP | -2.25 | 3.07E-02 | 8.06E-04 |
| tr\|B0SMC6\|B0SMC6_LEPBP | -2.22 | 4.80E-04 | 7.97E-04 |

| | | | |
|---|---|---|---|
| tr\|B0STX2\|B0STX2_LEPBP | -2.21 | 2.56E-02 | 7.76E-04 |
| tr\|B0SL41\|B0SL41_LEPBP | -2.21 | 3.17E-02 | 1.18E-03 |
| tr\|B0SSB2\|B0SSB2_LEPBP | -2.19 | 2.29E-02 | 2.25E-03 |
| tr\|B0SQC8\|B0SQC8_LEPBP | -2.14 | 6.27E-03 | 6.52E-04 |
| tr\|B0SMX9\|B0SMX9_LEPBP | -2.09 | 1.89E-02 | 1.19E-03 |
| tr\|B0SPU5\|B0SPU5_LEPBP | -2.05 | 8.18E-03 | 1.65E-03 |
| sp\|B0SSI6\|RL10_LEPBP | -2.02 | 2.86E-02 | 8.83E-04 |

| Signal(Top) | Description |
|---|---|
| 4.23E-02 | Uncharacterized protein OS=Leptospira biflexa serovar Patoc (strain Patoc 1 / ATCC 23582 / Paris) OX=456481 GN=LEPBI_I1597 PE=1 SV=1 |
| 1.08E-01 | Uncharacterized protein OS=Leptospira biflexa serovar Patoc (strain Patoc 1 / ATCC 23582 / Paris) OX=456481 GN=LEPBI_I0267 PE=1 SV=1 |
| 9.98E-03 | Uncharacterized protein OS=Leptospira biflexa serovar Patoc (strain Patoc 1 / ATCC 23582 / Paris) OX=456481 GN=LEPBI_I3081 PE=4 SV=1 |
| 3.16E-02 | Uncharacterized protein OS=Leptospira biflexa serovar Patoc (strain Patoc 1 / ATCC 23582 / Paris) OX=456481 GN=LEPBI_I2297 PE=4 SV=1 |
| 1.70E-03 | Uncharacterized protein OS=Leptospira biflexa serovar Patoc (strain Patoc 1 / ATCC 23582 / Paris) OX=456481 GN=LEPBI_I3291 PE=4 SV=1 |
| 1.04E-03 | Uncharacterized protein OS=Leptospira biflexa serovar Patoc (strain Patoc 1 / ATCC 23582 / Paris) OX=456481 GN=LEPBI_I2103 PE=4 SV=1 |
| 1.42E-03 | Putative serine-type endopeptidase putative signal peptide OS=Leptospira biflexa serovar Patoc (strain Patoc 1 / ATCC 23582 / Paris) OX=456481 GN=LEPBI_I0523 PE=4 SV=1 |
| 1.07E-03 | General secretion pathway protein C putative signal peptide OS=Leptospira biflexa serovar Patoc (strain Patoc 1 / ATCC 23582 / Paris) OX=456481 GN=gspC PE=4 SV=1 |
| 1.49E-03 | Uncharacterized protein OS=Leptospira biflexa serovar Patoc (strain Patoc 1 / ATCC 23582 / Paris) OX=456481 GN=LEPBI_I1347 PE=4 SV=1 |
| 4.18E-02 | Uncharacterized protein OS=Leptospira biflexa serovar Patoc (strain Patoc 1 / ATCC 23582 / Paris) OX=456481 GN=LEPBI_I0551 PE=4 SV=1 |
| 2.21E-03 | Uncharacterized protein OS=Leptospira biflexa serovar Patoc (strain Patoc 1 / ATCC 23582 / Paris) OX=456481 GN=LEPBI_I0666 PE=4 SV=1 |

| | |
|---|---|
| 6.82E-04 | AB hydrolase-1 domain-containing protein OS=Leptospira biflexa serovar Patoc (strain Patoc 1 / ATCC 23582 / Paris) OX=456481 GN=LEPBI_II0168 PE=4 SV=1 |
| 9.53E-04 | Glycerol kinase OS=Leptospira biflexa serovar Patoc (strain Patoc 1 / ATCC 23582 / Paris) OX=456481 GN=glpK PE=3 SV=1 |

| Signal(Top) | Description |
|---|---|
| 1.70E-04 | Putative adenylate/guanylate cyclase, family 3 putative signal peptide OS=Leptospira biflexa serovar Patoc (strain Patoc 1 / ATCC 23582 / Paris) OX=456481 GN=LEPBI_I0546 PE=4 SV=1 |
| 1.60E-04 | ATPase ClpC, ATP-binding subunit OS=Leptospira biflexa serovar Patoc (strain Patoc 1 / ATCC 23582 / Paris) OX=456481 GN=clpC PE=4 SV=1 |
| 3.60E-04 | Heat shock protein HtpG OS=Leptospira biflexa serovar Patoc (strain Patoc 1 / ATCC 23582 / Paris) OX=456481 GN=htpG PE=3 SV=1 |
| 2.49E-04 | Uncharacterized protein OS=Leptospira biflexa serovar Patoc (strain Patoc 1 / ATCC 23582 / Paris) OX=456481 GN=LEPBI_I2835 PE=4 SV=1 |
| 5.38E-04 | TPR-repeat-containing protein putative signal peptide OS=Leptospira biflexa serovar Patoc (strain Patoc 1 / ATCC 23582 / Paris) OX=456481 GN=LEPBI_I2897 PE=4 SV=1 |
| 2.52E-04 | Putative outer membrane efflux protein putative signal peptide OS=Leptospira biflexa serovar Patoc (strain Patoc 1 / ATCC 23582 / Paris) OX=456481 GN=LEPBI_I2058 PE=3 SV=1 |
| 2.71E-04 | Alginate_exp domain-containing protein OS=Leptospira biflexa serovar Patoc (strain Patoc 1 / ATCC 23582 / Paris) OX=456481 GN=LEPBI_I3035 PE=4 SV=1 |
| 8.36E-05 | Succinate dehydrogenase cytochrome B subunit SdhC putative membrane protein OS=Leptospira biflexa serovar Patoc (strain Patoc 1 / ATCC 23582 / Paris) OX=456481 GN=sdhC PE=4 SV=1 |
| 1.60E-04 | ABC-type transport system, ATP binding protein OS=Leptospira biflexa serovar Patoc (strain Patoc 1 / ATCC 23582 / Paris) OX=456481 GN=LEPBI_I0714 PE=4 SV=1 |
| 3.56E-04 | Putative molybdopterin oxidoreductase, iron-sulfur binding subunit OS=Leptospira biflexa serovar Patoc (strain Patoc 1 / ATCC 23582 / Paris) OX=456481 GN=LEPBI_I1369 PE=4 SV=1 |
| 8.45E-04 | Plug domain-containing protein OS=Leptospira biflexa serovar Patoc (strain Patoc 1 / ATCC 23582 / Paris) OX=456481 GN=LEPBI_I3017 PE=4 SV=1 |

| | |
|---|---|
| 1.92E-04 | PEGA domain-containing protein OS=Leptospira biflexa serovar Patoc (strain Patoc 1 / ATCC 23582 / Paris) OX=456481 GN=LEPBI_I2898 PE=4 SV=1 |
| 2.16E-04 | Ketol-acid reductoisomerase (NADP(+)) OS=Leptospira biflexa serovar Patoc (strain Patoc 1 / ATCC 23582 / Paris) OX=456481 GN=ilvC PE=3 SV=1 |
| 2.79E-04 | Uncharacterized protein OS=Leptospira biflexa serovar Patoc (strain Patoc 1 / ATCC 23582 / Paris) OX=456481 GN=LEPBI_I0859 PE=4 SV=1 |
| 8.36E-05 | 50S ribosomal protein L15 OS=Leptospira biflexa serovar Patoc (strain Patoc 1 / ATCC 23582 / Paris) OX=456481 GN=rplO PE=3 SV=1 |
| 4.14E-04 | Chaperone protein DnaK OS=Leptospira biflexa serovar Patoc (strain Patoc 1 / ATCC 23582 / Paris) OX=456481 GN=dnaK PE=3 SV=1 |
| 1.96E-04 | Ribosome-binding ATPase YchF OS=Leptospira biflexa serovar Patoc (strain Patoc 1 / ATCC 23582 / Paris) OX=456481 GN=ychF PE=3 SV=1 |
| 1.09E-04 | Uncharacterized protein OS=Leptospira biflexa serovar Patoc (strain Patoc 1 / ATCC 23582 / Paris) OX=456481 GN=LEPBI_II0097 PE=3 SV=1 |
| 2.83E-04 | Acriflavin resistance protein F putative membrane protein OS=Leptospira biflexa serovar Patoc (strain Patoc 1 / ATCC 23582 / Paris) OX=456481 GN=acrF PE=4 SV=1 |
| 3.02E-04 | DNA topoisomerase (ATP-hydrolyzing) OS=Leptospira biflexa serovar Patoc (strain Patoc 1 / ATCC 23582 / Paris) OX=456481 GN=gyrB2 PE=4 SV=1 |
| 3.66E-04 | Uncharacterized protein OS=Leptospira biflexa serovar Patoc (strain Patoc 1 / ATCC 23582 / Paris) OX=456481 GN=LEPBI_I0318 PE=4 SV=1 |
| 1.98E-04 | Fatty acid synthase subunit beta OS=Leptospira biflexa serovar Patoc (strain Patoc 1 / ATCC 23582 / Paris) OX=456481 GN=fabI PE=4 SV=1 |
| 4.61E-04 | Aconitate hydratase A OS=Leptospira biflexa serovar Patoc (strain Patoc 1 / ATCC 23582 / Paris) OX=456481 GN=acnA PE=4 SV=1 |
| 2.03E-04 | DNA gyrase subunit B OS=Leptospira biflexa serovar Patoc (strain Patoc 1 / ATCC 23582 / Paris) OX=456481 GN=gyrB1 PE=3 SV=1 |
| 3.87E-04 | 50S ribosomal protein L16 OS=Leptospira biflexa serovar Patoc (strain Patoc 1 / ATCC 23582 / Paris) OX=456481 GN=rplP PE=3 SV=1 |
| 3.13E-04 | Uncharacterized protein OS=Leptospira biflexa serovar Patoc (strain Patoc 1 / ATCC 23582 / Paris) OX=456481 GN=LEPBI_I2754 PE=4 SV=1 |

| | |
|---|---|
| 1.21E-03 | ATP synthase subunit alpha OS=Leptospira biflexa serovar Patoc (strain Patoc 1 / ATCC 23582 / Paris) OX=456481 GN=atpA PE=3 SV=1 |
| 1.37E-04 | Uncharacterized protein OS=Leptospira biflexa serovar Patoc (strain Patoc 1 / ATCC 23582 / Paris) OX=456481 GN=LEPBI_I2705 PE=4 SV=1 |
| 6.18E-04 | DNA helicase OS=Leptospira biflexa serovar Patoc (strain Patoc 1 / ATCC 23582 / Paris) OX=456481 GN=LEPBI_I2544 PE=3 SV=1 |
| 1.60E-04 | PlsC domain-containing protein OS=Leptospira biflexa serovar Patoc (strain Patoc 1 / ATCC 23582 / Paris) OX=456481 GN=LEPBI_I2441 PE=4 SV=1 |
| 1.69E-04 | Putative bifunctional enzyme: Glycerol-3-phosphate dehydrogenase [NAD(P)+] (NAD(P)H-dependent glycerol-3-phosphate dehydrogenase)/ Glycerol-3-phosphate acyltransferase (GPAT) OS=Leptospira biflexa serovar Patoc (strain Patoc 1 / ATCC 23582 / Paris) OX=456481 GN=LEPBI_I0227 PE=3 SV=1 |
| 3.88E-04 | Glutamate synthase (NADPH) OS=Leptospira biflexa serovar Patoc (strain Patoc 1 / ATCC 23582 / Paris) OX=456481 GN=gltD PE=4 SV=1 |
| 2.21E-04 | Uncharacterized protein OS=Leptospira biflexa serovar Patoc (strain Patoc 1 / ATCC 23582 / Paris) OX=456481 GN=LEPBI_I0860 PE=4 SV=1 |
| 2.43E-04 | Uncharacterized protein OS=Leptospira biflexa serovar Patoc (strain Patoc 1 / ATCC 23582 / Paris) OX=456481 GN=LEPBI_I0667 PE=4 SV=1 |
| 1.62E-04 | Putative metallo-dependent phosphatase OS=Leptospira biflexa serovar Patoc (strain Patoc 1 / ATCC 23582 / Paris) OX=456481 GN=LEPBI_I2350 PE=4 SV=1 |
| 3.25E-04 | Sigma factor SigB regulation protein RsbU putative phosphoserine phosphatase OS=Leptospira biflexa serovar Patoc (strain Patoc 1 / ATCC 23582 / Paris) OX=456481 GN=LEPBI_II0171 PE=4 SV=1 |
| 3.26E-04 | ATP synthase subunit b OS=Leptospira biflexa serovar Patoc (strain Patoc 1 / ATCC 23582 / Paris) OX=456481 GN=atpF PE=3 SV=1 |
| 2.36E-04 | 50S ribosomal protein L22 OS=Leptospira biflexa serovar Patoc (strain Patoc 1 / ATCC 23582 / Paris) OX=456481 GN=rplV PE=3 SV=1 |
| 5.26E-04 | General secretory pathway protein D OS=Leptospira biflexa serovar Patoc (strain Patoc 1 / ATCC 23582 / Paris) OX=456481 GN=gspD PE=3 SV=1 |
| 1.98E-04 | ABC_transp_aux domain-containing protein OS=Leptospira biflexa serovar Patoc (strain Patoc 1 / ATCC 23582 / Paris) OX=456481 GN=LEPBI_I0506 PE=4 SV=1 |
| 1.67E-04 | 50S ribosomal protein L29 OS=Leptospira biflexa serovar Patoc (strain Patoc 1 / ATCC 23582 / Paris) OX=456481 GN=rpmC PE=3 SV=1 |

| | |
|---|---|
| 1.69E-04 | ABC-type transport system, ATP binding protein OS=Leptospira biflexa serovar Patoc (strain Patoc 1 / ATCC 23582 / Paris) OX=456481 GN=LEPBI_I2731 PE=4 SV=1 |
| 6.32E-04 | ABC-type transport system, ATP-binding component OS=Leptospira biflexa serovar Patoc (strain Patoc 1 / ATCC 23582 / Paris) OX=456481 GN=ybiT PE=4 SV=1 |
| 7.66E-04 | Long-chain acyl-CoA synthetase, AMP-forming OS=Leptospira biflexa serovar Patoc (strain Patoc 1 / ATCC 23582 / Paris) OX=456481 GN=LEPBI_I1879 PE=4 SV=1 |
| 1.60E-04 | Acetyl-coenzyme A synthetase OS=Leptospira biflexa serovar Patoc (strain Patoc 1 / ATCC 23582 / Paris) OX=456481 GN=acsA PE=3 SV=1 |
| 5.31E-04 | Putative ATPase OS=Leptospira biflexa serovar Patoc (strain Patoc 1 / ATCC 23582 / Paris) OX=456481 GN=LEPBI_I2327 PE=4 SV=1 |
| 7.08E-04 | Translation initiation factor IF-2 OS=Leptospira biflexa serovar Patoc (strain Patoc 1 / ATCC 23582 / Paris) OX=456481 GN=infB PE=3 SV=1 |
| 5.65E-04 | Uncharacterized protein OS=Leptospira biflexa serovar Patoc (strain Patoc 1 / ATCC 23582 / Paris) OX=456481 GN=LEPBI_I1505 PE=4 SV=1 |
| 1.60E-04 | Hydroxyethylthiazole kinase OS=Leptospira biflexa serovar Patoc (strain Patoc 1 / ATCC 23582 / Paris) OX=456481 GN=thiM PE=3 SV=1 |
| 3.97E-04 | Uncharacterized protein OS=Leptospira biflexa serovar Patoc (strain Patoc 1 / ATCC 23582 / Paris) OX=456481 GN=LEPBI_I2981 PE=4 SV=1 |
| 4.95E-04 | Electron transfer flavoprotein beta-subunit (Beta-ETF Electron transfer flavoprotein small subunit) (ETFSS) OS=Leptospira biflexa serovar Patoc (strain Patoc 1 / ATCC 23582 / Paris) OX=456481 GN=etfB PE=4 SV=1 |
| 2.82E-04 | Putative dolichyl-phosphate-mannose--protein mannosyltransferase protein, family 39 putative membrane protein OS=Leptospira biflexa serovar Patoc (strain Patoc 1 / ATCC 23582 / Paris) OX=456481 GN=LEPBI_I1367 PE=4 SV=1 |
| 3.67E-04 | Uncharacterized protein OS=Leptospira biflexa serovar Patoc (strain Patoc 1 / ATCC 23582 / Paris) OX=456481 GN=LEPBI_I2426 PE=4 SV=1 |
| 2.15E-04 | Nitric-oxide reductase subunit B (Nitric oxide reductase cytochrome b subunit NOR large subunit) putative membrane protein OS=Leptospira biflexa serovar Patoc (strain Patoc 1 / ATCC 23582 / Paris) OX=456481 GN=norB PE=3 SV=1 |
| 1.92E-04 | Delta-aminolevulinic acid dehydratase OS=Leptospira biflexa serovar Patoc (strain Patoc 1 / ATCC 23582 / Paris) OX=456481 GN=hemB PE=3 SV=1 |
| 6.02E-04 | Putative sodium-dependent phosphate transport protein OS=Leptospira biflexa serovar Patoc (strain Patoc 1 / ATCC 23582 / Paris) OX=456481 GN=LEPBI_I3125 PE=4 SV=1 |
| 5.99E-04 | Putative ABC-type transport sytem, outer membrane protein OS=Leptospira biflexa serovar Patoc (strain Patoc 1 / ATCC 23582 / Paris) OX=456481 GN=LEPBI_I3390 PE=3 SV=1 |

| | |
|---|---|
| 3.11E-04 | Inosine-5'-monophosphate dehydrogenase OS=Leptospira biflexa serovar Patoc (strain Patoc 1 / ATCC 23582 / Paris) OX=456481 GN=guaB PE=3 SV=1 |
| 2.26E-04 | Putative methyltransferase OS=Leptospira biflexa serovar Patoc (strain Patoc 1 / ATCC 23582 / Paris) OX=456481 GN=LEPBI_I0533 PE=4 SV=1 |
| 4.71E-04 | Putative peroxidase OS=Leptospira biflexa serovar Patoc (strain Patoc 1 / ATCC 23582 / Paris) OX=456481 GN=LEPBI_II0146 PE=3 SV=1 |
| 3.03E-04 | Putative transcriptional regulator, TetR family OS=Leptospira biflexa serovar Patoc (strain Patoc 1 / ATCC 23582 / Paris) OX=456481 GN=LEPBI_I3036 PE=4 SV=1 |
| 2.16E-04 | Uncharacterized protein OS=Leptospira biflexa serovar Patoc (strain Patoc 1 / ATCC 23582 / Paris) OX=456481 GN=LEPBI_II0203 PE=4 SV=1 |
| 2.21E-04 | Iron-sulfur cluster carrier protein OS=Leptospira biflexa serovar Patoc (strain Patoc 1 / ATCC 23582 / Paris) OX=456481 GN=mrp PE=3 SV=1 |
| 2.53E-04 | Putative arylesterase OS=Leptospira biflexa serovar Patoc (strain Patoc 1 / ATCC 23582 / Paris) OX=456481 GN=LEPBI_I3323 PE=4 SV=1 |
| 3.34E-04 | Lon protease OS=Leptospira biflexa serovar Patoc (strain Patoc 1 / ATCC 23582 / Paris) OX=456481 GN=lon PE=2 SV=1 |
| 1.68E-03 | DNA-directed RNA polymerase subunit beta' OS=Leptospira biflexa serovar Patoc (strain Patoc 1 / ATCC 23582 / Paris) OX=456481 GN=rpoC PE=3 SV=1 |
| 3.02E-04 | Putative Zinc-exporting ATPase (Zn(II)-translocating P-type ATPase) putative membrane protein OS=Leptospira biflexa serovar Patoc (strain Patoc 1 / ATCC 23582 / Paris) OX=456481 GN=ziaA PE=3 SV=1 |
| 4.71E-04 | Uncharacterized protein OS=Leptospira biflexa serovar Patoc (strain Patoc 1 / ATCC 23582 / Paris) OX=456481 GN=LEPBI_I0178 PE=4 SV=1 |
| 2.43E-04 | ABC-type transport system associated with Fe-S cluster assembly, ATPase OS=Leptospira biflexa serovar Patoc (strain Patoc 1 / ATCC 23582 / Paris) OX=456481 GN=sufC PE=3 SV=1 |
| 4.00E-04 | ATP-dependent RNA helicase OS=Leptospira biflexa serovar Patoc (strain Patoc 1 / ATCC 23582 / Paris) OX=456481 GN=LEPBI_I2136 PE=3 SV=1 |
| 2.81E-04 | Putative UTP--glucose-1-phosphate uridylyltransferase OS=Leptospira biflexa serovar Patoc (strain Patoc 1 / ATCC 23582 / Paris) OX=456481 GN=galU2 PE=4 SV=1 |
| 2.49E-04 | Uncharacterized protein OS=Leptospira biflexa serovar Patoc (strain Patoc 1 / ATCC 23582 / Paris) OX=456481 GN=LEPBI_I2924 PE=4 SV=1 |

| | |
|---|---|
| 4.79E-04 | Ankyrin_rpt-contain_dom domain-containing protein OS=Leptospira biflexa serovar Patoc (strain Patoc 1 / ATCC 23582 / Paris) OX=456481 GN=LEPBI_I1206 PE=4 SV=1 |
| 7.36E-04 | 50S ribosomal protein L19 OS=Leptospira biflexa serovar Patoc (strain Patoc 1 / ATCC 23582 / Paris) OX=456481 GN=rplS PE=3 SV=1 |
| 4.23E-04 | Aerobic glycerol-3-phosphate dehydrogenase OS=Leptospira biflexa serovar Patoc (strain Patoc 1 / ATCC 23582 / Paris) OX=456481 GN=glpD PE=3 SV=1 |
| 4.06E-04 | 50S ribosomal protein L9 OS=Leptospira biflexa serovar Patoc (strain Patoc 1 / ATCC 23582 / Paris) OX=456481 GN=rplI PE=3 SV=1 |
| 2.68E-04 | Uncharacterized protein OS=Leptospira biflexa serovar Patoc (strain Patoc 1 / ATCC 23582 / Paris) OX=456481 GN=LEPBI_I1813 PE=4 SV=1 |
| 5.82E-04 | Acetolactate synthase OS=Leptospira biflexa serovar Patoc (strain Patoc 1 / ATCC 23582 / Paris) OX=456481 GN=ilvB2 PE=3 SV=1 |
| 3.33E-04 | PHB domain-containing protein OS=Leptospira biflexa serovar Patoc (strain Patoc 1 / ATCC 23582 / Paris) OX=456481 GN=LEPBI_I1602 PE=3 SV=1 |
| 1.85E-03 | Transcription elongation factor GreA OS=Leptospira biflexa serovar Patoc (strain Patoc 1 / ATCC 23582 / Paris) OX=456481 GN=greA PE=3 SV=1 |
| 8.86E-04 | S-adenosylmethionine synthase OS=Leptospira biflexa serovar Patoc (strain Patoc 1 / ATCC 23582 / Paris) OX=456481 GN=metK PE=3 SV=1 |
| 9.90E-04 | TonB-dependent receptor protein putative signal peptide OS=Leptospira biflexa serovar Patoc (strain Patoc 1 / ATCC 23582 / Paris) OX=456481 GN=LEPBI_I3362 PE=3 SV=1 |
| 2.73E-04 | GGDEF domain-containing protein OS=Leptospira biflexa serovar Patoc (strain Patoc 1 / ATCC 23582 / Paris) OX=456481 GN=LEPBI_I1337 PE=4 SV=1 |
| 3.96E-04 | Uncharacterized protein OS=Leptospira biflexa serovar Patoc (strain Patoc 1 / ATCC 23582 / Paris) OX=456481 GN=LEPBI_I1373 PE=4 SV=1 |
| 6.32E-04 | Putative ATPase, ParA family OS=Leptospira biflexa serovar Patoc (strain Patoc 1 / ATCC 23582 / Paris) OX=456481 GN=LEPBI_II0025 PE=4 SV=1 |
| 1.20E-03 | Membrane protein insertase YidC OS=Leptospira biflexa serovar Patoc (strain Patoc 1 / ATCC 23582 / Paris) OX=456481 GN=yidC PE=3 SV=1 |
| 3.09E-04 | Uncharacterized protein OS=Leptospira biflexa serovar Patoc (strain Patoc 1 / ATCC 23582 / Paris) OX=456481 GN=LEPBI_I1436 PE=4 SV=1 |
| 5.09E-04 | Uncharacterized protein OS=Leptospira biflexa serovar Patoc (strain Patoc 1 / ATCC 23582 / Paris) OX=456481 GN=LEPBI_I2788 PE=4 SV=1 |

| | |
|---|---|
| 8.15E-04 | RNA polymerase sigma factor SigA OS=Leptospira biflexa serovar Patoc (strain Patoc 1 / ATCC 23582 / Paris) OX=456481 GN=rpoD PE=3 SV=1 |
| 1.32E-03 | Transcriptional regulator, CarD family OS=Leptospira biflexa serovar Patoc (strain Patoc 1 / ATCC 23582 / Paris) OX=456481 GN=LEPBI_I2676 PE=4 SV=1 |
| 7.11E-04 | 50S ribosomal protein L20 OS=Leptospira biflexa serovar Patoc (strain Patoc 1 / ATCC 23582 / Paris) OX=456481 GN=rplT PE=3 SV=1 |
| 8.06E-04 | 50S ribosomal protein L14 OS=Leptospira biflexa serovar Patoc (strain Patoc 1 / ATCC 23582 / Paris) OX=456481 GN=rplN PE=3 SV=1 |
| 4.11E-04 | NAD(P) transhydrogenase subunit beta OS=Leptospira biflexa serovar Patoc (strain Patoc 1 / ATCC 23582 / Paris) OX=456481 GN=pntB PE=3 SV=1 |
| 4.27E-03 | Putative cell envelope biogenesis protein putative signal peptide OS=Leptospira biflexa serovar Patoc (strain Patoc 1 / ATCC 23582 / Paris) OX=456481 GN=LEPBI_I1733 PE=4 SV=1 |
| 8.08E-04 | Putative strictosidine synthase putative signal peptide OS=Leptospira biflexa serovar Patoc (strain Patoc 1 / ATCC 23582 / Paris) OX=456481 GN=LEPBI_I2637 PE=4 SV=1 |
| 4.71E-04 | 30S ribosomal protein S18 OS=Leptospira biflexa serovar Patoc (strain Patoc 1 / ATCC 23582 / Paris) OX=456481 GN=rpsR PE=3 SV=1 |
| 1.38E-03 | Protein translocase subunit SecD OS=Leptospira biflexa serovar Patoc (strain Patoc 1 / ATCC 23582 / Paris) OX=456481 GN=secD PE=3 SV=1 |
| 5.10E-04 | Putative peptidase, S41 family putative signal peptide OS=Leptospira biflexa serovar Patoc (strain Patoc 1 / ATCC 23582 / Paris) OX=456481 GN=LEPBI_I1490 PE=4 SV=1 |
| 1.56E-03 | Uncharacterized protein OS=Leptospira biflexa serovar Patoc (strain Patoc 1 / ATCC 23582 / Paris) OX=456481 GN=LEPBI_I0061 PE=4 SV=1 |
| 4.41E-04 | Uncharacterized protein OS=Leptospira biflexa serovar Patoc (strain Patoc 1 / ATCC 23582 / Paris) OX=456481 GN=LEPBI_I1329 PE=4 SV=1 |
| 1.87E-03 | DNA-directed RNA polymerase subunit beta OS=Leptospira biflexa serovar Patoc (strain Patoc 1 / ATCC 23582 / Paris) OX=456481 GN=rpoB PE=3 SV=1 |
| 3.58E-04 | Uncharacterized protein OS=Leptospira biflexa serovar Patoc (strain Patoc 1 / ATCC 23582 / Paris) OX=456481 GN=LEPBI_II0258 PE=4 SV=1 |
| 3.60E-04 | RNA polymerase sigma factor OS=Leptospira biflexa serovar Patoc (strain Patoc 1 / ATCC 23582 / Paris) OX=456481 GN=fliA PE=3 SV=1 |

| | |
|---|---|
| 3.51E-04 | Putative UDP-glucose 4-epimerase OS=Leptospira biflexa serovar Patoc (strain Patoc 1 / ATCC 23582 / Paris) OX=456481 GN=LEPBI_II0120 PE=4 SV=1 |
| 5.33E-04 | Uncharacterized protein OS=Leptospira biflexa serovar Patoc (strain Patoc 1 / ATCC 23582 / Paris) OX=456481 GN=LEPBI_I0716 PE=4 SV=1 |
| 1.03E-03 | 2-oxoglutarate reductase OS=Leptospira biflexa serovar Patoc (strain Patoc 1 / ATCC 23582 / Paris) OX=456481 GN=serA2 PE=3 SV=1 |
| 3.04E-04 | Uncharacterized protein OS=Leptospira biflexa serovar Patoc (strain Patoc 1 / ATCC 23582 / Paris) OX=456481 GN=LEPBI_I1479 PE=3 SV=1 |
| 5.68E-04 | Citrate synthase OS=Leptospira biflexa serovar Patoc (strain Patoc 1 / ATCC 23582 / Paris) OX=456481 GN=gltA PE=3 SV=1 |
| 8.04E-04 | Protein-export membrane protein SecF OS=Leptospira biflexa serovar Patoc (strain Patoc 1 / ATCC 23582 / Paris) OX=456481 GN=secF PE=3 SV=1 |
| 4.38E-04 | 50S ribosomal protein L10 OS=Leptospira biflexa serovar Patoc (strain Patoc 1 / ATCC 23582 / Paris) OX=456481 GN=rplJ PE=3 SV=1 |

**Table S3.** Label-free comparative proteomics of *L. biflexa* filaments from *wt*

| **Samples of *L. biflexa* filaments from wt and sheath mutant s** |
|---|

| ***L. biflexa wt*** | **Total Proteins** |
|---|---|
| R1 | 749 |
| R2 | 727 |
| R3 | 609 |

| ***L. biflexa flaA2-*** |
|---|
| R1 |
| R2 |
| R4 |

vs mutant strains.

**strains, used for label-free comparative proteomics:**

| Total Proteins |
|---|
| 1382 |
| 1564 |
| 1170 |

| ***L. biflexa fcpA-*** | **Total Proteins** |
|---|---|
| R1 | 990 |
| R2 | 624 |
| R4 | 964 |
